# Emergent Metabolic Niches for Marine Heterotrophs

**DOI:** 10.1101/2024.05.29.596556

**Authors:** R Reynolds, ACB Weiss, CC James, CY Kojima, JL Weissman, JC Thrash, NM Levine

## Abstract

Ocean microbial communities are made up of thousands of diverse taxa whose metabolic demands set the rates of both biomass production and degradation. Thus, these microscopic organisms play a critical role in ecosystem dynamics, global carbon cycling, and climate. While we have frameworks for relating phytoplankton diversity to rates of carbon fixation, our knowledge of how variations in heterotrophic microbial populations drive changes in carbon cycling is in its infancy. Here, we leverage global metagenomic datasets and metabolic models to identify a set of metabolic niches with distinct growth strategies. These groupings provide a simplifying framework for describing microbial communities in different oceanographic regions and for understanding how heterotrophic microbial populations function. This framework, predicated directly on metabolic capability rather than taxonomy, enables us to tractably link heterotrophic diversity directly to biogeochemical rates in large scale ecosystem models.

---

Classification of heterotrophic microbes into metabolic functional guilds can provide a framework for coalescing diverse microbial communities^1^ into more tractable units for incorporation into biogeochemical models^2^. Historically, we have grouped marine microbial heterotrophs into copiotrophic organisms, which thrive in high resource environments and generally have faster growth rates with flexible metabolisms, and oligotrophic organisms, which dominate resource poor environments and have slower growth rates^3^. While these broad categories are useful, they do not inherently facilitate defining metabolic niches or substrate preferences that are critical when considering rates of biogeochemical cycling. Specifically, there is no intrinsic linkage between fast or slow maximum growth rates and the substrate preferences for organisms^4^. In this analysis, we expand beyond the copiotroph-oligotroph paradigm and independently assess metabolic strategy and growth rates to generate a generalizable functional categorization of marine microbial heterotrophic metabolisms.

Genome-scale metabolic models (GEMs) provide a means for translating genomic information into cellular metabolisms^5^ but have historically been labor intensive to generate and have been generally restricted to cultured isolates^6^. The advent of fast automated metabolic model construction software such as CarveMe, ModelSEED, Agora^7–9^, etc., has enabled generating GEMs for large numbers of genomes and from uncultured organisms^10^. Metabolic potential from GEMs can further be explored through the use of flux balance analysis tools such as CobraPy^11^. These combined analyses provide insights into the minimal metabolic requirements for a cell and hypotheses about the preferred substrates for growth^12^.

Here we leveraged a large global dataset of marine microbial genomes (Ocean Microbial Database (OMD))^13^ to identify patterns in metabolic strategies among marine bacteria through GEMs. Testing model sensitivity to growth on different substrates allowed us to define unique clusters of marine heterotrophic bacteria with shared growth strategies. We identified a classic fast-growing copiotrophic cluster, four slow-growing oligotrophic clusters each with a unique metabolic strategy, and three intermediate growth clusters, also with unique metabolic strategies. These clusters are found globally but at varying abundances in different ecological regimes. While clear phylogenetic signals emerged distinguishing the clusters, our findings also suggest that similar metabolic niches are occupied by distinct taxonomic groups.

## Results

### CarveMe model quality

We generated GEMs for 3,918 high quality bacterial genomes (including cultured isolates, metagenomes, and single-cell genomes) from OMD using CarveMe^7^. This dataset spanned a wide diversity of marine bacteria representing 205 distinct taxonomic orders, fifteen of which had fifty or more genomes (Figure 1). Given the stochastic nature of the cutting algorithm in CarveMe, it is necessary to run ensembles of models for each genome^7,14^. To ensure we included only high quality models in our analysis, we generated a large model ensemble for each genome (N = 60) and assessed the robustness of the models using a consensus metric. Specifically, higher confidence can be placed in models where a consistent set of enzymatic reactions are included across the entire ensemble of models generated by CarveMe for a single genome (high consensus value).

**Figure 1:**
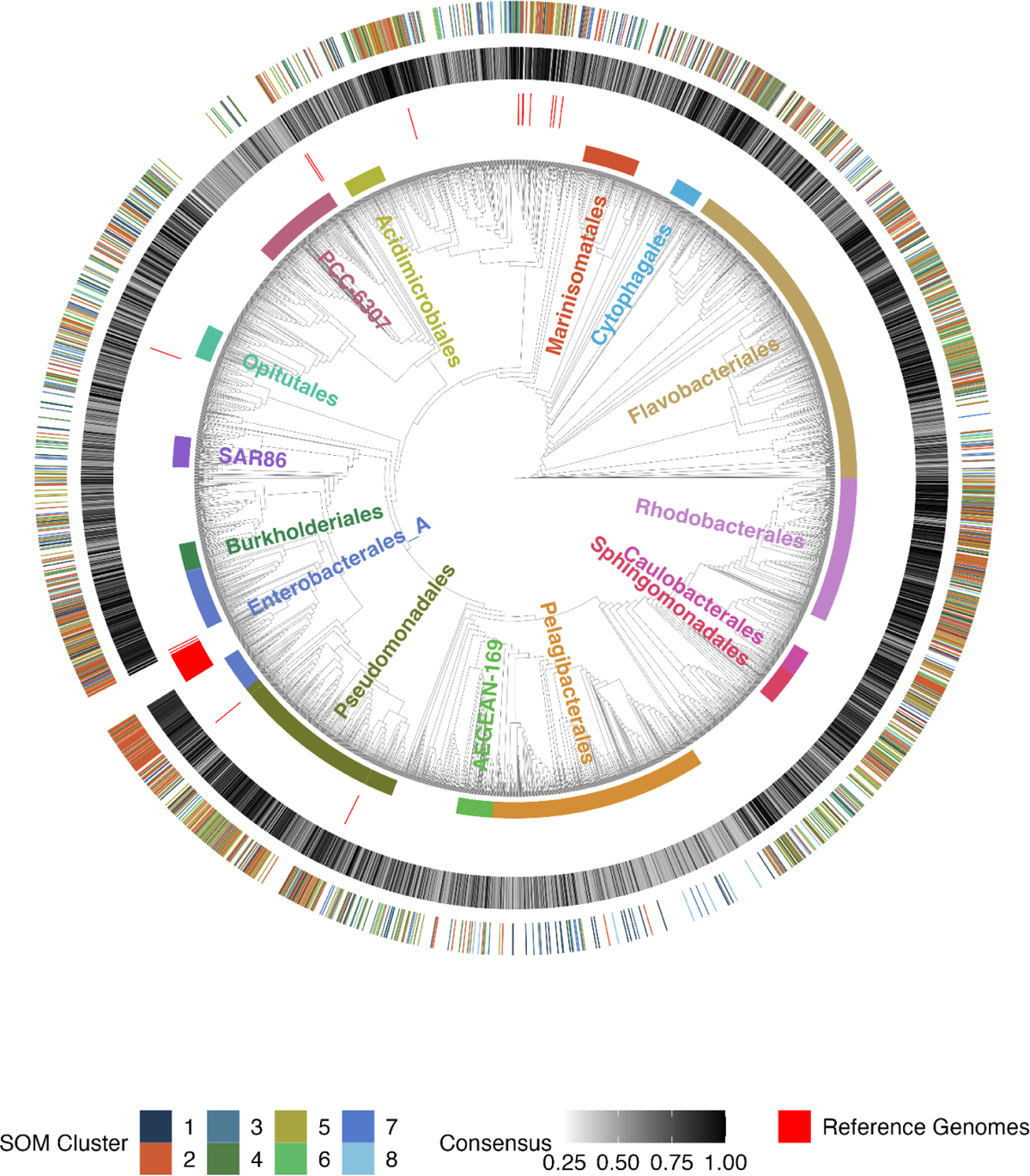
Diversity of dataset, quality of metabolic models, and designation of metabolic clusters. Phylogenetic tree of all 3,984 bacterial genomes included in this study (including the 66 reference genomes from the BiGG database). The tree is contextualized by several external rings that describe different qualitative and quantitative components of the genomes in this study. The first ring around the tree denotes both the position and density of high quality ensembles within the tree as well as the assignment of these genomes to each of our eight SOM clusters. The second ring shows the ensemble consensus score (Equation 1) for each genome in the tree. The third, sparse ring of red lines denotes the position of the 66 BiGG reference genomes present in the tree. Finally, the fourth and innermost ring shows the location of the top 15 most abundant orders.

Consensus values for the OMD genomes ranged from 1 (all model reactions were the same across all ensemble members) to 0.24 (only 24% of reactions were conserved between ensemble members thus providing low confidence in the CarveMe models). Systematic differences in consensus values were seen between phylogenetic groups (Figure 1, 3rd ring). The *Enterobacterales*, *Rhodobacterales*, *Cytophagales*, *Sphingomonadales*, and *Pseudomonadales* had the largest number of genomes with high consensus values: on average, 65.0% of genomes from each of these orders had consensus values above 0.8 (range 51.3%-81.3%). Genomes that were phylogenetically similar to the reference genomes used to develop CarveMe generally had higher consensus values (Figure 1). Several orders had a large proportion of genomes with high consensus values despite being phylogenetically distant from the reference genomes. The *Rhodobacterales*, for example, have no reference genome but 72.1% of these genomes had a consensus value greater than 0.8. CarveMe struggled to generate high consensus ensembles for several orders. Only 10.8% of *Pelagibacterales* and 4.1% of PCC-6307 genomes had consensus values above 0.8, while 66.5% and 49.7% of genomes in these groups had values at or below 0.5, respectively.

This analysis suggests that adding reference genomes (experimentally validated metabolic models used to improve the CarveMe tool) in these low consensus orders would substantially improve the ability for CarveMe to generate high consensus ensembles. This would greatly improve our ability to robustly apply CarveMe broadly to environmental datasets^15^. Our further analyses only used the 1,591 genomes with consensus values of 0.8 or greater. We tested a more conservative cutoff of 0.9, yielding a dataset of 983 genomes, and showed that the primary findings of this work remain unchanged (Supplemental Figure S1). The low number of PCC-6307 genomes with high quality models (N=7) was likely due to the fact that these are Cyanobacteria and so phototrophic or mixotrophic (capable of growing on or supplementing growth with organic compounds), whereas the CarveMe universal model is based on, and validated with, heterotrophic bacterial genomes. Given that CarveMe was designed for heterotrophic microbes and only 0.44% of the genomes used for subsequent analyses were from the PCC-6307 order, we focused our analyses on heterotrophic metabolic strategies.

### Metabolic strategy assessment

We defined metabolic strategy as the substrates that are preferred by an organism for growth. We assessed the metabolic strategy for each genome using a suite of sensitivity studies. Specifically, model growth dynamics were evaluated using CobraPy under ‘replete’ conditions (all potential growth substrates available), and then under ‘limiting’ conditions in which the availability of certain compound classes were substantially reduced. Here we used a threshold of an 80% reduction in growth rate under the limiting condition as the designation of substantial reduction in growth (Supplemental Figure S2). We then considered a genome to be sensitive to a compound class if growth was substantially reduced when the compound class was removed.

We validated our approach using an extensive culture-based analysis of carbon substrate preferences for 191 marine microbes^16^. Good agreement was observed between the CarveMe model predictions for these genomes and the experimentally validated growth preferences (Supplemental Figure S3a). Specifically, 51.0% of the comparisons showed exact agreement between the model predictions and experimental data and only 4.6% of models predicted no growth where growth was experimentally observed. For the remaining 44.4% of the comparisons, the models predicted that the substrate could be taken up by the organism but no growth was experimentally observed when that compound was provided as a sole carbon source. These cases suggest that the organisms might be able to use the substrate, but not as a sole carbon source or under the conditions tested. Good agreement was also seen between model-predicted compound sensitivities and the designation of acid versus sugar specialists identified by^16^ (Supplemental Figure S3b), suggesting that our framework is capturing substrate preferences observed experimentally (Supplement S1).

The highest growth sensitivity occurred under carboxylic acid limitation, with 39.9% of all models in the dataset demonstrating substantial growth reduction when the uptake of this compound class was limited (Supplemental Figure S4a). Reducing the availability of amino acids or carbohydrates resulted in substantial growth reduction in 29.2% and 17.6% of the models, respectively (Supplemental Figure S4a). In contrast, the models were generally insensitive to the reduction of amines/amides, ketones/aldehydes, or alcohols with only 0.13%, 0.2%, and 0.25% of models showing substantial growth reduction, respectively (Supplemental Figure S4a). When analyzed by taxonomic order, we found that the most sensitive taxa to substrate limitation across all compound classes were the *Pelagibacterales* (24.9% of models showed substantial growth reduction to at least one compound class), SAR86 (21.6% showed substantial growth reduction), and AEGEAN-169 (19.1% showed substantial growth reduction) (Supplemental Figure S5a). This is consistent with these groups being classically oligotrophic organisms with streamlined genomes and limited metabolic flexibility^17–19^. By contrast, the classically copiotrophic groups (the *Enterobacterales*, *Sphingomonadales*, and *Rhodobacterales*) showed the least growth sensitivity to substrate reduction, with only 5.9%, 9.1% and 9.4% of these models showing substantial growth reduction across all compound classes, respectively (Supplemental Figure S5a). This indicates that the classically designated copiotrophic orders may have more flexible metabolisms where they can achieve high growth rates using many different compound classes.

### Emergent metabolic clusters

Metabolic niches were identified based on the substrate preference profiles for all 1,591 genomes using Self Organizing Maps (SOMs). The SOMs method is an unsupervised clustering method that reduces large, high dimensional datasets to a topologically defined two-dimensional grid space^20^. Eight SOM clusters emerged with distinct growth sensitivities to different compound classes (Table 1). Differences in sensitivities to carbohydrates, carboxylic acids, amino acids, peptides, and B-vitamins drove the largest separations between the clusters (Figure 2, Supplemental Figure S4b). To further expand the analysis of growth strategy, we computed an estimate of maximum growth rate for each genome based on genomic optimization using codon usage bias (dCUB)^21,22^. We then assessed differences in genomic estimates of maximum growth rates between clusters (dCUB values were not used in the SOM clustering). Significant differences in estimated maximum growth rates were observed between the SOM clusters (Tukey’s HSD, ANOVA) with one fast-growing cluster (Cluster 2), four slow-growing clusters (Clusters 1, 3, 7, and 8), and three clusters with intermediate growth rates (Clusters 4, 5, and 6) (Supplemental Figure S6, Table 1).

**Figure 2:**
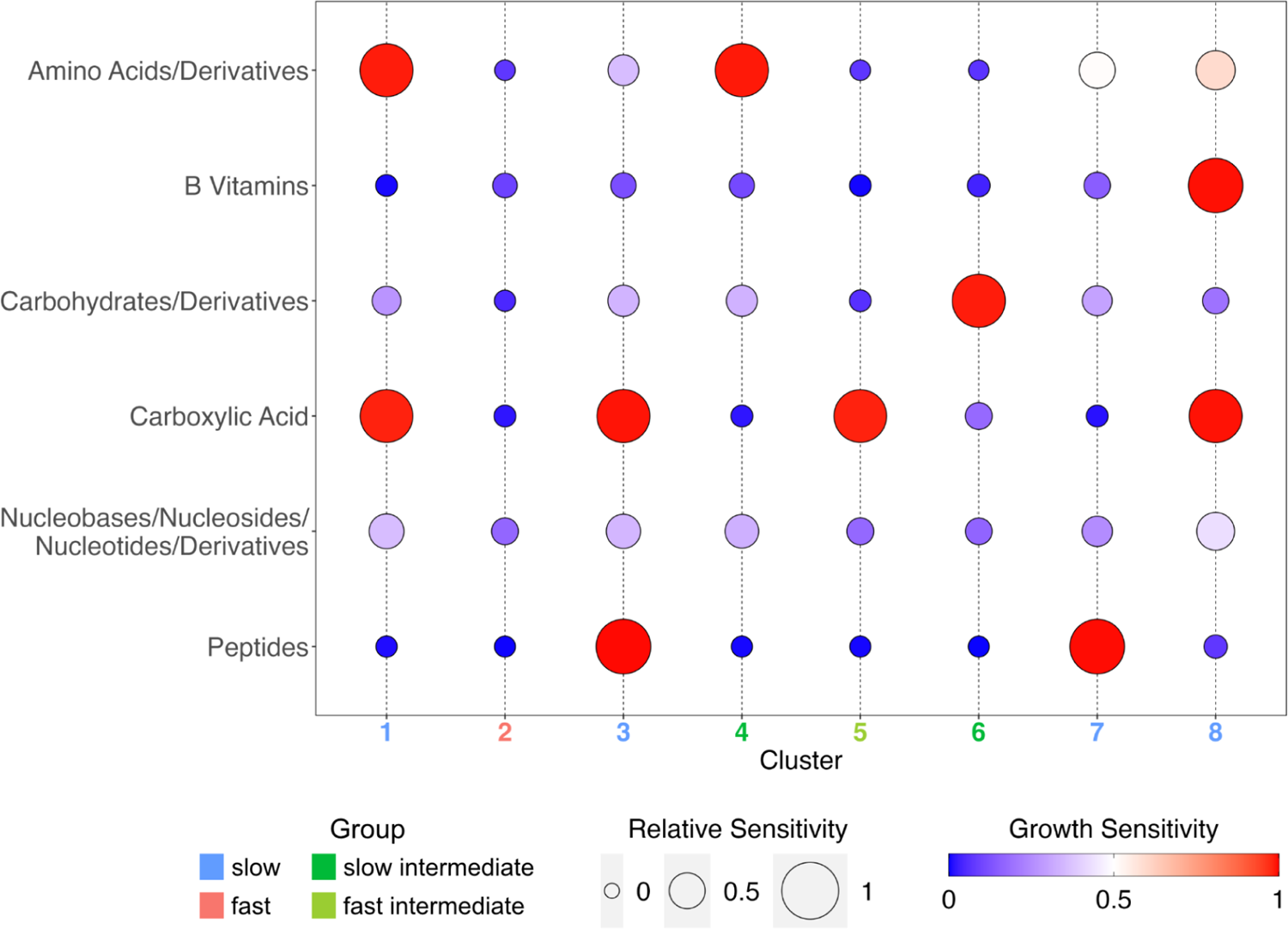
Substrate sensitivities for 8 SOM clusters. Bubble plot of the mean growth sensitivity values for genomes in each of our 8 SOM clusters. A growth sensitivity of 1 indicates high sensitivity to that substrate such that the modeled growth rate was reduced proportionally to the reduction in the substrate’s flux (e.g., 50% substrate reduction corresponded to 50% growth rate reduction). The size of the bubbles in this plot reflect the relative sensitivity of each of the 8 SOM clusters to a given compound class where larger bubbles indicate that cluster was more sensitive to that compound class than others. The 6 compound classes which resulted in significant growth reduction for at least one of the SOM clusters are shown here. The full results for all 11 substrate classes are provided in Supplemental Figures S2 & S3). Cluster numbers were colored based on maximal genomic growth rate (Supplemental Figure S6).

**Table 1:** Description of 8 SOM clusters including the number of genomes per cluster, the growth strategy as determined by the dCUB distributions, the number and names of the growth limiting substrate classes, as well as the 2 most numerically abundant orders.

| Cluster | Genomes | Growth Strategy | % fast growers | Limiting Classes |  | Top 2 orders |
| --- | --- | --- | --- | --- | --- | --- |
|  |  |  |  | # | Substrate |  |
| 1 | 144 | Slow | 50.00% | 2 | Carboxylic Acids, Amino Acids | Pelagibacterales, Rhodobacterales |
| 2 | 558 | Fast | 78.00% | 0 | None | Flavobacteriales, Enterobacteriales |
| 3 | 95 | Slow | 40.00% | 2 | Carboxylic Acids, Peptides | Flavobacteriales, Pelagibacterales |
| 4 | 211 | Slow Intermediate | 62.10% | 1 | Amino Acids | Flavobacteriales, Rhodobacterales |
| 5 | 299 | Fast Intermediate | 73.20% | 1 | Carboxylic Acids | Pseudomonadales, Rhodobacterales |
| 6 | 133 | Slow Intermediate | 66.20% | 1 | Carbohydrates | Flavobacteriales, Rhodobacterales |
| 7 | 77 | Slow | 42.90% | 2 | Peptides, Amino Acids | Flavobacteriales, Rhodobacterales |
| 8 | 74 | Slow | 48.60% | 3 | Carboxylic Acids, Amino Acids, B vitamins | Flavobacteriales, Rhodobacterales |

The eight SOM clusters had conserved phylogenetic signals with enrichment in specific taxonomic groups (Table 1). However, we simultaneously observed that many taxonomic groups appeared across multiple clusters (Supplemental Figure S5b). This suggests that diverse taxonomic groups have similar substrate preferences and growth sensitivities, and also that some taxonomic groups appear to have sublineages with wide variations in lifestyle. For example, the *Enterobacterales* and *Rhodobacterales* were enriched in our fast-growing Cluster 2 by 174.6% and 121.0%, respectively, relative to their abundances in the total dataset (Supplemental Figure S7). Similarly, SAR86 and the *Pelagibacterales* were on average enriched in the slow-growing Clusters 1, 3, 7, and 8 by 211% and 307%, respectively, relative to their abundances in the total dataset. This was compared to 74.1% and 80.6% reductions of these two classically oligotrophic orders in the fast-growing Cluster 2 relative to the total dataset. However, we also found that nine of the fifteen most abundant orders were present in all clusters, and only three orders were absent in more than one cluster (the *Sphingomonadales*, PCC-6307, and AEGEAN-169). The *Flavobacteriales*, for instance, were present across all eight clusters accounting for 9.7% of slow-growing Cluster 1 up to 33.1% of the intermediate growth Cluster 6 (Supplemental Figure S5b). Thus, although there was variation in the taxonomic composition of the clusters, the differences between clusters was not driven by taxonomy alone (Supplement S3.3). Below we provide an analysis of the four growth types that emerged from the SOM clusters: fast-growing, slow-growing, fast intermediate growth, and slow intermediate growth.

The fast-growing Cluster 2 was classically ‘copiotrophic’. 78.0% of genomes in this cluster had predicted maximum genomic growth rates that were higher (more negative dCUB) than the threshold of slow growth (*dCUB* =− 0. 08, where lower dCUB values correspond to faster growth). This threshold corresponds to a doubling time of approximately 5 hours for mesophilic organisms (optimal growth temperature between 20-60°C)^22^ (Supplemental Figure S6). Taxonomically, Cluster 2 consisted primarily of the *Enterobacterales*, *Flavobacteriales*, *Rhodobacterales*, and *Psuedomonadales* (Figure 1, Supplemental Figure S5a). Metabolically, this fast-growing cluster showed the least sensitivity to the removal of compounds with no significant growth sensitivity to the reduction of any of the 11 measured compound classes (Figure 2). This suggests that these organisms have flexible metabolisms capable of growing on a wide range of substrates and are able to synthesize or substitute essential metabolites when not available from the environment. Hereafter, we will refer to this as the fast-growing generalist cluster.

In contrast, the slow-growing clusters (Clusters 1, 3, 7, and 8) had significantly slower estimated maximum growth rates than the fast-growing generalist cluster (average dCUB of −0.106) (Supplemental Figure S6). For example, 60% of genomes in Cluster 3 had dCUB values within the ‘indistinguishable slow growth range’ (dCUB values above the −0.08 threshold). This cluster had a high proportion of known oligotrophic orders such as SAR86 and the *Pelagibacterales*, with these groups enriched in this cluster by 457% and 343% relative to the overall dataset (Supplemental Figure S6). The *Marinisomatales* was also found to be slightly enriched (139.6%) in this cluster relative to its abundance in the overall dataset. All four slow-growing clusters showed growth sensitivities to multiple (two or more) compound classes (Figure 2). This is in contrast to the intermediate growth clusters which demonstrated sensitivity to a single compound class (*described below*). We observed metabolic niche separation within the slow-growing clusters. For example, Cluster 1 exhibited high growth sensitivities to two classes of acids (carboxylic acids and amino acids/derivatives), whereas Cluster 3 models showed high growth sensitivities to carboxylic acids and peptides (Figure 2, Supplemental Figure S4b). Cluster 7 showed growth sensitivity to peptides and amino acids, while Cluster 8 models were sensitive to carboxylic acids, B vitamins, and amino acids.

The three intermediate growth clusters (Clusters 4, 5, and 6) showed growth sensitivity to a single compound class: amino acids, carboxylic acids, and carbohydrates, respectively (Figure 2B). All three intermediate growth clusters had predicted growth rates that were significantly slower than the fast-growing Cluster 2 (Supplemental Figure S6). Cluster 5 was also estimated to be significantly faster than the four slow-growing clusters – hereafter referred to as the fast intermediate growth cluster. Overall, these intermediate growth clusters appeared to be more flexible metabolically and faster-growing than the slow-growing specialist clusters but more specialized and slower growing than the fast-growing generalist cluster. The intermediate growth clusters corroborate a recent modeling study which suggested that the dominant heterotrophic group in the subsurface ocean might be slow-growing copiotrophs^2^.

### Biogeographic Distribution

To investigate the biogeographic distributions of our eight SOM clusters, we performed a competitive metagenomic recruitment and calculated normalized Reads Per Kilobase per Million mapped reads (RPKM) for 1,424 globally distributed samples. We compared the relative abundance of aggregate total RPKM for each of the 23 unique oceanographic regions in our global dataset using a bootstrapping approach (Supplemental Figure S8). We further grouped these 23 regions into 5 oceanographic categories and applied our bootstrapping approach, as well as a direct clustering on the raw RPKM values from each site. Clear biogeographical patterns emerged across both the 23 oceanographic regions and the 5 defined oceanographic categories (Figure 3, Supplemental Figures S8&S9). Estuarine sites showed the highest abundance of both the fast-growing generalist and the fast-growing intermediate clusters. The co-occurrence of these copiotrophic and metabolically flexible organisms in these frequently eutrophic and variable salinity environments aligns with our expectation that microbial communities present in these regions are dominated by fast-growing organisms. In contrast, these faster growing clusters were rare at open ocean oligotrophic sites where copiotrophs are primarily present at low abundances, occupying niches such as sinking particles^23^. We show that community compositions in the oligotrophic seas (Mediterranean and Red Sea) and open ocean sites are dominated by the four slow-growing clusters and the two slower-growing intermediate clusters. The slow-growing acid-specialist cluster (Cluster 1) was the most numerically dominant group of organisms across all samples and also had the greatest enrichment of the *Pelagibacterales*, the most numerically dominant order of marine heterotrophs^17^. Unfortunately, the dataset did not allow us to make further conclusions related to the co-occurrence of specific metabolic preferences and the biogeochemical environment in which these communities were found. However, expanding this analysis to samples where interdisciplinary information (e.g. metagenomic, metabolomic, organic matter composition, and rate measurements) are co-collected is an exciting avenue for future work.

**Figure 3:**
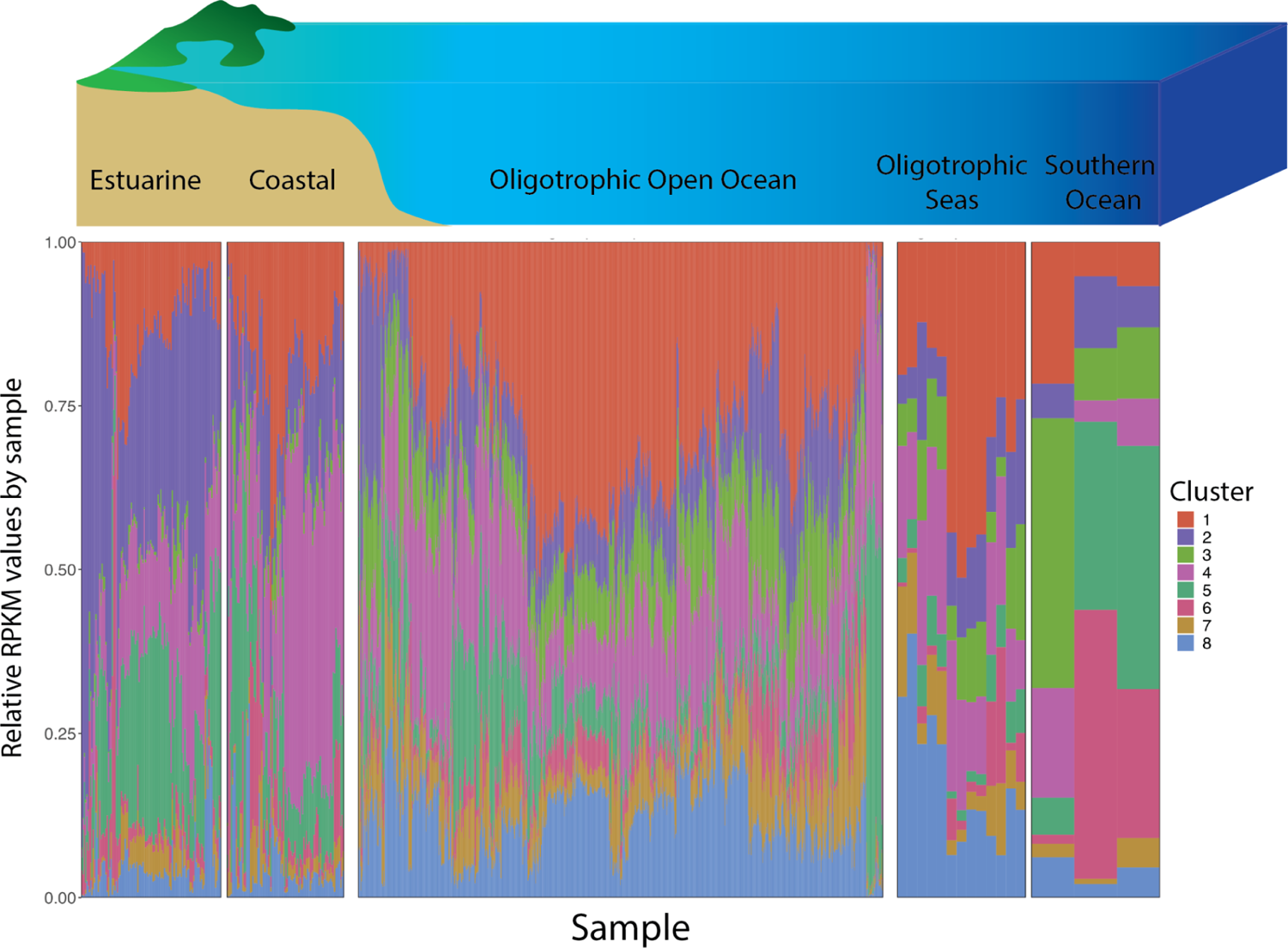
Biogeographical relative abundances of 8 SOM clusters. Clustered bar charts of the relative abundances of the 8 SOM clusters as determined by RPKM at each of the 1,203 stations assigned to one of the 23 oceanographic regions. Stations were grouped into our 5 defined oceanographic categories and then arranged based on a hierarchical clustering of the relative abundances.

## Discussion

Although marine microbial heterotrophs play a primary role in regulating organic matter cycling, biogeochemical cycles, and global climate outcomes, we have historically lacked an overarching framework for characterizing these diverse communities and assessing their functional metabolic niches. Since the majority of microbial heterotrophic diversity in the oceans remains uncultured^24^, we must rely on indirect methods to assess these metabolic strategies. By combining metagenomic information, numerical models, and statistical approaches, we identified eight distinct metabolic strategies for marine heterotrophic metabolism. Critically, our approach is a high throughput method for generating key metabolic and physiological insight that could previously only be obtained through labor intensive laboratory experiments restricted to cultured organisms. The hypothesized clusters generated by this analysis provide a set of microbial building blocks on which we can understand the assemblage of global heterotrophic microbial communities and how they differ by oceanographic region.

We demonstrated that, when applied correctly, the CarveMe tool provides fundamental insights into the metabolism of a highly diverse set of marine heterotrophic organisms. We additionally showed that there were clear biases in the quality of models generated by CarveMe with some orders, such as the *Pelagibacterales* that consistently produced poor quality models. There are likely several factors that result in poor quality models. We postulate that the current universal model and cutting algorithm used by CarveMe may struggle with streamlined genomes (e.g., the *Pelagibacterales*) and for marine heterotrophs that specialize in growth on more complex carbon substrates. We also hypothesize that issues with annotation, in particular for transporters, might also contribute to poor quality models for certain groups. We suggest that including additional reference genomes with validated metabolic models for certain orders (e.g., the *Pelagibacterales*) will substantially improve our ability to generate high quality metabolic models across diverse groups.

While the clusters identified in this analysis are robust, they are not necessarily complete. In particular, we demonstrated that the CarveMe tool was not successful at creating high quality models for the majority of genomes in many key microbial groups. Thus, we anticipate that once we can create high quality models for these groups and investigate their metabolic strategies, we will potentially identify additional meaningful clusters.

Here we used metabolic models to analyze the growth strategies for a large number of marine microbial genomes (the majority uncultured) via *in silico* methods. We identified eight clusters with distinct substrate preferences, growth strategies, taxonomic profiles, and biogeographic distributions (Table 1, Supplemental Table S1). We demonstrated that some growth strategies correspond strongly with phylogeny, suggesting that we can infer metabolism directly from phylogeny in some cases. However, the majority of the phylogenetic groups in our dataset were distributed across multiple clusters with distinct metabolic preferences, demonstrating that organisms from diverse taxonomic groups can occupy the same metabolic niche, consistent with the findings of widely varying growth rates within closely related organisms^25^. Our approach also provides a new resource for artificial media development by identifying key growth-limiting substrates whose absence in traditional media may currently be inhibiting culturing efforts. Finally, our metabolic clusters provide a framework for developing biogeochemical models that explicitly incorporate diverse microbial communities by specifying specific growth strategies and metabolic preferences that can be used to parameterize these groups.

## 2. Methods

### 2.1 Genomic Data

Genomic data was obtained from the Ocean Microbial Database (OMD) hosted at microbiomics.io^13^, which contains approximately 35,000 microbial genomes including metagenome-assembled genomes (MAGs), single amplified genomes (SAGs), and cultured isolates. We included only high quality bacterial genomes as defined by standard thresholds of > 80% completeness and < 5% contamination^26,27^. These estimates were determined based on the average of the CheckM^28^ and Anvi’o^29^ completeness and contamination scores. High quality genomes were then dereplicated using dRep^30^ with a 95% ANI threshold which was provided in the OMD metadata. We used the resulting 3,918 high-quality dereplicated bacterial genomes as our preliminary dataset for analysis.

### 2.2 Phylogeny

The phylogenetic tree of the 3,918 bacterial genomes was determined using GtoTree v1.7.0^31^ and IQ-TREE v2.0.3^32^. We also included the 66 unique bacterial reference genomes underlying the bacterial metabolic models in the BiGG database^33^ that was used to generate CarveMe’s universal reaction model^7^. From these 3,984 total genomes, we created a multiple sequence alignment (MSA) file using the predefined Bacteria single copy gene (SCG) set in GToTree v1.7.00^31^. During this process, eight genomes were excluded from the tree due to an insufficient number of hits to the target SCG set resulting in an alignment file of 3,976 genomes. However, these eight genomes were still included in our taxonomic analyses of the SOM clusters as we were able to assign their phylogeny using the Genome Taxonomy Database (GTDB). The MSA file was then passed to IQ-TREE v2.0.3 using the LG+R10 model with 3,554 amino-acid sites to generate a phylogenetic tree (Figure 1). For a current taxonomy of all genomes in the dataset, we overlaid full taxonomic assignments generated by GTDB-Tk v2.1.0^34^ with the GTDB r214 database^35^ onto this tree.

We also calculated a quantitative measure of phylogenetic relatedness, the UniFrac distance^36^, for subgroups of genomes we defined based on several external parameters (e.g., SOM cluster, ensemble consensus score, dCUB, etc.). For example, we created sub-datasets containing genomes assigned to each of our eight SOM clusters that we compared using UniFrac. UniFrac-Binaries^37^ was run using the Strided State UniFrac algorithm^38^ to compute unweighted UniFrac scores^39^. These results from this assessment are presented in Supplemental Section S3.3.

### 2.3 Model Generation and Quality Assessment

CarveMe v1.5.1^7^ was used to generate multiple metabolic models for each genome, called an ensemble. CarveMe’s ensemble function creates multiple models from a single genome by randomizing the weighting factors for unannotated reactions before generating each model using its mixed integer linear programming (MILP) algorithm. Annotated reactions receive weighting factors based on their gene-protein-reaction score, a metric that reflects the level of confidence in whether all proteins and subunits required for a reaction to take place are supported in the genome. We tested a variety of ensemble sizes ranging from 2 models to 100 models to assess the necessary number of model replicates to effectively capture the reaction space of each genome (Supplemental Figure S10). We found that the new reactions added to the total reaction space started to plateau around an ensemble size of 60 suggesting that 60 models were sufficient to capture the majority of possible model solutions.

For each of the 3,918 high quality genomes in our dataset, 60 models were generated using their protein fasta files as input. CarveMe^7^was run using python 3.7^40^ and IBM ILOG CPLEX Optimizer v20.1.0, using the native DIAMOND annotation procedure v0.9.14^41^. To assess the quality of the metabolic models generated by CarveMe, we developed a consensus score metric *C*. The consensus score is defined as:

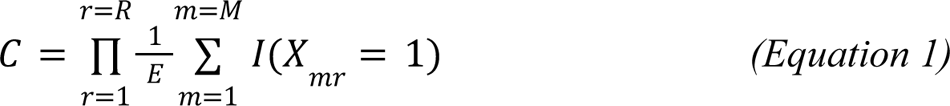

where *X_mr_* is the presence/absence matrix of ensemble model reactions across *M* individually generated models, *r* is an individual reaction, *R* is the total number of reactions in the ensemble, and *E* is the ensemble size. In this context, *I* is the indicator function for the case that reaction *r* is present in ensemble model *m*. In plain terms, *C* measures the consistency of the CarveMe model reactions across the ensemble generated for a single genome. If all models in the ensemble contained all the same reactions then the consensus score would be 1, if only half of the models had the same set of reactions then the consensus score would be 0.5. Similar to Machado et al. 2018, we equate the consensus score with overall ensemble quality because significant dissimilarities between ensemble models suggest that the cutting algorithm in CarveMe was forced to make more uninformed choices. On the other hand, an ensemble with highly consistent models suggests that the cutting procedure had sufficient knowledge to consistently include the correct pathways in the model. Only genomes with a consensus score greater than 0.8 were used for further analyses (N=1,591).

### 2.4 Compound Classification

To provide an assessment of broad metabolic strategies, we analyzed the CarveMe model growth sensitivities by compound classes. To do this, compounds that were used as substrates (imported into the cell by the model) were manually classified into the following 13 major categories: carboxylic acids, amino acids and derivatives, peptides, nucleobases/nucleosides/nucleotides and derivatives, carbohydrates and derivatives, ketones/aldehydes, organic sulfur, phospholipids/fatty acids and triglycerides, alcohols, amines and amides, B vitamins, inorganics, and “other” (Supplemental Table S2). We excluded inorganics and ‘other’ categories from our downstream analyses to focus on the eleven categories with organic substrates necessary for growth. References used in the categorization included ChEBI^42^, NIH PubChem^43^, BiGG Database^44,45^, HMDB^46^, BioCyc^47,48^, ChemSpider^49^, ECMDB^50,51^ and prior knowledge. There were 2,467 external exchange reactions in the universal model representing the acquisition of compounds from the environment or media. Only 633 of the 2,467 appeared as external reactions in any of our models. We then classified the 456 compounds that showed up the most frequently and accounted for the majority of the total flux into the models across all 95,460 CarveMe models generated for this study. Specifically, our classified compounds accounted for at least 90% of the total import flux in 98.3% models across all of our sensitivity tests (N=1,050,060). The 177 compounds that were not included each appeared in fewer than 10 of the 95,460 total models in the dataset.

### 2.5 Growth sensitivity analysis

We used the CobraPy v0.25.0^11^ software to test the CarveMe model growth sensitivities under a wide range of substrate availability. First, we assessed the type and quantity of compounds preferred for growth for each of the CarveMe models under replete conditions (where we define replete conditions as having maximum flux of all substrates available to the model). Specifically, we estimated the maximum model growth rate using the slim_optimize function in CobraPy with all possible media components turned on. We then determined the minimal set of compounds that allowed the previously determined maximum model growth rate using the CobraPy minimal media prediction (minimal_media function). This function solves a mixed integer linear programming (MILP) problem to minimize the import fluxes (external exchange reactions) while maintaining the maximum model growth rate.

Compound-specific growth sensitivities for each of our 11 growth compound classes were then determined for each model. For each substrate compound class (*defined above in section 2.4*), the available flux for that class was supplied at 50% of the import flux value in the ‘replete conditions’ while all other substrates were allowed to reach their maximum values. Any medium component from the limited growth compound class that was not originally predicted as part of the minimal medium of a given model was made unavailable to prevent models from circumventing the substrate limitation. We then assessed how the substrate import fluxes shifted under these limitation scenarios and the resulting change in predicted growth rate. Sensitivities were computed on a [0, 1] scale using the following equation:

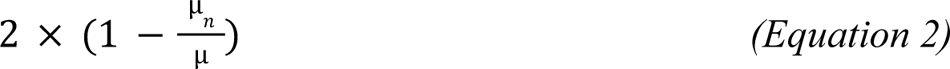

where µ*_n_* is the predicted growth rate under substrate limitation by compound class *n* and µ is the predicted growth rate in the ‘replete conditions’. The eleven compound-specific growth sensitivities that were estimated per model then served as the input data for the SOM clustering. The full enumeration of the compound specific growth sensitivities for each genome can be found in Supplemental Table S1.

### 2.6 Validation on Experimentally Characterized Genomes

To validate our CarveMe models and growth sensitivity analysis, we compared our model results to experimentally validated measurements on a shared set of genomes. Specifically, we constructed model ensembles for 176 marine bacterial genomes that were experimentally tested for their ability to grow on a variety of sugar/acid substrates^16^. The data from this study included binary measurements of growth/no growth on 118 compounds as the sole carbon substrate in the media and a prediction of sugar/acid preference (SAP). Of the 176 genomes, 146 generated high quality CarveMe models (above the consensus threshold of 0.8).

We conducted a paired comparison of the experimental and model predictions of substrate growth for the 146 high quality genomes that had been assessed for growth on the range of carbon substrates. Of the 118 experimentally measured compounds, 59 had corresponding reactions in the BiGG database. Our results exclude one of these 59 compounds, oxaloacetate, which is highly unstable and rapidly degrades to pyruvate^52^ and confounds the fidelity of the growth experiment with that compound as the sole carbon source. For presence in the model, we required the external exchange reaction for the substrate to be present in >80% of the ensemble models. For absence, we required that the exchange reaction be absent in >80% of the ensemble models. 1.5% of the model-substrate comparisons fell between these two cutoffs and so were not assigned an outcome. For each genome and each substrate, we assigned one of three outcomes: 1) agreement between the models and the data (either presence/growth or absence/no-growth); 2) disagreement between the models and the data (absence in the model/growth in the data); or 3) false positives where the models contained the exchange reaction but the organism was not able to grow on the substrate as a sole carbon source.

To assess the agreement between the modeled growth sensitivities of these organisms and the experimental findings for sugar/acid preference (SAP), we computed the growth sensitivities of the CarveMe model ensembles (see Section 2.5). We used the sensitivity to carbohydrates for the sugar preference assessment and the sensitivity to amino acids for the acid preference assessment. The 146 genomes with high quality models were grouped into two groups based on whether they were sugar-preferring organisms (SAP > 0, N=77) or acid-preferring organisms (SAP < 0, N=69). We then compared the relative sensitivities for each of these classes to their experimentally assigned SAP value by determining the average growth sensitivity of the models associated with the genomes in each group to our sugar and acid compound classes. The results from this assessment are presented in Supplemental Section S1.

### 2.7 Generation of Self-Organized Maps

To identify clusters of organisms with similar metabolic strategies, we employed Self-Organized Maps (SOMs) to the assessment of compound specific growth sensitivities. SOMs are an unsupervised machine learning dimension reduction method capable of handling large data formats^53^. SOMs are a non-parametric approach, capable of highlighting nonlinear, complex patterns in two-dimensional space from highly dimensional data. The map was built using the CobraPy growth sensitivity analysis for the 95,460 high-quality ensemble models (*C* ≥ 0. 8). These scaled compound flux predictions were clustered using kohonen v3.0.12^20^ and solved over 1,500 iterations with a learning rate vector of (0.025, 0.01) and default neighborhood radii on a 20-by-20 toroidal, hexagonal grid spatially described by standard Euclidean distance. Map parameters were determined using heuristics and metrics of error proposed in the SOMs literature^53–57^ and are discussed further in Supplemental Section S2. Each node in the grid was initialized with a random codebook vector of values for each independent variable. Data entries were then randomly drawn from the dataset-every entry in the dataset was drawn in each iteration - and the grid point values of the closest neighborhood of nodes were updated. After sufficient training, the values assigned to each grid point reflect the spatial topology of the data (e.g., density of data points, variation) as well as the full range of values in the original dataset. The final SOM map was then grouped into eight distinct clusters using k-means clustering^58^ based on the coherence of the growth compound sensitivity predictions. The full map and designation of the clusters is shown in Supplemental Figure S11a. After 1,500 iterations, the mean object distance to its closest map unit (the quantization error) was approximately ^−4^ (Supplemental Figure S11b).

As the SOM map and clusters were built using all of the ensemble models (60 per genome), we then needed to assign each genome to a cluster. This was done by assigning each model to its closest mapping unit, and determining the mapping unit possessing a simple majority of the 60 models generated from a single genome. This mapping unit was designated as the mapping node for the genome and the genome was assigned to the associated SOM cluster. We assessed the frequency with which each of the 60 ensemble models occurred in a single SOM cluster (Supplemental Figure S11c) and showed that 96.0% of the genomes had 90% of their models assigned to the same SOM cluster (and 68.6% had all 60 models assigned to the same SOM cluster). Of the 1,591 genomes, only one genome had models split equally between two SOM clusters. In this case, this genome was randomly assigned to one of the two clusters using a fixed random seed of 123. The parameter optimization of the SOM map developed in this study is discussed in further detail in Supplemental Section S2.

### 2.8 Maximum Growth Rate Estimations

To assess differences in maximum growth rates, we estimated the codon usage bias (dCUB) for all 1,591 genomes using the gRodon program^21^. dCUB is a metric that has been empirically linked with optimization for faster growth. gRodon measures codon usage bias of highly expressed genes, in this case ribosomal proteins, compared to the codon usage patterns across the whole genome. This genomic measure of maximum growth is a reasonable proxy and allows us to examine the differences in growth optimization for this set of uncultured organisms without needing to do extensive culturing and metabolic characterization efforts. Because estimating actual growth rates from codon usage bias requires correcting for temperature, we used the raw dCUB scores for this analysis to assess relative differences in genomic optimization for rapid growth. Previous work by Weissman^21,22^ suggests that differences in dCUB values are only reliable below the threshold of −0.08 (i.e., lower values of dCUB represent faster growth rates). We use this threshold to differentiate between ‘slow growth’ and ‘fast growth’ organisms. The results from these analyses are presented in Supplemental Section S3.2.

### 2.9 Global Distribution

To assess the global distribution of the genomes within each SOM cluster, we performed competitive metagenomic recruitment. Specifically, we calculated normalized Reads Per Kilobase per Million mapped reads (RPKM) with the pipeline RRAP v1.3.2^59^. RRAP uses bowtie2 v2.4.2^60^ to align reads and SAMTools v1.14^61^ to index and sort the read alignment data. RRAP takes read alignment statistics generated by SAMTools to calculate RPKM. A total of 1,424 metagenomes were used for the read recruitment from several metagenomics surveys including Tara Oceans, BioGeoTraces, and Malaspina^13^. Raw metagenome fastq files were aggregated by sample and by depth when multiple depths were present – e.g., the Tara Oceans dataset – and quality filtered using the iu-filter-quality-minoche script from the Illumina-utils library v2.10 with default parameters. This script follows the quality filtering approach outlined in^62^. After quality filtering, our genome set was recruited to the metagenomic reads, and reads per kilobase per million mapped reads (RPKM) values were calculated for each genome at each site.

We then partitioned our data into 23 oceanographic regions defined in Lanclos et al. 2023 and aggregated the raw RPKM values for the genomes in our study^63^. Of the 1,424 sampling sites in the metagenomic recruitment, we had oceanographic region assignments from the metadata for 1,203 sites. The 23 defined oceanographic regions in this metadata averaged 52.3 distinct samples per region (ranging from 3 samples/sites in the Southern Ocean to 299 samples at station ALOHA) (Supplemental Table S3). We then further clustered the 23 oceanographic regions into 5 categories: Estuarine, Coastal, Oligotrophic Seas, Oligotrophic Open Oceans, and the Southern Ocean (Supplemental Table S4). We used this categorization to group the sampling sites and compare the relative abundances determined from the raw RPKM values. For the sampling sites associated with each category, we clustered the relative abundances of the eight SOM clusters at each site using Euclidean distance and hierarchical clustering with McQuitty linkage distance.

To assess the relative abundance of genomes assigned to each SOM cluster per oceanographic region or category, we conducted a bootstrap recruitment of the individual genome abundance values at each station. We employed bootstrapping due to large variation both in the number of samples present in each region (ranging from 3 at SOC to 299 at ALOHA) and in the number of genomes assigned to each cluster (ranging from 74 genomes in Cluster 8 to 558 genomes in Cluster 2). For each region, we computed 1,000 independent bootstrap iterations (using the fixed random seed 123), drawing 10,000 data points from the pool of RPKM samples for each of our eight clusters. During each bootstrapping step, we calculated the cumulative RPKM of the sampled data for each cluster and then compared their magnitudes to determine the relative abundances of the clusters. Average values and 95% confidence intervals were then computed from the resulting distributions of relative abundances for each cluster/region combination.

### 2.10 Data Visualization

All data visualizations in R v4.2.3 were performed using ggplot v3.4.2, ggridges v0.5.4^64^, ggtree^65^, patchwork v1.1.2^66^, ragg v1.2.5, and plots native to kohonen v3.0.12^20^.

## Acknowledgement

This work was supported by grants from the Simons Foundation (542389 to NML), NSF Biological Oceanography Program (OCE-1945279 to JCT), NSF Emerging Frontiers Program (EF-2125191 to NML and JCT) and Simons Investigator in Aquatic Microbial Ecology awards (LS-SIAME-00001997 to JCT and LS-SIAME-00001961 to NML).

## Author contributions

NML and JCT designed the study, RR and AW developed the framework, generated the models, and conducted the numerical analysis; CCJ contributed to the statistical analyses; CYK conducted the read recruitment; JLW conducted the dCUB analysis; all authors contributed to the writing.

## Data availability

All data used in this study are from previously published sources and are publically available. Genome identifiers are provided in Supplemental Table 1.

## Code availability

All code has been previously published and is referenced.

## References

1. Reynolds, R., Hyun, S., Tully, B., Bien, J. & Levine, N. M. Identification of microbial metabolic functional guilds from large genomic datasets. Front. Microbiol. 14, 1197329 (2023).

2. Zakem, E. J., McNichol, J., Weissman, J. L., Raut, Y., Xu, L., Halewood, E. R., Carlson, C. A., Dutkiewicz, S., Fuhrman, J. A. & Levine, N. M. Predictable functional biogeography of marine microbial heterotrophs. bioRxiv 2024.02.14.580411 (2024). doi:10.1101/2024.02.14.580411

3. Koch, A. L. Oligotrophs versus copiotrophs. Bioessays 23, 657–661 (2001).

4. Liu, S., Parsons, R., Opalk, K., Baetge, N., Giovannoni, S., Bolaños, L. M., Kujawinski, E. B., Longnecker, K., Lu, Y., Halewood, E. & Carlson, C. A. Different carboxyl-rich alicyclic molecules proxy compounds select distinct bacterioplankton for oxidation of dissolved organic matter in the mesopelagic Sargasso Sea. Limnol. Oceanogr. 65, 1532–1553 (2020).

5. Oberhardt, M. A., Palsson, B. Ø. & Papin, J. A. Applications of genome-scale metabolic reconstructions. Mol. Syst. Biol. 5, (2009).

6. Gu, C., Kim, G. B., Kim, W. J., Kim, H. U. & Lee, S. Y. Current status and applications of genome-scale metabolic models. Genome Biol. 20, 1–18 (2019).

7. Machado, D., Andrejev, S., Tramontano, M. & Patil, K. R. Fast automated reconstruction of genome-scale metabolic models for microbial species and communities. Nucleic Acids Res. 46, 7542–7553 (2018).

8. Henry, C. S., DeJongh, M., Best, A. A., Frybarger, P. M., Linsay, B. & Stevens, R. L. High-throughput generation, optimization and analysis of genome-scale metabolic models. Nat. Biotechnol. 28, 977–982 (2010).

9. Magnúsdóttir, S., Heinken, A., Kutt, L., Ravcheev, D. A., Bauer, E., Noronha, A., Greenhalgh, K., Jäger, C., Baginska, J., Wilmes, P., Fleming, R. M. T. & Thiele, I. Generation of genome-scale metabolic reconstructions for 773 members of the human gut microbiota. Nat. Biotechnol. 35, 81–89 (2017).

10. Mendoza, S. N., Olivier, B. G., Molenaar, D. & Teusink, B. A systematic assessment of current genome-scale metabolic reconstruction tools. Genome Biol. 20, 158 (2019).

11. Ebrahim, A., Lerman, J. A., Palsson, B. O. & Hyduke, D. R. COBRApy: COnstraints-Based Reconstruction and Analysis for Python. BMC Syst. Biol. 7, 74 (2013).

12. Régimbeau, A., Budinich, M., Larhlimi, A., Pierella Karlusich, J. J., Aumont, O., Memery, L., Bowler, C. & Eveillard, D. Contribution of genome-scale metabolic modelling to niche theory. Ecol. Lett. (2022). doi:10.1111/ele.13954

13. Paoli, L., Ruscheweyh, H.-J., Forneris, C. C., Hubrich, F., Kautsar, S., Bhushan, A., Lotti, A., Clayssen, Q., Salazar, G., Milanese, A., Carlström, C. I., Papadopoulou, C., Gehrig, D., Karasikov, M., Mustafa, H., Larralde, M., Carroll, L. M., Sánchez, P., Zayed, A. A., Cronin, D. R., Acinas, S. G., Bork, P., Bowler, C., Delmont, T. O., Gasol, J. M., Gossert, A. D., Kahles, A., Sullivan, M. B., Wincker, P., Zeller, G., Robinson, S. L., Piel, J. & Sunagawa, S. Biosynthetic potential of the global ocean microbiome. Nature 607, 111–118 (2022).

14. Bernstein, D. B., Sulheim, S., Almaas, E. & Segrè, D. Addressing uncertainty in genome-scale metabolic model reconstruction and analysis. Genome Biol. 22, 64 (2021).

15. Giordano, N., Gaudin, M., Trottier, C., Delage, E., Nef, C., Bowler, C. & Chaffron, S. Genome-scale community modelling reveals conserved metabolic cross-feedings in epipelagic bacterioplankton communities. Nat. Commun. 15, 2721 (2024).

16. Gralka, M., Pollak, S. & Cordero, O. X. Genome content predicts the carbon catabolic preferences of heterotrophic bacteria. Nat Microbiol 8, 1799–1808 (2023).

17. Giovannoni, S. J. SAR11 Bacteria: The Most Abundant Plankton in the Oceans. Ann. Rev. Mar. Sci. 9, 231–255 (2017).

18. Swan, B. K., Tupper, B., Sczyrba, A., Lauro, F. M., Martinez-Garcia, M., González, J. M., Luo, H., Wright, J. J., Landry, Z. C., Hanson, N. W., Thompson, B. P., Poulton, N. J., Schwientek, P., Acinas, S. G., Giovannoni, S. J., Moran, M. A., Hallam, S. J., Cavicchioli, R., Woyke, T. & Stepanauskas, R. Prevalent genome streamlining and latitudinal divergence of planktonic bacteria in the surface ocean. Proc. Natl. Acad. Sci. U. S. A. 110, 11463–11468 (2013).

19. Getz, E. W., Lanclos, V. C., Kojima, C. Y., Cheng, C., Henson, M. W., Schön, M. E., Ettema, T. J. G., Faircloth, B. C. & Thrash, J. C. The AEGEAN-169 clade of bacterioplankton is synonymous with SAR11 subclade V (HIMB59) and metabolically distinct. mSystems 8, e0017923 (2023).

20. Wehrens, R. & Buydens, L. M. C. Self- and Super-organizing Maps in R: The kohonen Package. J. Stat. Softw. 21, 1–19 (2007).

21. Weissman, J. L., Hou, S. & Fuhrman, J. A. Estimating maximal microbial growth rates from cultures, metagenomes, and single cells via codon usage patterns. Proc. Natl. Acad. Sci. U. S. A. 118, (2021).

22. Weissman, J. L., Dimbo, E.-R. O., Krinos, A. I., Neely, C., Yagües, Y., Nolin, D., Hou, S., Laperriere, S., Caron, D. A., Tully, B., Alexander, H. & Fuhrman, J. A. Estimating global variation in the maximum growth rates of eukaryotic microbes from cultures and metagenomes via codon usage patterns. bioRxiv 2021.10.15.464604 (2022). doi:10.1101/2021.10.15.464604

23. Fuhrman, J. A. Microbial community structure and its functional implications. Nature 459, 193–199 (2009).

24. Lloyd, K. G., Steen, A. D., Ladau, J., Yin, J. & Crosby, L. Phylogenetically Novel Uncultured Microbial Cells Dominate Earth Microbiomes. mSystems 3, (2018).

25. Deulofeu-Capo, O., Sebastián, M., Auladell, A., Cardelús, C., Ferrera, I., Sánchez, O. & Gasol, J. M. Growth rates of marine prokaryotes are extremely diverse, even among closely related taxa. ISME Commun ycae066 (2024).

26. Parks, D. H., Rinke, C., Chuvochina, M., Chaumeil, P.-A., Woodcroft, B. J., Evans, P. N., Hugenholtz, P. & Tyson, G. W. Recovery of nearly 8,000 metagenome-assembled genomes substantially expands the tree of life. Nat Microbiol 2, 1533–1542 (2017).

27. Tully, B. J., Graham, E. D. & Heidelberg, J. F. The reconstruction of 2,631 draft metagenome-assembled genomes from the global oceans. Sci Data 5, 170203 (2018).

28. Parks, D. H., Imelfort, M., Skennerton, C. T., Hugenholtz, P. & Tyson, G. W. CheckM: assessing the quality of microbial genomes recovered from isolates, single cells, and metagenomes. Genome Res. 25, 1043–1055 (2015).

29. Eren, A. M., Esen, Ö. C., Quince, C., Vineis, J. H., Morrison, H. G., Sogin, M. L. & Delmont, T. O. Anvi’o: an advanced analysis and visualization platform for ‘omics data. PeerJ 3, e1319 (2015).

30. Olm, M. R., Brown, C. T., Brooks, B. & Banfield, J. F. dRep: a tool for fast and accurate genomic comparisons that enables improved genome recovery from metagenomes through de-replication. ISME J. 11, 2864–2868 (2017).

31. Lee, M. D. GToTree: a user-friendly workflow for phylogenomics. Bioinformatics 35, 4162–4164 (2019).

32. Minh, B. Q., Schmidt, H. A., Chernomor, O., Schrempf, D., Woodhams, M. D., von Haeseler, A. & Lanfear, R. IQ-TREE 2: New Models and Efficient Methods for Phylogenetic Inference in the Genomic Era. Mol. Biol. Evol. 37, 1530–1534 (2020).

33. King, Z. A., Lu, J., Dräger, A., Miller, P., Federowicz, S., Lerman, J. A., Ebrahim, A., Palsson, B. O. & Lewis, N. E. BiGG Models: A platform for integrating, standardizing and sharing genome-scale models. Nucleic Acids Res. 44, D515–22 (2016).

34. Chaumeil, P.-A., Mussig, A. J., Hugenholtz, P. & Parks, D. H. GTDB-Tk v2: memory friendly classification with the genome taxonomy database. Bioinformatics 38, 5315–5316 (2022).

35. Parks, D. H., Chuvochina, M., Rinke, C., Mussig, A. J., Chaumeil, P.-A. & Hugenholtz, P. GTDB: an ongoing census of bacterial and archaeal diversity through a phylogenetically consistent, rank normalized and complete genome-based taxonomy. Nucleic Acids Res. 50, D785–D794 (2022).

36. Lozupone, C., Lladser, M. E., Knights, D., Stombaugh, J. & Knight, R. UniFrac: an effective distance metric for microbial community comparison. ISME J. 5, 169–172 (2011).

37. unifrac-binaries. (Github). at <https://github.com/biocore/unifrac-binaries>

38. McDonald, D., Vázquez-Baeza, Y., Koslicki, D., McClelland, J., Reeve, N., Xu, Z., Gonzalez, A. & Knight, R. Striped UniFrac: enabling microbiome analysis at unprecedented scale. Nat. Methods 15, 847–848 (2018).

39. Lozupone, C. & Knight, R. UniFrac: a new phylogenetic method for comparing microbial communities. Appl. Environ. Microbiol. 71, 8228–8235 (2005).

40. Van Rossum, G. & Drake, F. L. Python 3 Reference Manual: (Python Documentation Manual Part 2). (CreateSpace Independent Publishing Platform, 2009).

41. Buchfink, B., Xie, C. & Huson, D. H. Fast and sensitive protein alignment using DIAMOND. Nat. Methods 12, 59–60 (2015).

42. Hastings, J., Owen, G., Dekker, A., Ennis, M., Kale, N., Muthukrishnan, V., Turner, S., Swainston, N., Mendes, P. & Steinbeck, C. ChEBI in 2016: Improved services and an expanding collection of metabolites. Nucleic Acids Res. 44, D1214–9 (2016).

43. Kim, S., Chen, J., Cheng, T., Gindulyte, A., He, J., He, S., Li, Q., Shoemaker, B. A., Thiessen, P. A., Yu, B., Zaslavsky, L., Zhang, J. & Bolton, E. E. PubChem 2023 update. Nucleic Acids Res. 51, D1373–D1380 (2023).

44. Schellenberger, J., Park, J. O., Conrad, T. M. & Palsson, B. Ø. BiGG: a Biochemical Genetic and Genomic knowledgebase of large scale metabolic reconstructions. BMC Bioinformatics 11, 213 (2010).

45. Norsigian, C. J., Pusarla, N., McConn, J. L., Yurkovich, J. T., Dräger, A., Palsson, B. O. & King, Z. BiGG Models 2020: multi-strain genome-scale models and expansion across the phylogenetic tree. Nucleic Acids Res. 48, D402–D406 (2020).

46. Wishart, D. S., Guo, A., Oler, E., Wang, F., Anjum, A., Peters, H., Dizon, R., Sayeeda, Z., Tian, S., Lee, B. L., Berjanskii, M., Mah, R., Yamamoto, M., Jovel, J., Torres-Calzada, C., Hiebert-Giesbrecht, M., Lui, V. W., Varshavi, D., Varshavi, D., Allen, D., Arndt, D., Khetarpal, N., Sivakumaran, A., Harford, K., Sanford, S., Yee, K., Cao, X., Budinski, Z., Liigand, J., Zhang, L., Zheng, J., Mandal, R., Karu, N., Dambrova, M., Schiöth, H. B., Greiner, R. & Gautam, V. HMDB 5.0: the Human Metabolome Database for 2022. Nucleic Acids Res. 50, D622–D631 (2022).

47. Karp, P. D., Billington, R., Caspi, R., Fulcher, C. A., Latendresse, M., Kothari, A., Keseler, I. M., Krummenacker, M., Midford, P. E., Ong, Q., Ong, W. K., Paley, S. M. & Subhraveti, P. The BioCyc collection of microbial genomes and metabolic pathways. Brief. Bioinform. 20, 1085–1093 (2019).

48. Caspi, R., Billington, R., Keseler, I. M., Kothari, A., Krummenacker, M., Midford, P. E., Ong, W. K., Paley, S., Subhraveti, P. & Karp, P. D. The MetaCyc database of metabolic pathways and enzymes - a 2019 update. Nucleic Acids Res. 48, D445–D453 (2020).

49. Pence, H. E. & Williams, A. ChemSpider: An Online Chemical Information Resource. J. Chem. Educ. 87, 1123–1124 (2010).

50. Guo, A. C., Jewison, T., Wilson, M., Liu, Y., Knox, C., Djoumbou, Y., Lo, P., Mandal, R., Krishnamurthy, R. & Wishart, D. S. ECMDB: the E. coli Metabolome Database. Nucleic Acids Res. 41, D625–30 (2013).

51. Sajed, T., Marcu, A., Ramirez, M., Pon, A., Guo, A. C., Knox, C., Wilson, M., Grant, J. R., Djoumbou, Y. & Wishart, D. S. ECMDB 2.0: A richer resource for understanding the biochemistry of E. coli. Nucleic Acids Res. 44, D495–501 (2016).

52. Wilcock, A. R. & Goldberg, D. M. Kinetic determination of malate dehydrogenase activity eliminating problems due to spontaneous conversion of oxaloacetate to pyruvate. Biochem. Med. 6, 116–126 (1972).

53. Kohonen, T. The self-organizing map. Proc. IEEE 78, 1464–1480 (1990).

54. Park, Y.-S., Céréghino, R., Compin, A. & Lek, S. Applications of artificial neural networks for patterning and predicting aquatic insect species richness in running waters. Ecol. Modell. 160, 265–280 (2003).

55. Céréghino, R. & Park, Y.-S. Review of the Self-Organizing Map (SOM) approach in water resources: Commentary. Environmental Modelling & Software 24, 945–947 (2009).

56. Kalteh, A. M., Hjorth, P. & Berndtsson, R. Review of the self-organizing map (SOM) approach in water resources: Analysis, modelling and application. Environmental Modelling & Software 23, 835–845 (2008).

57. Kiviluoto, K. Topology preservation in self-organizing maps. in Proceedings of International Conference on Neural Networks (ICNN’96) 1, 294–299 vol.1 (IEEE, 1996).

58. Hartigan, J. A. & Wong, M. A. Algorithm AS 136: A K-Means Clustering Algorithm. J. R. Stat. Soc. Ser. C Appl. Stat. 28, 100–108 (1979).

59. Kojima, C. Y., Getz, E. W. & Thrash, J. C. RRAP: RPKM Recruitment Analysis Pipeline. Microbiol Resour Announc 11, e0064422 (2022).

60. Langmead, B. & Salzberg, S. L. Fast gapped-read alignment with Bowtie 2. Nat. Methods 9, 357–359 (2012).

61. Danecek, P., Bonfield, J. K., Liddle, J., Marshall, J., Ohan, V., Pollard, M. O., Whitwham, A., Keane, T., McCarthy, S. A., Davies, R. M. & Li, H. Twelve years of SAMtools and BCFtools. Gigascience 10, (2021).

62. Minoche, A. E., Dohm, J. C. & Himmelbauer, H. Evaluation of genomic high-throughput sequencing data generated on Illumina HiSeq and genome analyzer systems. Genome Biol. 12, R112 (2011).

63. Lanclos, V. C., Rasmussen, A. N., Kojima, C. Y., Cheng, C., Henson, M. W., Faircloth, B. C., Francis, C. A. & Thrash, J. C. Ecophysiology and genomics of the brackish water adapted SAR11 subclade IIIa. ISME J. 17, 620–629 (2023).

64. Wilke, C. O. ggridges: Ridgeline Plots in ‘ggplot2’. Preprint at https://wilkelab.org/ggridges/ (2024)

65. Yu, G., Smith, D. K., Zhu, H., Guan, Y. & Lam, T. T.-Y. Ggtree: An r package for visualization and annotation of phylogenetic trees with their covariates and other associated data. Methods Ecol. Evol. 8, 28–36 (2017).

66. Pedersen, T. L. patchwork: The Composer of Plots. Preprint at https://patchwork.data-imaginist.com (2024)

