## Supplementary Information for "Emergent Metabolic Niches for Marine Heterotrophs"

**Supplemental Figures:**

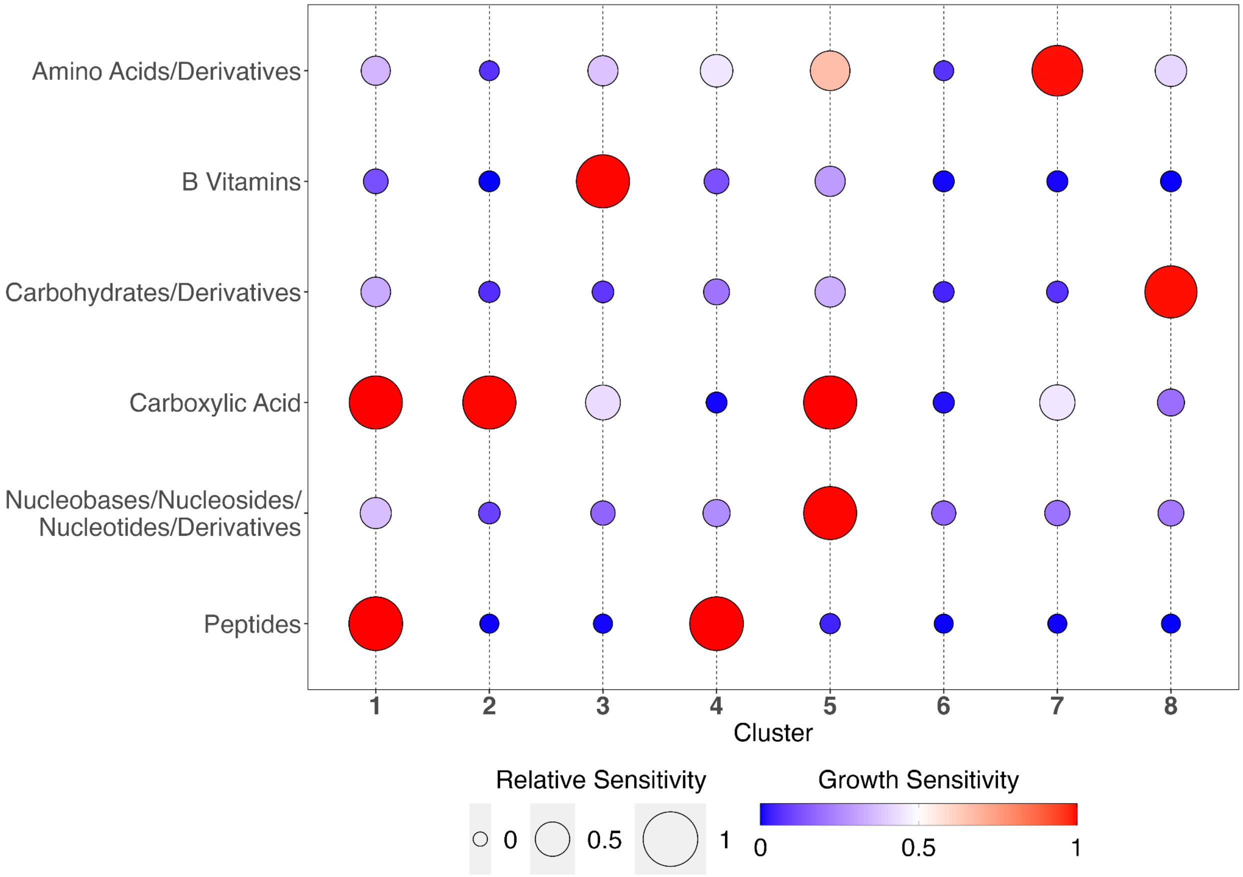


**Supplemental Figure S1:** Bubble plot of the mean growth sensitivity values (similar to Figure 2) for a new set of 8 SOM clusters generated on the 983 genomes with a consensus of 90% or greater. A growth sensitivity of 1 indicates high sensitivity to that substrate such that the modeled growth rate was reduced proportionally to the reduction in the substrate's flux (e.g., 50% substrate reduction corresponded to 50% growth rate reduction). The size of the bubbles in this plot reflect the relative sensitivity of each of the 8 SOM clusters to a given compound class where larger bubbles indicate that cluster was more sensitive to that compound class than others. The 6 compound classes which resulted in significant growth reduction for at least one of the SOM clusters are shown here. While the ordering of the clusters changed, we still observed the same overall patterns. We have one cluster with no growth sensitivities and multiple clusters with sensitivity to one compound and multiple with sensitivity to two compounds. The fast growth cluster and intermediate growth single sensitivity clusters from the primary analysis emerged in this higher consensus group of models. The slight shift for the multiple sensitivity clusters is consistent with the observation that the more classically oligotrophic orders generally had model ensembles with lower consensus values such that excluding these genomes from the SOM generation would be expected to have the largest impact on the slow growth clusters.

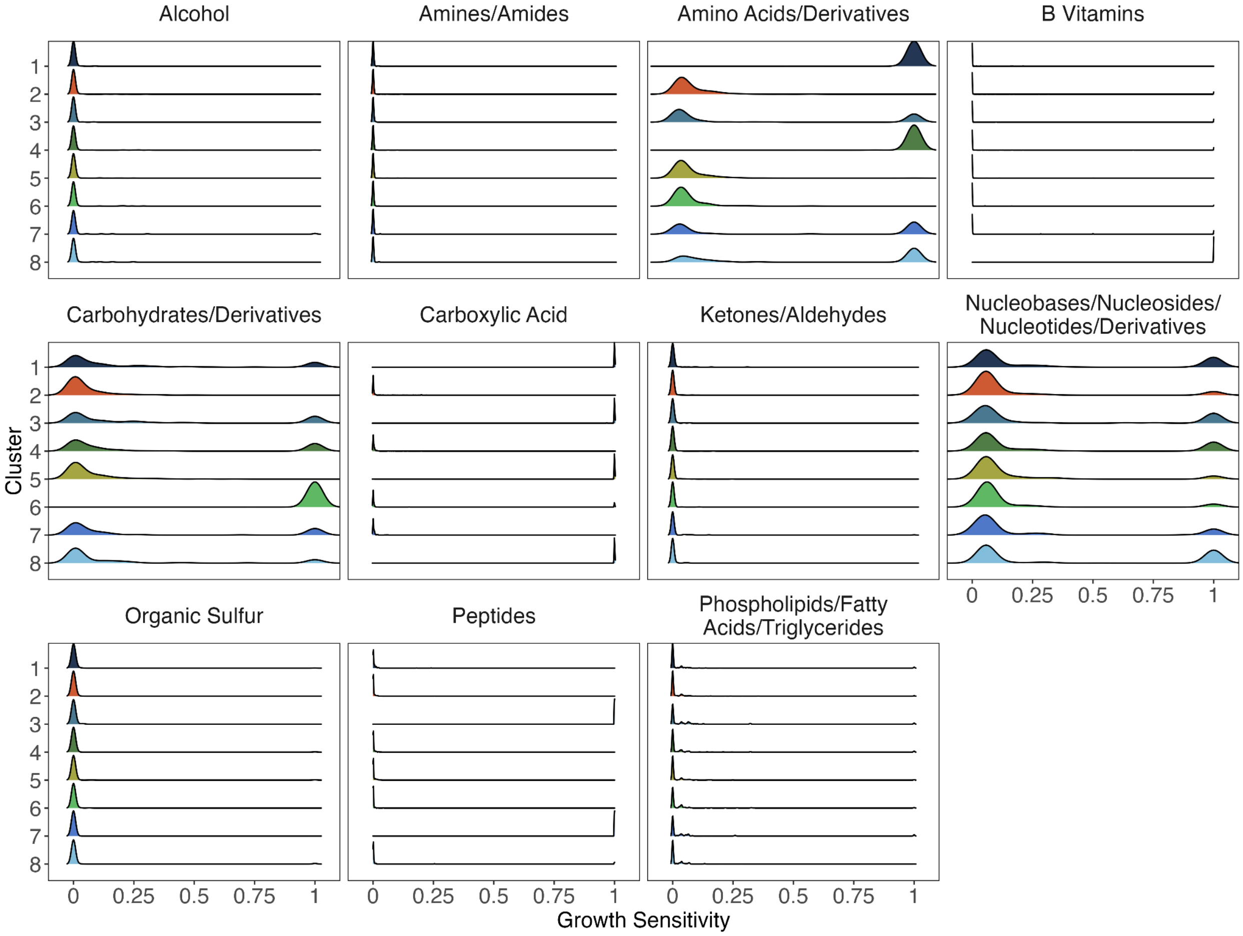


**Supplemental Figure S2: Distribution of growth sensitivity values by cluster.** Density plots of the growth sensitivity values for each model for each of the 11 compound classes grouped by SOM cluster (N=1,050,060).

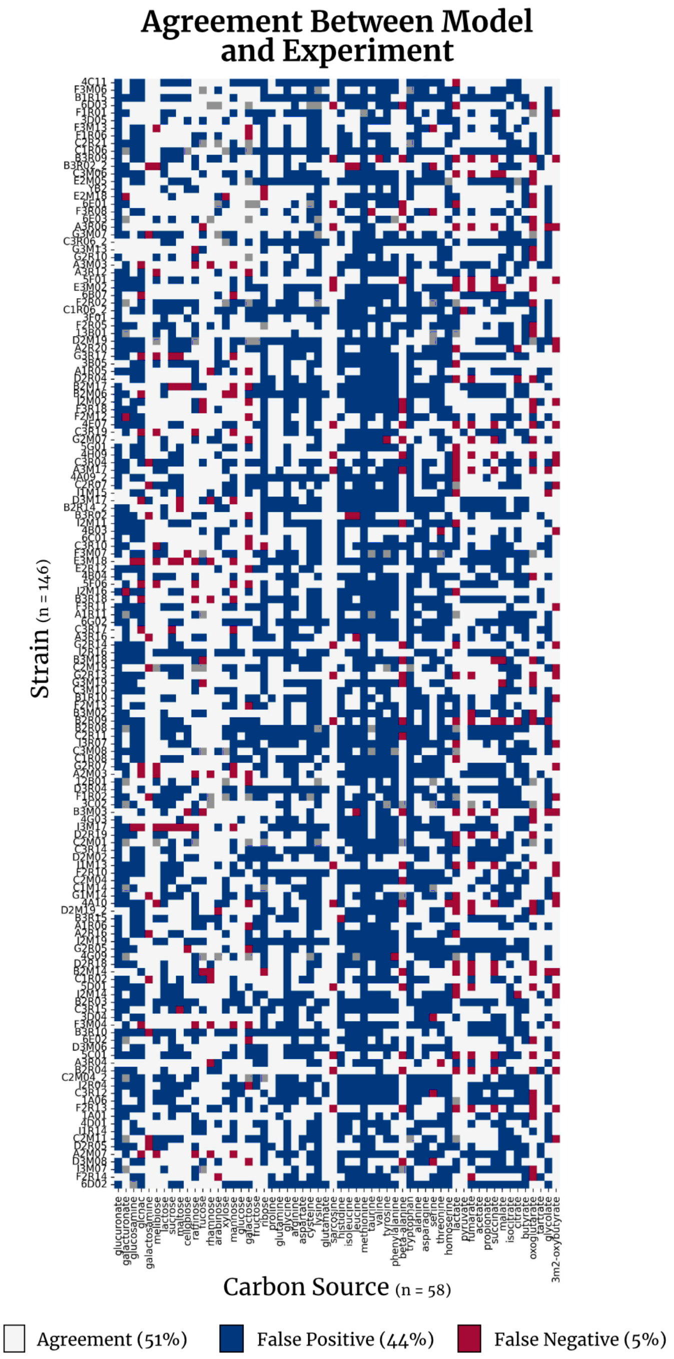


**Supplemental Figure S3: Comparison of results between CarveMe model ensembles and experimental growth studies performed in Gralka et al.** Heatmap of agreement between CarveMe model ensemble reactions and experimental growth data for a collection of 146 strains grown on 58 different sole carbon sources. White squares indicate direct agreement between the models and data (i.e., model includes the exchange reaction and growth was observed or model does not include the exchange reaction and no growth was observed), blue squares indicate a false positive (model includes the exchange reaction, experimental data does not) and red squares indicate a false negative (experimental data predicts growth, model does not include the exchange reaction). Gray squares indicate that the presence of the compound in the model exchange reactions was variable (between the consensus thresholds for "present" and "absent").

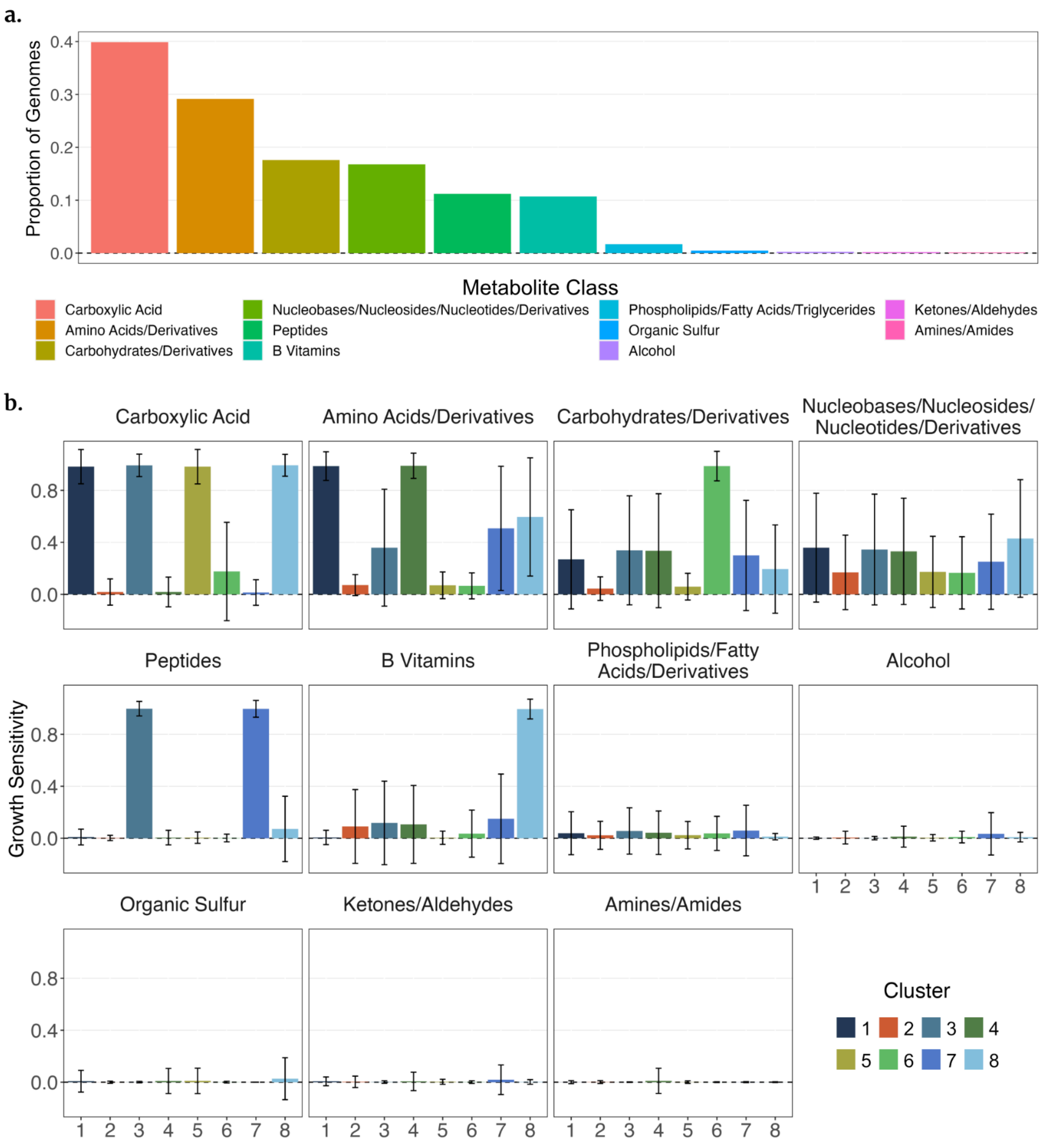


**Supplemental Figure S4: Relative growth sensitivities between SOM clusters.** (a) Ordered bar plot of the proportion of models across all clusters with substantial growth sensitivity to the reduction of each compound class (substantial is defined as >80% reduction in growth). (b) Bar plots of the relative mean growth sensitivity values for each of the 11 compound classes across the 8 SOM clusters. The error bars represent one standard deviation. Plot facets are ordered from the highest overall sensitivity (carboxylic acids) to the lowest (amines/amides).

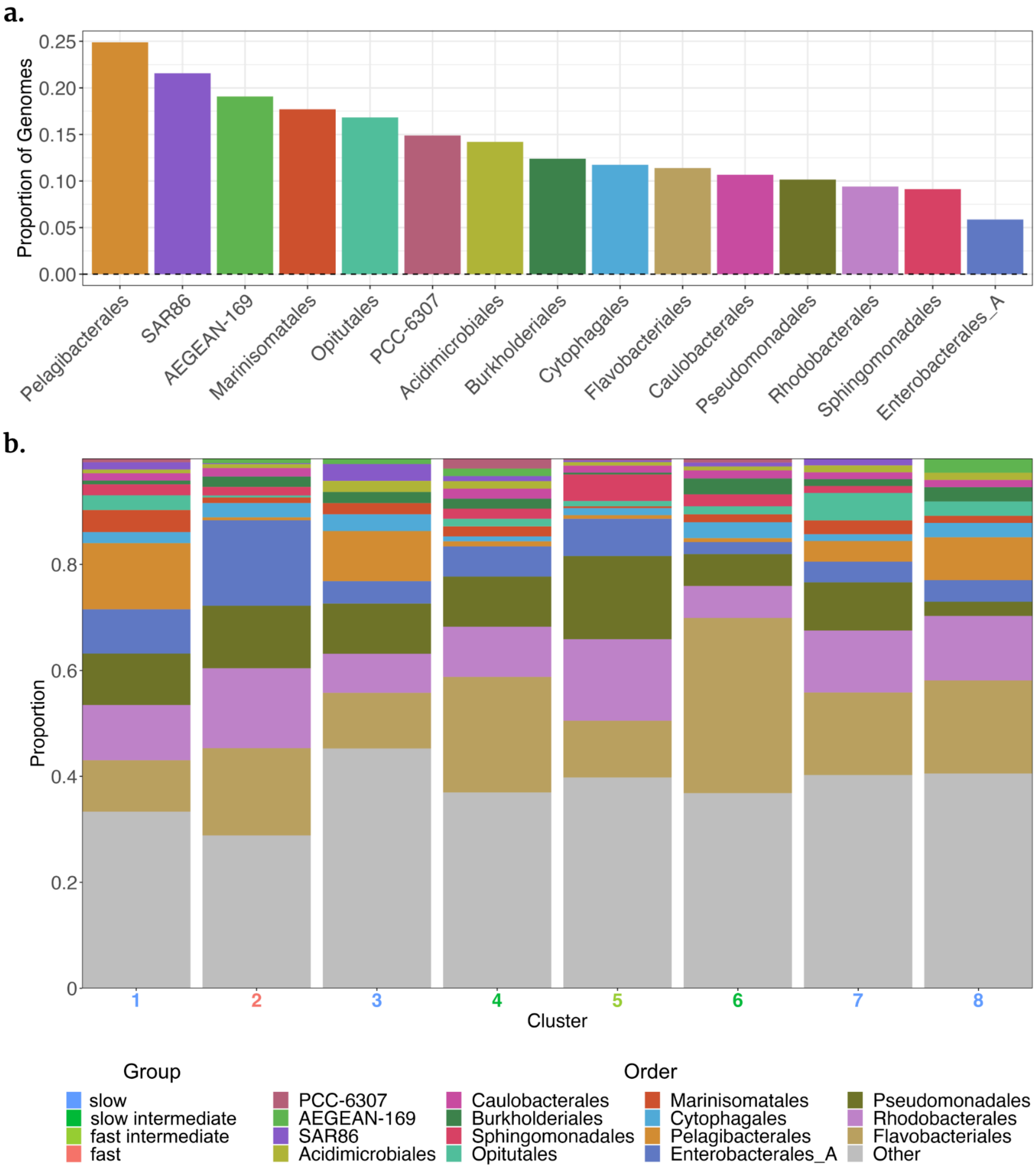


**Supplemental Figure S5: Taxonomy by cluster.** (a) Ordered bar plot of the proportion of models in each of the 15 most abundant orders with substantial growth sensitivity to the reduction of any compound class (substantial is defined as >80% reduction in growth). (b) Stacked bar plots of the relative abundances of the top 15 orders in each cluster.

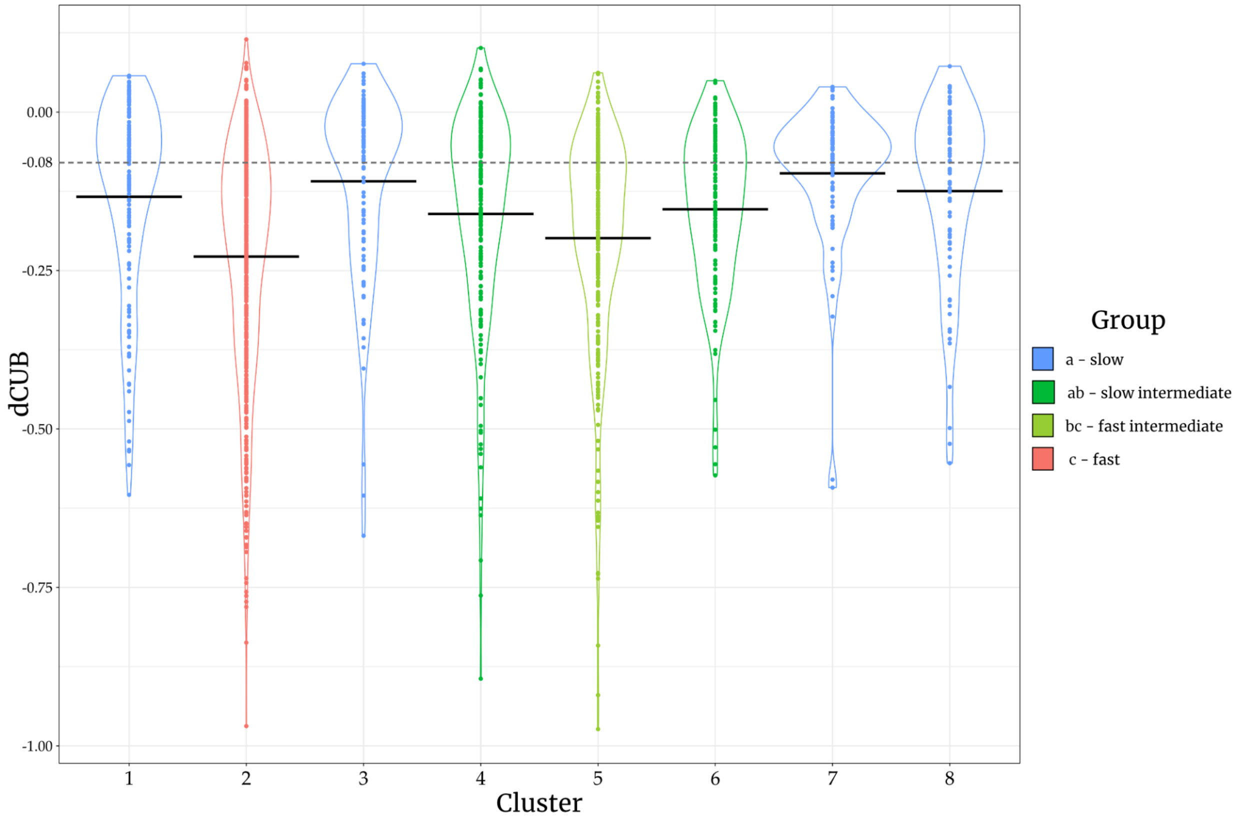


**Supplemental Figure S6: Codon usage bias (dCUB) by cluster.** The dCUB values fall into four statistically distinct groups designated with letters according to the key. Group *a* is the slow-growing group (Clusters 1, 3, 7 and 8) and statistically distinct from the other clusters. Group *c* (Cluster 2) is the fast-growing group and is distinct from all other clusters. Groups *ab and bc* represent our intermediate growers. Specifically, group *ab* (Clusters 4 and 6) are statistically distinct from fast-growing group *c* but not from slow-growing group *a*, whereas group *bc* (Cluster 5) is statistically distinct from group *a* but not from group *c*.

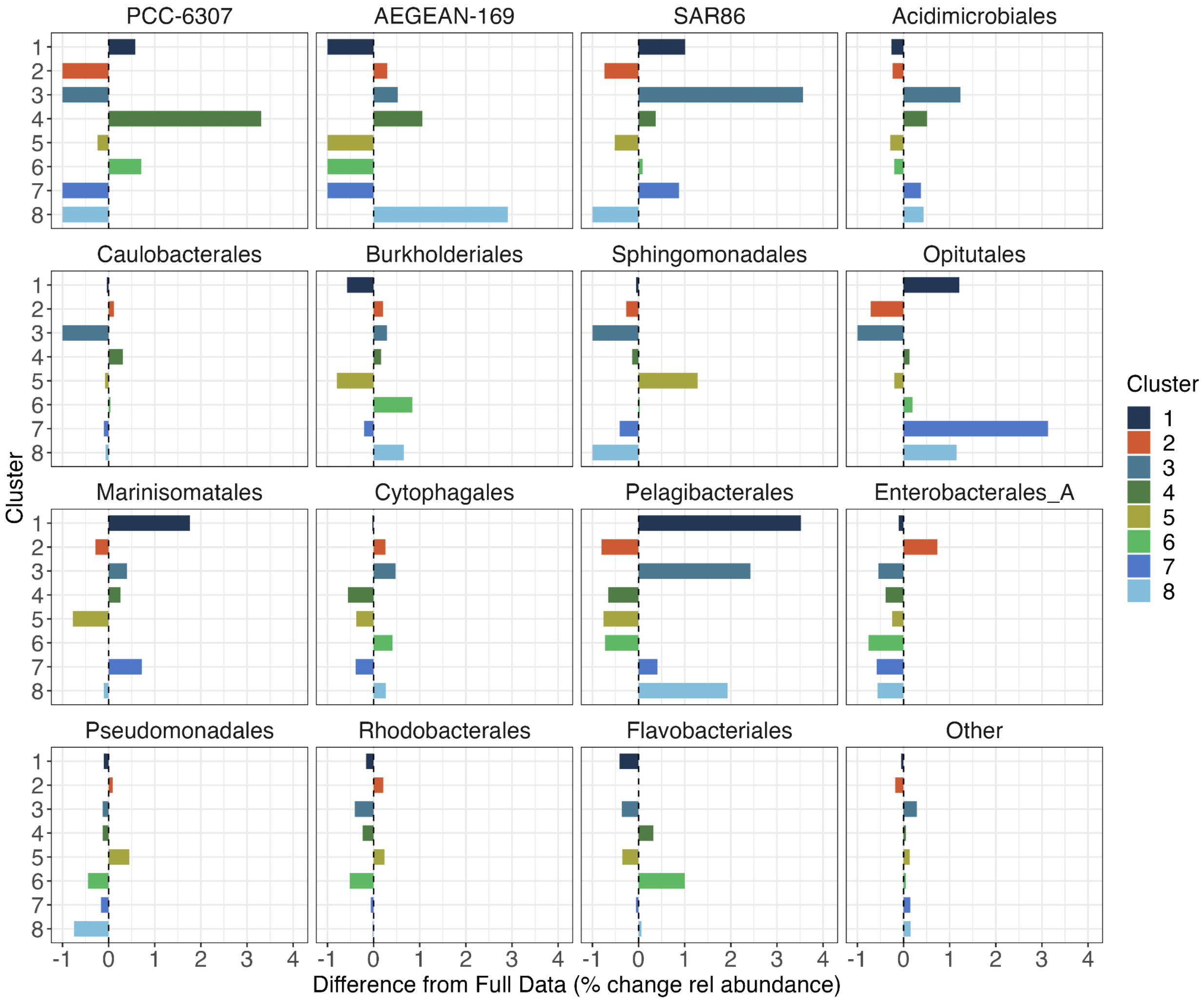


**Supplemental Figure S7: Taxonomic abundance enrichments of top 15 Orders by cluster.** Percentage enrichment in the relative abundance of the top 15 Orders (and Other) in each of the 8 SOM clusters compared to the relative abundances of each of these Orders in the full dataset.

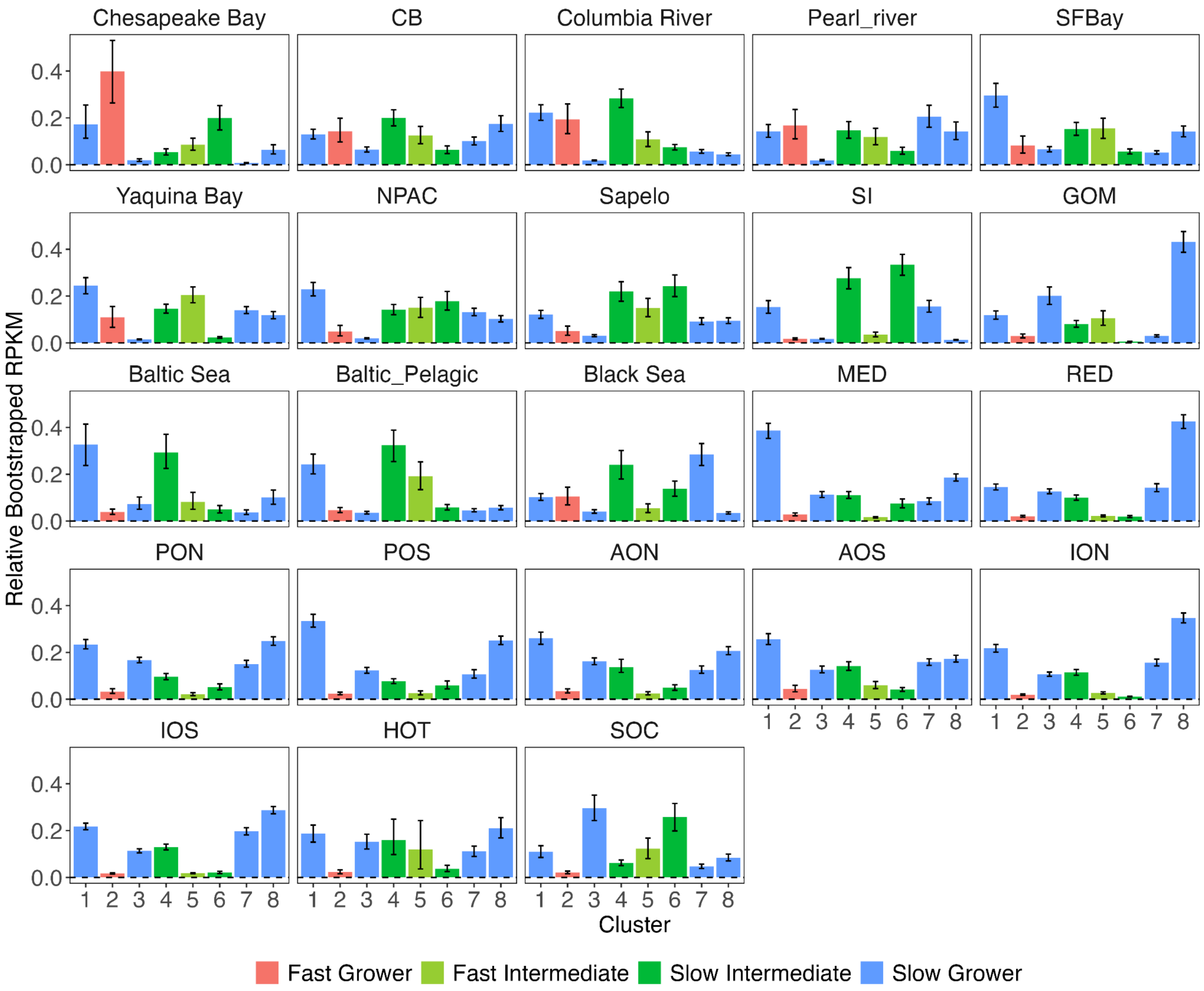


**Supplemental Figure S8: Relative abundances of SOM clusters by oceanographic region.** Sampling sites were grouped for bootstrapping according to the 23 oceanographic regions given in Table S3. For each region, bar plots of the average relative abundances of each of the 8 SOM clusters are shown. The relative abundances are calculated based on the bootstrap distributions of the raw RPKM values. The clusters are colored by their growth strategy (fast, fast-intermediate, slow-intermediate, and slow). Error bars represent the standard deviations of the bootstrapped distributions.

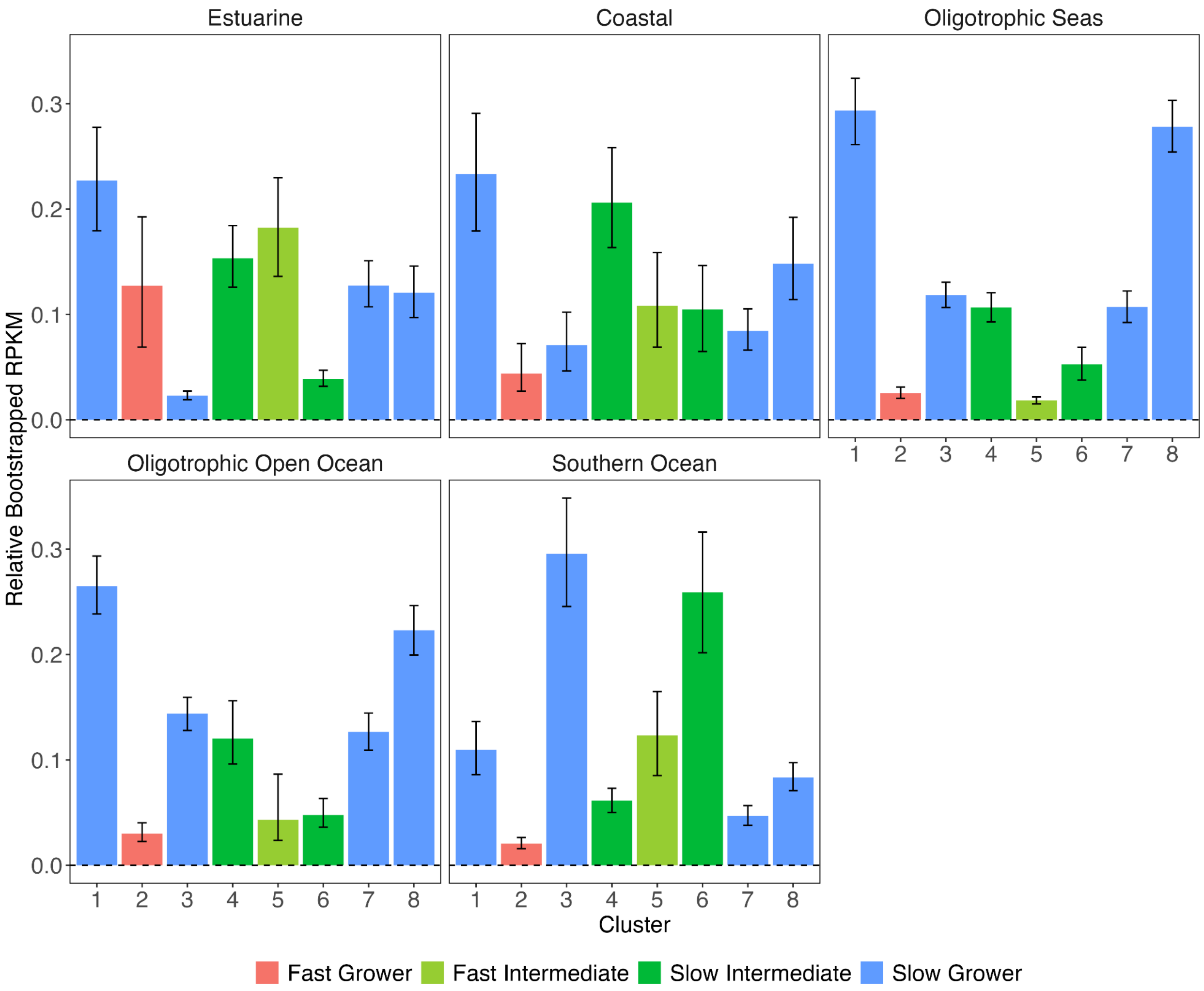


**Supplemental Figure S9: Relative abundances of SOM clusters by oceanographic category.** Sampling sites were grouped for bootstrapping according to the 5 oceanographic categories. Bar plots show the average relative abundances of each of the 8 SOM clusters in each category where the abundance is based on the bootstrap distributions of the raw RPKM values. The clusters are colored by their growth strategy (fast, fast-intermediate, slow-intermediate, and slow). Error bars represent the standard deviations of the bootstrapped distributions.

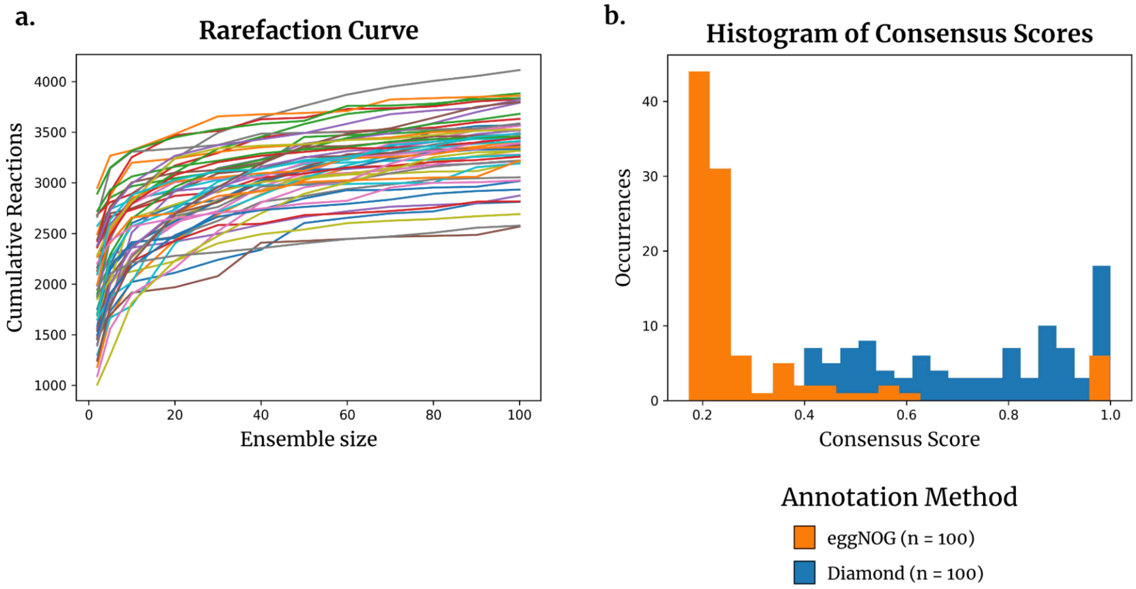


**Supplemental Figure S10: CarveMe run parameterizations.** (a) Rarefaction curve of the total number of unique reactions found in any model within an ensemble of models generated for a given genome. This curve was generated for ensemble sizes ranging from 2-100 models. At low ensemble sizes (e.g., in the range from 2-20) the model space rapidly identifies new unique reactions as more models are generated. The curves stabilize around ensemble sizes of 40-80 such that increasing the number of models in the ensemble does not add new reactions. (b) Histogram of the consensus scores for model ensembles when annotating reactions for CarveMe with eggNOG vs. the native Diamond (ensemble size = 60). Overall models generated with Diamond annotation produced significantly higher quality models than when eggNOG annotations were used for the same genomes.

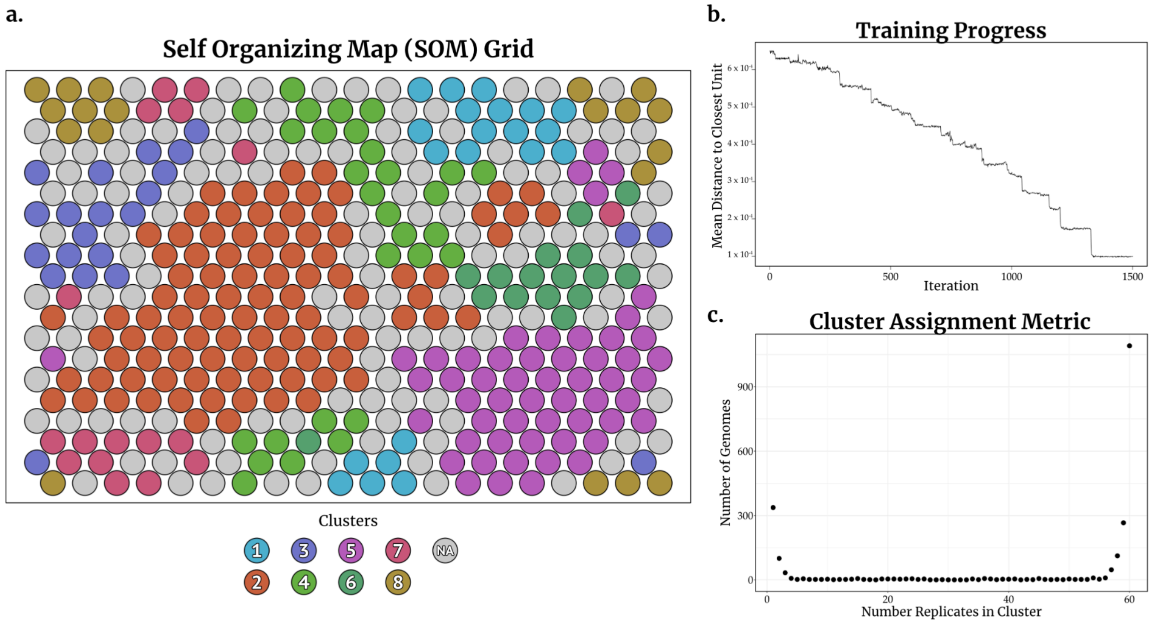


**Supplemental Figure S11: SOM metrics.** (a) The SOM grid is shown where each circle represents a grid point in the map (N=400). Grid points are colored by their assignment to the 8 defined SOM clusters. Grid points to which no genomes were assigned are colored gray to represent the absence of mapped data. It is important to note that the SOM uses a toroidal grid where the edges wrap around such that, for example, all nodes in Cluster 8 are in fact connected. (b) The training progress of the grid is shown for the duration of the map refinement process. (c) The number of models from each genome ensemble that were assigned to each SOM cluster, where a value of 60 denotes instances when all models from the ensemble were assigned to the same SOM cluster and a value of 0 denotes that no models from a specific ensemble were assigned to the cluster. Data is only shown for the 1,591 high consensus ensembles. The bimodal distribution of the data around 0 and 60 illustrates that all models from a given ensemble were almost always assigned to the same SOM cluster.

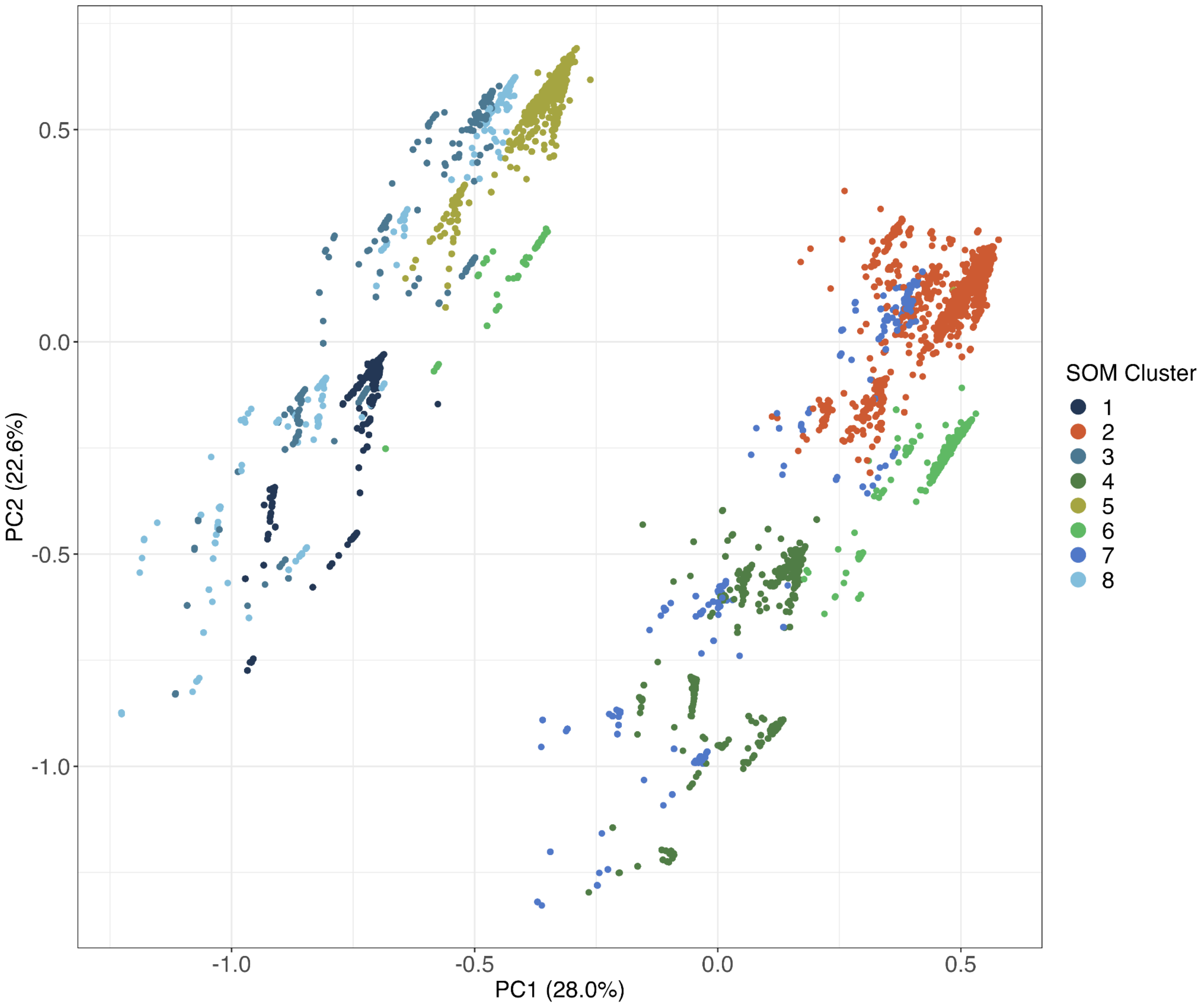


**Supplemental Figure S12: PCA plot of the growth sensitivity data.** The PCA captured 50.6% of the total variance on the first two principal component axes, and distinguished two major groups of data points, a slow growing and fast growing cluster. Of note, the estimates of maximum growth rate were not included in this clustering. The points in the PCA are colored by SOM cluster assignment to illustrate that both approaches identified similar clustering of the data but that the SOM method was able to better differentiate between the 8 clusters.

**Supplemental Table S1: All data for 1,591 high consensus genomes.** This table provides the unique identifiers for the genomes used in the SOM analysis as well as information on the SOM cluster they were assigned to, their specific value of the dCUB growth proxy, taxonomic information (order, genus, and species), as well as the raw growth sensitivity values computed for the 11 compound classes clustered in this study.

**Supplemental Table S2: Metabolite classification information.** This table provides information on the 456 compounds that were manually classified for this study including their names in plain English, the compound class they were assigned to, and the name of the corresponding external reaction in the CarveMe universal model.

**Supplemental Table S3: Biogeographical distribution of SOM clusters by oceanographic region.** This table provides the RPKM information for the 23 regions defined by Lanclos et al. (2023) including the full name of each region, the identifier for each region, the oceanographic category each region was assigned to, the number of stations assigned to each region, and the relative abundance of the 8 SOM clusters based on the bootstrapped RPKM values.

**Supplemental Table S4: Biogeographical distribution of SOM clusters by oceanographic category.** This table provides the RPKM information for the 5 oceanographic categories defined in this study including the number of stations assigned to each category and the RPKM relative abundance information for each category.

**Supplemental Table S5: Matrix of significance values for all pairs of SOM clusters based on dCUB distributions.** This table provides the p-values for all paired comparisons of the distributions of growth rates (estimated using dCUB) for our 8 SOM clusters.

**Supplemental Table S6: Unifrac distances for all paired comparisons of the SOM clusters.** This table provides the Unifrac distances for all paired comparisons of the genomes in SOM clusters. We report the Unifrac distances when comparing across the full phylogenetic tree as well as for subtrees of only the genomes in each of our 15 major taxonomic orders.

**Supplementary Information**

### S1 CarveMe Validation

We validated the CarveMe models and predictions of growth sensitivity to specific compounds by comparing our model predictions to an extensive experimental dataset where 176 marine bacterial strains were tested for their ability to grow on 118 substrates as a sole carbon source[1]. In 51.0% of cases, the CarveMe ensembles directly agreed with the growth predictions – either the models contained the exchange reaction and experimental data confirmed growth, or the models did not include the exchange reaction and the experiments showed no growth. Here, we required the exchange reaction to be present in >80% of the ensemble members to be considered present in the model predictions for a given strain. This agreement between the models and data was similar to the results reported by Gralka et al. of 58.0% agreement between CarveMe models and experimental data. We found that in just 4.6% of cases CarveMe predicted that a specific exchange reaction was not present in our CarveMe ensembles despite experimental evidence that that strain could grow on the specific substrate as a sole carbon source. Gralka et al found that these same instances occurred much more often in 16.1% of cases. Gralka et al's methodology differed from ours in several key ways including the fact that they did not use model ensembles nor exclude strains that generated poor quality models. Our low false negative rate offers strong evidence that the predictions of exchange reactions from CarveMe are robust when generated following our methodology.

In 44.4% of cases, our CarveMe ensembles predicted that the exchange reaction was present in the models despite experimental evidence demonstrating no growth on that substrate as the sole carbon source. The models generated by Gralka et al found these same instances in 26.0% of cases. This high false positive rate is not necessarily inconsistent with high quality CarveMe models. The experimental growth data from Gralka et al.[1] used each tested substrate as the sole carbon source. However, for this comparison, we only examined whether the exchange reaction was present in the model, not whether the model was able to grow on the substrate as a sole carbon source. For some compounds that are showing up as false positives, it is likely that the organisms could use the substrate but not as a sole carbon source. Amino acids are a good example of compounds that could be used in this way – an organism might have the ability to take up an amino acid from the environment which would be energetically favorable to synthesizing de novo however they might not be able to grow on that amino acid as the sole carbon source. Indeed, the false positive rate for amino acids in the Gralka et al dataset is higher than for the rest of the compounds (55.2%). When we looked at the carbohydrate compounds, which are more often used as primary carbon sources, the false positive rate decreased to 36.2%. Similarly, when we looked at only the carboxylic acids, the false positive rate decreased to 34.8%. This analysis suggests that the high false positive rates are in large part because the experiments and models are assessing different aspects of microbial metabolism – i.e. we are not comparing like to like. We also conducted a series of growth sensitivity tests on the models generated from the Gralka dataset which are described below (Supplement S3.2). These tests again are not a perfect comparison to the experimental design and so serve less as a validation and more as a point of comparison. Overall, these results provide support that the models generated by the CarveMe automated pipeline can be used to provide robust hypotheses about the metabolic capabilities of organisms.

### S2 SOMs clustering

The model growth sensitivity analysis generated 1,050,060 data points (1,591 genomes x 60 models x 11 compound class tests). To analyze this large dataset and identify overarching patterns we employed Self Organizing Maps (SOMs), a type of dimensional reduction method. In addition to SOMs, we analyzed the data using other traditional dimensional reduction approaches such as PCA, and clustering methods such as direct hierarchical clustering. The PCA was able to distinguish two broad clusters within the dataset with PC1 explaining 28.0% of the total variance and PC2 explaining 22.6% of the total variance (Supplemental Figure S12). However, PCA did not allow us to further differentiate within these groups while the SOM clustering provided clearly differentiated clusters. Direct hierarchical clustering was computationally infeasible on the full dataset, but was able to identify some patterns when the data values were averaged by ensemble.

We used a hexagonal, toroidal grid configuration to build the SOM in order to avoid the development of edge effects where the majority of data ends up in the corners of the map. Training a toroidal map connects these edges together fluidly to negate these edge effects (e.g., the top and bottom of the map are treated as adjacent). Adjusting for potential edge effects was necessary because, although the growth sensitivity results are continuous on a scale from 0-1, we observed bimodal distributions for most of the sensitivity tests where either a model was sensitive to the removal of a substrate (value between 0.8 and 1) or was insensitive to the removal (value between 0 and 0.2) (Supplemental Figure S2). Due to this bimodality, we also adjusted the learning rate vector from the default (0.05, 0.01) down to (0.025, 0.01). This reduced the learning rate for the map in early iterations to produce a smoother set of map values and combat the effect of bimodal data to tend towards extremes. We tested the number of iterations needed to achieve small quantization error values (discussed further below) (Supplemental Figure S11b) and increased the run length to 1500 iterations from the default setting of 100 iterations, in part due to the reduction in the initial learning rate which slows down convergence.

We also conducted a sensitivity analysis on the size of the map from 5-by-5 to 100-by-100. Determining map size for SOM clustering is an open problem, and there are no definitive theoretical bases for defining a “correct” size based on the size and characteristics of the input data[2–4]. Heuristic rules of thumb have been suggested[5] as well as field-specific guidelines[3]. However, the guidelines are not generalizable and remain dependent on data characteristics. Metrics of error have been proposed[6,7] to identify if a given set of map parameters generates a SOM with an appropriate level of resolution (quantization error) or topology preservation (topographic error). We computed quantization error which measures the resolution of a SOM by looking at the difference between a data vector and its best mapping unit (BMU). The final map size is a balance between a map that is large enough to fully differentiate all of the patterns in the dataset and one that does not contain too many unassigned nodes. We chose a 20-by-20 grid size such that the map had sufficient space to distribute the variability in the data without overfitting the map. Quantization error decreases with increasing map size so map size cannot be optimized to a global error minimum but only a local minimum[3]. Borders of unassigned nodes between clustered mapping units are one way that a sufficiently large map size is determined. The final map had 126 unassigned nodes (31.5%).

To identify clusters amongst the SOM nodes, we used k-means clustering. We also tried hierarchical clustering but elected to use k-means because it consistently performed better based on intra-cluster distances. The number of clusters (eight) was chosen to minimize the intra-cluster difference while not overfitting the map. The final map is shown in Supplemental Figure S11a colored by cluster. When visualized in two dimensional space this toroidal map appears as roughly square but the top and bottom, and left and right, sides of the map are continuously connected.

### S3 SOM cluster analyses

We performed a variety of analyses with the SOM clusters to identify patterns within the growth sensitivity profile data and between this data and other independent data measures. For each SOM cluster, we determined the set of compound classes that resulted in reduced growth when limited (hereafter called the growth sensitivity profile). We then calculated the genome estimated maximum growth rates for each cluster using codon usage bias (dCUB) and assessed the taxonomic composition of each cluster. For these analyses, we used the dCUB threshold value of $dCUB =- 0.08$ from[8,9] to differentiate between fast growing (more negative dCUB values) and slow growing (more positive dCUB value) organisms.

#### S3.1 dCUB growth estimates

We tested whether there were statistical differences in genome estimated maximum growth rates for the eight SOM clusters. Based on the results of our Tukey tests, we identified four statistically significant groups (Supplemental Figure S6). It is important to highlight that dCUB values were not included in building the SOM map and so any differences in dCUB values between clusters are emergent rather than prescribed. Genomes in Cluster 2 were predicted to be significantly faster growers (Supplemental Table S5) than all clusters except Cluster 5 based on codon usage bias (average dCUB of -0.228), while Clusters 1, 3, 7, and 8 all had significantly slower predicted growth rates (average dCUB ranging from -0.097 to -0.134). SOM Cluster 5 was statistically distinct from the slow growing clusters (average dCUB of -0.199) but overlapped with the fast growing cluster. Intermediate growth Clusters 4 and 6 were significantly slower than fast growing Cluster 2 (average dCUB of -0.161 and -0.153, respectively) but overlapped with the slow growing clusters. This suggests that we identified two distinct types of intermediate growers in our dataset – a fast intermediate growth cluster and slow intermediate growth clusters.

To further distinguish the intermediate growth clusters (Clusters 4-6) from one another, we calculated the fraction of genomes within each cluster with dCUB values below the -0.08 dCUB threshold (fraction of fast growth genomes). We found that 66.2% and 62.1% of genomes in Clusters 4 and 6 were 'fast growers' while 73.2% of genomes in Cluster 5 were 'fast growers'. For comparison, 78.0% of genomes in Cluster 2 were 'fast growers' while on average 45.4% of genomes in the slow growing clusters (range 40.0-50.0%) were 'fast growers' (Supplemental Figure S6). We posit that genomes in the 'intermediate growth' clusters could belong to some intermediary lifestyle phenotype(s) between copiotrophs and oligotrophs. Furthermore, our results hypothesize the existence of more than one of these intermediary phenotypes. Overall, our results suggest that there are several growth strategies for the 'intermediate' lifestyle and multiple growth strategies for the slow growing oligotrophs. In contrast, we identified only one growth strategy for the fast growing genomes. Our inability to differentiate multiple fast growing groups might be a bias in the model formulation as we are only able to differentiate the genomes based on the sensitivity to the 11 compound classes tested.

#### S3.2 Growth Sensitivity Profiles

We observed substantial differences in the metabolic strategies between all eight of our SOM clusters (Supplemental Figure S4b). Generally, we found that our SOM clusters fell into one of three distinct metabolic strategies, and that these strategies aligned with the four statistically different growth strategies identified above. The slow growing clusters demonstrated high growth sensitivity to two or more compound classes. For example, slow growing Cluster 3 demonstrated substantial sensitivity to both carboxylic acids and peptides. All four of the slow growth clusters had unique pairings of growth sensitive compound classes. By contrast, our fast growing Cluster 2 demonstrated low to zero sensitivity to any compound classes. Our three intermediate growth clusters each demonstrated a single compound class sensitivity, with our two slower intermediate growth Clusters 4 and 6 demonstrating sensitivity to amino acids and carbohydrates, respectively, while fast intermediate growth Cluster 5 showed sensitivity to carboxylic acids. Overall, carboxylic acids (4 clusters), amino acids/derivatives (2 clusters), and peptides (2 clusters) were the compound classes that caused the most significantly high sensitivities amongst our eight SOM clusters.

Overall, we observed that sugar and acid compounds were the primary drivers of our SOM clustering, which aligns with the proposed classification from the Gralka et al study through their sugar acid preference (SAP) metric. While our sensitivity test is not homologous to their SAP metric, we decided to compute growth sensitivities for the strains used in the Gralka study to examine the differences in sugar/acid sensitivity for organisms with a defined preference for sugars or acids based on their SAP. The sugar compounds from the Gralka et al study were primarily classified as carbohydrates with some carboxylic acids and nucleobases/nucleotides/nucleosides, while the acid compounds were primarily classified as amino acids, peptides, and carboxylic acids. Using the sugar/acid preference metric developed by Gralka et al, we grouped the 146 strains into sugar specialists (N=77) and acid specialists (N=69). For each group, we then assessed the growth sensitivity to removing carbohydrates, amino acids, or carboxylic acids (Supplemental Figure S3b). Reducing the availability of carbohydrates resulted in substantial growth reduction (growth sensitivity values of 0.8 or greater) in 14.3% of the sugar specialist strains, while there were no acid specialists that suffered substantial growth reduction. Given that the SAP is a continuum between sugar and acid preference such that some of the strains were only weak sugar preferrers, we also used a very liberal definition of growth sensitivity of 0.2 (i.e. a 20% reduction in growth due to limitation of the compound). With the lower threshold, 16.9% of the sugar specialists showed a response while we still observed 0% of the acid specialists with a response.

Reducing the availability of amino acids resulted in substantial growth limitation (0.8 threshold) in 15.9% of acid specialist strains compared to 11.7% of sugar specialist strains . We hypothesize that part of the challenge with the assessment of acid preference using amino acids is that amino acids are so central to growth that models often have multiple strategies for synthesizing amino acids when they are not available from the environment. When we assessed the growth reduction to the removal of carboxylic acids, we saw that 36.2% of acid specialist models were sensitive to the removal of carboxylic acids while 18.2% of sugar specialists were sensitive (0.8 threshold). With the liberal threshold of 0.2, we observed 42.0% sensitivity to the removal of carboxylic acids by acid specialists relative to 19.5% for sugar specialists.

It is important to note that the Gralka et al sugar/acid preference index is not an auxotrophy for these compounds and the preference metric is a continuum from -1 to 1. In contrast, our sensitivity test was not designed to determine the relative preference of sugar to acid but rather the response to removing the substrate. Thus we would not necessarily expect perfect agreement as these are two different tests. But the overall consistency in response is encouraging.

#### S3.3 Cluster phylogeny

To confirm that our SOM clusters were identifying *de novo* guilds of metabolic similarity rather than simply recapturing known phylogenetic groups, we performed phylogenetic and taxonomic analyses of the genomes assigned to each cluster. Qualitative analyses of taxonomic relative abundances within our SOM clusters showed that clusters were composed of many different taxonomic orders, and many orders were represented across multiple clusters, albeit at varying relative abundances (Supplemental Figures S5, S7). Thus, the metabolic niches captured by our SOM clusters could not be identified directly from taxonomy alone. To further investigate possible differences between the SOM cluster phylogenies, we computed the UniFrac distances between clusters to quantify their phylogenetic relatedness. We found that all pairwise comparisons of our clusters resulted in relatively high Unifrac distances (values ranging from 0.767-0.847 on a $[0, 1]$ scale). These values are reported in Supplemental Table S6. Typically, UniFrac distances in this range would suggest that the SOM clusters are taxonomically distinct from one another. Since our qualitative analyses demonstrated that our SOM clusters were not simply recapitulating taxonomy at the order level, we concluded that the UniFrac distances indicated phylogenetic distinction based on each cluster having unique members within each order. These high UniFrac distances were likely attributable in large part to the demonstrably different relative abundances of the major taxonomic orders between SOM clusters (Supplemental Figure S7). Another possibility was that the UniFrac scores reflected unique taxonomy below the order level (e.g., genus or species).

We tested this hypothesis by restricting our UniFrac analysis to individual orders and then re-computing the distances between SOM clusters (Supplemental Table S6). We completed all pairwise comparisons for the 8 SOM clusters across each of the top 15 orders. In general, the UniFrac distances between SOM clusters were lower when looking within an order than when looking at the entire phylogeny, which suggests that there are greater phylogenetic differences between SOM cluster assignments at the order level than there are at the sub-order level. This further supports the notion that the majority of these phylogenetic differences are likely driven by differences in the relative abundance of these orders in each SOM cluster. The average UniFrac distances between SOM clusters within the top 15 orders spanned a wide range; the *Flavobacteriales* were the most phylogenetically different between clusters (with the highest average UniFrac distance of 0.795) while *Caulobacterales,* AEGEAN-169, and PCC-6307 were the most similar between clusters (with average UniFrac distances of 0.293 0.240, and 0.180 respectively). This suggests that SOM cluster membership formed distinct clades at the sub-order level for the *Flavobacteriales*, *Psuedomonadales*, *Rhodobacterales*, and *Cytophagales*, all of which had UniFrac distances larger than 0.5 (Supplemental Table S6). For the remaining orders, the low average UniFrac distances could indicate that either the organisms in the clusters had a high level of metabolic flexibility or that metabolic differences could be accounted for by strain level variation in metabolic capability.

We also found that Cluster 2 had the highest average UniFrac distance to the other SOM clusters in 7 of the 15 orders, suggesting that it was the most phylogenetically distinct cluster. Cluster 2 was also our fast growing cluster and the only cluster without any compound sensitivities, which suggests that organisms occupying this metabolic niche were more distinct from organisms in other niches.

Overall these results demonstrate that, while in some cases there is a relationship between cluster membership and phylogeny, the clusters cannot be explained through phylogeny alone. They also emphasize the need for defining metabolic niches predicated directly from metabolism - like the one presented in this study - rather than defining metabolic niches directly from phylogeny.

#### S3.4 Geographic distribution

The SOM clusters showed unique biogeographic patterns based on metagenomic recruitment (Supplemental Figure S9). We found that slow growing Cluster 1 was the most numerically dominant of our eight SOM clusters with a mean relative abundance of 22.6%, while fast growing Cluster 2 was the least abundant with a mean relative abundance of 4.95%. Our fast intermediate growth Cluster 5 was second lowest in average relative abundance at 9.52%. Cluster 1 was found to be highly abundant relative to the other seven clusters in every oceanographic category except for the Southern Ocean where it was the fourth most abundant.

When we looked within specific oceanographic categories, further patterns emerged. While overall Cluster 2 and 5 had the lowest abundances, these two clusters were significantly more abundant at estuarine stations, reaching mean relative abundances of 12.7% for Cluster 2 (fifth highest abundance in estuarine regions amongst all clusters) and 18.2% for Cluster 5 (second highest abundance in estuarine regions among all clusters). The estuarine stations were highly diverse, with six of the eight SOM clusters present at relatively high abundances at statistically identical levels (Clusters 3 and 6 were significantly rarer than the other clusters in the estuarine samples). The relative evenness of the majority of the SOM clusters at estuarine stations suggests that the microbial community is being supplied with a diverse set of compounds at concentrations sufficiently high to support the metabolic requirements of a diverse group of organisms. This type of flexible environment with diverse compound availability favors more balanced metabolic strategies and higher maximum growth rates of the faster growing clusters which is consistent with the higher abundances of these clusters at estuarine stations.

The abundance of the clusters in coastal stations was more evenly distributed than in estuaries. However, unlike in the estuaries, the coastal stations were primarily dominated by the slow growing Cluster 1 (23.3% mean relative abundance) and slow intermediate growth Cluster 4 (20.6% mean relative abundance). Similarly to the estuarine stations, the overall taxonomic evenness suggests that the microbial community at coastal sites are also being supplied a diverse set of compounds at sufficiently high concentrations to support the growth of diverse metabolic strategies. In particular, since the two most dominant clusters were sensitive to amino acids, their abundance in coastal locations suggests that those stations had high enough concentrations of amino acids to sustain large total biomasses of these organisms. One possible cause for the greater evenness of our eight SOM clusters in coastal sites compared to estuaries is the effect of varying salinity on community membership at estuarine stations. Variations in the salinity levels could be an external environmental factor limiting the growth of organisms that could otherwise grow effectively, thereby suppressing the abundance of the SOM clusters they were assigned to.

The remaining three categories - oligotrophic seas, oligotrophic open oceans, and the southern ocean - were dominated by just two of the eight SOM clusters. The oligotrophic seas and oligotrophic open oceans showed similar distributions of relative abundance and were both dominated by slow growing Clusters 1 and 8. The high abundance of slow growing organisms in these categories is consistent with these oceanographic regions being resource limited. In resource limited environments, organisms often cope with consistently low nutrient concentrations by specializing in growth on specific compounds resulting in an environment with rigid, defined niches[10]. Organisms typically specialize in growth on certain compounds by using transporters with greater affinity for these compounds, and/or streamlining their genomes to reduce internal nutrient requirements. This sort of rigid niche structure, and low nutrient availability, is unfavorable to fast growing, metabolically flexible organisms as they can be outcompeted for compound acquisition by specialists. Resource limited conditions thus favor compound sensitive, specialist organisms occupying defined niches for growth on their specific compounds[11]. The significantly lower abundances of slow growing Clusters 3 and 7 suggest then that the environmental niches that organisms in these clusters occupy are not present. Specifically, Clusters 3 and 7 are the only two clusters sensitive to peptides which suggests that stations in these oligotrophic categories might have particularly low concentrations of peptide compounds available. This is consistent with the fact that peptides are one of the most labile forms of DOM available to heterotrophic organisms and would be drawn down rapidly under resource limited conditions.

In contrast to the oligotrophic categories, the Southern Ocean category was actually dominated by slow growing Cluster 3 and slow intermediate growth Cluster 6. There were only three samples from the Southern Ocean in our dataset so it is difficult to conclude if these relative abundances are representative of the entire region. However, the Southern Ocean is distinct from the oligotrophic open oceans in that it has higher nutrients and sustains higher productivity when iron limitation is alleviated[12]. Thus, it is possible that increased nutrients and the subsequent increased productivity promotes the abundance of intermediate growth strategy organisms like those found in Cluster 6.

### References


1. Gralka, M., Pollak, S. & Cordero, O. X. Genome content predicts the carbon catabolic preferences of heterotrophic bacteria. *Nat Microbiol* **8,** 1799–1808 (2023).
2. Park, Y.-S., Céréghino, R., Compin, A. & Lek, S. Applications of artificial neural networks for patterning and predicting aquatic insect species richness in running waters. *Ecol. Modell.* **160,** 265–280 (2003).
3. Céréghino, R. & Park, Y.-S. Review of the Self-Organizing Map (SOM) approach in water resources: Commentary. *Environmental Modelling & Software* **24,** 945–947 (2009).
4. Kalteh, A. M., Hjorth, P. & Berndtsson, R. Review of the self-organizing map (SOM) approach in water resources: Analysis, modelling and application. *Environmental Modelling & Software* **23,** 835–845 (2008).
5. Vesanto, J. *Neural Network Tool for Data Mining: SOM Toolbox*. (Helsinki University of Technology, 2000). at <https://citeseerx.ist.psu.edu/document?repid=rep1&type=pdf&doi=0e9bee375e885c4740ba0dab007167c485fa1e48>
6. Kiviluoto, K. Topology preservation in self-organizing maps. in *Proceedings of International Conference on Neural Networks (ICNN'96)* **1,** 294–299 vol.1 (IEEE, 1996).
7. Kohonen, T. The self-organizing map. *Neurocomputing* **21,** 1–6 (1998).
8. Weissman, J. L., Hou, S. & Fuhrman, J. A. Estimating maximal microbial growth rates from cultures, metagenomes, and single cells via codon usage patterns. *Proc. Natl. Acad. Sci. U. S. A.* **118,** (2021).
9. Weissman, J. L., Dimbo, E.-R. O., Krinos, A. I., Neely, C., Yagües, Y., Nolin, D., Hou, S., Laperriere, S., Caron, D. A., Tully, B., Alexander, H. & Fuhrman, J. A. Estimating global

variation in the maximum growth rates of eukaryotic microbes from cultures and metagenomes via codon usage patterns. *bioRxiv* 2021.10.15.464604 (2022). doi:10.1101/2021.10.15.464604

10. Gifford, S. M., Sharma, S., Booth, M. & Moran, M. A. Expression patterns reveal niche diversification in a marine microbial assemblage. *ISME J.* **7,** 281–298 (2013).
11. Sarmento, H., Morana, C. & Gasol, J. M. Bacterioplankton niche partitioning in the use of phytoplankton-derived dissolved organic carbon: quantity is more important than quality. *ISME J.* **10,** 2582–2592 (2016).
12. Venables, H. & Moore, C. M. Phytoplankton and light limitation in the Southern Ocean: Learning from high‐nutrient, high‐chlorophyll areas. *J. Geophys. Res.* **115,** (2010).
