## Supplemental Table S1 for "Emergent Metabolic Niches for Marine Heterotrophs"

| Genome | Cluster | dCUB | Order | Genus | Species | Alcohol | Amines/Amides | Amino Acids/Deri | B Vitamins | Carbohydrates/D | Carboxylic Acid | Ketones/Aldehyd | Nucleobases/Nu | Organic Sulfur | Peptides | Phospholipids/Fatty Acids/Triglycerides |
|---|---|---|---|---|---|---|---|---|---|---|---|---|---|---|---|---|
| BATS_SAMN07137086_METAG_HNJEHGAI | 1 | -0.04877634133 | Marinisomatales | GCA-002701945 | GCA-002701945 | 0 | 0 | 1 | 0.03333333333 | 8.94E-05 | 1 | 0 | 1 | 0 | 8.24E-05 | 0 |
| BATS_SAMN07137114_METAG_AICJLGGM | 2 | -0.1099505454 | Pseudomonadale | UBA9145 | UBA9145 sp019 | 0 | 0 | 0.03417139413 | 1.10E-04 | 0.003221708099 | 6.81E-06 | 0 | 0.0693100433 | 0 | 0.001833638321 | 0 |
| BATS_SAMN07137114_METAG_IAJLFLPG | 6 | -0.05430496608 | SAR202 | UBA11996 | UBA11996 sp002 | 0 | 0 | 0.032860455 | 5.00E-07 | 0.9999966669 | 0.9999966669 | 0 | 0 | 3.33E-08 | 0.01666666667 | 0 |
| BGEO_SAMN07136490_METAG_MFAPHJGH | 4 | -0.1295523918 | Flavobacteriales | SCGC-AAA160-F | SCGC-AAA160-F | 0 | 0 | 1 | 9.59E-05 | 0.005451292335 | 0 | 0.008684732576 | 1 | 0 | 0.001579564457 | 0.2093414724 |
| BGEO_SAMN07136496_METAG_NAJLEHLF | 7 | -0.09401928318 | Pseudomonadale | UBA4421 | UBA4421 sp0027 | 0 | 0 | 1 | 0.01685642332 | 0.1090120637 | 0.01667198198 | 0 | 0.06424157268 | 0 | 1 | 0.06615096859 |
| BGEO_SAMN07136507_METAG_JCDJIDOL | 7 | -0.01682046171 | Flavobacteriales | | | 0 | 1.86E-08 | 1.000000009 | 0.01674412322 | 1.000000009 | 0.002840831481 | 0 | 0.07321987906 | 0 | 0.9669912666 | 0.03439797437 |
| BGEO_SAMN07136523_METAG_NMOKDPNF | 3 | -0.2738337238 | Chitinophagales | | | 0 | 0 | 1.00000024 | 1.64E-04 | 1.00000024 | 1.00000024 | 0 | 1.00000024 | 0 | 1.00000024 | 0 |
| BGEO_SAMN07136562_METAG_LEGHEPPA | 3 | -0.04424182638 | Verrucomicrobial | EC70 | EC70 sp0139111 | 0 | 0 | 0.04965856207 | 2.15E-04 | 1.00000023 | 1.00000023 | 0 | 0.9950640987 | 0 | 1.00000023 | 0.07195522749 |
| BGEO_SAMN07136581_METAG_FOFJEIIN | 2 | -0.02395335436 | Arenicellales | REDSEA-S09-B1 | REDSEA-S09-B1 | 0 | 0 | 0.01761095967 | 1.61E-04 | 0.07585243218 | 5.28E-06 | 0 | 0.05030721276 | 0 | 0.001590762795 | 0 |
| BGEO_SAMN07136617_METAG_FKHCGNDB | 5 | -0.07105547837 | Rhizobiales | Methylobacterium | | 0 | 0 | 0.02281652429 | 0.01667072448 | 0.01681207345 | 1.000000001 | 0 | 0.3814991193 | 0 | 4.01E-05 | 0 |
| BGEO_SAMN07136635_METAG_NPBPCJHB | 1 | -0.04920494888 | Woeseiales | SP4260 | SP4260 sp00272 | 0 | 0 | 0.9999998046 | 2.81E-04 | 0.9999998046 | 0.9999998046 | 0 | 0.9994275998 | 0 | 8.07E-04 | 0.9999998046 |
| BGEO_SAMN07136637_METAG_DFMFBHGA | 1 | 0.01135336538 | Actinomarinales | Actinomarina | Actinomarina sp0 | 0 | 0 | 1.000000377 | 1.06E-04 | 1.000000377 | 1.000000377 | 0 | 1.000000377 | 7.28E-04 | 7.53E-04 | 0 |
| BGEO_SAMN07136638_METAG_DHFJAKOF | 4 | -0.00643303841 | Flavobacteriales | MED-G13 | MED-G13 sp002 | 0 | 0 | 1.000000304 | 1.91E-04 | 1.000000304 | 0.003696918273 | 0 | 1.000000304 | 0 | 0 | 0 |
| BGEO_SAMN07136638_METAG_PDOFCFGC | 4 | -0.09306977202 | Balneolales | UBA1275 | UBA1275 sp9025 | 0 | 0 | 0.9999997247 | 1.60E-04 | 0.9999997247 | 0.001368131779 | 0 | 0.9982588665 | 0 | 0.002809487512 | 0.9999997247 |
| BGEO_SAMN07136687_METAG_HGPPKMNF | 4 | -0.05582622399 | Pseudomonadales | | | 0 | 0 | 1 | 1.50E-06 | 0.1151572453 | 0.01951942449 | 0 | 0.06959003835 | 0 | 7.62E-05 | 0 |
| BGEO_SAMN07136710_METAG_BEJKEPKM | 4 | -0.2330213158 | Pseudomonadale | Alcanivorax | Alcanivorax sp00 | 0 | 0 | 1 | 5.59E-05 | 0.003334033901 | 0 | 0 | 0.03528997416 | 0 | 0.001219418596 | 0 |
| BGEO_SAMN07136710_METAG_IMMBNPOM | 4 | -0.2861531041 | Pseudomonadale | Alcanivorax | Alcanivorax sp90 | 0 | 0 | 1.000000306 | 8.61E-05 | 0.003153399462 | 0 | 0 | 0.03151927254 | 0 | 7.62E-04 | 0 |
| BGEO_SAMN07136710_METAG_NFNKEFAD | 5 | -0.5321797616 | Enterobacterales | Pseudoalteromor | Pseudoalteromor | 0 | 0 | 0.04792703897 | 1.24E-04 | 0.006438832544 | 1.000000037 | 0 | 0.113617999 | 0 | 5.36E-04 | 0 |
| BGEO_SAMN07136715_METAG_KOEDIAMF | 2 | -0.432063257 | Enterobacterales | Alteromonas | | 0 | 0 | 0.03652182934 | 0.999999993 | 1.39E-04 | 9.65E-05 | 0 | 0.2559639036 | 0 | 0.01576161052 | 0.1006527948 |
| BGEO_SAMN07136788_METAG_AFBPEAAE | 5 | -0.43785813 | Pseudomonadale | Pseudomonas_E | Pseudomonas_E | 0 | 0 | 0.1028471562 | 0.01666845362 | 0.3392766482 | 0.9500000345 | 0 | 0.06267698315 | 0 | 0 | 0.06231116404 |
| BGEO_SAMN07136900_METAG_NDDKEADO | 2 | -0.1907059959 | Burkholderiales | Pelomonas | | 0 | 0 | 0.02297375905 | 2.95E-05 | 0.09535096642 | 0.2166780853 | 0 | 0.07384664399 | 0 | 0.006116740199 | 2.33E-04 |
| BGEO_SAMN07136914_METAG_PMDBELBJ | 4 | -0.1706113371 | Flavobacteriales | UBA11663 | UBA11663 sp002 | 0 | 0 | 1.000000005 | 1.24E-04 | 1.000000005 | 0.01932574759 | 0 | 1.000000005 | 0 | 0.006333527778 | 0 |
| BGEO_SAMN07136920_METAG_MGPENMLJ | 4 | 0.02809935464 | Cytophagales | MED-G16 | | 0 | 0 | 0.9999997464 | 0.0167980096 | 0.9999997464 | 0.01592962415 | 0 | 0.9876151238 | 0 | 0.0127832963 | 0 |
| BGEO_SAMN07136937_METAG_ALPGGBFO | 4 | -0.08513223881 | Rhodobacterales | Planktomarina | Planktomarina sp | 0 | 0 | 0.9999998202 | 0.01666666339 | 0.06259921325 | 0.001162026083 | 0.002365449697 | 0.9999998202 | 1.48E-04 | 1.34E-05 | 0.03207421654 |
| BGEO_SAMN07136937_METAG_BMDGFMPD | 1 | -0.04097253704 | Pirellulales | UBA1268 | UBA1268 sp0028 | 0 | 0 | 0.9838674264 | 0.1334870418 | 2.65E-05 | 1.000000263 | 0 | 0.2113848232 | 0 | 0.01666667189 | 0.05957205789 |
| BGEO_SAMN07136957_METAG_NGKMKODO | 1 | -0.0432382834 | MED-G09 | MED-G09 | | 0.01072280667 | 0 | 1 | 0.01666666667 | 0.1340810576 | 0.9833333333 | 0 | 0.9840765305 | 5.32E-04 | 0.001302778345 | 2.01E-04 |
| BGEO_SAMN07136957_METAG_PDLAKDGF | 4 | -0.06709256812 | Flavobacteriales | UBA3537 | UBA3537 sp0026 | 0 | 0 | 1 | 0.01685006495 | 0.9833988054 | 7.65E-04 | 0 | 0.05985977835 | 0 | 0.003651582827 | 0 |
| BGEO_SAMN07136961_METAG_MADCCAJI | 7 | -0.05045655078 | Rhodobacterales | GCA-2697345 | | 0 | 0 | 0.0334657421 | 1 | 0.01232712421 | 1.71E-04 | 0.08288632583 | 0.05309216127 | 2.83E-05 | 1 | 0 |
| GORG_SAMEA6067721_SAGS_AH287C13 | 1 | 0.01828491831 | Pelagibacterales | Pelagibacter_A | Pelagibacter_A s | 0 | 0 | 1 | 3.16E-05 | 0.08132857463 | 1 | 0 | 0.06030884657 | 0 | 0.02236646375 | 0 |
| GORG_SAMEA6067750_SAGS_AH287E08 | 8 | 0.03125578999 | Pelagibacterales | Pelagibacter | Pelagibacter sp9 | 0 | 0 | 0.999999753 | 0.999999753 | 0.4750874809 | 0.999999753 | 0 | 0.9997136875 | 0 | 0.008422872882 | 0 |
| GORG_SAMEA6067751_SAGS_AH287E09 | 3 | 0.01601327585 | Pelagibacterales | Pelagibacter | Pelagibacter sp9 | 0 | 0.008527944493 | 1.000000005 | 3.99E-05 | 0.004076472406 | 1.000000005 | 0 | 0.210298438 | 1.01E-04 | 1.000000005 | 0 |
| GORG_SAMEA6067759_SAGS_AH287E19 | 1 | 0.02013654126 | Pelagibacterales | Pelagibacter_A | Pelagibacter_A s | 0 | 0 | 0.9999998074 | 0.01683369137 | 0.04916264474 | 0.9999998074 | 0.001479274822 | 0.9843954476 | 0 | 0.003173373045 | 0 |
| GORG_SAMEA6068347_SAGS_AH321N17 | 3 | -0.04741304274 | MED-G09 | AG-430-B22 | | 5.75E-07 | 0 | 1 | 1.47E-04 | 1 | 1 | 0.006685275878 | 1 | 0 | 1 | 0.07200280347 |
| GORG_SAMEA6068673_SAGS_AG365B19 | 1 | 0.02047884769 | Pelagibacterales | GCA-2704625 | | 0 | 0 | 0.9999997142 | 5.77E-05 | 0.4798320915 | 0.9999997142 | 5.72E-07 | 0.9999997142 | 0 | 0.2400057844 | 0 |
| GORG_SAMEA6069045_SAGS_AG359O02 | 1 | 0.0164270232 | Pelagibacterales | Pelagibacter | Pelagibacter sp0 | 0 | 0 | 1.000000266 | 1.15E-04 | 0.3379749998 | 1.000000266 | 0.002797799256 | 0.1345487579 | 9.03E-06 | 7.70E-04 | 0 |
| GORG_SAMEA6069416_SAGS_AG891A05 | 7 | 0.0374353844 | Pelagibacterales | Pelagibacter | Pelagibacter sp9 | 1.14E-05 | 0 | 1 | 5.42E-05 | 1 | 0.007488246246 | 0 | 0.07692915601 | 0 | 1 | 0.04096534819 |
| GORG_SAMEA6069450_SAGS_AG891C11 | 2 | 0.03748703107 | AEGEAN-169 | AG-337-I02 | AG-337-I02 sp90 | 0 | 0 | 0.09689367213 | 4.27E-05 | 0.006170156993 | 0.01462549281 | 0 | 1 | 0 | 0 | 0.03410524886 |
| GORG_SAMEA6069522_SAGS_AG891I06 | 6 | -0.05910983233 | Flavobacteriales | MS024-2A | MS024-2A sp902 | 0 | 0 | 0.02758479057 | 1.33E-04 | 1 | 1 | 0 | 0.2015179056 | 0 | 6.00E-04 | 0.0295228361 |
| GORG_SAMEA6069550_SAGS_AG891J21 | 4 | 0.02778469553 | Flavobacteriales | MED-G14 | MED-G14 sp002 | 0 | 0 | 1 | 1.70E-04 | 1 | 0 | 0 | 0.08315108227 | 0 | 0.002838476238 | 0 |
| GORG_SAMEA6069648_SAGS_AG892A22 | 1 | 0.01666555529 | Pelagibacterales | Pelagibacter | Pelagibacter sp9 | 0 | 0 | 0.9999997595 | 2.22E-04 | 0.2520011594 | 0.9999997595 | 0 | 0.06067784094 | 0 | 0.005196916871 | 0 |
| GORG_SAMEA6069733_SAGS_AG892G16 | 3 | 1.82E-04 | SAR86 | SAR86A | SAR86A sp9025 | 0 | 0 | 0.1393940097 | 1.000000197 | 0 | 1.000000197 | 0 | 0.06798878403 | 0 | 1.000000197 | 0 |
| GORG_SAMEA6069825_SAGS_AG892O07 | 4 | 0.007654860196 | Puniceispirillales | AAA536-G10 | AAA536-G10 sp9 | 7.49E-08 | 0 | 0.9999999962 | 0.01666748663 | 0.3784532762 | 0.07554385919 | 0 | 0.04917407193 | 0.002305574267 | 1.35E-05 | 0 |
| GORG_SAMEA6069857_SAGS_AG893A06 | 4 | 0.06851447565 | AEGEAN-169 | AG-337-I02 | | 7.52E-06 | 0 | 1 | 1.34E-04 | 0.03573106147 | 0.004055510948 | 0 | 1 | 0 | 0.001549435206 | 0 |
| GORG_SAMEA6069936_SAGS_AG893I03 | 4 | 0.04512986631 | AEGEAN-169 | AG-337-I02 | AG-337-I02 sp90 | 0 | 0.002713723139 | 0.9999997209 | 0.9999997209 | 0.1189952527 | 0.004604175684 | 0 | 0.04851659197 | 0 | 7.54E-04 | 0 |
| GORG_SAMEA6069959_SAGS_AG893J17 | 2 | -0.1133953281 | Puniceispirillales | UBA3439 | | 0 | 0 | 0.03792921652 | 1.000000271 | 0.1199940049 | 6.50E-06 | 0 | 0.9982586148 | 0 | 0.001938729561 | 0.04193405048 |
| GORG_SAMEA6070007_SAGS_AG893M20 | 8 | 0.02379047633 | AEGEAN-169 | AG-337-I02 | AG-337-I02 sp90 | 0 | 0 | 0.1627843546 | 1.000000246 | 0.2014697434 | 1.000000246 | 0 | 1.000000246 | 0 | 0.001436687632 | 0 |
| GORG_SAMEA6070209_SAGS_AG894L20 | 1 | 0.03939725917 | Flavobacteriales | MED-G14 | MED-G14 sp902 | 0 | 0 | 0.9999999951 | 1.58E-04 | 0.281221209 | 0.9833333333 | 0 | 0.9999999951 | 0 | 0.001559703634 | 0 |
| GORG_SAMEA6070235_SAGS_AG894N13 | 2 | 0.0181683655 | AEGEAN-169 | AG-337-I02 | AG-337-I02 sp90 | 0 | 0 | 0.01792525052 | 1 | 0.1229227964 | 6.34E-06 | 0 | 0.05694363067 | 0 | 0.001591336199 | 0 |
| GORG_SAMEA6070272_SAGS_AG894P23 | 3 | 0.04446567958 | Pelagibacterales | Pelagibacter | Pelagibacter sp9 | 0 | 0 | 0.9999997648 | 8.89E-05 | 0.007623716672 | 0.9999997648 | 0 | 0.1601173328 | 0 | 0.9999997648 | 0.06146333503 |
| GORG_SAMEA6070327_SAGS_AG895E15 | 4 | -0.005436614149 | Pelagibacterales | Pelagibacter_A | Pelagibacter_A s | 9.87E-05 | 0 | 0.999999796 | 0.01666666667 | 0.08971678103 | 0.001963931487 | 0 | 0.999999796 | 1.70E-04 | 0.001846669379 | 0.002675554923 |
| GORG_SAMEA6071019_SAGS_AG899A06 | 1 | 0.03254495682 | Pelagibacterales | Pelagibacter | Pelagibacter sp9 | 0 | 0 | 1 | 2.07E-04 | 9.86E-04 | 1 | 0 | 1 | 0.9990792556 | 0.005944446892 | 0.03258544655 |
| GORG_SAMEA6071089_SAGS_AG899G02 | 1 | 0.03470599064 | Pelagibacterales | Pelagibacter | Pelagibacter sp9 | 0 | 0 | 1 | 8.09E-05 | 0.2469492984 | 1 | 0 | 0.05420133525 | 0 | 0.003660230127 | 0.01970208983 |
| GORG_SAMEA6071177_SAGS_AG899M14 | 3 | 0.03311605798 | SAR86 | AG-339-G14 | AG-339-G14 sp9 | 0 | 0 | 0.04796018487 | 0 | 0 | 1 | 0 | 1 | 0 | 1 | 0.1269623538 |
| GORG_SAMEA6071336_SAGS_AG900I13 | 2 | 0.0679553398 | Pelagibacterales | Pelagibacter | Pelagibacter sp9 | 0 | 0 | 0.03417650461 | 0 | 0.1876362034 | 4.80E-06 | 0 | 0.0528202233 | 0 | 0.008032929997 | 0.01646553215 |
| GORG_SAMEA6071497_SAGS_AG901E05 | 1 | -0.06542601707 | Flavobacteriales | MS024-2A | MS024-2A sp902 | 0 | 0 | 1 | 0.01673428062 | 1 | 1 | 0 | 1 | 0 | 0.02114186495 | 0 |
| GORG_SAMEA6071777_SAGS_AG903M21 | 8 | -0.03316917555 | Pelagibacterales | Pelagibacter_A | Pelagibacter_A s | 0 | 0 | 0.9999998801 | 0.9999998801 | 0.00921067546 | 0.9999998801 | 0 | 0.9999998801 | 0 | 9.46E-04 | 0.0289658291 |
| GORG_SAMEA6071779_SAGS_AG903N02 | 4 | -0.03899572754 | Flavobacteriales | MED-G14 | | 0 | 0 | 1.000000287 | 1.98E-04 | 1.000000287 | 0.005033285883 | 0 | 1.000000287 | 0 | 0.001647716492 | 1.000000287 |
| GORG_SAMEA6072083_SAGS_AG905E11 | 8 | 0.01790270126 | Pelagibacterales | Pelagibacter | Pelagibacter sp9 | 0 | 0 | 1 | 1 | 0.98478326 | 1 | 0 | 0.08367632822 | 0 | 0.001899177005 | 0.02812518408 |
| GORG_SAMEA6073085_SAGS_AG911C08 | 8 | -0.02396443873 | Puniceispirillales | GCA-002684695 | | 3.31E-06 | 0 | 0.07116794887 | 1 | 0.01282819523 | 1 | 0 | 0.07627813612 | 3.82E-04 | 0 | 0.0665544303 |
| GORG_SAMEA6073323_SAGS_AG912F03 | 4 | 0.01483649749 | AEGEAN-169 | AG-337-I02 | AG-337-I02 sp90 | 0 | 0 | 0.9999997465 | 1.05E-04 | 0.9999997465 | 4.56E-06 | 0 | 0.9999997465 | 0 | 0.01473052378 | 0.03059755721 |
| GORG_SAMEA6073345_SAGS_AG912G22 | 1 | 0.03011653515 | Pelagibacterales | Pelagibacter | Pelagibacter sp9 | 0 | 0.01833216789 | 1 | 1.47E-04 | 0.3324764468 | 1 | 0 | 0.09050060391 | 0 | 8.11E-04 | 0 |
| GORG_SAMEA6073557_SAGS_AG913I06 | 3 | 0.01141605673 | Pelagibacterales | Pelagibacter_A | | 0 | 0 | 0.9999999956 | 2.57E-06 | 0.1771979772 | 0.9999999956 | 0 | 0.9999999956 | 0 | 0.9670180782 | 0.01701880928 |
| GORG_SAMEA6073589_SAGS_AG913K21 | 8 | 9.22E-04 | Pelagibacterales | Pelagibacter | | 0 | 0 | 1 | 1 | 0.2643450722 | 1 | 0 | 0.05558945229 | 0.001829795122 | 3.20E-05 | 0 |
| GORG_SAMEA6073608_SAGS_AG913M07 | 2 | 0.03651366833 | AEGEAN-169 | AG-337-I02 | AG-337-I02 sp90 | 0 | 0 | 0.04172524241 | 0.9999997316 | 0.1364711289 | 0.004361129803 | 0 | 0.9999997316 | 0 | 0 | 0.002913770748 |
| GORG_SAMEA6073834_SAGS_AG914N11 | 1 | 0.05552394156 | Pelagibacterales | Pelagibacter_A | Pelagibacter_A s | 0 | 0 | 1.000000333 | 1.62E-04 | 0.4441498517 | 1.000000333 | 0 | 1.000000333 | 0 | 0.009097716473 | 0 |
| GORG_SAMEA6074186_SAGS_AG916G20 | 7 | 0.03417005445 | Flavobacteriales | MED-G14 | MED-G14 sp902 | 1 | 0 | 1 | 1.11E-04 | 1 | 0 | 0 | 0.0471611436 | 0 | 1 | 0 |
| GORG_SAMEA6074373_SAGS_AG917F02 | 1 | -0.05366165832 | Kiritimatiellales | GCA-2712105 | GCA-2712105 sp | 4.78E-08 | 0 | 1 | 0.01674099973 | 0.1226435629 | 1 | 0.004654733648 | 0.9476115259 | 0 | 0.1937361967 | 0 |
| GORG_SAMEA6074416_SAGS_AG917J06 | 8 | 0.03286453398 | Pelagibacterales | Pelagibacter | | 0 | 0 | 0.9999997049 | 0.9999997049 | 0.9999997049 | 0.9999997049 | 0 | 0.9999997049 | 0 | 0.003290305729 | 0 |
| GORG_SAMEA6074543_SAGS_AG918C15 | 3 | 0.01885081177 | Pelagibacterales | Pelagibacter | Pelagibacter sp9 | 0 | 0 | 1.000000284 | 1.63E-04 | 0.03565726073 | 1.000000284 | 0 | 0.08817843778 | 0 | 1.000000284 | 1.000000284 |
| GORG_SAMEA6074616_SAGS_AG918K06 | 1 | 0.04649769139 | SAR86 | MED-G85 | MED-G85 sp003 | 0 | 0 | 1 | 1.16E-04 | 0 | 1 | 0 | 1 | 0 | 0.00157127067 | 0 |
| GORG_SAMEA6074649_SAGS_AG918M19 | 1 | 0.01539713798 | Pelagibacterales | Pelagibacter | | 0 | 0 | 1.000000191 | 1.35E-04 | 1.000000191 | 1.000000191 | 0.1599999236 | 1.000000191 | 7.72E-04 | 7.93E-04 | 0 |

| | | | | | | | | | | | | | | | | |
|---|---|---|---|---|---|---|---|---|---|---|---|---|---|---|---|---|
| GORG_SAMEA6074891_SAGS_AG920B11 | 6 | -0.02133314691 | Flavobacteriales | MS024-2A | MS024-2A sp902 | 0.1967054732 | 0 | 0.03887294181 | 0.01666742517 | 1.000000142 | 0.03598887305 | 0.002605721807 | 0.07122415979 | 0.001785821403 | 0.00276897547 | 0.006983076537 |
| GORG_SAMEA6075144_SAGS_AG349C11 | 1 | 0.03570905577 | Pelagibacterales | Pelagibacter_A | Pelagibacter_A s | 0 | 0 | 1 | 7.86E-05 | 0.04580226453 | 1 | 0.08838930041 | 0.03189235974 | 0 | 9.51E-04 | 0 |
| GORG_SAMEA6075158_SAGS_AG349D08 | 1 | -4.72E-04 | Pelagibacterales | Pelagibacter_A | Pelagibacter_A s | 0 | 0 | 1 | 0.03531119976 | 9.65E-04 | 1 | 0.004150655297 | 1 | 0 | 0.001791187955 | 0 |
| GORG_SAMEA6075358_SAGS_AG404A02 | 7 | 0.006120964142 | SAR202 | GCA-002694895 | GCA-002694895 | 7.86E-07 | 0 | 0.02068727729 | 6.92E-05 | 0.9839273096 | 1.22E-06 | 0.07037881164 | 0.02423087973 | 0 | 0.9999995752 | 5.23E-04 |
| GORG_SAMEA6075523_SAGS_AG404M02 | 5 | 0.04783741411 | TMED127 | GCA-2711515 | GCA-2711515 sp | 0 | 0 | 0.03314678114 | 4.47E-05 | 0.5306780035 | 1 | 0.05579946996 | 0.0392297924 | 0 | 0 | 0.05194677562 |
| GORG_SAMEA6075696_SAGS_AG390G04 | 2 | -0.02426632218 | AEGEAN-169 | AG-337-I02 | | 0 | 0 | 0.03761977478 | 1.20E-04 | 0.219331324 | 0.001966211127 | 0 | 0.07602135611 | 6.94E-04 | 0.004806293866 | 1.49E-06 |
| GORG_SAMEA6075741_SAGS_AG390J14 | 4 | 0.006922938661 | Flavobacteriales | MED-G13 | MED-G13 sp902 | 0 | 0 | 1 | 0.01680503369 | 1 | 0.03110549055 | 0.01666666667 | 1 | 0 | 0.0213466993 | 1 |
| GORG_SAMEA6075810_SAGS_AG390M23 | 4 | -0.03858740716 | Flavobacteriales | MAG-121220-bin | MAG-121220-bin | 0 | 0 | 0.9999998165 | 1.32E-06 | 0.9836724626 | 0.005221149648 | 0 | 0.2463577426 | 0 | 0 | 0.02941361425 |
| GORG_SAMEA6075916_SAGS_AG414E04 | 1 | 0.01700712195 | Actinomarinales | Actinomarina | Actinomarina sp( | 0 | 0 | 0.9999996701 | 1.79E-04 | 0.9999996701 | 0.9999996701 | 0 | 0.03750085355 | 0.001394711641 | 0.005551796813 | 0 |
| GORG_SAMEA6076030_SAGS_AG414N23 | 8 | -0.00284242223! | AEGEAN-169 | AG-337-I02 | AG-337-I02 sp90 | 0 | 0 | 1 | 1 | 1 | 1 | 0 | 0.1011749106 | 0 | 0.001617950627 | 0.06881745682 |
| GORG_SAMEA6076353_SAGS_AG422M03 | 8 | -0.08963073752 | Rhodobacterales | CACIJG01 | CACIJG01 sp902 | 0.001050306308 | 0 | 1 | 1 | 0.122930958 | 1 | 0 | 1 | 0 | 2.94E-06 | 0 |
| GORG_SAMEA6076385_SAGS_AG422O05 | 7 | 0.02160610588 | SAR86 | SAR86A | SAR86A sp0032 | 0 | 0 | 0.9999998009 | 0.9999998009 | 0.003250904247 | 0.0215269175 | 0 | 0.1036923594 | 0 | 0.9999998009 | 0.01932466529 |
| GORG_SAMEA6076629_SAGS_AG447M16 | 4 | -0.00788764509! | SAR86 | D2472 | D2472 sp902595 | 0 | 0.9833333333 | 0.9838187407 | 0.01673724387 | 0 | 1.13E-05 | 0 | 0.9846929123 | 0 | 0.02349549388 | 0 |
| GORG_SAMEA6077196_SAGS_AG319P17 | 1 | 0.03273982387 | Flavobacteriales | Marisimplicoccus | Marisimplicoccus | 0 | 0 | 1.000000276 | 2.20E-04 | 1.000000276 | 1.000000276 | 0 | 1.000000276 | 0 | 0 | 0 |
| GORG_SAMEA6077297_SAGS_AG325E20 | 1 | 7.79E-04 | Pelagibacterales | MED727 | MED727 sp0026 | 0 | 0 | 1 | 7.64E-05 | 0.1347594161 | 1 | 0.002246189839 | 0.05646153255 | 0 | 7.66E-04 | 0 |
| GORG_SAMEA6077398_SAGS_AG325L02 | 8 | 0.03828892378 | Pelagibacterales | Pelagibacter | Pelagibacter sp9 | 1.39E-07 | 0 | 1 | 1 | 0.727166058 | 1 | 0 | 1 | 0 | 0.004410036284 | 0 |
| GORG_SAMEA6077561_SAGS_AG333D04 | 1 | -0.05351754887 | Woeseiales | SP4260 | SP4260 sp00271 | 0 | 0.007430768893 | 0.9214921828 | 0.01671199828 | 0.02577563461 | 1.000000004 | 0.007093116265 | 0.07184524491 | 0 | 1.33E-05 | 0.006220054709 |
| GORG_SAMEA6077629_SAGS_AG333G15 | 1 | 0.04187107376 | Actinomarinales | Actinomarina | Actinomarina sp( | 0 | 0 | 1 | 1.64E-04 | 1 | 1 | 0 | 1 | 0.001456220311 | 0.005229464061 | 0 |
| GORG_SAMEA6078463_SAGS_AG453G04 | 1 | 0.005535628584 | Pelagibacterales | Pelagibacter_A | Pelagibacter_A s | 0 | 0 | 1.000000156 | 0.0167650005 | 0.3056758115 | 1.000000156 | 1.67E-04 | 0.9833836078 | 2.64E-05 | 0.001869131394 | 0.03510603791 |
| GORG_SAMEA6079413_SAGS_AG470J09 | 5 | -0.00228022682! | UBA1144 | GCA-002715585 | GCA-002715585 | 0 | 0 | 0.03922038777 | 4.73E-05 | 0.2669299731 | 1 | 0 | 8.18E-04 | 0 | 7.40E-04 | 0.06301260201 |
| GORG_SAMEA6079738_SAGS_AG426P08 | 1 | 0.03429051385 | Pelagibacterales | Pelagibacter | Pelagibacter sp9 | 0 | 0 | 0.9999997317 | 1.47E-04 | 0.2432635688 | 0.9999997317 | 0.01372090646 | 0.1315817222 | 0 | 0 | 0.03646477116 |
| GORG_SAMEA6079747_SAGS_AG426P19 | 1 | 0.01610809813 | Cytophagales | MED-G16 | MED-G16 sp902 | 0 | 0 | 1.000000196 | 1.29E-04 | 1.000000196 | 1.000000196 | 0 | 1.000000196 | 0 | 0 | 0.04775884011 |
| GORG_SAMEA6079844_SAGS_AG430F16 | 1 | 0.02877774509 | Pelagibacterales | Pelagibacter | Pelagibacter sp0 | 0 | 0 | 1 | 0 | 0.005909409876 | 1 | 0.3090808813 | 1 | 0.01521450749 | 7.88E-04 | 0 |
| GORG_SAMEA6080234_SAGS_AG439C18 | 1 | 0.0573998665 | Pelagibacterales | Pelagibacter | Pelagibacter sp9 | 0 | 0 | 1 | 4.20E-05 | 0.7511468002 | 1 | 0 | 0.06085901548 | 0 | 0.005841246783 | 0 |
| GORG_SAMEA6080277_SAGS_AG439G19 | 1 | 0.04779849938 | Pelagibacterales | Pelagibacter | Pelagibacter sp9 | 0 | 0 | 0.9999998336 | 0 | 0.2750462998 | 0.9999998336 | 0 | 0.935333376 | 0 | 0 | 0 |
| HOTS_SAMN07137059_METAG_EICKKFBA | 5 | -0.01852112365 | UBA1151 | GCA-002708145 | GCA-002708145 | 0 | 0 | 0.01453277959 | 1.45E-04 | 3.84E-06 | 1.00000032 | 0 | 1.00000032 | 0 | 0.001477736157 | 0 |
| HOTS_SAMN07137059_METAG_PMKAHHED | 3 | -0.07339085276 | Pirellulales | Bythopirellula | Bythopirellula sp( | 0 | 0 | 0.02637104469 | 1.67E-04 | 1.000000324 | 1.000000324 | 0.009284001271 | 0.05877082983 | 0 | 1.000000324 | 0 |
| MALA_SAMN05421697_METAG_OILIAGNK | 7 | -0.09899132363 | Chlamydiales | JABDGQ01 | | 0 | 0 | 1 | 1 | 0.6966023923 | 3.66E-06 | 0 | 1 | 0 | 1 | 0 |
| MALA_SAMN05421699_METAG_ACDJADOH | 1 | -0.3478152028 | Micavibrionales | | | 0 | 0 | 1 | 1.26E-04 | 0 | 1 | 0 | 0.04241623562 | 0 | 0.002767902501 | 0 |
| MALA_SAMN05421901_METAG_NCIHGEED | 7 | -0.06680770862 | SAR324 | Arctic96AD-7 | Arctic96AD-7 sp( | 0 | 0 | 0.04288534238 | 0.08343262237 | 0.06274884572 | 6.25E-06 | 0 | 0.9298117377 | 0 | 0.9999998045 | 0.03234087742 |
| MALA_SAMN05422107_METAG_EOEMECJL | 4 | -0.1534667863 | Flavobacteriales | Mesonia | Mesonia mobilis | 0 | 0 | 1 | 1.51E-04 | 0.06489932531 | 0 | 0 | 0.03382540419 | 0 | 0.001616151023 | 0 |
| MALA_SAMN05422113_METAG_ACPCAKPA | 2 | -0.1424908739 | Nevskiales | Abyssibacter | Abyssibacter sp0 | 0 | 0 | 0.02228687539 | 3.00E-06 | 0.001642076335 | 0.01998365143 | 0 | 0.04565402329 | 0 | 3.97E-05 | 0.03201966219 |
| MALA_SAMN05422114_METAG_NAJFPNCO | 2 | -0.552608519 | Pseudomonadale | Acinetobacter | Acinetobacter joh | 0 | 0 | 0.05474369176 | 1.13E-04 | 0.4606182177 | 0.01666728429 | 0 | 0.06124948684 | 0 | 0 | 0 |
| MALA_SAMN05422120_METAG_BOFCDAAO | 2 | -0.4562438883 | Enterobacterales | Alteromonas | Alteromonas con | 0 | 0 | 0.1873436392 | 1.23E-07 | 0.1362322567 | 3.74E-05 | 0 | 0.05336046052 | 0 | 0.009161343081 | 0.001036418774 |
| MALA_SAMN05422120_METAG_MALJNMLA | 5 | -0.2301584105 | Sphingomonadal | Qipengyuania | | 0 | 0 | 0.0244805245 | 1.82E-04 | 0 | 1 | 0 | 0.08285830695 | 0 | 0.02020648536 | 0 |
| MALA_SAMN05422120_METAG_NDMHKPKD | 5 | -0.599604848 | Enterobacterales | Alteromonas | | 0 | 0 | 0.008254175962 | 1.01E-04 | 0.01352966012 | 1 | 0 | 0.06587414238 | 0 | 0.01677056576 | 0.1428570404 |
| MALA_SAMN05422120_METAG_PINPMALI | 4 | -0.09945677778 | Flavobacteriales | Aquaticitalea | Aquaticitalea lipo | 1.15E-07 | 0 | 1.000000249 | 0.01666854065 | 0.985278079 | 0 | 0 | 0.9844149387 | 0 | 6.32E-04 | 0.03097835911 |
| MALA_SAMN05422121_METAG_GMGNEBDN | 4 | -0.01671921994 | Propionibacterial | Nocardioides | Nocardioides ma | 0 | 0 | 1.000000007 | 0.02449493191 | 0 | 9.76E-04 | 0 | 0.9844011417 | 0 | 7.63E-04 | 1.32E-04 |
| MALA_SAMN05422121_METAG_PMNELIKC | 4 | -0.4512274819 | Bacteriovoracale | GCA-2712005 | | 0 | 0 | 0.9999997084 | 0.06669619971 | 0.007828736248 | 0.03342833589 | 0 | 0.4188846354 | 0 | 7.64E-04 | 0.003488061272 |
| MALA_SAMN05422122_METAG_CKBPOFGO | 2 | -0.404023326 | Rhodobacterales | Sulfitobacter | Sulfitobacter pon | 2.86E-06 | 0 | 0.02758433361 | 0.9833341591 | 0.124166472 | 0.01667273965 | 0 | 0.0616360193 | 2.93E-04 | 1.54E-04 | 0.04292954549 |
| MALA_SAMN05422123_METAG_MFNFJBLI | 8 | -0.1851110534 | Pseudomonadale | Salinicola | Salinicola salariu | 2.78E-05 | 0 | 1.000000005 | 0.9842082786 | 0.001325209604 | 1.000000005 | 0 | 0.9374008866 | 0 | 5.58E-04 | 0.06024942892 |
| MALA_SAMN05422133_METAG_KFHCJPLA | 5 | -0.2431100563 | Rhodobacterales | Salipiger | Salipiger thioxid | 0 | 0 | 0.0306332583 | 0.01666666667 | 0.09702509556 | 1.000000205 | 0 | 0.06493532545 | 0 | 0 | 0 |
| MALA_SAMN05422133_METAG_LICJKDFC | 2 | -0.1254266839 | Rhizobiales | Oricola | Oricola sp00270( | 0 | 0 | 0.1709625707 | 1.25E-04 | 0.0120058887 | 0 | 0 | 0.07480060171 | 0 | 7.67E-04 | 0 |
| MALA_SAMN05422136_METAG_MLBOFNBE | 6 | -0.00140957119 | Propionibacterial | Nocardioides | Nocardioides sp( | 0 | 0 | 0.06451014253 | 0.006186756528 | 0.9833329906 | 1.38E-07 | 0 | 0.0794501701 | 0 | 7.26E-04 | 0.05564032436 |
| MALA_SAMN05422148_METAG_BICMPFEL | 5 | -0.04620920741 | UBA3495 | UBA11650 | UBA11650 sp002 | 0 | 0 | 0.003086745205 | 6.59E-05 | 0.02345536955 | 0.9999997911 | 0 | 0.04571693286 | 0 | 0.02948814707 | 0.06218349355 |
| MALA_SAMN05422148_METAG_IFLPOAIP | 2 | -0.2013970636 | Pseudomonadale | Ketobacter | Ketobacter sp00: | 0 | 0 | 0.1529798391 | 4.23E-05 | 0.001033062312 | 0.002302674387 | 0 | 0.02951089197 | 0 | 7.61E-05 | 0 |
| MALA_SAMN05422149_METAG_ADPGENHC | 4 | -0.03965548241 | Flavobacteriales | CAJJDF01 | | 0 | 0 | 1.000000235 | 9.55E-05 | 1.000000235 | 0.003669526155 | 0 | 1.000000235 | 0 | 0 | 0 |
| MALA_SAMN05422149_METAG_IOGIMBKE | 8 | -0.04851784547 | Arenicellales | REDSEA-S09-B1 | REDSEA-S09-B1 | 1.75E-05 | 2.84E-04 | 0.02632581635 | 0.999999747 | 0.05637524161 | 0.999999747 | 6.09E-05 | 0.06452579835 | 0 | 0.001571901576 | 0 |
| MALA_SAMN05422153_METAG_AGJKPNIP | 4 | -0.3155686004 | Pseudomonadale | Psychrobacter | Psychrobacter sp | 0 | 0 | 0.9999999957 | 2.23E-06 | 0.1012600304 | 0.01666666233 | 0 | 0.0585258127 | 0 | 0 | 0 |
| MALA_SAMN05422165_METAG_NBPIDFID | 3 | -0.2434561008 | Rhodobacterales | Marinovum | Marinovum algico | 0 | 0 | 0.02203722542 | 3.55E-05 | 0.021630556 | 0.9999998777 | 0.00295202773 | 0.0437365384 | 0 | 0.9999998777 | 0.008823160582 |
| MALA_SAMN05422166_METAG_LIPHNDPD | 1 | 0.0232129723 | Propionibacterial | Nocardioides | Nocardioides sal | 0 | 0 | 1 | 0 | 0.2777657229 | 0.9833333333 | 0 | 0.1101911127 | 0 | 0 | 0.9837833915 |
| MALA_SAMN05422166_METAG_OAKDMEED | 2 | -0.0185077094 | Acidimicrobiales | UBA9410 | UBA9410 sp012 | 8.17E-08 | 0 | 0.05568135389 | 0.1084009229 | 0 | 0.0166693022 | 0 | 0.01350968054 | 0 | 3.25E-05 | 0 |
| MALA_SAMN05422183_METAG_NNKMKILK | 8 | -0.09263515215 | Marinisomatales | GCA-002701945 | GCA-002701945 | 0 | 0 | 0.09944694735 | 0.9999997582 | 0.9999997582 | 0.9999997582 | 0 | 0.9999997582 | 0.02422346896 | 2.45E-04 | 0 |
| MALA_SAMN05422191_METAG_MKDPFEDB | 5 | -0.3309574577 | Pseudomonadale | Marinobacter | Marinobacter vini | 0 | 0 | 0.06740981792 | 7.31E-05 | 0.1069108455 | 1 | 0 | 1 | 0 | 0 | 0 |
| MARD_SAMD00000349_REFG_MMP00000349 | 4 | -0.5025372972 | Lactobacillales | Streptococcus | Streptococcus pa | 0 | 0 | 1.000000003 | 1.000000003 | 0.3425819346 | 0 | 0 | 0.07161843897 | 0 | 0.007462096645 | 0 |
| MARD_SAMD00000371_REFG_MMP00000371 | 2 | -0.3828872597 | Bacillales_H | C254 | C254 sp0027973 | 0 | 0 | 0.2828935316 | 1.56E-04 | 0.003433333726 | 6.49E-06 | 0 | 0.06528397364 | 0 | 0 | 0 |
| MARD_SAMD00000372_REFG_MMP00000372 | 2 | -0.3430760384 | Bacillales_H | Halalkalibacter | Halalkalibacter sp | 0 | 0 | 0.04188016933 | 1.85E-04 | 0.1687557945 | 0.09636521101 | 0 | 0.05675659718 | 9.53E-04 | 0.001253262831 | 0.01666666002 |
| MARD_SAMD00000404_REFG_MMP00000404 | 2 | -0.7570061796 | Enterobacterales | Edwardsiella | Edwardsiella ang | 0 | 0 | 0.09312369349 | 5.00E-05 | 0.006332508244 | 0.1986793267 | 0 | 0.04179651884 | 0 | 4.13E-05 | 0 |
| MARD_SAMD00000449_REFG_MMP00000449 | 6 | -0.104016572 | Bacteroidales | Geofilum | Geofilum rubicun | 0 | 0 | 0.1362075698 | 1.12E-04 | 0.9999998169 | 0 | 0 | 0.080878176 | 0 | 0 | 0 |
| MARD_SAMD00000547_REFG_MMP00000547 | 4 | 0.0662019218 | Rhodothermales | Rhodothermus | Rhodothermus m | 8.00E-08 | 0 | 0.9999997217 | 0.01666667066 | 0.9999997217 | 6.24E-05 | 0 | 0.05081583842 | 0 | 0.007291564097 | 0.02704117336 |
| MARD_SAMD00000560_REFG_MMP00000560 | 2 | -0.5630391372 | Enterobacterales | Vibrio | Vibrio alginolyticu | 0 | 0.0370944511 | 0.2063153515 | 5.66E-05 | 0.01301266576 | 0.002037929933 | 0 | 0.06541075194 | 0 | 0 | 0 |
| MARD_SAMD00006053_REFG_MMP00006053 | 2 | -0.3561888422 | Bacillales_H | Bacillus_H | Bacillus_H misca | 0 | 0.02412239458 | 0.06048243203 | 1.75E-04 | 0.005882163222 | 3.57E-06 | 0 | 0.1667874981 | 0 | 0 | 0.7666666667 |
| MARD_SAMD00009117_REFG_MMP00009117 | 2 | -0.4451545285 | Bacillales_H | Bacillus_H | Bacillus_H xiaoxi | 0 | 0 | 0.07154966087 | 1.35E-04 | 0.007022125546 | 6.14E-06 | 0 | 0.2535641505 | 0 | 0 | 1 |
| MARD_SAMD00011086_REFG_MMP00011086 | 2 | -0.6545970604 | Enterobacterales | Shewanella | Shewanella indic | 0 | 0 | 0.04066251917 | 8.84E-05 | 0.01533983801 | 0.1233333486 | 0 | 0.110750894 | 0 | 7.84E-04 | 0 |
| MARD_SAMD00017647_REFG_MMP00017647 | 2 | -0.1694804269 | Streptosporangia | Nocardiopsis | Nocardiopsis alb | 0 | 0 | 0.1335131862 | 0.007292479537 | 0.04594005255 | 2.74E-05 | 0 | 0.04603931422 | 0 | 0.003406897658 | 0.008243927484 |
| MARD_SAMD00017759_REFG_MMP00017759 | 2 | -0.08059890691 | Parvibaculales | Tepidicaulis | Tepidicaulis mari | 5.13E-08 | 0 | 0.02689260968 | 0 | 1.10E-04 | 1.60E-04 | 0 | 0.06272072729 | 0 | 2.80E-04 | 0.0276676101 |
| MARD_SAMD00018367_REFG_MMP00018367 | 2 | -0.4311581564 | Enterobacterales | Pseudoalteromor | Pseudoalteromor | 0 | 0 | 0.1873698742 | 1.93E-05 | 0 | 4.80E-06 | 0 | 0.1082016128 | 0 | 0.01135242998 | 0 |
| MARD_SAMD00018622_REFG_MMP00018622 | 8 | 0.01357597338 | Thermoanaeroba | Thermoanaeroba | Thermoanaeroba | 0 | 0 | 0.08375382542 | 0.9999998398 | 0.04622029143 | 0.8499998398 | 0 | 0.06009579332 | 0 | 4.35E-04 | 0 |
| MARD_SAMD00018681_REFG_MMP00018681 | 2 | -0.5769939129 | Enterobacterales | Vibrio | Vibrio diazotroph | 0 | 0 | 0.1564049294 | 1.18E-04 | 0 | 0 | 0 | 0.05579230747 | 0 | 0 | 0 |
| MARD_SAMD00018689_REFG_MMP00018689 | 5 | 0.02375467656 | Actinomycetales | Piscicoccus | Piscicoccus intes | 0.01796398704 | 0 | 0.09807147983 | 1.36E-04 | 0.04075108683 | 1.000000218 | 0 | 0.05272132525 | 0 | 7.81E-04 | 0 |
| MARD_SAMD00019133_REFG_MMP00019133 | 2 | -0.5121648431 | Enterobacterales | Vibrio | Vibrio hyugaensi | 0 | 0 | 0.06288372091 | 2.91E-05 | 0.005740357064 | 5.87E-07 | 0 | 0.06625056461 | 0 | 4.00E-05 | 0 |
| MARD_SAMD00019143_REFG_MMP00019143 | 2 | -0.4456174276 | Enterobacterales | Vibrio | Vibrio rotiferianus | 0 | 0 | 0.0547779415 | 1.67E-05 | 0.002444152799 | 0.0450964938 | 0 | 0.0624550745 | 0 | 2.47E-05 | 0 |
| MARD_SAMD00019870_REFG_MMP00019870 | 2 | -0.2119150281 | Sphingomonadal | Novosphingobiur | Novosphingobiur | 0 | 0 | 0.03058206587 | 7.38E-05 | 2.95E-04 | 0 | 0 | 0.06088367336 | 0 | 0.008570938858 | 0.01450872302 |

| | | | | | | | | | | | | | | | | |
|---|---|---|---|---|---|---|---|---|---|---|---|---|---|---|---|---|
| MARD_SAMD00020220_REFG_MMP00020220 | 2 | -0.1195454 | Flavobacteriales | Jejuia | Jejuia pallidilutea | 1.08E-07 | 0 | 0.0172680138 | 0.03333333058 | 0.04634595711 | 0 | 0 | 0.04781436156 | 0 | 2.67E-05 | 5.04E-04 |
| MARD_SAMD00025187_REFG_MMP00025187 | 2 | -0.2761325136 | Pseudomonadale | Cellvibrio | Cellvibrio sp0009 | 2.58E-06 | 0 | 0.08083130421 | 0 | 0.06708887443 | 0.01667093364 | 0 | 0.02053683497 | 0 | 0.006784076531 | 0.06020407411 |
| MARD_SAMD00036551_REFG_MMP00036551 | 4 | -0.5605582345 | Enterobacterales | Photobacterium | Photobacterium I | 0 | 0 | 0.9874188788 | 2.41E-05 | 0.002262271486 | 0.04883931937 | 0 | 0.06245986505 | 0 | 1.12E-04 | 0 |
| MARD_SAMD00036589_REFG_MMP00036589 | 5 | -0.3837643504 | Enterobacterales | Pseudoalteromor | Pseudoalteromor | 2.28E-04 | 0 | 0.02712382429 | 1.70E-06 | 3.83E-05 | 1 | 0 | 0.10797965 | 0 | 1.31E-05 | 0 |
| MARD_SAMD00036590_REFG_MMP00036590 | 1 | -0.4405569034 | Enterobacterales | Pseudoalteromor | Pseudoalteromor | 0 | 0 | 0.6132166375 | 0 | 0.01044322038 | 0.9999998976 | 0 | 0.0484319788 | 0 | 0 | 0 |
| MARD_SAMD00036591_REFG_MMP00036591 | 5 | -0.3541422458 | Enterobacterales | Pseudoalteromor | Pseudoalteromor | 0 | 0 | 0.05970959869 | 0 | 0.01197705848 | 0.9999998278 | 0 | 0.1166873094 | 0 | 0 | 0 |
| MARD_SAMD00036594_REFG_MMP00036594 | 5 | -0.3925428138 | Enterobacterales | Pseudoalteromor | Pseudoalteromor | 0 | 0 | 0.04323119234 | 3.45E-05 | 0.002463506351 | 1 | 0 | 0.1069023115 | 0 | 0.003832159696 | 0.001956662214 |
| MARD_SAMD00036597_REFG_MMP00036597 | 2 | -0.3826307763 | Enterobacterales | Paraglaciecola | Paraglaciecola cl | 0 | 0 | 0.03553162262 | 2.94E-07 | 0 | 0.03333333325 | 0 | 0.06224007575 | 0 | 6.35E-07 | 0.1536072592 |
| MARD_SAMD00036600_REFG_MMP00036600 | 2 | -0.4632047906 | Enterobacterales | Aliiglaciecola | Aliiglaciecola lipo | 0 | 0 | 0.0485427659 | 0 | 0.08123358667 | 5.44E-06 | 0 | 0.07451298737 | 0 | 4.84E-05 | 7.62E-04 |
| MARD_SAMD00036601_REFG_MMP00036601 | 2 | -0.3474627444 | Enterobacterales | Paraglaciecola | Paraglaciecola ai | 0 | 0 | 0.02737019779 | 3.98E-06 | 4.76E-04 | 2.66E-06 | 0 | 0.05503814062 | 0 | 2.43E-05 | 0.03089668343 |
| MARD_SAMD00036602_REFG_MMP00036602 | 5 | -0.380503556 | Enterobacterales | Paraglaciecola | Paraglaciecola m | 0 | 0 | 0.02955603163 | 1.42E-05 | 0 | 1 | 0.008125261296 | 0.06202322156 | 0 | 3.78E-05 | 0.1596661084 |
| MARD_SAMD00036604_REFG_MMP00036604 | 2 | -0.3897890049 | Enterobacterales | Paraglaciecola | Paraglaciecola p | 0 | 0 | 0.03931315429 | 9.45E-06 | 0.002372970355 | 0 | 0 | 0.06562421155 | 0 | 0 | 0.1564348405 |
| MARD_SAMD00036605_REFG_MMP00036605 | 2 | -0.3178981682 | Enterobacterales | Paraglaciecola | Paraglaciecola p | 0 | 0 | 1.27E-04 | 1.49E-07 | 1.68E-04 | 0.01666666667 | 0 | 0.04682162507 | 0 | 0.005511911336 | 0.2180256034 |
| MARD_SAMD00036617_REFG_MMP00036617 | 2 | -0.599246339 | Enterobacterales | Rheinheimera | Rheinheimera na | 0 | 0 | 0.03846767573 | 8.65E-05 | 0.005423387409 | 5.09E-06 | 0 | 0.2780913357 | 0 | 0.008651967133 | 0 |
| MARD_SAMD00036654_REFG_MMP00036654 | 2 | -0.5321680631 | Enterobacterales | Vibrio | Vibrio jasicida | 0 | 0 | 0.0474137308 | 2.97E-05 | 0.009361300789 | 0.09101513445 | 0 | 0.07197201258 | 0 | 2.68E-05 | 0 |
| MARD_SAMD00036658_REFG_MMP00036658 | 2 | -0.4962609639 | Enterobacterales | Vibrio | Vibrio owensii | 0 | 0 | 0.06530631523 | 2.50E-05 | 0.07733507252 | 0.004547713728 | 0 | 0.1583547091 | 0 | 4.68E-06 | 0 |
| MARD_SAMD00036665_REFG_MMP00036665 | 2 | -0.09135184551 | Acidimicrobiales | Ilumatobacter_A | Ilumatobacter_A | 0 | 0 | 0.01142238035 | 0.9999996915 | 0.08307062311 | 0 | 0 | 0 | 0 | 7.28E-04 | 0 |
| MARD_SAMD00036772_REFG_MMP00036772 | 2 | -0.18859865 | Cyanobacteriales | Limnothrix | Limnothrix sp002 | 0 | 0 | 0.05981153653 | 1.07E-07 | 0.216607657 | 4.32E-04 | 0 | 0.04199581556 | 0 | 0.006138145625 | 0 |
| MARD_SAMD00039863_REFG_MMP00039863 | 5 | -0.1114641687 | Desulfobacterale | Desulfatitalea | Desulfatitalea tep | 0 | 0 | 0.02546594395 | 1.02E-05 | 0.03497539818 | 0.95 | 0 | 0.02824640376 | 0 | 0.02242561344 | 0 |
| MARD_SAMD00039894_REFG_MMP00039894 | 5 | -0.7361724053 | Enterobacterales | Pseudoalteromor | Pseudoalteromor | 0 | 0 | 0.04542609276 | 8.51E-07 | 0.001435912918 | 1 | 0 | 0.04747774491 | 0 | 1.29E-05 | 1.23E-06 |
| MARD_SAMD00041794_REFG_MMP00041794 | 2 | -0.550349836 | Enterobacterales | Vibrio | Vibrio halioticoli | 0 | 0 | 0.05624416463 | 3.11E-05 | 5.60E-04 | 2.61E-07 | 0 | 0.03054467329 | 0 | 4.01E-04 | 0 |
| MARD_SAMD00041798_REFG_MMP00041798 | 2 | -7.31E-04 | Actinomycetales | Mobilicoccus | Mobilicoccus pela | 0 | 0 | 0.03844725194 | 0.01668221581 | 0.494035364 | 0 | 0 | 0.9841165033 | 0 | 4.04E-05 | 2.27E-04 |
| MARD_SAMD00041816_REFG_MMP00041816 | 2 | -0.9688281422 | Enterobacterales | Vibrio | Vibrio proteolytic | 0 | 0 | 0.1049782177 | 1.11E-04 | 0.004761702148 | 3.68E-06 | 0 | 0.06294165376 | 0 | 0 | 0 |
| MARD_SAMD00046480_REFG_MMP00046480 | 4 | -0.00718732575 | Actinomycetales | Janibacter | Janibacter coralli | 0 | 0 | 0.8080933355 | 1.59E-04 | 0.06651188606 | 0.2000046855 | 0 | 0.05414105993 | 0 | 0 | 0 |
| MARD_SAMD00046500_REFG_MMP00046500 | 2 | -0.1074474171 | Actinomycetales | Kribbia | Kribbia dieselivor | 0 | 0 | 0.03891319448 | 1.37E-04 | 0.2158354881 | 5.46E-06 | 0 | 0.05113357409 | 0 | 8.33E-04 | 0 |
| MARD_SAMD00046757_REFG_MMP00046757 | 2 | -0.6713642157 | Enterobacterales | Vibrio | Vibrio natriegens | 0 | 0.06990115245 | 0.1229962142 | 3.92E-05 | 0 | 0.1416680847 | 0 | 0.05878926754 | 0 | 3.16E-05 | 0 |
| MARD_SAMD00046758_REFG_MMP00046758 | 2 | -0.630853829 | Enterobacterales | Vibrio | Vibrio nereis | 0 | 0 | 0.1619332268 | 5.52E-05 | 0.006626115836 | 0.01117692019 | 0 | 0.06992731986 | 0 | 2.89E-05 | 0 |
| MARD_SAMD00046775_REFG_MMP00046775 | 2 | -0.1682772645 | Mycobacteriales | Rhodococcus | Rhodococcus ma | 0 | 0 | 0.1211681483 | 1.55E-04 | 0.3098438761 | 0.1231930104 | 0 | 0.04825691627 | 0 | 8.14E-04 | 0 |
| MARD_SAMD00050834_REFG_MMP00050834 | 4 | -0.0898802098 | Flavobacteriales | Tenacibaculum | Tenacibaculum o | 0 | 0 | 0.9710917386 | 1.70E-06 | 0.9999999969 | 6.24E-05 | 0 | 0.0635401049 | 0 | 0 | 0.3111192308 |
| MARD_SAMD00053328_REFG_MMP00053328 | 5 | -0.6131117824 | Brevibacillales | Brevibacillus | Brevibacillus port | 0 | 0 | 0.1039234899 | 1.58E-04 | 0.005689250388 | 1 | 0 | 0.08268786277 | 0 | 0 | 0 |
| MARD_SAMD00056695_REFG_MMP00056695 | 2 | -0.2035705476 | Mycobacteriales | Rhodococcus | Rhodococcus eq | 0 | 0 | 0.03503766293 | 0.09733229166 | 0.05010397166 | 4.69E-05 | 0 | 0.04591538016 | 0 | 8.03E-04 | 0 |
| MARD_SAMD00065689_REFG_MMP00065689 | 5 | -0.1747825477 | Campylobacteral | Halarcobacter | Halarcobacter sp | 0.1042864332 | 0 | 0.1803616647 | 4.21E-05 | 0.04067200962 | 0.9999999217 | 0 | 0.05686708579 | 1.52E-04 | 0 | 0 |
| MARD_SAMD00113762_REFG_MMP00113762 | 8 | -0.1105754665 | Aggregatilineales | Aggregatilinea | Aggregatilinea le | 0 | 0 | 0.03827029869 | 1.00000023 | 0.04427389799 | 1.00000023 | 0 | 1.00000023 | 0 | 0 | 0 |
| MARD_SAMD00129508_REFG_MMP00129508 | 2 | -0.1376995917 | Streptomycetales | Streptomyces | Streptomyces sp | 0 | 0 | 0.0308641517 | 0.06048344186 | 0.001613617746 | 1.07E-05 | 0 | 0.04095256244 | 0 | 7.65E-04 | 0 |
| MARD_SAMEA104307712_REFG_MMP10430771 | 4 | -0.3301425493 | Flavobacteriales | Chryseobacteriur | Chryseobacteriur | 0 | 0 | 1 | 1.93E-06 | 0.06531122913 | 0.002755774994 | 0 | 0.07001155678 | 0 | 5.16E-08 | 0 |
| MARD_SAMEA104692609_REFG_MMP10469260 | 2 | -0.3418459362 | Rhodobacterales | Ruegeria | Ruegeria sp9003 | 0 | 0 | 0.07977148791 | 0 | 0.007247607936 | 0 | 0 | 0.03247193913 | 0 | 0.001825827547 | 0 |
| MARD_SAMEA104692611_REFG_MMP10469261 | 2 | -0.3066287332 | Rhodobacterales | Sulfitobacter | Sulfitobacter mec | 3.78E-07 | 0 | 0.1143547054 | 1.54E-04 | 0.003531282134 | 0 | 0 | 1.000000189 | 0 | 0.008545937201 | 0 |
| MARD_SAMEA104692614_REFG_MMP10469261 | 5 | -0.3534029089 | Kiloniellales | Kiloniella | Kiloniella sp9003 | 0 | 0 | 0.03171919434 | 5.07E-07 | 0.006811153602 | 0.9834207676 | 0 | 0.03940315424 | 0 | 0 | 1.08E-04 |
| MARD_SAMEA1059311_REFG_MMP1059311 | 2 | -0.321438509 | Rhodobacterales | Roseovarius | Roseovarius sp9 | 0 | 0 | 0.03579323195 | 1.31E-04 | 7.13E-04 | 0.02373371211 | 0 | 0.9999998131 | 0 | 0 | 0 |
| MARD_SAMEA18850168_REFG_MMP18850168 | 2 | -0.6216377052 | Enterobacterales | Vibrio | Vibrio quintilis | 0 | 0.1139922427 | 0.2482515208 | 1.21E-04 | 0.06007162276 | 4.08E-06 | 0 | 0.06930740609 | 0 | 0 | 0 |
| MARD_SAMEA2271987_REFG_MMP2271987 | 5 | -0.1946781587 | Mycobacteriales | Rhodococcus | Rhodococcus sp | 0.006417356876 | 0 | 0.06300309923 | 0.00359524655 | 0.03309642852 | 1.000000163 | 0.04262198874 | 0.05978952753 | 0 | 0 | 0 |
| MARD_SAMEA2272100_REFG_MMP2272100 | 2 | -0.5193179712 | Enterobacterales | Vibrio | Vibrio harveyi | 0 | 0 | 0.0644573853 | 3.90E-05 | 0.003132558562 | 0.003141167375 | 0 | 0.06217339115 | 0 | 2.45E-05 | 0 |
| MARD_SAMEA2272178_REFG_MMP2272178 | 5 | -0.204571548 | Streptomycetales | Streptomyces | Streptomyces sp | 0.01889924702 | 0 | 0.04611566489 | 5.67E-05 | 0.1437270367 | 0.9999998122 | 0 | 0.06143072088 | 0.006747021796 | 2.03E-05 | 0 |
| MARD_SAMEA2272395_REFG_MMP2272395 | 6 | -0.1935804027 | Cytophagales | Fibrisoma | Fibrisoma limi | 0 | 0 | 0.03157813522 | 2.58E-06 | 0.9501948454 | 5.12E-05 | 0 | 0.07010969719 | 0 | 0 | 0.02490518605 |
| MARD_SAMEA2272452_REFG_MMP2272452 | 1 | -0.2028214005 | Xanthomonadale | Stenotrophomona | Stenotrophomona | 0 | 0 | 0.9837623428 | 7.38E-05 | 0.05002468645 | 0.9833333333 | 0 | 0.0742833505 | 0 | 0.02221597484 | 0 |
| MARD_SAMEA2272636_REFG_MMP2272636 | 2 | -0.6865262318 | Enterobacterales | Pseudoalteromor | Pseudoalteromor | 0 | 0 | 0.04265243498 | 2.37E-05 | 0.009130898746 | 5.49E-04 | 1.19E-04 | 0.09238122703 | 0 | 1.90E-04 | 2.37E-05 |
| MARD_SAMEA2272788_REFG_MMP2272788 | 5 | -0.4465458606 | Pseudomonadale | Pseudomonas_E | Pseudomonas_E | 0 | 0 | 0.03875839459 | 4.23E-05 | 0.3404558096 | 1.000000007 | 0 | 0.06075357495 | 0 | 0 | 0 |
| MARD_SAMEA2591298_REFG_MMP2591298 | 2 | -0.3800367011 | Enterobacterales | Vibrio | Vibrio crassostre | 0 | 0 | 0.1509889302 | 1.86E-05 | 0.001915820633 | 8.33E-07 | 0 | 0.06287212213 | 0 | 1.86E-05 | 0 |
| MARD_SAMEA2594004_REFG_MMP2594004 | 2 | -0.3770659997 | Enterobacterales | Vibrio | Vibrio coralliirubr | 0 | 0 | 0.07216297994 | 3.60E-05 | 0.003185724892 | 0 | 0 | 0.05584762375 | 0 | 4.26E-05 | 0 |
| MARD_SAMEA2752412_REFG_MMP2752412 | 2 | -0.6053785479 | Enterobacterales | Aeromonas | Aeromonas salm | 0 | 0 | 0.02234833518 | 9.76E-05 | 0.009599089321 | 1.71E-06 | 0 | 0.06759883056 | 0 | 0 | 0 |
| MARD_SAMEA2752415_REFG_MMP2752415 | 2 | -0.6502133139 | Enterobacterales | Aeromonas | Aeromonas pisci | 0 | 0 | 0.1810337183 | 4.75E-05 | 0.0032161878 | 0.1666679684 | 0 | 0.1721495448 | 0 | 0 | 0 |
| MARD_SAMEA3221090_REFG_MMP3221090 | 2 | -0.6941302748 | Enterobacterales | Shewanella | Shewanella alga | 0 | 0 | 0.04127160722 | 9.14E-05 | 5.36E-05 | 0.1933334793 | 0 | 0.1086045849 | 0 | 7.62E-04 | 0 |
| MARD_SAMEA3297057_REFG_MMP3297057 | 2 | -0.1306121147 | Mycobacteriales | Rhodococcus | Rhodococcus sp | 0 | 0 | 0.01849614769 | 9.49E-06 | 0.08647830567 | 1.14E-04 | 0 | 0.04675867233 | 0 | 0 | 0 |
| MARD_SAMEA3355991_REFG_MMP3355991 | 2 | -0.4869576129 | Enterobacterales | Aliivibrio | Aliivibrio fischeri | 0 | 0 | 0.04494857285 | 9.62E-05 | 0.003377657371 | 0 | 0 | 0.007778231383 | 0 | 0 | 0 |
| MARD_SAMEA3393042_REFG_MMP3393042 | 2 | -0.4567235691 | Rhodobacterales | Phaeobacter | Phaeobacter itali | 0 | 0 | 0.0245496494 | 6.58E-06 | 0.001571442442 | 2.04E-07 | 0 | 0.05147798119 | 0 | 5.59E-05 | 0 |
| MARD_SAMEA3492688_REFG_MMP3492688 | 2 | -0.2646406183 | Rhizobiales | Roseibium | Roseibium aggre | 0 | 0.04275541484 | 0.05229219211 | 1.11E-05 | 0.003617461689 | 5.81E-06 | 0 | 0.03859938888 | 0 | 0 | 0.007224017654 |
| MARD_SAMEA3493278_REFG_MMP3493278 | 2 | -0.2198840726 | Rhizobiales | Roseibium | Roseibium albun | 0 | 0 | 0.06685623624 | 9.52E-05 | 0.003384338245 | 5.02E-06 | 0 | 0.07967578007 | 0 | 8.04E-04 | 0 |
| MARD_SAMEA3495629_REFG_MMP3495629 | 2 | -0.3673484724 | Rhizobiales | Roseibium | Roseibium albun | 3.47E-05 | 0 | 0.01581251855 | 3.19E-06 | 0.0905128943 | 0.002116152815 | 0.02283609158 | 0.02008805815 | 0 | 1.27E-05 | 0 |
| MARD_SAMEA3495742_REFG_MMP3495742 | 2 | -0.2890211829 | Rhizobiales | Roseibium | Roseibium alexa | 0 | 0 | 0.09859113007 | 3.76E-06 | 1.48E-04 | 5.10E-05 | 0 | 0.1052834928 | 0 | 0.002638602362 | 0.006513030877 |
| MARD_SAMEA3500511_REFG_MMP3500511 | 2 | -0.1803388503 | Pseudomonadale | Alcanivorax | Alcanivorax xeno | 1.03E-04 | 0 | 0.03272421837 | 5.30E-05 | 0.002857593488 | 0.00590868647 | 0 | 0.1083581136 | 4.79E-04 | 0.001176089748 | 0 |
| MARD_SAMEA3528420_REFG_MMP3528420 | 5 | -0.305216885 | Rhodobacterales | Ruegeria | Ruegeria atlantic | 0 | 0 | 4.97E-04 | 4.26E-07 | 0 | 0.98333318 | 0 | 0.01675263831 | 0 | 7.44E-05 | 0 |
| MARD_SAMEA3528422_REFG_MMP3528422 | 8 | -0.4336358707 | Rhodobacterales | Shimia | Shimia marina | 0 | 0 | 1.000000156 | 0.9833345152 | 0.006086050454 | 1.000000156 | 0 | 0.04824234338 | 0 | 0.002422855279 | 0 |
| MARD_SAMEA3531082_REFG_MMP3531082 | 2 | -0.3993315327 | Rhodobacterales | Thalassobius | Thalassobius aut | 0.002751960455 | 0 | 0.1136080888 | 1.96E-04 | 0.07333529078 | 0.007462911277 | 0 | 0.1311539827 | 0 | 8.02E-04 | 0.001510185646 |
| MARD_SAMEA3532031_REFG_MMP3532031 | 2 | -0.3514703293 | Rhodobacterales | Thalassobius | Thalassobius me | 0 | 0 | 0.09447027615 | 1.71E-04 | 0.07068041342 | 0 | 0 | 0.04410185209 | 0 | 7.45E-04 | 0 |
| MARD_SAMEA3532112_REFG_MMP3532112 | 8 | -0.364899689 | Rhodobacterales | Tropicibacter | Tropicibacter nap | 0 | 0 | 0.03746902833 | 0.9999998267 | 0.1454262071 | 0.9999998267 | 0 | 0.04362308026 | 0 | 7.81E-04 | 0.0272697961 |
| MARD_SAMEA3535146_REFG_MMP3535146 | 5 | -0.3534733594 | Rhodobacterales | Pseudophaeobac | Pseudophaeobac | 1.36E-06 | 0 | 0.2017914501 | 8.46E-05 | 0.006443131263 | 1 | 0 | 0.04614541932 | 0 | 8.28E-04 | 0.04084127002 |
| MARD_SAMEA3539551_REFG_MMP3539551 | 2 | -0.3370483487 | Rhodobacterales | Leisingera | Leisingera aquae | 0 | 0 | 0.03085180345 | 2.47E-05 | 0.03331483965 | 0.0118893832 | 0.01831492298 | 0.07005328839 | 0 | 0.004132897599 | 0 |
| MARD_SAMEA3542132_REFG_MMP3542132 | 2 | -0.3083161575 | Rhodobacterales | Ruegeria | Ruegeria denitrifi | 0 | 0 | 0.04014289252 | 9.95E-06 | 0.0149593147 | 0.001788643201 | 0 | 0.06641008549 | 0 | 7.85E-04 | 0 |
| MARD_SAMEA3542137_REFG_MMP3542137 | 1 | -0.4301239815 | Rhodobacterales | Shimia | Shimia thalassica | 0 | 0 | 0.9999997512 | 3.68E-05 | 0.02048946474 | 0.9999997512 | 0 | 0.06784979569 | 0 | 8.16E-04 | 0.03670375879 |
| MARD_SAMEA3542246_REFG_MMP3542246 | 2 | -0.4825700829 | Rhodobacterales | Cognatishimia | Cognatishimia ac | 0 | 0 | 0.1855577434 | 1.34E-04 | 0.01217499027 | 0 | 0 | 0.07117873957 | 0 | 7.78E-04 | 4.87E-06 |
| MARD_SAMEA3872802_REFG_MMP3872802 | 2 | -0.5288299759 | Enterobacterales | Enterovibrio | Enterovibrio cele | 0 | 0.1146097647 | 0.06810896977 | 4.16E-05 | 0 | 0.003304962388 | 0 | 0.06470719824 | 0 | 5.22E-04 | 0 |
| MARD_SAMEA3872803_REFG_MMP3872803 | 2 | -0.5308809348 | Enterobacterales | Enterovibrio | Enterovibrio mari | 0 | 0 | 0.05485186689 | 2.95E-05 | 5.71E-05 | 7.73E-08 | 0 | 0.159338861 | 0 | 4.75E-05 | 0 |
| MARD_SAMEA3925393_REFG_MMP3925393 | 2 | -0.3305305095 | Enterobacterales | Shewanella | Shewanella sp00 | 0 | 0 | 0.04870581367 | 9.49E-05 | 0 | 0 | 0 | 0.0415724814 | 0 | 1.28E-05 | 0 |

| | | | | | | | | | | | | | | | | |
|---|---|---|---|---|---|---|---|---|---|---|---|---|---|---|---|---|
| MARD_SAMEA4019367_REFG_MMP4019367 | 5 | -0.5658263888 | Pseudomonadale | Marinomonas | Marinomonas aq | 0 | 0 | 0.0336741228 | 1.01E-04 | 0 | 1 | 0 | 0.03544931239 | 0 | 0.004023930696 | 0 |
| MARD_SAMEA4029000_REFG_MMP4029000 | 5 | -0.4685043231 | Pseudomonadale | Marinomonas | Marinomonas ga | 0 | 0 | 0.03564803443 | 1.38E-04 | 0.00920983887 | 1 | 0 | 0.083280154 | 0 | 0.003635610097 | 0 |
| MARD_SAMEA4034450_REFG_MMP4034450 | 6 | -0.1889065194 | Flavobacteriales | Aquimarina | Aquimarina sp90 | 0 | 0 | 0.07983866807 | 1.10E-04 | 1.000000229 | 0 | 0 | 0.06906776698 | 0 | 0 | 0 |
| MARD_SAMEA44539918_REFG_MMP44539918 | 7 | -0.1758586292 | Enterobacterales | Phocoenobacter | Phocoenobacter | 0 | 0 | 0.9999998791 | 0.9999998791 | 7.01E-05 | 0.05389454592 | 0 | 0.9999998791 | 0 | 0.9999998791 | 0 |
| MARD_SAMEA4520411_REFG_MMP4520411 | 2 | -0.3934380287 | Enterobacterales | Moritella | Moritella viscosa | 0 | 0 | 0.04084381088 | 4.81E-05 | 0 | 5.25E-06 | 0 | 0.05097451406 | 0 | 0 | 0 |
| MARD_SAMEA4644770_REFG_MMP4644770 | 4 | -0.1905714101 | Flavobacteriales | Aequorivita | Aequorivita lipoly | 0 | 0 | 1 | 0 | 2.94E-07 | 0.007748643026 | 0 | 0.2825593741 | 0 | 0.001610961584 | 0.0302310679 |
| MARD_SAMEA859881_REFG_MMP859881 | 2 | -0.3752705991 | Enterobacterales | Yersinia | Yersinia massilie | 0 | 0 | 0.05365531549 | 1.25E-04 | 0.003278517079 | 0.2000012161 | 0 | 0.01982505658 | 0 | 0 | 0 |
| MARD_SAMEA859932_REFG_MMP859932 | 2 | -0.3679264291 | Enterobacterales | Yersinia | Yersinia kristense | 0 | 0 | 0.05211321389 | 2.61E-05 | 0.006555653042 | 1.24E-05 | 0 | 0.03047137603 | 0 | 0 | 0 |
| MARD_SAMEA859946_REFG_MMP859946 | 2 | -0.3497724675 | Enterobacterales | Yersinia | Yersinia intermec | 0 | 0 | 0.2558738729 | 5.67E-05 | 3.22E-04 | 0.1270828018 | 0 | 0.06283762894 | 0 | 0 | 0 |
| MARD_SAMN00622969_REFG_MMP00622969 | 5 | -0.8417074151 | Bacillales | Bacillus_AB | Bacillus_AB infar | 0.02033084879 | 0 | 0.03327932247 | 0.001220253468 | 0.004733764693 | 0.7669085637 | 0 | 0.1294487238 | 0 | 1.35E-05 | 0.7666668107 |
| MARD_SAMN00622971_REFG_MMP00622971 | 2 | -0.07624930354 | Actinomycetales | Knoellia | Knoellia sp00015 | 0 | 0 | 0.04728038872 | 1.51E-04 | 0.2122591052 | 0.01666910108 | 0 | 0.08437011727 | 0 | 7.64E-04 | 0 |
| MARD_SAMN00622973_REFG_MMP00622973 | 5 | -0.4187165825 | Enterobacterales | Pseudoalteromor | Pseudoalteromor | 0 | 0 | 0.02682751261 | 8.35E-07 | 0 | 0.9666666127 | 0 | 0.1026333499 | 0 | 0 | 0 |
| MARD_SAMN00622974_REFG_MMP00622974 | 2 | -0.4667530557 | Enterobacterales | Psychromonas | Psychromonas s | 0 | 0 | 0.1350480826 | 0 | 0.001482906923 | 0.02619217584 | 0 | 0.154566866 | 0 | 0 | 0 |
| MARD_SAMN02261247_REFG_MMP02261247 | 3 | -0.4046293215 | Burkholderiales | Methylophilus | Methylophilus sp | 0 | 0 | 0.05714637017 | 9.14E-05 | 0.4478301465 | 1 | 0 | 0.04565809433 | 0 | 1 | 0 |
| MARD_SAMN02261258_REFG_MMP02261258 | 4 | -0.167351531 | Rhodobacterales | Paracoccus | Paracoccus amir | 0 | 0.06128505399 | 1.000000195 | 1.11E-04 | 0.07714027929 | 2.24E-05 | 0 | 0.06948308229 | 0 | 8.29E-04 | 0.03747259118 |
| MARD_SAMN02261354_REFG_MMP02261354 | 2 | -0.05931452992 | Cyanobacteriales | Xenococcus | Xenococcus sp0 | 0 | 0 | 0.03854446656 | 4.29E-05 | 0.009330200008 | 2.67E-04 | 0 | 0.1896811297 | 0 | 0 | 0 |
| MARD_SAMN02436080_REFG_MMP02436080 | 4 | -0.3364762708 | Rhodobacterales | Roseovarius | Roseovarius nub | 5.33E-04 | 0 | 0.9677542406 | 0.001237546936 | 4.78E-04 | 1.37E-05 | 5.52E-04 | 0.03732555846 | 0 | 1.19E-04 | 1.16E-04 |
| MARD_SAMN02436081_REFG_MMP02436081 | 2 | -0.02468374431 | Rhodobacterales | Pseudooceanicol | Pseudooceanicol | 0 | 0 | 0.0462970835 | 1.41E-04 | 0.00344102066 | 6.28E-06 | 0 | 0.0172835818 | 0 | 0.000814606837 | 0.04786613033 |
| MARD_SAMN02436084_REFG_MMP02436084 | 1 | -0.06623264491 | Rhizobiales | Nitrobacter | Nitrobacter sp00 | 0 | 0 | 1.000000205 | 7.72E-05 | 0.01804718108 | 1.000000205 | 0 | 0.2326724083 | 0 | 0 | 0 |
| MARD_SAMN02436087_REFG_MMP02436087 | 2 | -0.443238159 | Enterobacterales | Photobacterium | Photobacterium p | 0 | 0 | 0.05419797451 | 2.23E-05 | 0.04295075595 | 0.002680143695 | 0 | 0.1025075086 | 0 | 1.36E-05 | 0 |
| MARD_SAMN02436090_REFG_MMP02436090 | 2 | -0.02826899975 | Cyanobacteriales | Limnoraphis | Limnoraphis sp0 | 0 | 0 | 0.01306353706 | 1.78E-04 | 0.1875764683 | 3.01E-08 | 0 | 0.08831760133 | 0 | 1.65E-07 | 0 |
| MARD_SAMN02436096_REFG_MMP02436096 | 2 | -0.4159735319 | Enterobacterales | Shewanella | Shewanella bent | 0 | 0 | 0.1189020211 | 9.32E-05 | 0.2176018899 | 2.60E-08 | 0 | 0.06274983469 | 0 | 7.41E-04 | 0 |
| MARD_SAMN02436105_REFG_MMP02436105 | 4 | -0.2026256023 | Rhodobacterales | Salipiger | Salipiger bermud | 0 | 0.01043639324 | 1.000000179 | 1.35E-06 | 0.03310452292 | 0.06666665728 | 0 | 0.05469837584 | 0.009203313365 | 4.06E-05 | 0.002954106851 |
| MARD_SAMN02436109_REFG_MMP02436109 | 1 | -0.2054900773 | Rhodobacterales | Yoonia | Yoonia vestfolder | 0 | 0 | 0.9999998365 | 0.01713472717 | 0 | 0.9999998365 | 0 | 0.9999998365 | 0 | 0.004449529684 | 0.03402129556 |
| MARD_SAMN02436110_REFG_MMP02436110 | 3 | -0.1882928419 | Rhodobacterales | Maritimibacter | Maritimibacter all | 8.15E-06 | 0 | 0.1182072249 | 6.99E-05 | 0.00366790793 | 1 | 0 | 0.09718238776 | 0 | 1 | 0 |
| MARD_SAMN02436111_REFG_MMP02436111 | 2 | -0.2515804367 | Flavobacteriales | Leeuwenhoekiell | Leeuwenhoekiell | 0 | 0 | 0.1606766278 | 1.43E-04 | 0 | 0 | 0 | 0.0733744708 | 0 | 3.34E-06 | 0.03733038102 |
| MARD_SAMN02436112_REFG_MMP02436112 | 3 | -0.1749394211 | Pirellulales | Blastopirellula | Blastopirellula ma | 4.96E-06 | 0 | 0.03132779822 | 8.97E-05 | 0.1172752153 | 1 | 0 | 0.03007858816 | 0 | 1 | 0 |
| MARD_SAMN02436113_REFG_MMP02436113 | 5 | -0.1890714875 | Pseudomonadale | Congregibacter | Congregibacter li | 0 | 0.002666991523 | 0.03187797041 | 3.96E-07 | 0.09248491185 | 0.7669189981 | 0 | 0.06000216169 | 0 | 2.46E-04 | 2.70E-08 |
| MARD_SAMN02436114_REFG_MMP02436114 | 2 | -0.124525585 | Flavobacteriales | Polaribacter | Polaribacter irger | 0 | 3.17E-04 | 0.1251310289 | 0 | 0.02858798962 | 0 | 0 | 0.05475086556 | 0 | 0 | 0 |
| MARD_SAMN02436115_REFG_MMP02436115 | 2 | -0.5284709324 | Enterobacterales | Photobacterium | Photobacterium s | 0 | 0 | 0.2007340014 | 1.78E-05 | 0.003255924309 | 1.10E-07 | 0 | 0.06633538749 | 0 | 3.01E-05 | 0 |
| MARD_SAMN02436126_REFG_MMP02436126 | 2 | -0.1055528546 | Rhizobiales | Fulvimarina | Fulvimarina pela | 0 | 0 | 0.1288958514 | 2.15E-04 | 0.01228140207 | 0.01809234685 | 7.86E-05 | 0.01353060428 | 0 | 7.38E-04 | 0.01036896891 |
| MARD_SAMN02436127_REFG_MMP02436127 | 3 | -0.2038239731 | Mariprofundales | Mariprofundus | Mariprofundus fe | 0 | 0 | 1.000000273 | 9.38E-05 | 0.2289772675 | 1.000000273 | 0 | 0.0340094377 | 0 | 1.000000273 | 0 |
| MARD_SAMN02436130_REFG_MMP02436130 | 2 | -0.1800742169 | Cytophagales | Microscilla | Microscilla marin | 0 | 0 | 0.02509635546 | 0.9833349725 | 0.1417073637 | 7.68E-05 | 0 | 0.05889911138 | 0 | 0 | 3.41E-04 |
| MARD_SAMN02436132_REFG_MMP02436132 | 3 | -0.3343143112 | Pseudomonadale | Bermanella | Bermanella maris | 0 | 0 | 0.999999878 | 5.82E-05 | 0.003019707669 | 0.999999878 | 0 | 0.0605387917 | 0 | 0.999999878 | 0 |
| MARD_SAMN02436135_REFG_MMP02436135 | 5 | -0.2460889893 | Sphingomonadal | Erythrobacter | Erythrobacter sp | 0 | 0 | 0.02682990141 | 0.491667615 | 0 | 1.000000313 | 0 | 0.07872931724 | 0 | 0.007114358274 | 0 |
| MARD_SAMN02436137_REFG_MMP02436137 | 2 | -0.4022402178 | Rhodobacterales | Pseudophaeobac | Pseudophaeobac | 0 | 0 | 0.04544653499 | 1.45E-04 | 0.1146739739 | 5.60E-06 | 0 | 0.06557600823 | 0 | 8.26E-04 | 0.03413367017 |
| MARD_SAMN02436140_REFG_MMP02436140 | 5 | -0.4308789412 | Pseudomonadale | Reinekea | Reinekea blande | 0 | 0 | 0.03479270984 | 7.48E-05 | 0.0121713948 | 0.9999998808 | 0 | 0.08209762975 | 0 | 7.94E-04 | 0 |
| MARD_SAMN02436141_REFG_MMP02436141 | 5 | -0.05396542398 | Nitrococcales | Nitrococcus | Nitrococcus mob | 0 | 0.003436208324 | 0.03281212221 | 3.08E-06 | 5.28E-04 | 0.9667072034 | 0 | 0.03577801824 | 0 | 0.004730824216 | 0 |
| MARD_SAMN02436142_REFG_MMP02436142 | 2 | -0.05390421856 | Rhodobacterales | Oceanicola | Oceanicola granu | 4.18E-04 | 0.001449669009 | 0.03120754029 | 1 | 0.1339511789 | 0.02085024378 | 0 | 0.07895030871 | 0 | 0.001640713101 | 0.00119158638 |
| MARD_SAMN02436148_REFG_MMP02436148 | 1 | -0.3359835078 | Lentisphaerales | Lentisphaera | Lentisphaera ara | 0 | 0 | 1 | 1.45E-04 | 0.123155218 | 1 | 0 | 0.08215694654 | 0 | 0 | 0 |
| MARD_SAMN02436150_REFG_MMP02436150 | 2 | -0.4528764542 | Enterobacterales | Moritella | Moritella sp0001 | 0 | 0 | 0.198224113 | 6.12E-06 | 1.27E-04 | 0.005453184219 | 0 | 0.06779085061 | 0 | 5.39E-06 | 0 |
| MARD_SAMN02436155_REFG_MMP02436155 | 2 | -0.1651087272 | Pseudomonadale | Oceanicoccus | Oceanicoccus sp | 0 | 0 | 0.05267707504 | 1.95E-06 | 1.19E-04 | 3.20E-08 | 0 | 0.0738882586 | 0 | 0.007156958049 | 0.05325144285 |
| MARD_SAMN02436161_REFG_MMP02436161 | 2 | -0.1196161893 | Rhizobiales | Aurantimonas | Aurantimonas ma | 0 | 0 | 0.1892482569 | 0.03350037409 | 0.05979016805 | 4.88E-05 | 0 | 0.07265833899 | 0 | 8.01E-04 | 0.0362980219 |
| MARD_SAMN02436164_REFG_MMP02436164 | 2 | -0.1208788853 | Rhodobacterales | Roseovarius | Roseovarius sp0 | 2.60E-04 | 0 | 0.06671307504 | 0.01669897201 | 0.02695671147 | 1.08E-04 | 2.20E-04 | 0.983808961 | 0 | 0.004309180396 | 0 |
| MARD_SAMN02436165_REFG_MMP02436165 | 2 | -0.3805279925 | Pseudomonadale | Marinomonas | Marinomonas sp | 0 | 0 | 0.09643956597 | 1.00E-04 | 0.008514192309 | 0.001431516538 | 0 | 0.05753571818 | 0 | 7.72E-04 | 0 |
| MARD_SAMN02436179_REFG_MMP02436179 | 5 | -0.05150128963 | Nitrosococcales | Nitrosococcus | Nitrosococcus oc | 0 | 0 | 0.02493291038 | 1.43E-04 | 0.08627751465 | 0.9999996485 | 0 | 0.03855645282 | 0 | 0.01683822043 | 0 |
| MARD_SAMN02436225_REFG_MMP02436225 | 2 | -0.2525506591 | Rhodobacterales | Planktotalea | Planktotalea sp0 | 0.02282844299 | 0 | 0.02921591402 | 1.37E-04 | 0.002509576798 | 0.01120646081 | 0.02282844299 | 1 | 2.89E-05 | 0.001940456317 | 0 |
| MARD_SAMN02440876_REFG_MMP02440876 | 5 | -0.2377809864 | Enterobacterales | Glaciecola | Glaciecola pallidu | 0 | 0 | 0.1164548997 | 9.38E-05 | 0.1429804201 | 0.9999999846 | 0 | 0.1838656317 | 0 | 6.75E-05 | 0 |
| MARD_SAMN02441415_REFG_MMP02441415 | 2 | -0.1219553472 | Cyanobacteriales | Crocosphaera | Crocosphaera wa | 0 | 0 | 0.03935155737 | 1.64E-04 | 0.2297972413 | 0.00343430299 | 0 | 0.04181274027 | 0 | 0 | 0 |
| MARD_SAMN02441806_REFG_MMP02441806 | 2 | -0.186338914 | Desulfuromonada | Desulfuromonas | Desulfuromonas | 0 | 0 | 0.06337839412 | 3.20E-07 | 0.004457769372 | 0.01810795818 | 0 | 0.2683122831 | 0 | 0.003935549133 | 0.02152560769 |
| MARD_SAMN02469978_REFG_MMP02469978 | 2 | -0.06169444043 | Bacteroidales | Marinilabilia | Marinilabilia salm | 0 | 0 | 0.04832888838 | 5.82E-06 | 0.07386447677 | 7.71E-05 | 0.001982934823 | 0.0619426394 | 0 | 0 | 4.13E-05 |
| MARD_SAMN02470171_REFG_MMP02470171 | 1 | -0.4732146887 | Enterobacterales | Pseudoalteromor | Pseudoalteromor | 0 | 0 | 0.7578456053 | 7.28E-05 | 0.004200438441 | 0.7666679157 | 0 | 0.05180383516 | 0 | 8.80E-04 | 6.26E-04 |
| MARD_SAMN02470761_REFG_MMP02470761 | 2 | -0.3205601834 | Enterobacterales | Catenovulum | Catenovulum aga | 0 | 0 | 0.04192752674 | 9.84E-05 | 0.007829883586 | 0 | 0 | 0.05405333708 | 0 | 0.007734790395 | 0.159289258 |
| MARD_SAMN02470762_REFG_MMP02470762 | 2 | -0.648861719 | Enterobacterales | Pseudoalteromor | Pseudoalteromor | 0 | 0 | 0.03482560843 | 8.77E-05 | 0.00328508801 | 0 | 0 | 0.04818510504 | 0 | 0 | 0 |
| MARD_SAMN02470816_REFG_MMP02470816 | 7 | -0.2440524478 | Pseudomonadale | Zhongshania | Zhongshania alip | 0.1607898555 | 0 | 0.02991155439 | 3.37E-05 | 0.001581865095 | 0.00328994283 | 0 | 0.03188383725 | 0 | 1 | 0.06517915253 |
| MARD_SAMN02603920_REFG_MMP02603920 | 2 | -0.09589808942 | Flavobacteriales | Polaribacter | Polaribacter sp0 | 0 | 0 | 0.5729530604 | 1.37E-04 | 0.04096544774 | 0 | 0 | 0.05358677325 | 0 | 0.001660096363 | 0 |
| MARD_SAMN02744645_REFG_MMP02744645 | 5 | -0.245543841 | Pseudomonadale | Stutzerimonas | Stutzerimonas xa | 0.01142108768 | 0.001739698148 | 0.09389844618 | 1.85E-05 | 0.07772657239 | 0.999999901 | 0.01128739416 | 0.07444936681 | 0 | 0.007336850153 | 0.038702848 |
| MARD_SAMN02745129_REFG_MMP02745129 | 2 | -0.6706763459 | Enterobacterales | Ferrimonas | Ferrimonas marir | 0 | 0 | 0.03431950027 | 3.25E-05 | 0.0297203769 | 0.004721494583 | 0 | 0.1083804941 | 0 | 8.09E-04 | 0 |
| MARD_SAMN02745135_REFG_MMP02745135 | 1 | -0.1135443311 | Tissierellales | Caloranaerobact | Caloranaerobact | 0 | 0 | 0.999999779 | 5.35E-05 | 0.2855926657 | 0.999999779 | 0 | 0.1851920366 | 0 | 0.001467522338 | 0 |
| MARD_SAMN02745148_REFG_MMP02745148 | 1 | -0.1664088522 | Pseudomonadale | Modicisalibacter | Modicisalibacter | 0 | 0 | 0.9999999978 | 4.05E-05 | 0.01307002053 | 0.9833333333 | 0 | 0.2333044806 | 0 | 8.36E-04 | 1.07E-04 |
| MARD_SAMN02745151_REFG_MMP02745151 | 1 | -0.3847814998 | Lachnospirales | Anaerotignum | Anaerotignum pr | 0 | 0 | 1 | 1.22E-04 | 0.01270352505 | 1 | 0 | 0.2139398775 | 0 | 0 | 0 |
| MARD_SAMN02745161_REFG_MMP02745161 | 2 | -0.1992058949 | Desulfovibrionale | Halodesulfovibric | Halodesulfovibric | 0 | 0 | 0.1495435325 | 1.39E-04 | 0.006151757981 | 0 | 0 | 0.07113345834 | 0 | 7.72E-04 | 0.2397977341 |
| MARD_SAMN02745164_REFG_MMP02745164 | 4 | -0.02532250159 | Petrotogales | Marinitoga | Marinitoga hydro | 0 | 0 | 1.000000259 | 0.01691809332 | 0.02119986803 | 0 | 0.1674066682 | 0.0942152454 | 0 | 1.32E-04 | 0 |
| MARD_SAMN02745165_REFG_MMP02745165 | 2 | -0.1037311117 | Desulfuromonada | Malonomonas | Malonomonas ru | 0 | 0 | 0.01226205114 | 7.62E-05 | 0 | 1.28E-05 | 0 | 0.2934716182 | 0 | 7.95E-04 | 0.08543567678 |
| MARD_SAMN02745181_REFG_MMP02745181 | 2 | -0.4029721038 | Verrucomicrobial | Rubritalea | Rubritalea squale | 0 | 0 | 0.01164475027 | 9.76E-05 | 0.003529334371 | 0 | 0 | 1 | 0 | 0.01768689696 | 0.07300689819 |
| MARD_SAMN02745216_REFG_MMP02745216 | 2 | -0.05097239706 | Desulfobacterale | Desulfatibacillum | Desulfatibacillum | 0 | 0 | 0.03907492436 | 1.43E-06 | 0.006390177161 | 8.14E-07 | 6.03E-04 | 0.05852993468 | 0 | 1.36E-05 | 0 |
| MARD_SAMN02745220_REFG_MMP02745220 | 1 | -0.09976221666 | Desulfobulbales | Desulfopila | Desulfopila aestu | 0 | 0 | 0.9839163968 | 1.09E-04 | 0.06937150004 | 0.9999998369 | 0 | 0.03727573017 | 0 | 7.62E-04 | 0 |
| MARD_SAMN02745246_REFG_MMP02745246 | 6 | -0.1601084336 | Flavobacteriales | Leeuwenhoekiell | Leeuwenhoekiell | 0 | 0 | 0.03289430802 | 4.48E-06 | 0.9837396728 | 0 | 0 | 0.1012303578 | 0 | 2.94E-08 | 0.03326322179 |
| MARD_SAMN02745681_REFG_MMP02745681 | 2 | -0.2154066031 | Pseudomonadale | Teredinibacter | | 0 | 0 | 0.1478892379 | 1.40E-06 | 0.05709247134 | 7.09E-04 | 0.008124306792 | 0.05477598343 | 0 | 0.02675595722 | 0.05219606493 |
| MARD_SAMN02745724_REFG_MMP02745724 | 4 | -0.524237963 | Enterobacterales | Pseudoalteromor | Pseudoalteromor | 0 | 0 | 0.7596806352 | 9.75E-05 | 0 | 0.5833340936 | 0 | 0.06018967057 | 0 | 5.35E-04 | 0 |
| MARD_SAMN02745729_REFG_MMP02745729 | 2 | -0.2776872906 | Pseudomonadale | Marinobacterium | Marinobacterium | 6.35E-05 | 5.18E-04 | 0.0437823604 | 1.14E-04 | 0.1771007591 | 2.95E-06 | 0.01666666667 | 0.04538367294 | 0 | 7.73E-04 | 0 |
| MARD_SAMN02745751_REFG_MMP02745751 | 2 | -0.2018693083 | Tissierellales | Dethiosulfatibact | Dethiosulfatibact | 0.1734640438 | 0 | 0.04834042506 | 1.52E-04 | 0.07595556322 | 0.04242413583 | 0.001658452756 | 0.02568286782 | 1.08E-06 | 1.82E-04 | 0.003262520236 |
| MARD_SAMN02745753_REFG_MMP02745753 | 2 | -0.2153694964 | Pseudomonadale | Marinomonas | Marinomonas po | 0 | 0 | 0.03193668552 | 5.41E-05 | 0.01879036733 | 0.1966706029 | 0 | 0.05647675953 | 0 | 7.94E-04 | 0 |

| | | | | | | | | | | | | | | | | |
|---|---|---|---|---|---|---|---|---|---|---|---|---|---|---|---|---|
| MARD_SAMN02745781_REFG_MMP02745781 | 2 | -0.6826229404 | Enterobacterales | Vibrio | Vibrio gazogenes | 0 | 0 | 0.05816303292 | 1.84E-05 | 0.001856243735 | 0 | 0 | 0.06885612824 | 0 | 0 | 0 |
| MARD_SAMN02745824_REFG_MMP02745824 | 2 | -0.1397382347 | Sphingomonadal | Parasphingorhab | Parasphingorhab | 0 | 0 | 0.0371340885 | 1.66E-06 | 0.337552217 | 0.01666666667 | 0 | 0.07336458796 | 0 | 1.32E-05 | 5.49E-05 |
| MARD_SAMN02745912_REFG_MMP02745912 | 5 | -0.05738578172 | Peptostreptococc | Paramaledivibac | Paramaledivibac | 0.02079871237 | 0.03007169659 | 0.03627444763 | 2.18E-04 | 0.1711245694 | 0.9999996291 | 0 | 0.06085995112 | 0.08203475751 | 0.006460889919 | 0 |
| MARD_SAMN02745945_REFG_MMP02745945 | 5 | -0.4003545739 | Peptostreptococc | Peptoclostridium | Peptoclostridium | 0 | 0 | 0.0867551192 | 2.74E-04 | 0.2663093922 | 0.7499997493 | 0 | 0.06215727055 | 0 | 4.75E-05 | 0.05551303745 |
| MARD_SAMN02927921_REFG_MMP02927921 | 6 | -0.3138694053 | Flavobacteriales | Sinomicrobium | Sinomicrobium o | 0 | 0 | 0.0263195193 | 9.80E-07 | 0.9999999989 | 0.004426976493 | 0 | 0.1977484543 | 0 | 0 | 1.67E-04 |
| MARD_SAMN02927937_REFG_MMP02927937 | 4 | -0.3301190089 | Flavobacteriales | Flavobacterium | Flavobacterium r | 0 | 0 | 1 | 6.26E-05 | 1 | 0 | 0 | 0.2674534703 | 0 | 0 | 0 |
| MARD_SAMN02927977_REFG_MMP02927977 | 5 | -0.3639729373 | Rhodospirillales_ | Thalassospira | Thalassospira sp | 0 | 0.003069658981 | 0.04099744885 | 9.85E-05 | 0 | 0.9999998134 | 0 | 0.1633987013 | 0 | 0.003046149558 | 0 |
| MARD_SAMN02927978_REFG_MMP02927978 | 5 | -0.4618741029 | Rhodospirillales_ | Thalassospira | Thalassospira po | 0 | 0 | 0.03193478813 | 9.77E-05 | 0.003336875817 | 1 | 0 | 0.1235798472 | 0 | 7.92E-04 | 2.26E-06 |
| MARD_SAMN02927979_REFG_MMP02927979 | 5 | -0.2911337505 | Rhodospirillales_ | Thalassospira | Thalassospira luc | 0 | 0 | 0.03414833817 | 9.48E-05 | 0.0900179238 | 0.9999998111 | 0 | 0.05749177525 | 0 | 7.71E-04 | 8.82E-04 |
| MARD_SAMN02927981_REFG_MMP02927981 | 2 | -0.4088147206 | Rhodospirillales_ | Thalassospira | Thalassospira pr | 0 | 0 | 0.03960922117 | 2.98E-06 | 0.001655344386 | 2.34E-05 | 0 | 0.1479084803 | 6.30E-05 | 1.29E-05 | 5.87E-05 |
| MARD_SAMN02927983_REFG_MMP02927983 | 5 | -0.3036788303 | Rhodospirillales_ | Thalassospira | Thalassospira pr | 0 | 0 | 0.03884764925 | 6.73E-05 | 0.1044317111 | 1 | 0 | 0.05014690476 | 0 | 7.87E-04 | 7.56E-08 |
| MARD_SAMN02927984_REFG_MMP02927984 | 5 | -0.3974660944 | Rhodospirillales_ | Thalassospira | Thalassospira pr | 0 | 0.004375916732 | 0.04111650245 | 1.10E-04 | 0 | 1 | 0 | 0.1707756021 | 0 | 7.96E-04 | 0 |
| MARD_SAMN03079717_REFG_MMP03079717 | 2 | -0.5472609599 | Enterobacterales | Salmonella | Salmonella enter | 0 | 0 | 0.08452901768 | 4.82E-04 | 0 | 0.06670750207 | 0 | 0.140633761 | 0 | 0 | 0 |
| MARD_SAMN03080594_REFG_MMP03080594 | 6 | -0.2705381411 | Flavobacteriales | Arenibacter | Arenibacter palla | 0 | 0 | 0.02720073413 | 8.15E-05 | 1 | 0.002742407637 | 0 | 0.05380118701 | 0 | 0 | 0 |
| MARD_SAMN03080599_REFG_MMP03080599 | 5 | -0.2276651461 | Peptostreptococc | Acidaminobacter | Acidaminobacter | 0.1776946078 | 0 | 0.1012807087 | 2.37E-04 | 0.202757901 | 1 | 0 | 0.06246284096 | 0 | 9.46E-06 | 0.08060481996 |
| MARD_SAMN03080610_REFG_MMP03080610 | 2 | -0.06663656868 | Rhizobiales | Afifella | Afifella marina | 0.040000191 | 0 | 0.0398806221 | 3.29E-05 | 0.04364723354 | 0 | 0 | 0.05301142685 | 0 | 7.93E-04 | 0 |
| MARD_SAMN03080615_REFG_MMP03080615 | 2 | -0.4827934463 | Pseudomonadale | Amphritea | Amphritea atlanti | 0 | 0 | 0.3850967911 | 1.01E-04 | 0.07648552634 | 3.83E-05 | 0 | 0.06505651943 | 0 | 7.86E-04 | 0.02221336698 |
| MARD_SAMN03339836_REFG_MMP03339836 | 5 | -0.3632678129 | Rhodobacterales | Epibacterium | Epibacterium mo | 0 | 0 | 0.1861996041 | 1.35E-04 | 0.007867876527 | 1 | 0 | 0.02993991677 | 0 | 0.001543842051 | 2.45E-06 |
| MARD_SAMN03729479_REFG_MMP03729479 | 6 | 0.0212581145 | Rubrobacterales | Rubrobacter_A | Rubrobacter_A a | 0 | 0.009035408492 | 0.05239629861 | 1.04E-04 | 1.000000202 | 2.02E-06 | 0 | 0.0558893042 | 0 | 0 | 0.0374722642 |
| MARD_SAMN03732514_REFG_MMP03732514 | 6 | -0.07294994341 | Actinomycetales | Kocuria | Kocuria turfanens | 0 | 0 | 0.04830410318 | 2.54E-05 | 1 | 2.46E-05 | 0 | 0.07051882216 | 0 | 7.45E-04 | 0 |
| MARD_SAMN03764910_REFG_MMP03764910 | 4 | -0.3668326105 | Flavobacteriales | Chryseobacteriu | Chryseobacteriu | 0 | 0 | 1 | 1.19E-04 | 1 | 0.003747568691 | 3.19E-06 | 0.0762935839 | 0 | 0 | 0 |
| MARD_SAMN03941568_REFG_MMP03941568 | 1 | -0.3113695372 | Pseudomonadale | Halopseudomona | Halopseudomona | 0 | 0 | 0.9837436742 | 6.25E-05 | 0.1635251412 | 1.000000003 | 0 | 0.2053022483 | 0 | 2.84E-04 | 0.05589113932 |
| MARD_SAMN03981068_REFG_MMP03981068 | 2 | -0.5311784067 | Enterobacterales | Vibrio | Vibrio sp0026085 | 0 | 0 | 0.09655792162 | 1.20E-05 | 0.001762063891 | 0.003335427642 | 0 | 0.07817073644 | 6.02E-07 | 2.85E-04 | 0 |
| MARD_SAMN03999371_REFG_MMP03999371 | 5 | -0.149728435 | Streptomycetales | Streptomyces | Streptomyces sp | 0 | 0 | 0.03192092868 | 2.07E-05 | 0.1455070202 | 1 | 0 | 0.05295362853 | 0 | 8.01E-04 | 0 |
| MARD_SAMN03999384_REFG_MMP03999384 | 5 | -0.1892208421 | Streptomycetales | Streptomyces | Streptomyces sp | 0.007254654429 | 0 | 0.08492941793 | 3.33E-05 | 0.07108957415 | 1 | 0 | 0.06466119954 | 0.005719448493 | 4.56E-05 | 0 |
| MARD_SAMN03999393_REFG_MMP03999393 | 5 | -0.1625668688 | Streptomycetales | Streptomyces | Streptomyces sp | 0 | 0 | 0.02448446918 | 2.49E-05 | 0 | 0.9999997693 | 0 | 0.1268746549 | 0 | 8.02E-04 | 0 |
| MARD_SAMN04026261_REFG_MMP04026261 | 6 | -0.08457547453 | Streptomycetales | Streptomyces | Streptomyces ab | 0 | 0 | 0.04195150007 | 4.00E-05 | 0.9557897828 | 1.00000001 | 0 | 0.2010719052 | 0 | 8.28E-04 | 0.07307556292 |
| MARD_SAMN04219563_REFG_MMP04219563 | 2 | -0.837156703 | Enterobacterales | Vibrio | Vibrio vulnificus | 0 | 0 | 0.06803227103 | 3.93E-05 | 0.005303485738 | 0.05666856584 | 0 | 0.06071042257 | 0 | 2.39E-05 | 0 |
| MARD_SAMN04311937_REFG_MMP04311937 | 8 | -0.4985316218 | Lactobacillales | Streptococcus | Streptococcus pe | 0 | 0 | 1.000000067 | 1.000000067 | 0.2134595975 | 1.000000067 | 0.003129478217 | 1.000000067 | 0 | 0.004506909422 | 0 |
| MARD_SAMN04320402_REFG_MMP04320402 | 2 | -0.320769526 | Enterobacterales | Yersinia | Yersinia ruckeri | 0 | 0 | 0.2953334664 | 4.66E-05 | 0.006530074931 | 0.1489778444 | 0 | 0.08383774924 | 0 | 0.003944906987 | 0 |
| MARD_SAMN04390145_REFG_MMP04390145 | 6 | -0.05009408851 | Flavobacteriales | Polaribacter | Polaribacter vadi | 0 | 0 | 0.02575499032 | 1.07E-04 | 1.000000192 | 0.002793256857 | 0 | 0.07111241772 | 0 | 0 | 0.04689316072 |
| MARD_SAMN04390147_REFG_MMP04390147 | 4 | -0.1670591588 | Burkholderiales | Hydrogenophaga | Hydrogenophaga | 0 | 0 | 0.9999997323 | 3.00E-05 | 0.1249337118 | 4.82E-06 | 0 | 0.05218197651 | 0 | 0.004401992622 | 0 |
| MARD_SAMN04390149_REFG_MMP04390149 | 6 | -0.0889330412 | Flavobacteriales | Polaribacter | Polaribacter reich | 4.07E-08 | 0 | 0.06593142864 | 2.67E-06 | 0.9833333333 | 0.01666666667 | 0 | 0.07262673031 | 0 | 2.34E-05 | 0.2627367005 |
| MARD_SAMN04390154_REFG_MMP04390154 | 6 | -0.06089503983 | Flavobacteriales | Polaribacter | Polaribacter atrin | 0 | 0 | 0.04431183428 | 1.55E-04 | 1 | 0 | 0 | 0.103952161 | 0 | 0 | 0.05086677922 |
| MARD_SAMN04427046_REFG_MMP04427046 | 5 | -0.2035547752 | Sphingomonadal | Croceicoccus | Croceicoccus bis | 0 | 0 | 0.03946696933 | 1.97E-06 | 1.23E-05 | 0.9833331186 | 0 | 0.0793729599 | 0 | 1.31E-05 | 0 |
| MARD_SAMN04449598_REFG_MMP04449598 | 6 | -0.3316631684 | Flavobacteriales | Epilithonimonas | Epilithonimonas ( | 0 | 0 | 0.05065188807 | 1.05E-04 | 0.9999997037 | 2.37E-06 | 2.96E-06 | 0.05903280229 | 0 | 0 | 0 |
| MARD_SAMN04450243_REFG_MMP04450243 | 3 | -0.1434506552 | Rickettsiales | Wolbachia | Wolbachia sp013 | 0 | 0 | 0.999999719 | 0.01678919226 | 0.01666666667 | 0.999999719 | 0 | 0.9861882485 | 0 | 0.999999719 | 0.05367407679 |
| MARD_SAMN04487765_REFG_MMP04487765 | 2 | -0.1037447381 | Flavobacteriales | Tenacibaculum | Tenacibaculum s | 0 | 0 | 0.03773028314 | 0.01666867787 | 0.05000000003 | 2.60E-05 | 0 | 0.9529417215 | 0 | 0 | 3.41E-08 |
| MARD_SAMN04487806_REFG_MMP04487806 | 2 | -0.2716576959 | Mycobacteriales | Rhodococcus | Rhodococcus qir | 0 | 0 | 0.05135779058 | 4.09E-05 | 3.78E-04 | 0.007636798263 | 0 | 0.04970956148 | 0 | 8.38E-04 | 0.03707204978 |
| MARD_SAMN04487808_REFG_MMP04487808 | 5 | -0.6374801066 | Bacillales_G | Fictibacillus_C | Fictibacillus_C er | 0 | 0 | 0.1506197476 | 1.73E-04 | 0.05730704957 | 0.9999998404 | 5.83E-04 | 0.1189731713 | 0 | 0 | 0 |
| MARD_SAMN04487859_REFG_MMP04487859 | 5 | -0.2072214997 | Rhodobacterales | Roseovarius | Roseovarius lutir | 1.28E-06 | 0 | 0.1167507144 | 1.72E-04 | 0.02721598342 | 1 | 0 | 0.04293980069 | 0 | 7.82E-04 | 0 |
| MARD_SAMN04487867_REFG_MMP04487867 | 2 | -0.1938920008 | Pseudomonadale | Halomonas | Halomonas titani | 0 | 0 | 0.04057091516 | 6.73E-05 | 0.003419020158 | 1.75E-06 | 0 | 0.04967839204 | 0 | 0 | 0 |
| MARD_SAMN04487868_REFG_MMP04487868 | 2 | -0.2710370311 | Pseudomonadale | Marinobacter | Marinobacter sal | 0 | 0 | 0.09468070138 | 1.06E-04 | 0 | 0.001745540891 | 0 | 0.06534255059 | 0 | 0.001983401134 | 0 |
| MARD_SAMN04487906_REFG_MMP04487906 | 2 | -0.2263219166 | Flavobacteriales | Zhouia | Zhouia amylolytic | 0 | 0 | 0.04081534288 | 1.14E-04 | 0.0410756166 | 0 | 0 | 0.06935585939 | 0 | 0 | 0 |
| MARD_SAMN04487910_REFG_MMP04487910 | 6 | -0.09824265154 | Flavobacteriales | Aquimarina | Aquimarina ampl | 0 | 0 | 0.04119193362 | 1.03E-04 | 1 | 0 | 0 | 0.06894933583 | 0 | 0 | 0 |
| MARD_SAMN04487911_REFG_MMP04487911 | 6 | -0.3068233569 | Flavobacteriales | Arenibacter | Arenibacter nanh | 0 | 0 | 0.02812446957 | 1.67E-06 | 0.9841662273 | 2.51E-04 | 0 | 0.07409477364 | 0 | 0 | 0.03291506942 |
| MARD_SAMN04487931_REFG_MMP04487931 | 2 | -0.09973423302 | Desulfobacterale | Desulfobacula | Desulfobacula to | 8.81E-09 | 0 | 0.01260229749 | 0 | 0.3453324742 | 0.01666667107 | 0 | 0.01671099971 | 0 | 3.23E-06 | 0 |
| MARD_SAMN04487960_REFG_MMP04487960 | 2 | -0.3365124661 | Pseudomonadale | Marinobacter | Marinobacter mo | 0 | 0 | 0.1147472303 | 8.57E-05 | 0.01567341924 | 0.00314486411 | 0 | 0.06315294114 | 0 | 0.004193076366 | 0.04195596221 |
| MARD_SAMN04487961_REFG_MMP04487961 | 5 | -0.2530937354 | Pseudomonadale | Marinobacter | Marinobacter pel | 0 | 0 | 0.2762164758 | 1.63E-04 | 0.00341511619 | 0.9999998204 | 0 | 0.05683480251 | 0 | 8.11E-04 | 0.02886870371 |
| MARD_SAMN04487965_REFG_MMP04487965 | 5 | -0.3212082404 | Pseudomonadale | Microbulbifer | Microbulbifer dor | 0 | 0 | 0.0369399201 | 3.03E-05 | 0.2351877494 | 0.8172053044 | 0 | 0.04015227186 | 0 | 7.94E-04 | 0.006760136974 |
| MARD_SAMN04487972_REFG_MMP04487972 | 1 | -0.1044585451 | Rhodobacterales | Paracoccus | Paracoccus halo | 0 | 0 | 0.9999997798 | 3.48E-05 | 0.1145519799 | 0.9999997798 | 0.04970527126 | 0.05586867205 | 0 | 7.71E-04 | 0 |
| MARD_SAMN04487987_REFG_MMP04487987 | 2 | -0.1479621217 | Flavobacteriales | Algibacter_B | Algibacter_B pec | 0 | 0 | 0.01997225631 | 1.16E-04 | 0 | 4.75E-06 | 0 | 0.9999997034 | 0 | 0 | 0 |
| MARD_SAMN04487991_REFG_MMP04487991 | 4 | -0.5059183547 | Rhodobacterales | Celeribacter | Celeribacter nept | 0.01585034632 | 0 | 0.9847377116 | 9.24E-05 | 0.01559498797 | 0.01666666453 | 0.01584729353 | 0.07200663539 | 0 | 0.002747930999 | 0.02144986079 |
| MARD_SAMN04487992_REFG_MMP04487992 | 6 | -0.181537342 | Flavobacteriales | Cellulophaga | Cellulophaga bal | 0 | 0 | 0.02633131984 | 0 | 0.9999997292 | 0 | 0 | 0.0579959629 | 0 | 5.15E-05 | 0.03569794844 |
| MARD_SAMN04487993_REFG_MMP04487993 | 4 | -0.2228446084 | Rhodobacterales | Salipiger | Salipiger marinus | 0.01839831585 | 0 | 0.9999998102 | 0.01701004305 | 2.19E-04 | 0 | 0 | 0.9999998102 | 0 | 8.10E-04 | 0.03773809097 |
| MARD_SAMN04487999_REFG_MMP04487999 | 6 | -0.2693451039 | Flavobacteriales | Leeuwenhoekiell | Leeuwenhoekiell | 0 | 0 | 0.2149230413 | 0.01666666187 | 0.966798566 | 0 | 0 | 0.06422500156 | 0 | 0 | 0.001378311742 |
| MARD_SAMN04488001_REFG_MMP04488001 | 2 | -0.3073982113 | Rhodobacterales | Litoreibacter | Litoreibacter albi | 0 | 0 | 0.02257418357 | 0.05002923905 | 0.008579004677 | 0.03335351958 | 0.01666666357 | 0.0509423962 | 0 | 0.00270383372 | 0 |
| MARD_SAMN04488002_REFG_MMP04488002 | 2 | -0.2714507191 | Rhodobacterales | Litoreibacter | Litoreibacter jant | 0 | 0 | 0.1533645238 | 3.42E-05 | 0.01358777634 | 0.003243042177 | 1.06E-04 | 0.07131474193 | 0 | 0.001238548771 | 0 |
| MARD_SAMN04488005_REFG_MMP04488005 | 2 | -0.3775169535 | Rhodobacterales | Yoonia | Yoonia tamlensis | 0 | 0 | 0.1094477336 | 0.01683404529 | 0.01562574893 | 0.06099372482 | 0 | 1 | 0 | 0.003079433506 | 0 |
| MARD_SAMN04488006_REFG_MMP04488006 | 2 | -0.03244871102 | Flavobacteriales | Lutibacter | Lutibacter maritir | 2.08E-04 | 0 | 0.02949310835 | 2.19E-06 | 0.08450278993 | 6.16E-04 | 0 | 0.06585241592 | 0 | 2.58E-05 | 0.002210019258 |
| MARD_SAMN04488007_REFG_MMP04488007 | 6 | -0.09053212219 | Flavobacteriales | Maribacter | Maribacter aquivi | 0 | 0 | 0.03253015037 | 1.06E-04 | 1 | 0 | 0 | 0.06399082582 | 0 | 0 | 0 |
| MARD_SAMN04488008_REFG_MMP04488008 | 2 | -0.1716256841 | Flavobacteriales | Maribacter | Maribacter orient | 0 | 0 | 0.0351395963 | 1.36E-04 | 0.2510977928 | 0 | 0 | 0.05816178678 | 0 | 0 | 0 |
| MARD_SAMN04488012_REFG_MMP04488012 | 8 | -0.07711729202 | Rhodobacterales | Palleronia | Palleronia salina | 5.52E-07 | 0 | 0.9999997238 | 0.9999997238 | 0.003159699058 | 0.9999997238 | 0 | 0.08991302402 | 0 | 0.001711618759 | 0.03756168913 |
| MARD_SAMN04488013_REFG_MMP04488013 | 4 | -0.3756228734 | Lactobacillales | Marinilactibacillus | Marinilactibacillus | 0 | 0 | 1.00000024 | 1.00000024 | 0.0101049922 | 0.0337330331 | 0 | 0.8983505753 | 0 | 0.007369889405 | 0 |
| MARD_SAMN04488018_REFG_MMP04488018 | 4 | -0.1579480541 | Flavobacteriales | Flavobacterium | Flavobacterium r | 0 | 0 | 0.999999756 | 0.1271309801 | 0 | 5.37E-06 | 0 | 0.1554250417 | 0.999999756 | 8.67E-04 | 0.0314074236 |
| MARD_SAMN04488020_REFG_MMP04488020 | 2 | -0.09488761848 | Rhodobacterales | Palleronia | Palleronia marisr | 6.72E-06 | 0 | 0.02210856682 | 5.86E-05 | 0.09732986521 | 5.81E-04 | 0 | 0.06557895837 | 0 | 7.80E-04 | 0 |
| MARD_SAMN04488026_REFG_MMP04488026 | 5 | -0.1615471273 | Rhodobacterales | Tropicimonas | Tropicimonas luti | 0 | 0 | 0.104462653 | 1.57E-06 | 0.01311934195 | 0.9666666667 | 0 | 0.0718499334 | 0 | 1.32E-05 | 0 |
| MARD_SAMN04488031_REFG_MMP04488031 | 4 | 0.002197271014 | Rhodobacterales | Roseovarius | Roseovarius indi | 0 | 0 | 1 | 5.55E-05 | 0.003137996092 | 0.002409093387 | 0 | 0.05359870988 | 0 | 7.86E-04 | 0 |
| MARD_SAMN04488032_REFG_MMP04488032 | 2 | -0.4492821789 | Rhodobacterales | Pacificibacter | Pacificibacter ma | 0.007335422682 | 0 | 0.01607757441 | 9.98E-05 | 0.02199022022 | 0.02466003069 | 0 | 0.04464662097 | 0.03236142212 | 0.001422385466 | 0 |
| MARD_SAMN04488033_REFG_MMP04488033 | 6 | -0.1510147553 | Flavobacteriales | Salegentibacter | Salegentibacter a | 1.66E-06 | 0 | 0.03369442486 | 1.35E-04 | 0.9999997229 | 0 | 0 | 0.05994017675 | 0 | 0 | 0.03818269227 |
| MARD_SAMN04488035_REFG_MMP04488035 | 4 | -0.1329492819 | Actinomycetales | Flavimobilis | Flavimobilis mari | 1.16E-05 | 0 | 0.9864016277 | 0.06069498236 | 0.09037480118 | 0.04081785185 | 0 | 0.05139619556 | 0 | 2.70E-05 | 0 |
| MARD_SAMN04488036_REFG_MMP04488036 | 2 | -0.3181504051 | Rhodobacterales | Shimia | Shimia haliotis | 0 | 0 | 0.02034757877 | 1.43E-05 | 0.05340281115 | 0.00361450032 | 0 | 0.05470373142 | 0 | 7.87E-04 | 0.06357438577 |
| MARD_SAMN04488039_REFG_MMP04488039 | 2 | -0.2499466966 | Rhodobacterales | Sulfitobacter | Sulfitobacter dub | 0 | 0 | 0.02592578862 | 2.46E-06 | 0.09587000944 | 0.003253025736 | 0 | 0.06132496523 | 0 | 1.51E-05 | 9.13E-05 |

| | | | | | | | | | | | | | | | | |
|---|---|---|---|---|---|---|---|---|---|---|---|---|---|---|---|---|
| MARD_SAMN04488040_REFG_MMP04488040 | 4 | -0.2604181975 | Rhodobacterales | Sulfitobacter | Sulfitobacter mar | 0 | 0 | 0.9844442643 | 0.9833361373 | 0.06546674358 | 0.001227642246 | 6.17E-04 | 0.0523197311 | 0 | 7.72E-04 | 1.12E-05 |
| MARD_SAMN04488042_REFG_MMP04488042 | 2 | -0.2723497481 | Rhodobacterales | Shimia | Shimia aestuarii | 0.02435041629 | 0 | 0.02370364764 | 4.50E-05 | 0.01685806194 | 4.20E-04 | 0.02435041629 | 0.07665187664 | 0 | 0.001308621513 | 0.01295204504 |
| MARD_SAMN04488043_REFG_MMP04488043 | 1 | -0.2388958211 | Rhodobacterales | Thalassobius | Thalassobius gel | 6.42E-06 | 0 | 0.9840621304 | 0.01666666667 | 0.07044910365 | 1.000000172 | 0 | 0.07798137402 | 0 | 1.31E-05 | 0.0356866026 |
| MARD_SAMN04488044_REFG_MMP04488044 | 1 | -0.4732300518 | Rhodobacterales | Cognatishimia | Cognatishimia m | 0 | 0 | 0.9999997954 | 9.86E-05 | 0.01627613106 | 0.9999997954 | 0 | 0.05021215901 | 7.50E-04 | 0.001487312922 | 0 |
| MARD_SAMN04488047_REFG_MMP04488047 | 2 | -0.05189613315 | Rhodobacterales | Tranquillimonas | Tranquillimonas | 0 | 0 | 0.07590229552 | 3.29E-05 | 0.03879208674 | 6.15E-06 | 0 | 1 | 0 | 7.82E-04 | 0 |
| MARD_SAMN04488049_REFG_MMP04488049 | 2 | -0.4208824031 | Rhodobacterales | Epibacterium | Epibacterium mu | 3.24E-06 | 0 | 0.1326928827 | 1.73E-04 | 0.02069602576 | 6.07E-06 | 0 | 0.0699396194 | 1.95E-04 | 7.81E-04 | 0.0156986335 |
| MARD_SAMN04488050_REFG_MMP04488050 | 5 | -0.2567362227 | Rhodobacterales | Salipiger | Salipiger pacificu | 0 | 0 | 0.122461675 | 1.41E-04 | 0.03219131263 | 0.9999998447 | 0 | 0.0779826239 | 0 | 8.13E-04 | 0.03798842038 |
| MARD_SAMN04488054_REFG_MMP04488054 | 5 | -0.1066170839 | Bacillales_H | Salibacterium | Salibacterium qir | 0 | 0 | 0.0578499164 | 1.51E-04 | 0.005775568666 | 1 | 0 | 0.3708483048 | 0 | 0 | 0 |
| MARD_SAMN04488056_REFG_MMP04488056 | 5 | -0.2969553106 | Rhizobiales | Cohaesibacter | Cohaesibacter m | 0.195049395 | 0 | 0.07073557367 | 1.28E-06 | 0.001310424912 | 0.9999998801 | 0 | 0.03444900885 | 0 | 2.59E-05 | 0 |
| MARD_SAMN04488057_REFG_MMP04488057 | 2 | -0.1831195183 | Cytophagales | Cyclobacterium | Cyclobacterium li | 0.1796764313 | 0 | 0.273569043 | 1 | 0.007686999285 | 0.00291092741 | 0 | 0.05621546387 | 0 | 0 | 0 |
| MARD_SAMN04488070_REFG_MMP04488070 | 1 | -0.3450810843 | Enterobacterales | Pseudidiomarina | Pseudidiomarina | 0 | 0 | 1 | 3.29E-05 | 0.1179134383 | 1 | 0.09833319153 | 0.07300824323 | 0 | 0.01424585788 | 2.11E-04 |
| MARD_SAMN04488081_REFG_MMP04488081 | 2 | -0.4243512253 | Bacillales_D | Salimicrobium | Salimicrobium all | 0 | 0 | 0.1604564637 | 0.999999995 | 0.1140835294 | 0 | 0 | 0.05531323707 | 0 | 0 | 0 |
| MARD_SAMN04488087_REFG_MMP04488087 | 4 | -0.1093603663 | Rhodothermales | Rhodothermus | Rhodothermus pr | 0 | 0 | 1 | 9.73E-05 | 1 | 0.01696120437 | 0 | 0.1393712477 | 0 | 0.00277000013 | 0 |
| MARD_SAMN04488089_REFG_MMP04488089 | 6 | -0.1935637323 | Flavobacteriales | Flavobacterium | Flavobacterium p | 0 | 0 | 0.02676254464 | 0.03333470664 | 0.9666914903 | 0 | 8.75E-05 | 0.1759575997 | 0 | 4.04E-05 | 3.94E-05 |
| MARD_SAMN04488092_REFG_MMP04488092 | 8 | -0.1972890037 | Rhodobacterales | Litorimicrobium | Litorimicrobium ta | 0 | 0 | 0.1470774635 | 1 | 0.01278109726 | 1 | 0 | 0.06202202528 | 0 | 8.70E-04 | 0.04598463731 |
| MARD_SAMN04488094_REFG_MMP04488094 | 2 | -0.07682642638 | Rhodobacterales | Tropicimonas | Tropicimonas iso | 0 | 0 | 0.06605526497 | 0.9666670834 | 0.06496160326 | 1.27E-04 | 0 | 0.06801320876 | 0 | 6.44E-05 | 0 |
| MARD_SAMN04488095_REFG_MMP04488095 | 5 | -0.08630522084 | Rhodobacterales | Jannaschia | Jannaschia poha | 0 | 0 | 0.02803518432 | 0.01666770666 | 0.00150378051 | 0.9834716444 | 0 | 0.07502882584 | 0 | 2.03E-05 | 0.03379271804 |
| MARD_SAMN04488096_REFG_MMP04488096 | 8 | -0.07652706281 | Flavobacteriales | Mesonia | Mesonia phycico | 0 | 0 | 0.03434953197 | 1 | 0.1090936544 | 1 | 0 | 0.06573192186 | 0 | 0 | 0 |
| MARD_SAMN04488098_REFG_MMP04488098 | 4 | -0.4516699867 | Lactobacillales | Alkalibacterium | Alkalibacterium tl | 0 | 0 | 1 | 1.71E-04 | 0.001590695145 | 0.001604920193 | 0 | 1 | 0 | 0.002264293036 | 0 |
| MARD_SAMN04488100_REFG_MMP04488100 | 8 | -0.5539331836 | Lactobacillales | Alkalibacterium | Alkalibacterium p | 0.01598785002 | 0 | 1.000000137 | 1.000000137 | 0.1590040618 | 1.000000137 | 0 | 1.000000137 | 0 | 0.005984746426 | 0 |
| MARD_SAMN04488102_REFG_MMP04488102 | 1 | -0.5569591581 | Lactobacillales | Alkalibacterium | Alkalibacterium s | 0.0340628418 | 0 | 1 | 1.61E-04 | 0.1621800256 | 1 | 0 | 1 | 9.20E-07 | 0.005535936211 | 0 |
| MARD_SAMN04488104_REFG_MMP04488104 | 2 | -0.2907715973 | Cytophagales | Algoriphagus | Algoriphagus fae | 0 | 0 | 0.0371771965 | 5.29E-05 | 0.07920225728 | 5.46E-06 | 0 | 0.06023187898 | 0 | 1.58E-05 | 0 |
| MARD_SAMN04488108_REFG_MMP04488108 | 2 | -0.2672916515 | Cytophagales | Algoriphagus | Algoriphagus zha | 0 | 0 | 0.02771209071 | 9.01E-05 | 0.1136757094 | 5.21E-06 | 0 | 0.05948168785 | 0 | 0 | 0 |
| MARD_SAMN04488110_REFG_MMP04488110 | 4 | -0.04500731496 | Mycobacteriales | Micromonospora | Micromonospora | 0 | 0 | 1.000000407 | 0.1143333678 | 0.009911902647 | 0.03333341997 | 0 | 0.04937506245 | 0 | 7.27E-07 | 0 |
| MARD_SAMN04488117_REFG_MMP04488117 | 5 | -0.3362013711 | Rhodobacterales | Celeribacter | Celeribacter bael | 5.86E-08 | 0 | 0.06745819524 | 2.24E-06 | 1.66E-05 | 0.9999998315 | 0 | 0.2059679503 | 0 | 7.63E-04 | 0 |
| MARD_SAMN04488125_REFG_MMP04488125 | 5 | -0.07180995688 | Rhizobiales | Methylobacteriun | Methylobacteriun | 0 | 9.11E-05 | 0.03520337409 | 4.50E-06 | 0.007226294105 | 0.9999997133 | 0 | 0.06668321091 | 0 | 2.03E-04 | 0.02967120689 |
| MARD_SAMN04488129_REFG_MMP04488129 | 5 | -0.110686375 | Pseudomonadale | Halomonas | Halomonas daqia | 0 | 0 | 0.2326547847 | 5.36E-07 | 0.007070018374 | 0.9999999975 | 0 | 0.2053826974 | 0 | 0 | 0.09722035997 |
| MARD_SAMN04488134_REFG_MMP04488134 | 8 | -0.346512731 | Bacillales_D | Amphibacillus_D | Amphibacillus_D | 0 | 0 | 0.03743470979 | 0.9833364516 | 0.417543839 | 0.9999999994 | 0.06147529536 | 0.06256352656 | 5.92E-04 | 4.43E-05 | 0 |
| MARD_SAMN04488139_REFG_MMP04488139 | 1 | -0.3549120883 | Enterobacterales | Pseudidiomarina | Pseudidiomarina | 0 | 0 | 0.9999999508 | 1.12E-04 | 5.84E-05 | 0.9999999508 | 0 | 0.2728013494 | 0 | 0.008515324012 | 0 |
| MARD_SAMN04488238_REFG_MMP04488238 | 7 | -0.122623425 | Rhodobacterales | Roseicitreum | Roseicitreum ant | 9.53E-04 | 0 | 0.02019973665 | 1 | 0.01096980126 | 0.01007255755 | 0 | 0.04315286416 | 0 | 0.9833461566 | 2.31E-04 |
| MARD_SAMN04488239_REFG_MMP04488239 | 7 | -0.2503510552 | Rhodobacterales | Ruegeria_B | Ruegeria_B mari | 0 | 0 | 0.05574438994 | 1.04E-06 | 0.08734783757 | 0.01666666667 | 0 | 0.07461382141 | 0 | 1.0000002 | 1.04E-04 |
| MARD_SAMN04488512_REFG_MMP04488512 | 2 | -0.2491844443 | Rhodobacterales | Sulfitobacter | Sulfitobacter litor | 0 | 0 | 0.06016906749 | 1 | 0.1477041513 | 0 | 0 | 0.03393554707 | 0 | 0.001428088332 | 0.006185544437 |
| MARD_SAMN04488513_REFG_MMP04488513 | 6 | -0.2634696312 | Flavobacteriales | Pseudozobellia | Pseudozobellia tl | 1.70E-06 | 0 | 0.03783073438 | 9.45E-05 | 1.000000213 | 0 | 0 | 0.1526964344 | 2.60E-05 | 0 | 0 |
| MARD_SAMN04488514_REFG_MMP04488514 | 2 | -0.1270032012 | Flavobacteriales | Kriegella | Kriegella aquima | 0 | 0.00669974839 | 0.03188715708 | 0.004747620294 | 0 | 0 | 0 | 1 | 0 | 0 | 0 |
| MARD_SAMN04488515_REFG_MMP04488515 | 2 | -0.186541371 | Rhodobacterales | Cognatiyoonia | Cognatiyoonia kc | 0 | 0 | 0.03029241723 | 3.66E-05 | 0.1141300143 | 1.31E-05 | 0 | 0.06162716542 | 0 | 8.05E-04 | 0.03328820399 |
| MARD_SAMN04488518_REFG_MMP04488518 | 5 | -0.2524937186 | Rhizobiales | Pseudovibrio | Pseudovibrio asc | 0 | 0 | 0.02746543827 | 1.17E-04 | 0.007222184203 | 1.000000128 | 0 | 0.06095457755 | 0 | 8.16E-04 | 0.02120748767 |
| MARD_SAMN04488519_REFG_MMP04488519 | 5 | -0.3328213853 | Cytophagales | Algoriphagus | Algoriphagus orn | 0 | 0 | 0.006083069786 | 0.01666758594 | 0.01840219304 | 0.001348462857 | 0 | 0.3675529967 | 0.9333335178 | 0 | 5.79E-04 |
| MARD_SAMN04488526_REFG_MMP04488526 | 2 | -0.1241173589 | Rhodobacterales | Jannaschia | Jannaschia helgc | 0 | 0 | 0.06399828245 | 0.9833340583 | 0.001894827482 | 0.01666666667 | 0 | 0.06855547233 | 0 | 2.87E-04 | 0.03548363519 |
| MARD_SAMN04488527_REFG_MMP04488527 | 5 | -0.4934843716 | Rhodobacterales | Aliiroseovarius | Aliiroseovarius cr | 0 | 0 | 0.01916299891 | 0 | 0.007756603915 | 0.9999999034 | 0.05947445095 | 0.02636295558 | 0 | 1.39E-05 | 0.004107300339 |
| MARD_SAMN04488540_REFG_MMP04488540 | 4 | -0.8940833047 | Enterobacterales | Ferrimonas | Ferrimonas sedir | 0 | 0 | 1.000000023 | 0.01669465105 | 5.32E-04 | 0.1166671536 | 0 | 0.2113196437 | 0 | 2.31E-04 | 0 |
| MARD_SAMN04488567_REFG_MMP04488567 | 5 | -0.1022954683 | Rhodobacterales | Limimaricola | Limimaricola pyo | 0 | 0 | 0.02856298173 | 2.96E-07 | 0.1183166155 | 1.00000026 | 0 | 0.05370737403 | 0 | 1.30E-05 | 0 |
| MARD_SAMN04488569_REFG_MMP04488569 | 8 | -0.3428054514 | Lactobacillales | Marinilactibacillu | Marinilactibacillu | 0 | 0 | 1 | 1 | 0.2077525205 | 1 | 0 | 0.2478127854 | 0 | 0.003592300063 | 0 |
| MARD_SAMN04489724_REFG_MMP04489724 | 2 | -0.3327668963 | Cytophagales | Algoriphagus | Algoriphagus loci | 0 | 0 | 0.04237690828 | 1.75E-05 | 0.001768790221 | 0 | 0 | 0.03342810339 | 0 | 0 | 0.03751765474 |
| MARD_SAMN04489759_REFG_MMP04489759 | 2 | -0.1926866318 | Rhodobacterales | Sulfitobacter | Sulfitobacter deli | 0 | 0 | 0.03035398129 | 0.01666666286 | 0.008119640763 | 0.09723843918 | 0 | 0.06791429864 | 0 | 2.04E-05 | 2.55E-04 |
| MARD_SAMN04489796_REFG_MMP04489796 | 2 | -0.1543400685 | Flavobacteriales | Winogradskyella | Winogradskyella | 0 | 0 | 0.01942259301 | 0.01666666667 | 5.60E-05 | 0 | 0 | 0.9845144581 | 0 | 2.59E-05 | 0 |
| MARD_SAMN04489800_REFG_MMP04489800 | 2 | -0.4376660378 | Pseudomonadale | Pseudomonas_E | Pseudomonas_E | 0 | 0 | 0.0474051142 | 6.44E-06 | 0.01157020595 | 0.1666666698 | 0 | 0.06296600633 | 0 | 2.07E-06 | 0.01613374804 |
| MARD_SAMN04489858_REFG_MMP04489858 | 2 | -0.2984651432 | Rhodobacterales | Paracoccus | Paracoccus hom | 0.01918614757 | 0 | 0.02625949747 | 3.23E-05 | 0.001618957367 | 5.83E-06 | 0.00447115293 | 0.05439185363 | 0 | 8.00E-04 | 0 |
| MARD_SAMN04490244_REFG_MMP04490244 | 5 | -0.03720246522 | Rhodobacterales | Tranquillimonas | Tranquillimonas r | 0 | 0 | 0.03403490169 | 3.61E-05 | 0.08792979882 | 1.000000003 | 0.03746015114 | 0.06143502436 | 0 | 0.001266848017 | 0.03641951322 |
| MARD_SAMN04490248_REFG_MMP04490248 | 5 | -0.1035895784 | Rhodobacterales | Salinihabitans | Salinihabitans fla | 0.04510516991 | 0 | 0.1015907114 | 6.44E-07 | 0.1132306032 | 0.9836620483 | 0 | 0.968436687 | 0 | 1.31E-05 | 0.03492925211 |
| MARD_SAMN04490369_REFG_MMP04490369 | 4 | -0.2284071294 | Pseudomonadale | Halomonas | Halomonas aqua | 0 | 0 | 1.000000028 | 4.41E-06 | 0.002950618984 | 0.250645099 | 0 | 0.188032498 | 0 | 8.57E-05 | 0.02031065131 |
| MARD_SAMN04495211_REFG_MMP04495211 | 5 | -0.1608296805 | Mycobacteriales | Rhodococcus | Rhodococcus sp | 0 | 0 | 0.1261398846 | 1.47E-04 | 0.06104133506 | 1 | 0.06996502589 | 0.06767582815 | 0 | 7.73E-04 | 0 |
| MARD_SAMN04515623_REFG_MMP04515623 | 1 | -0.2767981525 | Rhodospirillales_ | Thalassospira | Thalassospira xia | 0 | 0.002520077947 | 0.9999998097 | 1.03E-05 | 0.1084101608 | 0.9999998097 | 0 | 0.02804948491 | 0 | 7.65E-04 | 0 |
| MARD_SAMN04515658_REFG_MMP04515658 | 1 | -0.2190696258 | Enterobacterales | Idiomarina | Idiomarina zobell | 0 | 0 | 1 | 2.93E-05 | 0 | 1 | 0 | 0.02376119012 | 0 | 0.009634534972 | 0 |
| MARD_SAMN04515673_REFG_MMP04515673 | 5 | -0.1597375252 | Rhodobacterales | Poseidonocella | Poseidonocella s | 3.10E-04 | 0 | 0.05305628452 | 0.01666667089 | 0.06935868912 | 0.9999998367 | 0 | 0.9865557031 | 0 | 1.29E-05 | 0 |
| MARD_SAMN04570274_REFG_MMP04570274 | 4 | -0.3118848027 | Flavobacteriales | Flavobacterium | Flavobacterium c | 0 | 0 | 1 | 6.12E-05 | 0.01575181335 | 0.002907672111 | 0 | 0.05703369415 | 0 | 0.001604752478 | 0.01410727168 |
| MARD_SAMN04571751_REFG_MMP04571751 | 5 | -0.1443836028 | Streptomycetales | Streptomyces | Streptomyces sp | 0 | 0 | 0.02068573316 | 0.01860132705 | 0.02355588784 | 0.9999998159 | 0 | 0.0370424851 | 0 | 7.35E-04 | 0 |
| MARD_SAMN04867941_REFG_MMP04867941 | 4 | -0.6361539392 | Enterobacterales | Vibrio | Vibrio sp000287 | 0 | 0 | 1.000000137 | 3.64E-07 | 5.35E-05 | 1.66E-07 | 0 | 0.07260945293 | 0 | 1.82E-07 | 0 |
| MARD_SAMN04903162_REFG_MMP04903162 | 2 | -0.4411604792 | Enterobacterales | Pseudoalteromor | Pseudoalteromor | 0 | 0 | 0.01535035534 | 1.15E-06 | 0.01966549495 | 0 | 0 | 0.05515480164 | 0 | 1.65E-04 | 5.19E-05 |
| MARD_SAMN04915612_REFG_MMP04915612 | 2 | -0.1619824352 | Rhodobacterales | Paracoccus | Paracoccus mari | 0 | 0 | 0.1241911662 | 4.02E-05 | 0.09353122418 | 0.003547001877 | 0 | 0.06411990187 | 0 | 0.003743514923 | 0.04477489196 |
| MARD_SAMN04957305_REFG_MMP04957305 | 2 | -0.2357317922 | Rhodobacterales | Sulfitobacter | Sulfitobacter geo | 0 | 0 | 0.03540470893 | 1.06E-04 | 0.01029619985 | 0 | 0 | 0.0610709054 | 0.003448394555 | 7.80E-04 | 0.04558308284 |
| MARD_SAMN04957306_REFG_MMP04957306 | 5 | -0.1548832191 | Sphingomonadal | Qipengyuania | Qipengyuania fla | 7.75E-04 | 0 | 0.02810133547 | 4.31E-06 | 1.02E-04 | 0.983333345 | 2.14E-05 | 0.06455856305 | 0 | 0.007341120449 | 0.03355600548 |
| MARD_SAMN04957307_REFG_MMP04957307 | 5 | -0.1746134655 | Rhodobacterales | Gymnodinialimor | Gymnodinialimor | 5.96E-05 | 7.30E-05 | 0.04811098248 | 0.06666815659 | 0.002636190417 | 0.9667880179 | 3.67E-04 | 0.904959956 | 7.03E-04 | 0.001541400121 | 5.59E-06 |
| MARD_SAMN04957308_REFG_MMP04957308 | 2 | -0.1273679677 | Balneolales | Balneola | Balneola sp0016 | 9.42E-08 | 0 | 0.03632034297 | 0.05000000277 | 0.02358155667 | 0.002660697708 | 0 | 0.08821492566 | 0 | 2.22E-08 | 2.87E-05 |
| MARD_SAMN04965930_REFG_MMP04965930 | 2 | -0.2875196358 | Pseudomonadale | Marinobacter | Marinobacter sp0 | 0 | 0 | 0.03528214863 | 8.26E-05 | 0 | 0.002779780345 | 0 | 0.06698983005 | 0 | 8.16E-04 | 0.04684477302 |
| MARD_SAMN04965932_REFG_MMP04965932 | 2 | -0.1950003193 | Rhodobacterales | Sulfitobacter | Sulfitobacter sp0 | 0 | 0 | 0.02823407056 | 5.36E-07 | 0.2425687347 | 0.02716832397 | 0 | 0.03950496397 | 0 | 1.26E-05 | 0 |
| MARD_SAMN04965933_REFG_MMP04965933 | 5 | -0.1899060276 | Rhodobacterales | EhC02 | EhC02 sp001650 | 0.008438031587 | 0 | 0.03875401475 | 3.32E-05 | 0.01992056776 | 1.000000202 | 0 | 0.3056120233 | 0 | 8.08E-04 | 0.04005354282 |
| MARD_SAMN04999933_REFG_MMP04999933 | 5 | -0.7272445764 | Enterobacterales | Pseudoalteromor | Pseudoalteromor | 0 | 0 | 0.03793013212 | 3.21E-05 | 0 | 1.000000173 | 0 | 0.09170907777 | 0 | 8.41E-04 | 0.00276969557 |
| MARD_SAMN05001254_REFG_MMP05001254 | 6 | -0.04560962612 | Flavobacteriales | Algibacter_C | Algibacter_C aqu | 3.52E-06 | 0 | 0.02838727717 | 0.05224836127 | 1 | 0 | 0 | 0.04071210093 | 0 | 0 | 0 |
| MARD_SAMN05172613_REFG_MMP05172613 | 2 | -0.3770340005 | Enterobacterales | Vibrio | Vibrio cyclitrophic | 0 | 0 | 0.02807337051 | 7.86E-06 | 0.002724319783 | 0.0355747686 | 0 | 0.08528923077 | 0 | 2.21E-05 | 0 |
| MARD_SAMN05216175_REFG_MMP05216175 | 5 | -0.2959433452 | Pseudomonadale | Neptunomonas | Neptunomonas q | 0 | 0 | 0.05182088103 | 1.15E-04 | 0 | 1.000000205 | 0 | 0.1969136384 | 0 | 0.01389045489 | 0 |
| MARD_SAMN05216203_REFG_MMP05216203 | 5 | -0.287516606 | Pseudomonadale | Marinobacter | Marinobacter dac | 0 | 0 | 0.0502534959 | 1.92E-06 | 0.09945557741 | 0.9833333265 | 0 | 0.06156906885 | 0 | 1.30E-05 | 0 |
| MARD_SAMN05216207_REFG_MMP05216207 | 2 | 0.04895346248 | Mycobacteriales | Pseudonocardia | Pseudonocardia | 0 | 0 | 0.03349855419 | 8.73E-05 | 0.08315224753 | 8.18E-06 | 0 | 0.05119654129 | 0 | 0 | 0 |
| MARD_SAMN05216208_REFG_MMP05216208 | 2 | -0.1214670512 | Rhodobacterales | Roseovarius | Roseovarius nan | 0 | 0 | 0.02744333104 | 2.13E-04 | 7.81E-04 | 8.70E-05 | 0 | 0.04364399382 | 0 | 7.40E-04 | 0 |

| | | | | | | | | | | | | | | | | |
|---|---|---|---|---|---|---|---|---|---|---|---|---|---|---|---|---|
| MARD_SAMN05216212_REFG_MMP05216212 | 2 | -0.3427789013 | Pseudomonadale | Microbulbifer | Microbulbifer yue | 0 | 0 | 0.03373325403 | 1.21E-05 | 0.002310020805 | 0.05644539384 | 0 | 0.005677295267 | 0 | 0.003491348337 | 0.002279291348 |
| MARD_SAMN05216216_REFG_MMP05216216 | 2 | -0.3731898172 | Staphylococcales | Salinicoccus | Salinicoccus qing | 0 | 0 | 0.03401907873 | 4.13E-06 | 0.001441676102 | 7.53E-05 | 0 | 0.04739769357 | 0 | 1.30E-05 | 0 |
| MARD_SAMN05216224_REFG_MMP05216224 | 2 | -0.2404458842 | Rhodobacterales | Thioclava | Thioclava daliane | 0 | 0 | 0.1425942811 | 0 | 0.08334321236 | 4.14E-07 | 0 | 0.04915448347 | 0 | 1.30E-05 | 5.05E-08 |
| MARD_SAMN05216234_REFG_MMP05216234 | 4 | -0.2622050194 | Campylobacteral | Hydrogenimonas | Hydrogenimonas | 0 | 0 | 1 | 2.06E-04 | 0.1904259783 | 0.07211300279 | 0 | 0.05922250017 | 0 | 0.03271731917 | 0.06749169834 |
| MARD_SAMN05216236_REFG_MMP05216236 | 2 | -0.1044937049 | Rhodobacterales | Sedimentitalea | Sedimentitalea n | 0 | 0 | 0.1091854137 | 1.61E-05 | 0.0530048296 | 6.18E-06 | 0 | 0.06004797475 | 0 | 7.72E-04 | 0.00598249882 |
| MARD_SAMN05216256_REFG_MMP05216256 | 1 | -0.2117890204 | Pseudomonadale | Halopseudomona | Halopseudomona | 0 | 0.01666666309 | 0.9850340208 | 3.06E-05 | 0.03839203199 | 0.9999998595 | 0 | 0.3147218399 | 0 | 0 | 0.04186059273 |
| MARD_SAMN05216258_REFG_MMP05216258 | 2 | -0.04535129042 | Rhodobacterales | Albimonas | Albimonas pacific | 0 | 0 | 0.04177813524 | 0 | 0.001532339784 | 9.70E-04 | 0 | 0.0589881756 | 0 | 0 | 0.04024926502 |
| MARD_SAMN05216261_REFG_MMP05216261 | 6 | -0.1469220268 | Flavobacteriales | Arenitalea | Arenitalea lutea | 1.09E-06 | 0 | 0.04220106353 | 1.51E-04 | 1.000000272 | 0 | 0 | 0.05005122623 | 3.43E-05 | 0 | 0 |
| MARD_SAMN05216262_REFG_MMP05216262 | 5 | -0.4507148764 | Enterobacterales | Cognaticolwellia | Cognaticolwellia | 0 | 0.001341020892 | 0.1392915931 | 9.79E-05 | 0.02358754215 | 0.9999998143 | 0 | 0.3521793874 | 0 | 0.02151599038 | 0 |
| MARD_SAMN05216264_REFG_MMP05216264 | 5 | -0.356550118 | Pseudomonadale | Pseudomonas_E | Pseudomonas_E | 0 | 0 | 0.05974711716 | 3.08E-07 | 0.001607900805 | 1.000000232 | 4.07E-04 | 0.05398480783 | 0 | 1.36E-05 | 0 |
| MARD_SAMN05216266_REFG_MMP05216266 | 5 | -0.09256952171 | Mycobacteriales | Amycolatopsis_D | Amycolatopsis_D | 0 | 0 | 0.0317968125 | 1.41E-04 | 0.3339518623 | 0.9999999964 | 0.08746730269 | 0.05549176857 | 0 | 7.30E-04 | 3.62E-04 |
| MARD_SAMN05216283_REFG_MMP05216283 | 6 | -0.1727821014 | Bacteroidales | Sunxiuqinia | Sunxiuqinia ellipt | 0 | 0 | 0.0378945968 | 9.77E-05 | 1 | 0 | 0 | 0.07619137034 | 0 | 0 | 0 |
| MARD_SAMN05216284_REFG_MMP05216284 | 3 | -0.01980504224 | Mycobacteriales | Micromonospora | Micromonospora | 0 | 0.008947968637 | 0.02818031232 | 0.09881054868 | 0.003875247785 | 1 | 0 | 0.04073641286 | 0 | 1 | 0.06187578202 |
| MARD_SAMN05216293_REFG_MMP05216293 | 2 | -0.3326308634 | Flavobacteriales | Muricauda | Muricauda taean | 2.01E-07 | 0 | 0.02588440368 | 0.03547080423 | 0.1180605278 | 4.25E-05 | 3.60E-06 | 0.05701005507 | 0 | 2.71E-05 | 0 |
| MARD_SAMN05216294_REFG_MMP05216294 | 2 | -0.2249416952 | Flavobacteriales | Muricauda | Muricauda zhang | 0.002350116297 | 0.03310752958 | 0.02561786514 | 1.21E-04 | 0.1070895988 | 0 | 0 | 0.08215422594 | 0 | 0 | 0 |
| MARD_SAMN05216342_REFG_MMP05216342 | 4 | -0.6095951016 | Exiguobacterales | Exiguobacterium | Exiguobacterium | 0 | 0 | 1 | 1.53E-04 | 0.006596340507 | 0 | 0 | 0.2191828409 | 0 | 0 | 0 |
| MARD_SAMN05216369_REFG_MMP05216369 | 5 | -0.1404925574 | Pseudomonadale | Marinobacter | Marinobacter ant | 0 | 0 | 0.1115158783 | 5.99E-05 | 0.02407506685 | 0.9833331363 | 0 | 0.05474650115 | 0 | 8.08E-04 | 7.49E-04 |
| MARD_SAMN05216376_REFG_MMP05216376 | 5 | -0.1735743828 | Rhodobacterales | Mameliella | Mameliella alba | 0 | 0 | 0.029286103 | 7.96E-07 | 3.54E-05 | 0.9999998068 | 0 | 0.05345372613 | 0 | 0.004304182996 | 0.03527370648 |
| MARD_SAMN05216480_REFG_MMP05216480 | 2 | -0.1820172296 | Flavobacteriales | Pustulibacterium | Pustulibacterium | 0 | 0 | 0.1269852573 | 1.91E-06 | 0.1048498153 | 7.03E-06 | 0 | 0.05490468031 | 0 | 2.75E-05 | 0 |
| MARD_SAMN05216562_REFG_MMP05216562 | 5 | -0.2884777494 | Pseudomonadale | Microbulbifer | Microbulbifer mai | 0 | 0 | 0.09978350864 | 0 | 0.001431729619 | 0.9999999932 | 0 | 0.04512980326 | 0 | 1.39E-05 | 0.02751994716 |
| MARD_SAMN05216571_REFG_MMP05216571 | 5 | -0.2134123805 | Pseudomonadale | Halomonas_E | Halomonas_E ta | 0 | 3.49E-04 | 0.03179808592 | 7.34E-05 | 0.1053568292 | 0.6013937831 | 0 | 0.0643310673 | 0 | 7.87E-04 | 0.01200974148 |
| MARD_SAMN05216575_REFG_MMP05216575 | 1 | -0.3811182284 | Pseudomonadale | Pseudomonas_E | Pseudomonas_E | 8.90E-05 | 0 | 0.5392252555 | 4.39E-06 | 0.5227359048 | 0.9833691261 | 0 | 0.05585797709 | 9.13E-04 | 9.31E-05 | 0.1754650168 |
| MARD_SAMN05216600_REFG_MMP05216600 | 2 | -0.3369474654 | Pseudomonadale | Pseudomonas_E | Pseudomonas_E | 0 | 0 | 0.1604541201 | 1.22E-06 | 0.00288303823 | 0.03462719804 | 7.42E-04 | 0.1846525123 | 0.001795491883 | 1.24E-05 | 0 |
| MARD_SAMN05225367_REFG_MMP05225367 | 2 | -0.5693701143 | Enterobacterales | Vibrio | Vibrio scophthalm | 0 | 0 | 0.04907145536 | 1.43E-04 | 0 | 0 | 0 | 0.01280836091 | 0 | 0 | 0 |
| MARD_SAMN05375008_REFG_MMP05375008 | 5 | -0.9199573219 | Bacillales | Bacillus | Bacillus velezens | 0 | 0 | 0.02795320943 | 1.33E-04 | 0.006670158716 | 1 | 0 | 0.3033678259 | 0 | 0 | 0.5833333333 |
| MARD_SAMN05410694_REFG_MMP05410694 | 2 | -0.2462193666 | Lactobacillales | Listeria | Listeria monocyto | 0 | 0 | 0.169342623 | 0.9999998255 | 0.01091018925 | 0.008124972845 | 0 | 0.0458033915 | 0 | 0.001119530663 | 0 |
| MARD_SAMN05410695_REFG_MMP05410695 | 2 | -0.2587063479 | Lactobacillales | Listeria | Listeria monocyto | 0 | 0 | 0.2466101071 | 1 | 0.01092225622 | 0.01177273929 | 0 | 0.05516805521 | 0 | 0.001619188934 | 0.05587791933 |
| MARD_SAMN05417990_REFG_MMP05417990 | 2 | -0.3501037397 | Enterobacterales | Vibrio | Vibrio splendidus | 0 | 0 | 0.1076159514 | 1.47E-05 | 0.001226159908 | 0.1225950765 | 0 | 0.1261697786 | 0 | 1.58E-05 | 0 |
| MARD_SAMN05418027_REFG_MMP05418027 | 2 | -0.5174470674 | Enterobacterales | Enterovibrio | Enterovibrio norv | 0 | 0 | 0.05217267517 | 7.93E-05 | 0.004245510325 | 0.1000016046 | 0 | 0.1462402698 | 0 | 8.07E-04 | 0 |
| MARD_SAMN05418038_REFG_MMP05418038 | 2 | -0.373920411 | Enterobacterales | Vibrio | Vibrio sp0028757 | 0 | 0 | 0.1389152278 | 3.09E-05 | 0.004170893933 | 0.02562615966 | 0 | 0.06936665901 | 0 | 3.73E-05 | 0 |
| MARD_SAMN05418053_REFG_MMP05418053 | 2 | -0.372441764 | Enterobacterales | Vibrio | Vibrio lentus_A | 0 | 0 | 0.150794171 | 3.60E-05 | 0.003185726244 | 0 | 0 | 0.06020215376 | 0 | 0.00549633205 | 0 |
| MARD_SAMN05418058_REFG_MMP05418058 | 2 | -0.3308203358 | Enterobacterales | Vibrio | Vibrio echinoidec | 0 | 0 | 0.145052049 | 3.82E-05 | 0 | 0.04532437482 | 0 | 0.08945431993 | 0 | 4.22E-05 | 0 |
| MARD_SAMN05418065_REFG_MMP05418065 | 2 | -0.5548076232 | Enterobacterales | Vibrio | Vibrio breoganii | 0 | 0.1171664276 | 0.06440945411 | 1.11E-04 | 0.002949462413 | 4.02E-04 | 0 | 0.07866407762 | 0 | 0 | 0 |
| MARD_SAMN05418067_REFG_MMP05418067 | 2 | -0.3557500345 | Enterobacterales | Vibrio | Vibrio kanaloae | 0 | 0 | 0.1691008798 | 3.16E-05 | 0 | 0 | 0 | 0.06091026796 | 0 | 0 | 0 |
| MARD_SAMN05418100_REFG_MMP05418100 | 2 | -0.3385720874 | Enterobacterales | Vibrio | Vibrio splendidus | 0 | 0 | 0.1600801678 | 3.96E-05 | 0.003185980738 | 0 | 0 | 0.05587883045 | 0 | 4.26E-05 | 0 |
| MARD_SAMN05418109_REFG_MMP05418109 | 2 | -0.3517628564 | Enterobacterales | Vibrio | Vibrio crassostre | 0 | 0 | 0.0966392968 | 3.91E-05 | 0.003187585181 | 0.006570931172 | 0 | 0.05621716344 | 0 | 4.25E-05 | 0 |
| MARD_SAMN05418116_REFG_MMP05418116 | 2 | -0.614418947 | Enterobacterales | Vibrio | Vibrio sp0028773 | 0 | 0 | 0.1452437761 | 5.77E-05 | 1.59E-04 | 0.005555555556 | 0 | 0.05882995564 | 0 | 4.29E-05 | 0 |
| MARD_SAMN05418161_REFG_MMP05418161 | 2 | -0.3546823642 | Enterobacterales | Shewanella | Shewanella sp00 | 0 | 0 | 0.04054347934 | 7.81E-05 | 0.008685826587 | 0 | 0 | 0.06903812171 | 0 | 1.27E-05 | 0 |
| MARD_SAMN05418174_REFG_MMP05418174 | 2 | -0.6607919555 | Enterobacterales | Vibrio | Vibrio sp0028733 | 0 | 0 | 0.15874332 | 1.10E-04 | 0.006375793947 | 0 | 0 | 0.06402435977 | 0 | 0 | 0 |
| MARD_SAMN05421511_REFG_MMP05421511 | 4 | -0.06022086312 | Oceanibaculales | Oceanibaculum | Oceanibaculum i | 0 | 0.002688826538 | 0.9999997274 | 0.9999997274 | 0.003249384981 | 6.00E-06 | 0 | 0.05187564873 | 0 | 7.72E-04 | 0 |
| MARD_SAMN05421538_REFG_MMP05421538 | 2 | -0.1040471639 | Rhodobacterales | Paracoccus | Paracoccus isopo | 3.90E-04 | 0 | 0.02004573074 | 1 | 0 | 5.68E-06 | 0 | 0.05262091642 | 0 | 0.0072508157 | 0.009084149543 |
| MARD_SAMN05421636_REFG_MMP05421636 | 2 | -0.07084163112 | Flavobacteriales | Pricia | Pricia antarctica | 0 | 0 | 0.04139303875 | 0.004637855776 | 0.1018670888 | 0.005295178583 | 0 | 0.04858241897 | 0 | 0.01056889213 | 0.07440239662 |
| MARD_SAMN05421668_REFG_MMP05421668 | 1 | -0.4285241046 | Bacillales_D | Halolactibacillus | Halolactibacillus | 0 | 0 | 1 | 4.22E-05 | 0.009061670798 | 1 | 0 | 0.0310346922 | 0 | 8.03E-04 | 0.01213031116 |
| MARD_SAMN05421688_REFG_MMP05421688 | 7 | -0.1782150719 | Rhodobacterales | Poseidonocella | Poseidonocella p | 0 | 0 | 0.1238656394 | 1.19E-06 | 0.03106833201 | 0.007566379443 | 0 | 0.0919652118 | 0 | 0.916692211 | 0 |
| MARD_SAMN05421734_REFG_MMP05421734 | 5 | -0.2536616875 | Bacillales_D | Pelagirhabdus | Pelagirhabdus al | 0 | 0 | 0.03506300405 | 4.05E-06 | 0 | 1.000000225 | 0 | 0.1531605576 | 7.69E-04 | 7.89E-04 | 0 |
| MARD_SAMN05421742_REFG_MMP05421742 | 5 | -0.02167768844 | Rhodospirillales | Roseospirillum | Roseospirillum pa | 3.80E-07 | 0 | 0.1150469211 | 2.03E-04 | 0.01621301318 | 0.9999998099 | 0 | 0.05801811674 | 0 | 7.81E-04 | 0.05783675547 |
| MARD_SAMN05421762_REFG_MMP05421762 | 5 | -0.2950065194 | Rhodobacterales | Pseudooceanicol | Pseudooceanicol | 0 | 0 | 0.1923507326 | 0.000125735604 | 0.01461047718 | 1 | 0 | 0.06459808614 | 0 | 7.78E-04 | 4.87E-06 |
| MARD_SAMN05421767_REFG_MMP05421767 | 1 | -0.3856715484 | Lactobacillales | Granulicatella | Granulicatella ba | 0 | 0 | 1.00000023 | 1.06E-04 | 1.00000023 | 1.00000023 | 0 | 0.06850242247 | 0 | 0 | 0 |
| MARD_SAMN05421769_REFG_MMP05421769 | 8 | -0.3579294794 | Flavobacteriales | Chryseobacteriu | Chryseobacteriu | 0.006304200192 | 0 | 1 | 1 | 3.24E-05 | 1 | 0 | 1 | 3.44E-04 | 0.01236772157 | 0.0511430923 |
| MARD_SAMN05421791_REFG_MMP05421791 | 5 | -0.2671542911 | Lactobacillales | Facklamia | Facklamia mirour | 0 | 0 | 0.04147919603 | 1.60E-05 | 0.1139203588 | 1 | 0 | 0.06655730472 | 0 | 0 | 0 |
| MARD_SAMN05421793_REFG_MMP05421793 | 2 | -0.3397156272 | Flavobacteriales | Epilithonimonas | Epilithonimonas l | 0 | 0 | 0.2140126245 | 1.13E-04 | 0.004142157019 | 0 | 0.04741489946 | 0.05945129029 | 0 | 8.03E-04 | 0.04493959885 |
| MARD_SAMN05421798_REFG_MMP05421798 | 5 | -0.3309921671 | Rhizobiales | Pseudovibrio | Pseudovibrio axi | 0 | 0 | 0.03174875789 | 3.21E-05 | 0.1962845539 | 1 | 0 | 0.0576132652 | 0 | 7.69E-04 | 0 |
| MARD_SAMN05421803_REFG_MMP05421803 | 5 | -0.2168586064 | Streptosporangia | Nocardiopsis | Nocardiopsis flav | 0 | 0 | 0.05538054273 | 2.40E-05 | 0.05074002898 | 1 | 0 | 0.1880947414 | 0 | 2.43E-05 | 0 |
| MARD_SAMN05421806_REFG_MMP05421806 | 2 | -0.1924932187 | Streptomycetales | Streptomyces | Streptomyces inc | 0 | 0 | 0.2482974105 | 1.48E-04 | 0.007093883322 | 0.03579412987 | 0 | 0.1343939995 | 0.001195404589 | 0 | 0 |
| MARD_SAMN05421823_REFG_MMP05421823 | 2 | -0.3755434071 | Cytophagales | Catalinimonas | Catalinimonas all | 0 | 0 | 0.02608281416 | 0.00344532954 | 0 | 0 | 1 | 0.2006021393 | 0 | 0.001566748776 | 5.78E-04 |
| MARD_SAMN05421831_REFG_MMP05421831 | 5 | -0.2406907706 | Pseudomonadale | Pseudospirillum | Pseudospirillum j | 3.13E-08 | 0 | 0.00154798047 | 0 | 0.08568505887 | 0.9850830763 | 0 | 6.97E-04 | 0 | 1.20E-04 | 0 |
| MARD_SAMN05421838_REFG_MMP05421838 | 5 | -0.04877232085 | Aquificales | Hydrogenivirga | Hydrogenivirga c | 0 | 0 | 0.08776982068 | 0.01666719998 | 0.001624816996 | 1.000000161 | 0 | 0.08933828466 | 1.23E-05 | 3.80E-05 | 0.0345447347 |
| MARD_SAMN05421843_REFG_MMP05421843 | 2 | -0.2235716415 | Rhodobacterales | Litoreibacter | Litoreibacter med | 0 | 0 | 0.0378096425 | 3.93E-05 | 0.2557321082 | 2.01E-05 | 0 | 0.068614719 | 0 | 0.003738461945 | 0 |
| MARD_SAMN05421848_REFG_MMP05421848 | 5 | -0.1943894628 | Pseudomonadale | Kushneria | Kushneria avicer | 0 | 0 | 0.03340599334 | 0.01666666667 | 0.001509280509 | 1 | 0 | 0.0404447403 | 0 | 1.31E-05 | 0.04824345732 |
| MARD_SAMN05421850_REFG_MMP05421850 | 4 | -0.1302457525 | Rhodobacterales | Lutimaribacter | Lutimaribacter sa | 2.52E-06 | 0 | 1 | 1.20E-04 | 1 | 0.008310140566 | 0 | 0.06436835986 | 0 | 0 | 0 |
| MARD_SAMN05421855_REFG_MMP05421855 | 2 | -0.1670592778 | Flavobacteriales | Ulvibacter | Ulvibacter litoralis | 0.01430359572 | 0 | 0.04884659556 | 0.04212156037 | 0.2938215494 | 0.007231293406 | 0 | 0.05183674804 | 0 | 0.002000237343 | 0 |
| MARD_SAMN05421867_REFG_MMP05421867 | 2 | 0.07017401674 | Actinomycetales | Cellulomonas | Cellulomonas ma | 0 | 0 | 0.02545155533 | 1.61E-04 | 0.2129491657 | 8.67E-06 | 0 | 0.05056716196 | 0 | 0 | 0 |
| MARD_SAMN05443431_REFG_MMP05443431 | 2 | -0.03938844724 | Flavobacteriales | Olleya | Olleya namhaens | 3.88E-07 | 0 | 0.1294516659 | 1 | 0.005394595165 | 6.59E-06 | 0 | 0.06819461813 | 0 | 8.34E-04 | 0 |
| MARD_SAMN05443551_REFG_MMP05443551 | 2 | -0.1430380409 | Rhodobacterales | Marivita | Marivita hallyeon | 8.77E-07 | 0 | 0.07454201964 | 4.30E-05 | 0.004185645751 | 0 | 0 | 0.05132983463 | 0 | 7.96E-04 | 0 |
| MARD_SAMN05443573_REFG_MMP05443573 | 2 | -0.1793892571 | Rhodobacterales | Celeribacter | Celeribacter indic | 0.03921230545 | 0.003690457386 | 0.1028616948 | 3.78E-05 | 0.00323799851 | 2.39E-07 | 0.04013324579 | 0.1045091521 | 0.001192128698 | 0.003399659293 | 0.01006840571 |
| MARD_SAMN05443635_REFG_MMP05443635 | 2 | -0.1462418609 | Rhodobacterales | Roseobacter | Roseobacter den | 0 | 0 | 0.1156765593 | 1.54E-06 | 0 | 8.45E-05 | 0 | 0.08082565225 | 0 | 1.27E-05 | 0.0274777583 |
| MARD_SAMN05443667_REFG_MMP05443667 | 2 | -0.1584331926 | Flavobacteriales | Flavobacterium | Flavobacterium g | 0 | 0 | 0.01943324957 | 3.18E-05 | 0.00280423042 | 0.009555454385 | 0 | 0.04543794749 | 0 | 0.005547937882 | 0.009889451454 |
| MARD_SAMN05443999_REFG_MMP05443999 | 5 | -0.08735394367 | Rhodobacterales | Roseovarius | Roseovarius azo | 0 | 0 | 0.04270436109 | 1.49E-04 | 0.003289798595 | 1 | 0 | 0.06364902836 | 0 | 7.81E-04 | 0 |
| MARD_SAMN05444000_REFG_MMP05444000 | 2 | -0.3165310393 | Rhodobacterales | Shimia | Shimia gijangens | 7.42E-09 | 0 | 0.08794066804 | 0.03333817158 | 0.09887301762 | 0.07788187497 | 0 | 0.952762344 | 0 | 6.67E-04 | 9.36E-05 |
| MARD_SAMN05444002_REFG_MMP05444002 | 2 | -0.06203333304 | Rhodobacterales | Vannielia | Vannielia litorea | 0 | 0 | 0.04442835012 | 2.66E-06 | 0.1647833184 | 0.01666666234 | 0 | 0.2249464115 | 0 | 1.22E-05 | 0 |
| MARD_SAMN05444003_REFG_MMP05444003 | 2 | -0.337550582 | Rhodobacterales | Cognatiyoonia | Cognatiyoonia se | 0 | 0 | 0.01340563806 | 2.43E-05 | 0.00665911766 | 0 | 0 | 0.05647608985 | 0 | 0.001182526703 | 0.01768938124 |
| MARD_SAMN05444004_REFG_MMP05444004 | 5 | -0.150108045 | Rhodobacterales | Jannaschia | Jannaschia faeci | 0 | 0 | 0.04719658314 | 4.58E-06 | 0.01134958615 | 1 | 0 | 0.06317514577 | 0 | 7.88E-04 | 0.01282446637 |
| MARD_SAMN05444006_REFG_MMP05444006 | 5 | -0.07161203606 | Rhodobacterales | Acidimangrovimo | Acidimangrovimo | 4.87E-06 | 0 | 0.02228089041 | 1.57E-05 | 0.003548422848 | 1 | 0 | 0.04938025224 | 0 | 8.48E-04 | 0.03934042843 |

| | | | | | | | | | | | | | | | | |
|---|---|---|---|---|---|---|---|---|---|---|---|---|---|---|---|---|
| MARD_SAMN05444007_REFG_MMP05444007 | 4 | -0.09311278885 | Rhodobacterales | Cribrihabitans | Cribrihabitans ma | 0 | 0 | 1 | 1.39E-04 | 0.01297076688 | 0.002693902144 | 0 | 0.07165956607 | 0 | 7.91E-04 | 0 |
| MARD_SAMN05444141_REFG_MMP05444141 | 2 | -0.3081093549 | Rhizobiales | Pseudovibrio | Pseudovibrio der | 0 | 0 | 0.05903937248 | 2.03E-06 | 0.04748782461 | 1.15E-04 | 0 | 0.07171347886 | 0 | 0 | 0 |
| MARD_SAMN05444143_REFG_MMP05444143 | 4 | -0.2743976093 | Flavobacteriales | Flavobacterium | Flavobacterium s | 0 | 0 | 1.000000214 | 0.114353794 | 1.000000214 | 0 | 0 | 0.04182558521 | 0 | 0 | 0 |
| MARD_SAMN05444148_REFG_MMP05444148 | 2 | -0.143589701 | Flavobacteriales | Winogradskyella | Winogradskyella | 0 | 0 | 0.03656264853 | 1.78E-04 | 0.01068706783 | 0 | 0 | 0.05635069657 | 0 | 7.97E-04 | 0 |
| MARD_SAMN05444149_REFG_MMP05444149 | 6 | -0.217328736 | Rhodobacterales | Pseudosulfitobac | Pseudosulfitobac | 0 | 0 | 0.1323464738 | 4.05E-05 | 1.000000162 | 0 | 0.1075350059 | 0.05061340598 | 0 | 1.05E-05 | 0.0381573905 |
| MARD_SAMN05444274_REFG_MMP05444274 | 4 | -0.08314916793 | Bacteroidales | Mariniphaga | Mariniphaga ana | 0 | 0 | 1 | 1.32E-04 | 0.09124168895 | 0.006452560704 | 0 | 0.3780223788 | 0 | 0 | 0 |
| MARD_SAMN05444278_REFG_MMP05444278 | 2 | -0.1073372398 | Flavobacteriales | Psychroflexus | Psychroflexus sa | 0 | 0 | 0.0380705021 | 4.36E-06 | 0.067374872 | 1.97E-08 | 0 | 0.05270741041 | 0 | 0 | 0.007425627291 |
| MARD_SAMN05444285_REFG_MMP05444285 | 2 | -0.2130176975 | Bacteroidales | Draconibacterium | Draconibacterium | 0 | 0 | 0.1089723765 | 0.03333333914 | 0.01302040747 | 0.03333333914 | 0 | 0.02066608452 | 0 | 1.25E-05 | 4.78E-05 |
| MARD_SAMN05444287_REFG_MMP05444287 | 8 | -0.3186662669 | Rhodobacterales | Octadecabacter | Octadecabacter t | 1.36E-05 | 0 | 0.9715259237 | 0.9999998042 | 0.02263123894 | 0.9999998042 | 0 | 0.0564378437 | 0 | 2.57E-05 | 0.01799187215 |
| MARD_SAMN05444336_REFG_MMP05444336 | 5 | -0.08837458339 | Rhodobacterales | Albimonas | Albimonas dongh | 0 | 0 | 0.02720781004 | 3.18E-05 | 0.0999940421 | 1.000000005 | 0 | 0.09739369285 | 0 | 7.70E-04 | 0 |
| MARD_SAMN05444339_REFG_MMP05444339 | 2 | -0.09980772125 | Rhodobacterales | Loktanella | Loktanella atrilute | 0 | 0 | 0.09732825176 | 0.9833333303 | 0.03532485677 | 0.002759875676 | 0.0166666636 | 0.08809995626 | 0 | 1.31E-05 | 0.0365270715 |
| MARD_SAMN05444340_REFG_MMP05444340 | 2 | -0.03823204119 | Rhodobacterales | Citreimonas | Citreimonas salir | 0 | 0 | 0.01426906935 | 1.10E-04 | 0 | 0 | 0 | 0.07827681583 | 0 | 0 | 0 |
| MARD_SAMN05444343_REFG_MMP05444343 | 2 | -0.3090633301 | Bacteroidales | Mangrovibacteriu | Mangrovibacteriu | 0 | 0 | 0.03502612252 | 7.88E-05 | 0 | 0.002802643808 | 0 | 0.04977948123 | 0 | 0 | 0.04199089381 |
| MARD_SAMN05444353_REFG_MMP05444353 | 8 | -0.06088677291 | Flavobacteriales | Polaribacter | Polaribacter doko | 0 | 0 | 0.1159127269 | 1 | 0.003395962977 | 1 | 0 | 0.04283704947 | 0 | 8.08E-04 | 8.84E-07 |
| MARD_SAMN05444358_REFG_MMP05444358 | 2 | -0.2849418101 | Rhodobacterales | Ruegeria | Ruegeria halocyr | 5.92E-08 | 0 | 0.113836968 | 9.77E-07 | 0.004166619564 | 0 | 0 | 0.2982611405 | 0 | 0 | 0 |
| MARD_SAMN05444359_REFG_MMP05444359 | 2 | -0.2661168662 | Chitinophagales | Lewinella_A | Lewinella_A agar | 0 | 0 | 0.05247282601 | 0.2748266727 | 0.004212710237 | 0.01666749942 | 0 | 0.06809073744 | 0 | 3.67E-05 | 0.06314909989 |
| MARD_SAMN05444365_REFG_MMP05444365 | 2 | 0.001532293622 | Mycobacteriales | Micromonospora | Micromonospora | 6.28E-06 | 0 | 0.09632925304 | 1.04E-04 | 0.039328891 | 0.001173814882 | 0 | 0.06330735543 | 0 | 0.02128522575 | 0 |
| MARD_SAMN05444380_REFG_MMP05444380 | 8 | -0.02694594037 | Bacteroidales | Thermophagus | Thermophagus x | 0.003133086989 | 0 | 0.2170908484 | 1 | 0.1339609097 | 1 | 0 | 0.04550222619 | 0 | 7.95E-04 | 0.002317518174 |
| MARD_SAMN05444398_REFG_MMP05444398 | 2 | -0.1541256079 | Rhodobacterales | Roseovarius | Roseovarius pac | 0 | 0 | 0.01949453737 | 2.52E-06 | 0.0166666633 | 0 | 0 | 0.06115569257 | 0 | 0 | 0.008056562462 |
| MARD_SAMN05444411_REFG_MMP05444411 | 6 | 0.002481104466 | Flavobacteriales | Lutibacter | Lutibacter oricola | 1.37E-08 | 0 | 0.04227260752 | 0.01671853036 | 0.9525898447 | 3.13E-08 | 0 | 0.07188527767 | 0 | 0 | 0 |
| MARD_SAMN05444412_REFG_MMP05444412 | 3 | -0.2488618603 | Cytophagales | Rhodonellum | Rhodonellum psy | 0 | 0 | 0.01365528644 | 0.9999999964 | 0.01723432112 | 0.9999999964 | 0 | 0.09646679124 | 0 | 0.9999999964 | 0.01000401248 |
| MARD_SAMN05444413_REFG_MMP05444413 | 3 | -0.01585431439 | Rhodobacterales | Roseivivax | Roseivivax marin | 0 | 0 | 0.06123345338 | 1.11E-04 | 0.06747489766 | 0.9999998195 | 0 | 0.04597301921 | 0 | 0.9999998195 | 0.04311134932 |
| MARD_SAMN05444414_REFG_MMP05444414 | 5 | -0.2536838301 | Rhodobacterales | Roseovarius | Roseovarius mar | 0 | 0 | 0.03922005338 | 1.43E-04 | 0.1210099635 | 1 | 0 | 0.01758123572 | 0 | 0.001517949699 | 0 |
| MARD_SAMN05444415_REFG_MMP05444415 | 5 | -0.08546372206 | Rhodobacterales | Salipiger | Salipiger profund | 0 | 0 | 0.02634139485 | 5.39E-05 | 0.1764236347 | 1 | 0 | 1 | 3.73E-04 | 0.001912692501 | 0 |
| MARD_SAMN05444481_REFG_MMP05444481 | 2 | -0.2376118581 | Flavobacteriales | Flavobacterium | Flavobacterium fi | 0 | 0 | 0.03036364805 | 9.88E-05 | 0.0013568661 | 0.001238636246 | 0 | 1 | 0 | 1.59E-06 | 0 |
| MARD_SAMN05444482_REFG_MMP05444482 | 2 | -0.07693038194 | Flavobacteriales | Gillisia | Gillisia mitskevicl | 0 | 0 | 0.03043468486 | 1.88E-04 | 0.1312059129 | 0.004878237081 | 0 | 0.07403382273 | 0 | 0 | 0.04321821118 |
| MARD_SAMN05444483_REFG_MMP05444483 | 6 | -0.1863172194 | Flavobacteriales | Salegentibacter | Salegentibacter e | 3.51E-06 | 0 | 0.03597191045 | 3.82E-05 | 1 | 1.71E-05 | 0 | 1 | 0 | 0 | 0.01062025353 |
| MARD_SAMN05444484_REFG_MMP05444484 | 2 | -0.2489249138 | Flavobacteriales | Flavobacterium | Flavobacterium c | 0 | 0 | 0.09652766843 | 0.9833331806 | 0.2398435598 | 0.008042722626 | 1.59E-04 | 0.05935565621 | 1.59E-04 | 1.21E-05 | 8.59E-04 |
| MARD_SAMN05444486_REFG_MMP05444486 | 5 | -0.1407403022 | Rhodobacterales | Lentibacter | Lentibacter algar | 0 | 0 | 0.03561241877 | 1.77E-04 | 0.04586251132 | 0.9833336066 | 0 | 0.05620495519 | 0 | 0 | 0 |
| MARD_SAMN05444487_REFG_MMP05444487 | 5 | -0.3459778256 | Thermoactinomy | Marininema | Marininema mes | 0 | 0 | 0.1863296021 | 8.62E-05 | 0.003144553741 | 0.9999997887 | 0 | 0.04903475091 | 0 | 0 | 0 |
| MARD_SAMN05444714_REFG_MMP05444714 | 5 | -0.2925904701 | Rhodobacterales | Yoonia | Yoonia litorea | 5.02E-04 | 0 | 0.1414942763 | 1.74E-04 | 0.01087622066 | 1 | 0 | 0.05050402387 | 0 | 7.92E-04 | 0.001013536843 |
| MARD_SAMN05444851_REFG_MMP05444851 | 5 | -0.2393504522 | Rhodobacterales | Aliiroseovarius | Aliiroseovarius se | 0 | 0 | 0.2191332281 | 6.47E-06 | 2.02E-04 | 0.999999826 | 0 | 0.07932970743 | 0 | 3.90E-05 | 0.04096146811 |
| MARD_SAMN05504940_REFG_MMP05504940 | 8 | -0.03442804889 | Peptostreptococ | Caminicella | Caminicella spor | 0 | 0 | 1 | 1 | 0.004643370114 | 1 | 0 | 0.3154292428 | 0 | 0 | 0 |
| MARD_SAMN05504969_REFG_MMP05504969 | 2 | -0.08582083312 | Tissierellales | Thermohalobacte | Thermohalobacte | 0 | 0 | 0.2105356 | 4.96E-05 | 0.003198397743 | 0.09333337969 | 0 | 0.05943787402 | 0 | 4.27E-05 | 0 |
| MARD_SAMN05505091_REFG_MMP05505091 | 2 | -0.7355644011 | Enterobacterales | Vibrio | Vibrio anguillarur | 0 | 0 | 0.06568241291 | 8.99E-05 | 0.001864147843 | 9.44E-08 | 0 | 0.3269789543 | 0 | 1.38E-05 | 0.1666666984 |
| MARD_SAMN05526266_REFG_MMP05526266 | 5 | -0.4301596664 | Bacillales_I | Bacillus_AV | Bacillus_AV solin | 0 | 0 | 0.01635172987 | 0.01666666272 | 0.05065635667 | 0.9833333333 | 0 | 0.07161029956 | 0 | 0 | 0 |
| MARD_SAMN05567582_REFG_MMP05567582 | 8 | -0.1932257425 | Cytophagales | Fabibacter | Fabibacter misak | 0 | 0 | 0.02668228024 | 0.9833335218 | 0.01722492301 | 0.9834066794 | 0 | 0.07395636543 | 0 | 0.001560588685 | 4.47E-04 |
| MARD_SAMN05582966_REFG_MMP05582966 | 2 | -0.3840512496 | Rhodobacterales | Amylibacter | Amylibacter sedi | 0 | 0.00134407229 | 0.04172364293 | 7.51E-05 | 0.04208736786 | 0.001193694517 | 0 | 0.07270552486 | 0 | 0 | 0 |
| MARD_SAMN05660282_REFG_MMP05660282 | 6 | -0.33763512 | Mycobacteriales | Corynebacterium | Corynebacterium | 4.92E-06 | 0 | 0.0420079912 | 4.04E-05 | 1 | 0 | 0 | 0.07120202782 | 0 | 1.58E-05 | 0 |
| MARD_SAMN05660313_REFG_MMP05660313 | 5 | -0.0320494341 | Flavobacteriales | Cellulophaga | Cellulophaga fuc | 0.002820817503 | 0 | 0.1298276759 | 1.22E-04 | 4.65E-05 | 1.000000003 | 0 | 0.2965195352 | 0 | 7.63E-04 | 0 |
| MARD_SAMN05660337_REFG_MMP05660337 | 6 | -0.2144799179 | Desulfovibrionale | Maridesulfovibrio | Maridesulfovibrio | 0 | 0 | 0.04082037801 | 1.22E-04 | 1 | 0 | 0 | 0.0713648196 | 0 | 0 | 0.01103619466 |
| MARD_SAMN05660413_REFG_MMP05660413 | 2 | -0.193846865 | Flavobacteriales | Salegentibacter | Salegentibacter f | 0 | 0 | 0.1096804944 | 1.23E-04 | 0.1715366718 | 0.005445317044 | 0 | 0.1616563736 | 0 | 0.001602104123 | 0 |
| MARD_SAMN05660420_REFG_MMP05660420 | 5 | -0.1765986652 | Desulfuromonada | Desulfuromusa | Desulfuromusa k | 0 | 0 | 0.3848651434 | 1.63E-05 | 0.00228701281 | 0.9500000238 | 0 | 0.1620696874 | 0 | 0.009776736535 | 0 |
| MARD_SAMN05660429_REFG_MMP05660429 | 1 | -0.4874382055 | Enterobacterales | Thalassotalea_D | Thalassotalea_D | 5.48E-05 | 0 | 0.9999999967 | 7.17E-06 | 0.0404460573 | 0.6846711293 | 0 | 0.04928835195 | 0 | 1.92E-04 | 0.07886069327 |
| MARD_SAMN05660479_REFG_MMP05660479 | 2 | -0.2101337614 | Pseudomonadale | Microbulbifer | Microbulbifer ther | 0 | 1.85E-04 | 0.04427518389 | 3.21E-06 | 0.2178755095 | 0.03398038641 | 0 | 0.04449817844 | 0 | 4.90E-06 | 0 |
| MARD_SAMN05660649_REFG_MMP05660649 | 5 | 0.03070455796 | Desulfotomacula | Desulfallas | Desulfallas arctic | 0 | 0 | 0.07666461346 | 7.80E-05 | 0 | 1 | 0 | 0.2768311627 | 0 | 8.26E-04 | 0.02962757472 |
| MARD_SAMN05660691_REFG_MMP05660691 | 5 | -0.4705442962 | Enterobacterales | Rheinheimera | Rheinheimera pa | 0 | 0 | 0.04941554221 | 1.18E-04 | 0.01856374865 | 1.000000149 | 0 | 0.05319265125 | 0 | 7.54E-04 | 0.1576120748 |
| MARD_SAMN05660836_REFG_MMP05660836 | 5 | -0.1148028764 | Syntrophobacter | Thermodesulforh | Thermodesulforh | 0 | 0.05742049927 | 0.06832480972 | 5.95E-05 | 0 | 1 | 0 | 0.1377679642 | 0 | 8.11E-04 | 0 |
| MARD_SAMN05660971_REFG_MMP05660971 | 4 | -0.2520586859 | Pseudomonadale | Halomonas | Halomonas cupic | 0.001868811416 | 0 | 0.9837684454 | 9.03E-05 | 0.03675729854 | 0.005488036569 | 0 | 0.04683721541 | 0 | 1.25E-05 | 0 |
| MARD_SAMN05661223_REFG_MMP05661223 | 2 | -0.1925303819 | Pseudomonadale | Teredinibacter | Teredinibacter ha | 1.65E-07 | 0 | 0.3133171268 | 6.68E-06 | 0.03594187378 | 8.46E-07 | 0 | 0.06620214694 | 0 | 0.01092919129 | 5.11E-04 |
| MARD_SAMN05687753_REFG_MMP05687753 | 4 | -0.3179489821 | Cytophagales | Reichenbachiella | Reichenbachiella | 0 | 0 | 0.7408440253 | 6.51E-05 | 0.004390869777 | 0.09228241648 | 0 | 0.2006033875 | 0 | 5.87E-04 | 4.32E-04 |
| MARD_SAMN05687755_REFG_MMP05687755 | 5 | -0.6414541016 | Enterobacterales | Pseudoalteromor | Pseudoalteromor | 0 | 0 | 0.04333235657 | 1.30E-04 | 0.09520253974 | 1 | 0.04797228613 | 0.06847662887 | 0 | 0 | 0.0573964079 |
| MARD_SAMN05728473_REFG_MMP05728473 | 2 | -0.2657007223 | Flavobacteriales | Flavobacterium | Flavobacterium ti | 0 | 0 | 0.07808240293 | 0.01674998496 | 0.05038407082 | 0.05000728153 | 0 | 0.07158177288 | 0 | 0.003760841097 | 0.05083980173 |
| MARD_SAMN05791069_REFG_MMP05791069 | 2 | -0.1476421562 | Pseudomonadale | Kushneria | Kushneria phosp | 0 | 0 | 0.02276383484 | 3.25E-05 | 0.1474625277 | 0 | 0 | 0.05623695405 | 0 | 0.004241136304 | 0 |
| MARD_SAMN05792885_REFG_MMP05792885 | 2 | -0.2038416447 | Streptomycetales | Streptomyces | Streptomyces gri | 0 | 0.005756215993 | 0.170196936 | 9.61E-05 | 0.01938184128 | 1.65E-04 | 0 | 0.03704174468 | 0 | 0.001016022013 | 0 |
| MARD_SAMN05818643_REFG_MMP05818643 | 2 | -0.075358426 | Rhizobiales | Acuticoccus | Acuticoccus yang | 0 | 0 | 0.02841319052 | 2.78E-06 | 0.01522106723 | 0 | 0 | 0.05125914208 | 0 | 0.007366590074 | 0 |
| MARD_SAMN05897974_REFG_MMP05897974 | 2 | -0.1500778855 | Mycobacteriales | Mycobacterium | Mycobacterium a | 0 | 0 | 0.02451594094 | 3.58E-04 | 0.5684345934 | 0.002192622443 | 0 | 0.04970941533 | 0 | 7.97E-04 | 0.003132989681 |
| MARD_SAMN05897984_REFG_MMP05897984 | 5 | -0.1772524328 | Mycobacteriales | Mycobacterium | Mycobacterium s | 0 | 0 | 0.048968761 | 2.30E-04 | 0.01959838659 | 0.9666672571 | 0 | 0.06141482344 | 0 | 7.28E-04 | 0 |
| MARD_SAMN05897987_REFG_MMP05897987 | 2 | -0.1570476428 | Mycobacteriales | Mycobacterium | Mycobacterium c | 8.15E-08 | 0 | 0.03720935391 | 4.53E-05 | 0.02236302378 | 0 | 0 | 0.09973980114 | 0 | 0.001619345869 | 0.02907688882 |
| MARD_SAMN05915707_REFG_MMP05915707 | 2 | -0.1381992676 | Flavobacteriales | Formosa | Formosa algae | 0 | 0.0541581765 | 0.0643302129 | 0.01666667008 | 0.001040896789 | 0.004407382015 | 0 | 0.1174346253 | 0 | 1.31E-05 | 0.03524410409 |
| MARD_SAMN05928563_REFG_MMP05928563 | 5 | -0.189405322 | Rhodobacterales | Nioella | Nioella nitratiredu | 6.53E-09 | 0 | 0.1470562867 | 5.68E-07 | 0.1455213328 | 1 | 0 | 0.06828149576 | 0 | 1.29E-05 | 2.00E-06 |
| MARD_SAMN06006700_REFG_MMP06006700 | 2 | -0.2396629993 | Actinomycetales | Boudabousia | Boudabousia ma | 0 | 0 | 0.08240787297 | 4.94E-05 | 0.006225159956 | 0.1984710671 | 0 | 0.009960243674 | 0 | 0 | 0 |
| MARD_SAMN06028227_REFG_MMP06028227 | 4 | -0.7628232597 | Enterobacterales | Edwardsiella | Edwardsiella pisc | 3.49E-06 | 0 | 1.000000004 | 0.01675928156 | 0.9840137151 | 0 | 0 | 0.06199730086 | 0 | 5.68E-08 | 0 |
| MARD_SAMN06075351_REFG_MMP06075351 | 6 | -0.1043130614 | Flavobacteriales | Nonlabens | Nonlabens xylani | 0 | 0 | 0.02523966226 | 1.48E-04 | 0.9852527175 | 0.003438530812 | 4.66E-07 | 0.058390863 | 0 | 1.33E-05 | 0 |
| MARD_SAMN06075357_REFG_MMP06075357 | 2 | -0.1826195769 | Flavobacteriales | Tenacibaculum | Tenacibaculum s | 0 | 0 | 0.02998673611 | 1.72E-06 | 0.01666667155 | 0 | 0 | 0.07191944228 | 0 | 0 | 0 |
| MARD_SAMN06090799_REFG_MMP06090799 | 6 | -0.375446034 | Verrucomicrobial | Rubritalea | Rubritalea profur | 2.10E-06 | 0 | 0.05039510568 | 1.37E-04 | 1 | 0 | 0.004269838323 | 0.06032217871 | 0 | 1.47E-05 | 0.005459125246 |
| MARD_SAMN06091621_REFG_MMP06091621 | 5 | -0.01939089661 | Rhodothermales | Salinibacter | Salinibacter sp00 | 0 | 0 | 0.2725190013 | 0 | 0.006769702321 | 1.000000175 | 0 | 0.04154775281 | 0 | 0 | 0 |
| MARD_SAMN06133338_REFG_MMP06133338 | 2 | -0.5232297785 | Enterobacterales | Photobacterium | Photobacterium a | 0 | 0 | 0.04616252667 | 1.11E-05 | 0 | 1.02E-05 | 0 | 0.05830026841 | 0 | 5.96E-06 | 0 |
| MARD_SAMN06133354_REFG_MMP06133354 | 2 | -0.2520050683 | Enterobacterales | Psychrosphaera | Psychrosphaera | 0 | 0 | 0.06347854706 | 2.47E-05 | 0.002290507166 | 0.05422792805 | 0 | 0.07611717074 | 0 | 3.06E-05 | 0 |
| MARD_SAMN06133355_REFG_MMP06133355 | 2 | -0.3816160112 | Enterobacterales | Vibrio | Vibrio chagasii | 0 | 0 | 0.09650266797 | 1.43E-05 | 0.007715434974 | 5.33E-06 | 0 | 0.06504780247 | 0 | 1.67E-05 | 0 |
| MARD_SAMN06133456_REFG_MMP06133456 | 6 | -0.1313717629 | Flavobacteriales | Polaribacter | Polaribacter gang | 0 | 0 | 0.02801309751 | 5.04E-05 | 0.9833333333 | 3.08E-06 | 0 | 0.04093939278 | 0 | 0 | 0.04533602214 |
| MARD_SAMN06134493_REFG_MMP06134493 | 2 | -0.5612203101 | Enterobacterales | Photobacterium | Photobacterium p | 0 | 0 | 0.04914703389 | 2.23E-06 | 0.003352454938 | 0.007334873317 | 0 | 0.1450646482 | 0 | 1.67E-04 | 0 |
| MARD_SAMN06134494_REFG_MMP06134494 | 2 | -0.4336557908 | Enterobacterales | Aliivibrio | Aliivibrio sifiae_A | 0 | 0 | 0.04649140196 | 7.72E-06 | 0.003377548503 | 8.26E-07 | 0 | 0.03474733979 | 0 | 1.85E-05 | 0 |

| | | | | | | | | | | | | | | | | |
|---|---|---|---|---|---|---|---|---|---|---|---|---|---|---|---|---|
| MARD_SAMN06134495_REFG_MMP06134495 | 5 | -0.4417398825 | Enterobacterales | Aliivibrio | Aliivibrio sifiae | 0 | 0 | 0.1332516186 | 5.06E-06 | 0.01014875225 | 0.9833333356 | 7.66E-04 | 0.05925558283 | 0 | 2.25E-06 | 0.005360382761 |
| MARD_SAMN06140193_REFG_MMP06140193 | 2 | -0.4527382071 | Pseudomonadale | Pseudomonas_E | Pseudomonas_E | 0 | 0 | 0.03539850539 | 3.61E-06 | 0.07632803029 | 5.95E-05 | 0 | 0.1098534471 | 0 | 0 | 0.027920137 |
| MARD_SAMN06264851_REFG_MMP06264851 | 6 | -0.2019458298 | Flavobacteriales | Ulvibacter | Ulvibacter antarc | 0 | 0 | 0.01195518262 | 2.33E-06 | 0.9833332054 | 0.004060261112 | 0 | 0.2669369984 | 0 | 1.29E-05 | 0 |
| MARD_SAMN06264853_REFG_MMP06264853 | 4 | -0.2017525672 | Bacteroidales | Marinifilum | Marinifilum flexuo | 4.83E-08 | 0 | 0.9999999966 | 0.01666666322 | 0.4281301539 | 0 | 0 | 0.0661610372 | 0 | 0 | 0.02494258999 |
| MARD_SAMN06265172_REFG_MMP06265172 | 2 | -0.08517331377 | Flavobacteriales | Ichthyenterobacte | Ichthyenterobact | 0 | 0 | 0.03196854533 | 0.9999997388 | 0.01666667146 | 0.002679558497 | 0.009748443409 | 0.05350920459 | 0 | 2.74E-05 | 0 |
| MARD_SAMN06265217_REFG_MMP06265217 | 2 | -0.3050371434 | Cytophagales | Fabibacter | Fabibacter pacifi | 0 | 0 | 0.02912703856 | 4.26E-05 | 7.70E-04 | 9.06E-06 | 0 | 0.04010904512 | 0 | 0.004008865184 | 0 |
| MARD_SAMN06265352_REFG_MMP06265352 | 5 | -0.05523311688 | Sphingomonadal | Eilatimonas | Eilatimonas mille | 7.41E-07 | 0 | 0.1344684677 | 1.51E-04 | 0.002762038598 | 1.000000185 | 0.02656749301 | 0.05996268026 | 0 | 7.87E-04 | 0 |
| MARD_SAMN06265372_REFG_MMP06265372 | 6 | -0.077467534 | Rhodobacterales | Roseovarius | Roseovarius halo | 0 | 0 | 0.03618886429 | 1.45E-04 | 1 | 0.002326933683 | 0 | 0.060249998 | 0 | 0 | 0 |
| MARD_SAMN06265796_REFG_MMP06265796 | 8 | 0.001106872521 | Flavobacteriales | Lutibacter | Lutibacter oceani | 0 | 0 | 0.1586446335 | 0.9333374538 | 0.001218752163 | 0.9500930287 | 4.24E-04 | 0.09180286701 | 0 | 0.001605545724 | 0 |
| MARD_SAMN06299115_REFG_MMP06299115 | 5 | -0.1897743829 | Pseudomonadale | Salinicola | Salinicola haloph | 0 | 0 | 0.03945414274 | 4.59E-05 | 0 | 1.000000044 | 0 | 0.2075983288 | 0 | 5.88E-04 | 0.01340306964 |
| MARD_SAMN06299116_REFG_MMP06299116 | 5 | -0.2075118199 | Pseudomonadale | Salinicola | Salinicola acropo | 0 | 0 | 0.1194132199 | 9.01E-05 | 0.003208348672 | 0.9999999975 | 0 | 0.04681622145 | 0 | 0.003674223524 | 0 |
| MARD_SAMN06299117_REFG_MMP06299117 | 5 | -0.1834791698 | Pseudomonadale | Salinicola | Salinicola peritric | 0 | 1.01E-04 | 0.04600074161 | 4.42E-05 | 0 | 1 | 0 | 0.2655006131 | 0 | 8.06E-04 | 0 |
| MARD_SAMN06299118_REFG_MMP06299118 | 2 | -0.2094742569 | Pseudomonadale | Salinicola | Salinicola salariu | 0 | 0 | 0.02556917591 | 3.22E-05 | 0.003593481823 | 1.22E-05 | 0 | 1 | 0 | 8.59E-04 | 0.03815800784 |
| MARD_SAMN06644953_REFG_MMP06644953 | 7 | -0.1334448721 | Rhodobacterales | Marinibacterium | Marinibacterium : | 2.43E-07 | 0 | 0.9838394162 | 9.92E-06 | 0.9833608197 | 7.96E-05 | 0 | 3.67E-04 | 0 | 0.999999995 | 5.34E-04 |
| MARD_SAMN06699246_REFG_MMP06699246 | 5 | 0.03889177763 | Rhodothermales | Salinibacter | Salinibacter rube | 0 | 0 | 0.1042169601 | 1.53E-04 | 0.0178712679 | 1 | 0 | 0.05426102896 | 0 | 7.78E-04 | 0 |
| MARD_SAMN06948887_REFG_MMP06948887 | 2 | -0.2141061933 | Burkholderiales | Alcaligenes | Alcaligenes aqua | 0.009177682285 | 0 | 0.03867010768 | 0.9999999337 | 0.007136866032 | 0.3500066771 | 0.005758818748 | 0.06607742904 | 0.001282046293 | 4.62E-04 | 0 |
| MARD_SAMN07201002_REFG_MMP07201002 | 2 | -0.4399269827 | Staphylococcales | Staphylococcus | Staphylococcus ( | 9.42E-04 | 0 | 0.05977502475 | 0.9999999967 | 0.1170796749 | 0 | 0 | 0.08035784093 | 0 | 0 | 0 |
| MARD_SAMN07249100_REFG_MMP07249100 | 2 | -0.5695821021 | Bacillales_D | Terribacillus | Terribacillus sacc | 0 | 0 | 0.06830540819 | 2.08E-04 | 0.007499217754 | 0.03333607591 | 0 | 0.2513303055 | 0 | 0 | 0 |
| MARD_SAMN07327725_REFG_MMP07327725 | 2 | -0.5683968703 | Enterobacterales | Photobacterium | Photobacterium I | 0 | 0 | 0.07038835205 | 2.02E-05 | 0.003883041552 | 0.05890047326 | 0 | 0.07209027315 | 0 | 2.98E-05 | 0 |
| MARD_SAMN07327728_REFG_MMP07327728 | 4 | -0.5394152452 | Enterobacterales | Photobacterium | Photobacterium a | 0 | 0 | 0.8036697598 | 1.25E-04 | 0.0031772663 | 0.003237270535 | 0 | 0.06389050025 | 0 | 1.98E-06 | 0 |
| MARD_SAMN07327780_REFG_MMP07327780 | 2 | -0.5332702302 | Enterobacterales | Photobacterium | Photobacterium s | 0 | 0 | 0.0253320036 | 1.22E-04 | 0.003206409586 | 0 | 0 | 0.05617651978 | 0 | 0 | 0 |
| MARD_SAMN07357117_REFG_MMP07357117 | 3 | -0.6685350876 | Enterobacterales | Aeromonas | Aeromonas veroi | 0 | 0 | 0.09543255984 | 5.54E-05 | 0.04341053432 | 1.000000071 | 0 | 0.0670031905 | 0 | 1.000000071 | 0 |
| MARD_SAMN07415057_REFG_MMP07415057 | 2 | -0.3536745152 | Pseudomonadale | Agaribacterium | Agaribacterium s | 0 | 0 | 0.03964615802 | 3.52E-06 | 0.1200339434 | 9.62E-05 | 0 | 0.02595969612 | 9.14E-06 | 2.84E-05 | 0 |
| MARD_SAMN07424250_REFG_MMP07424250 | 2 | -0.02762159244 | Rhizobiales | Rhodobium | Rhodobium orien | 5.07E-05 | 0 | 0.2124030473 | 9.19E-07 | 0.008296243218 | 0.01666666667 | 0 | 0.1018025589 | 0 | 2.63E-05 | 0 |
| MARD_SAMN07621379_REFG_MMP07621379 | 2 | -0.1082810034 | Flavobacteriales | Lacinutrix | Lacinutrix veneru | 1.19E-06 | 0 | 0.03917370283 | 1.12E-04 | 0 | 0 | 0 | 0.0682293598 | 0 | 0 | 0 |
| MARD_SAMN07621896_REFG_MMP07621896 | 2 | -0.05015545968 | Flavobacteriales | Maribacter_B | Maribacter_B vac | 0 | 0 | 0.0371622341 | 3.04E-05 | 0.008032119142 | 0.0166681659 | 0 | 0.04726165889 | 0 | 0.003240292164 | 0.1717795259 |
| MARD_SAMN07626136_REFG_MMP07626136 | 5 | -0.4366599966 | Enterobacterales | Alteromonas | Alteromonas grac | 0 | 1.50E-05 | 0.1859215187 | 1.20E-04 | 0.003707563367 | 1.000000006 | 0 | 0.07427680719 | 0 | 0 | 0.0364619966 |
| MARD_SAMN07739585_REFG_MMP07739585 | 2 | -0.04931930398 | Campylobacteral | Campylobacter_I | Campylobacter_I | 0 | 0 | 0.02866534047 | 2.58E-05 | 0.004877288158 | 0.02975461186 | 0 | 0.05718630361 | 0 | 0.01671622881 | 0.07262223873 |
| MARD_SAMN07811444_REFG_MMP07811444 | 2 | -0.06369805064 | Mycobacteriales | Mycobacterium | Mycobacterium n | 0 | 0 | 0.06273529359 | 2.49E-05 | 0.003240658841 | 0.006142219516 | 0 | 0.06237304792 | 0 | 2.13E-06 | 0 |
| MARD_SAMN07818924_REFG_MMP07818924 | 2 | -0.6360806501 | Enterobacterales | Vibrio | Vibrio parahaem | 0 | 0 | 0.03102083108 | 1.68E-07 | 0.04878181328 | 7.68E-08 | 0 | 0.05699569712 | 0 | 0.007766137217 | 2.56E-05 |
| MARD_SAMN07818930_REFG_MMP07818930 | 2 | -0.4701454733 | Enterobacterales | Bowmanella | Bowmanella deni | 0 | 0 | 0.02820782694 | 0.003573440481 | 0.2018784284 | 0.01673993473 | 0 | 0.04632402604 | 0 | 0.01432286272 | 0.03921879238 |
| MARD_SAMN07961447_REFG_MMP07961447 | 2 | 0.002312245966 | Nevskiales | Salinisphaera | Salinisphaera ore | 0 | 0 | 0.01789787341 | 1.95E-06 | 0.05689214507 | 0.03970377195 | 0 | 0.0725750144 | 0 | 0.03333333333 | 0.06536842974 |
| MARD_SAMN07961448_REFG_MMP07961448 | 6 | -0.1309850772 | Nevskiales | Salinisphaera | Salinisphaera jap | 0 | 0 | 0.03230653483 | 0.05196630885 | 0.9853551459 | 0 | 0 | 0.03427346988 | 0 | 8.12E-04 | 0.06919699311 |
| MARD_SAMN07980965_REFG_MMP07980965 | 7 | -0.08835529957 | Euzebyales | Euzebya | Euzebya rosea | 0 | 0 | 0.9999997918 | 0.9500103481 | 0.003248120922 | 0.05000000633 | 0 | 0.1353327302 | 0 | 0.9502692844 | 0.05246105445 |
| MARD_SAMN07998568_REFG_MMP07998568 | 6 | -0.5556542783 | Lactobacillales | Streptococcus | Streptococcus ag | 0.2901872706 | 0.0100420375 | 0.03768137996 | 1.17E-04 | 0.9999998013 | 0 | 0 | 0.05745674425 | 0 | 0 | 0 |
| MARD_SAMN08099983_REFG_MMP08099983 | 6 | -0.2344527033 | Flavobacteriales | Arenibacter | Arenibacter catal | 0.108552163 | 0 | 0.1077265138 | 4.61E-05 | 0.9668013042 | 2.72E-06 | 0.01257965502 | 0.07990860744 | 0 | 0 | 6.25E-07 |
| MARD_SAMN08100074_REFG_MMP08100074 | 2 | -0.3261846227 | Bacillales_D | Oceanobacillus | Oceanobacillus a | 0 | 0 | 0.05106906081 | 1.00000021 | 0.06965447036 | 0 | 0 | 1.00000021 | 0 | 0.01321581558 | 0 |
| MARD_SAMN08100097_REFG_MMP08100097 | 5 | -0.3817131279 | Bacillales_D | Oceanobacillus | Oceanobacillus c | 0.004108385606 | 1.84E-05 | 0.4204159183 | 0.001806582624 | 0.05976110462 | 0.95 | 0 | 0.05495974231 | 0.01666666667 | 4.47E-04 | 0 |
| MARD_SAMN08122818_REFG_MMP08122818 | 2 | -0.3305517166 | Pseudomonadale | Stutzerimonas | Stutzerimonas st | 0 | 0 | 0.0441074532 | 5.21E-05 | 0.00348008116 | 0.00390594781 | 1.91E-06 | 0.05832419941 | 0 | 0 | 0.03688418802 |
| MARD_SAMN08131153_REFG_MMP08131153 | 1 | -0.04364307803 | Caulobacterales | Marinicaulis | Marinicaulis flavu | 3.85E-04 | 0 | 0.7888188242 | 0 | 0.08398858673 | 0.633333329 | 0 | 0.05989350368 | 0 | 0.005747866012 | 0 |
| MARD_SAMN08149906_REFG_MMP08149906 | 2 | -0.3650990461 | Pseudomonadale | Stutzerimonas | Stutzerimonas st | 0 | 0 | 0.1296855959 | 1.38E-04 | 0.003658794755 | 0 | 0 | 1.00000019 | 0 | 7.95E-04 | 2.32E-04 |
| MARD_SAMN08202528_REFG_MMP08202528 | 2 | -0.2672263401 | Rhodobacterales | Yoonia | Yoonia maritima_ | 0 | 0 | 0.0618781679 | 1.77E-05 | 0.004978575958 | 0.003236232714 | 0 | 0.06383145127 | 0 | 4.22E-05 | 0 |
| MARD_SAMN08204758_REFG_MMP08204758 | 2 | -0.6045638178 | Enterobacterales | Vibrio | Vibrio diabolicus | 0.001566985599 | 0 | 0.02143112011 | 2.68E-06 | 0.001245477622 | 0.1166666756 | 3.78E-04 | 0.9064498545 | 0 | 2.75E-06 | 0.06649251429 |
| MARD_SAMN08271789_SAGS_MMP08271789 | 4 | -0.01433095381 | Opitutales | PNEW01 | PNEW01 sp0029 | 0 | 0 | 0.8412956142 | 0 | 0 | 0.001871876594 | 0 | 0.1669761369 | 0 | 0.01103599678 | 0 |
| MARD_SAMN08271940_REFG_MMP08271940 | 4 | -0.1120667293 | Xanthomonadale | Stenotrophomona | Stenotrophomona | 0 | 0 | 0.9999997053 | 6.78E-05 | 0.0352108214 | 5.89E-06 | 0 | 0.06579779122 | 0 | 8.03E-04 | 0 |
| MARD_SAMN08295223_REFG_MMP08295223 | 2 | -0.4434742581 | Pseudomonadale | Acinetobacter | Acinetobacter sp | 7.77E-07 | 0 | 0.03920574096 | 1.67E-06 | 0.01666666386 | 3.81E-07 | 0 | 0.03682712688 | 0 | 0 | 0 |
| MARD_SAMN08295361_REFG_MMP08295361 | 6 | -0.1475995467 | Flavobacteriales | Aquimarina | Aquimarina sedir | 0 | 0 | 0.1358071694 | 9.17E-05 | 1 | 1 | 0 | 0.08198541146 | 0 | 0 | 0 |
| MARD_SAMN08324236_REFG_MMP08324236 | 6 | -0.4542713291 | Flavobacteriales | Chryseobacteriur | Chryseobacteriur | 3.19E-06 | 0 | 0.05097717955 | 1.13E-04 | 1 | 1 | 0 | 0.06843829366 | 0 | 0 | 0 |
| MARD_SAMN08324238_REFG_MMP08324238 | 2 | -0.4741893905 | Flavobacteriales | Chryseobacteriur | Chryseobacteriur | 0 | 0 | 0.06018155298 | 1.18E-04 | 0 | 0.002002573836 | 0 | 0.055938001 | 0 | 0 | 0 |
| MARD_SAMN08329936_REFG_MMP08329936 | 2 | -0.5554209457 | Enterobacterales | Vibrio | Vibrio plantispon: | 0 | 0 | 0.03047826813 | 1.54E-05 | 0.08213024533 | 4.05E-04 | 0 | 0.00135375186 | 0 | 7.17E-04 | 0 |
| MARD_SAMN08372522_REFG_MMP08372522 | 2 | -0.00325717776 | Desulfuromonad | PPFX01 | PPFX01 sp00289 | 7.92E-07 | 0 | 0.1843354106 | 2.61E-05 | 0.002078912705 | 6.74E-06 | 0 | 0.01563206058 | 0 | 7.97E-04 | 0.03644060191 |
| MARD_SAMN08378603_REFG_MMP08378603 | 3 | -0.02787290325 | Rhodobacterales | Roseovarius | Roseovarius con | 0 | 0 | 1 | 1.72E-04 | 0.003199325985 | 1 | 0 | 1 | 0 | 1 | 0 |
| MARD_SAMN08383634_REFG_MMP08383634 | 4 | -0.2915972542 | Pseudomonadale | Marinobacter | Marinobacter flav | 0 | 0 | 1.000000187 | 1.35E-04 | 1.000000187 | 0.003350503464 | 0 | 0.05652062613 | 0 | 4.46E-04 | 0.001144758467 |
| MARD_SAMN08457632_REFG_MMP08457632 | 2 | -0.2374081526 | Flavobacteriales | Cloacibacterium | Cloacibacterium | 0 | 0 | 0.2128724835 | 1.10E-04 | 0.003219720272 | 2.48E-08 | 0 | 0.0709943263 | 0 | 0 | 0 |
| MARD_SAMN08457750_REFG_MMP08457750 | 3 | -0.6051721003 | Enterobacterales | Aeromonas | Aeromonas janda | 0 | 0 | 0.006859645849 | 1.21E-05 | 0.003009962911 | 1 | 0 | 0.06424643521 | 0 | 1 | 0 |
| MARD_SAMN08463771_REFG_MMP08463771 | 2 | -0.2626798825 | Pseudomonadale | Microbulbifer | Microbulbifer pac | 0.002692876718 | 0 | 0.06008711876 | 1 | 0.1396207223 | 0.005339477303 | 0 | 0.0865039713 | 0 | 7.92E-04 | 0.9833333333 |
| MARD_SAMN08470889_REFG_MMP08470889 | 5 | -0.06917789438 | Cyanobacteriales | Crocosphaera | Crocosphaera sp | 0 | 0 | 0.0483044366 | 1.41E-04 | 0.1302657531 | 1 | 0 | 0.07031562307 | 0 | 0 | 0 |
| MARD_SAMN08568348_REFG_MMP08568348 | 2 | -0.6325279021 | Bacillales | Bacillus | Bacillus pumilus | 0 | 0 | 0.06096234805 | 5.18E-05 | 0.005993102453 | 0 | 0 | 0.05804772975 | 0 | 2.11E-05 | 0 |
| MARD_SAMN08634427_REFG_MMP08634427 | 4 | -0.6257770863 | Enterobacterales | Vibrio | Vibrio mediterran | 5.42E-08 | 0 | 1.000000213 | 2.11E-04 | 0.01666666667 | 0.01355147191 | 0 | 0.9861770195 | 0 | 0 | 0 |
| MARD_SAMN08683338_REFG_MMP08683338 | 5 | -0.6545942784 | Enterobacterales | Salinivibrio | Salinivibrio kushr | 0 | 0 | 0.03774209562 | 9.29E-07 | 0.00137038236 | 1 | 0 | 0.04166150709 | 0 | 0.01096166689 | 0 |
| MARD_SAMN08707727_REFG_MMP08707727 | 2 | -0.1813494787 | Burkholderiales | Nitrosomonas | Nitrosomonas su | 0 | 0 | 0.108797964 | 9.04E-05 | 0 | 8.08E-06 | 0 | 0.06640355223 | 0 | 0 | 0 |
| MARD_SAMN08707949_REFG_MMP08707949 | 1 | -0.5196459316 | Enterobacterales | Zobellella_D | Zobellella_D taiw | 0 | 0 | 0.9999999966 | 1.91E-06 | 0.9834945023 | 0.9669226633 | 0 | 0.9999999966 | 0 | 2.58E-05 | 9.74E-06 |
| MARD_SAMN08730605_REFG_MMP08730605 | 2 | -0.209668125 | Rhodobacterales | Mangrovicoccus | Mangrovicoccus | 3.28E-04 | 0 | 0.06357253633 | 0.0167897285 | 0.001146575494 | 5.58E-06 | 1.61E-04 | 0.2235323681 | 1.61E-04 | 8.16E-04 | 0.03806463308 |
| MARD_SAMN08730821_REFG_MMP08730821 | 5 | -0.05091647048 | Rhizobiales | Nitratireductor_D | Nitratireductor_D | 0 | 0 | 0.1595208364 | 1.59E-04 | 0.2608900666 | 1 | 0 | 0.06588594789 | 0 | 0.008477690908 | 0.05943858946 |
| MARD_SAMN08769611_REFG_MMP08769611 | 2 | -0.1215648061 | Desulfurobacteri | Thermovibrio | Thermovibrio gua | 0.2835215731 | 0 | 0.0455229715 | 1.25E-07 | 0.001043114472 | 0.03333332854 | 0 | 0.06310932938 | 0 | 2.88E-05 | 0 |
| MARD_SAMN08769778_REFG_MMP08769778 | 2 | -0.1563697417 | Caulobacterales | Maricaulis | Maricaulis maris | 0 | 0 | 0.1634656923 | 2.37E-06 | 0.1710779629 | 0.05470645991 | 0 | 0.09143695639 | 0 | 2.61E-05 | 0 |
| MARD_SAMN08769784_REFG_MMP08769784 | 2 | -0.253377343 | Pseudomonadale | Marinobacter | Marinobacter nau | 1.69E-08 | 0 | 0.5663110901 | 1.13E-06 | 0.05000000424 | 0 | 0 | 0.07630965598 | 0 | 0 | 0.001049827107 |
| MARD_SAMN08770175_REFG_MMP08770175 | 6 | -0.09075605643 | Flavobacteriales | Tenacibaculum | Tenacibaculum g | 5.15E-08 | 0 | 0.02204492048 | 2.17E-06 | 1 | 5.20E-04 | 0 | 0.05725960455 | 0 | 0 | 0.00164519933 |
| MARD_SAMN08770194_REFG_MMP08770194 | 5 | -0.3616638901 | Cytophagales | Marinoscillum | Marinoscillum fur | 0 | 0 | 0.03385501827 | 1.04E-04 | 0.04235770858 | 0.9833961997 | 0 | 0.07790354615 | 0 | 0.003096324633 | 0.03460121895 |
| MARD_SAMN08776247_REFG_MMP08776247 | 4 | -0.2586616201 | Pseudomonadale | Kushneria | Kushneria indalir | 0 | 0 | 1.000000262 | 1.77E-06 | 0.9878151586 | 0 | 0 | 0.04514629368 | 0 | 2.57E-05 | 0 |
| MARD_SAMN08776299_REFG_MMP08776299 | 5 | -0.2057814035 | Flavobacteriales | Marinirhabdus | Marinirhabdus ge | 0 | 0 | 0.05060568922 | 1.21E-04 | 0.1457138896 | 1 | 0 | 0.07045521953 | 0 | 0.003892625226 | 0.03836736294 |
| MARD_SAMN08776300_REFG_MMP08776300 | 6 | -0.2099298417 | Pseudomonadale | Kushneria | Kushneria marisf | 0 | 0 | 0.04235377418 | 8.49E-05 | 1.000000259 | 0 | 0 | 0.05994588082 | 0 | 2.59E-06 | 0 |
| MARD_SAMN08778569_REFG_MMP08778569 | 2 | -0.1242049255 | Flavobacteriales | Tenacibaculum | Tenacibaculum lu | 0 | 0 | 0.03002463928 | 9.60E-05 | 0 | 0 | 0 | 0.9999996982 | 0 | 0 | 0 |

| | | | | | | | | | | | | | | | | |
|---|---|---|---|---|---|---|---|---|---|---|---|---|---|---|---|---|
| MARD_SAMN08779097_REFG_MMP08779097 | 5 | -0.1474150723 | Flavobacteriales | Winogradskyella | Winogradskyella | 4.13E-07 | 0 | 0.1362197359 | 1.25E-04 | 0.01196069279 | 1.000000207 | 0 | 0.05372021476 | 0 | 7.84E-04 | 9.51E-06 |
| MARD_SAMN08783578_REFG_MMP08783578 | 5 | -0.09156987711 | Rhodobacterales | Paracoccus | Paracoccus sigai | 0 | 0 | 0.1278589935 | 9.62E-07 | 0.00129839183 | 0.9999999992 | 7.05E-06 | 0.06140794969 | 0 | 0 | 0.07005064267 |
| MARD_SAMN08794446_REFG_MMP08794446 | 6 | -0.5289250146 | Pseudomonadale | Pseudomonas_E | Pseudomonas_E | 6.40E-08 | 0 | 0.02583560333 | 6.57E-06 | 0.9685698292 | 0 | 0 | 0.0526319352 | 0 | 2.57E-05 | 0.3113429394 |
| MARD_SAMN08813454_REFG_MMP08813454 | 2 | 0.01147338305 | Flavobacteriales | Lutibacter | Lutibacter citreus | 0 | 0 | 0.06787940533 | 1 | 0 | 0.00534145755 | 0 | 0.04410970874 | 0 | 7.59E-04 | 0 |
| MARD_SAMN08828797_REFG_MMP08828797 | 2 | -0.2951727473 | Bacteriovoracale | Halobacteriovora | Halobacteriovora | 0 | 0 | 0.05123136574 | 1.36E-04 | 0.009747182437 | 3.86E-06 | 0 | 0.06613503014 | 0 | 6.20E-04 | 0 |
| MARD_SAMN08848607_REFG_MMP08848607 | 2 | -0.4850704401 | Enterobacterales | Oceanimonas | Oceanimonas ma | 0 | 0 | 0.03273373313 | 1.05E-04 | 0.3056920013 | 0 | 0 | 0.06791527063 | 0 | 7.90E-04 | 0 |
| MARD_SAMN08888560_REFG_MMP08888560 | 4 | -0.1847350137 | Sphingomonadal | Alteripontixantho | Alteripontixantho | 0 | 0 | 0.9999997031 | 8.60E-07 | 2.45E-04 | 0.01666678842 | 0 | 0.0369318104 | 0 | 1.46E-05 | 0.005383341992 |
| MARD_SAMN08906339_REFG_MMP08906339 | 1 | -0.2475412704 | Flavobacteriales | Muricauda | Muricauda chong | 2.33E-05 | 0 | 0.9999997164 | 4.99E-05 | 0.004068533974 | 0.983333053 | 0 | 0.09007520308 | 0 | 7.61E-04 | 0.001936874035 |
| MARD_SAMN08912472_REFG_MMP08912472 | 2 | -0.2148758815 | Rhodobacterales | Pseudooceanicol | Pseudooceanicol | 0 | 0 | 0.06227085133 | 2.39E-04 | 0.03109760245 | 2.38E-06 | 0.0203530115 | 0.05124204437 | 0 | 8.10E-04 | 0.35 |
| MARD_SAMN08913591_REFG_MMP08913591 | 5 | -0.5832801482 | Bacillales | Bacillus_BN | Bacillus_BN acar | 0 | 0 | 0.009949574111 | 3.37E-05 | 0.001097570215 | 1.000000228 | 0 | 0.148320324 | 0 | 0.01088924347 | 0 |
| MARD_SAMN08917673_REFG_MMP08917673 | 2 | -0.04331311133 | Kiloniellales | Algihabitans | Algihabitans albu | 0 | 0 | 0.05819857236 | 8.43E-05 | 0.006253228349 | 0.006385117582 | 0 | 0.04765108348 | 0 | 0 | 0 |
| MARD_SAMN08917691_REFG_MMP08917691 | 3 | -0.3569309235 | Enterobacterales | Alginatibacterium | Alginatibacterium | 0 | 0 | 0.1093036756 | 4.80E-05 | 0.00316752867 | 0.9999997692 | 0 | 0.02758927434 | 0 | 0.9999997692 | 0.03609189121 |
| MARD_SAMN08918410_REFG_MMP08918410 | 4 | -0.2131648535 | Rhodobacterales | Sulfitobacter | Sulfitobacter sp0 | 0 | 0 | 1.000000296 | 1.90E-05 | 0.003363248964 | 7.11E-06 | 0 | 0.06028779336 | 0 | 7.99E-04 | 0 |
| MARD_SAMN08918501_REFG_MMP08918501 | 1 | -0.1714023937 | Rhodobacterales | Pseudosulfitobac | Pseudosulfitobac | 0 | 0 | 1 | 1.32E-04 | 0.004431133612 | 1 | 0 | 0.04884074204 | 0 | 8.00E-04 | 0 |
| MARD_SAMN08918503_REFG_MMP08918503 | 2 | -0.1876579496 | Rhodobacterales | Pseudosulfitobac | Pseudosulfitobac | 0 | 0 | 0.02220559831 | 2.32E-06 | 0.01676338866 | 5.98E-05 | 0 | 0.07625413105 | 0 | 0 | 7.21E-04 |
| MARD_SAMN08924931_REFG_MMP08924931 | 8 | -0.09258875579 | Cytophagales | Pleomorphovibric | Pleomorphovibric | 0 | 0 | 0.9999998603 | 0.9999998603 | 0.2316565149 | 0.9999998603 | 0 | 0.06056933059 | 0 | 7.82E-04 | 0 |
| MARD_SAMN08934404_REFG_MMP08934404 | 2 | -0.07421780028 | Rhodobacterales | Roseovarius | Roseovarius sed | 0 | 0 | 0.03116083288 | 4.60E-05 | 0.007060675922 | 0.003254893467 | 0.0414635449 | 0.0693071755 | 0 | 0.001116613407 | 0 |
| MARD_SAMN08934405_REFG_MMP08934405 | 2 | -0.3121750937 | Rhodobacterales | Thalassorhabdor | Thalassorhabdor | 0 | 0 | 0.04271478086 | 1.83E-04 | 0.1390125786 | 3.83E-04 | 0 | 0.06903056123 | 5.79E-05 | 0 | 0 |
| MARD_SAMN08944605_REFG_MMP08944605 | 2 | -0.09674723707 | Lachnospirales | Vallitalea | Vallitalea guayma | 0 | 0 | 0.04202148799 | 7.64E-05 | 0.03333333533 | 0.001722877384 | 0 | 0.2709053282 | 0 | 0 | 0 |
| MARD_SAMN08954522_REFG_MMP08954522 | 2 | -0.1782310005 | Flavobacteriales | Maribacter | Maribacter litorali | 0 | 0 | 0.07022757896 | 1.04E-04 | 0 | 2.13E-06 | 0 | 0.2973859287 | 0 | 0 | 0 |
| MARD_SAMN08954523_REFG_MMP08954523 | 7 | -0.5798930017 | Enterobacterales | Zobellella_A | Zobellella_A mar | 0 | 0 | 0.02313047424 | 0.2840156825 | 0 | 0.005914827568 | 0 | 0.03721830442 | 0 | 1 | 0 |
| MARD_SAMN08969185_REFG_MMP08969185 | 2 | -0.03359028588 | Rhodobacterales | Meridianimarinicc | Meridianimarinicc | 0 | 0 | 0.03064732593 | 0.01671508909 | 0.02218938537 | 6.85E-06 | 6.50E-05 | 0.06349318201 | 0 | 8.18E-04 | 0 |
| MARD_SAMN08970161_REFG_MMP08970161 | 2 | -0.2074480721 | Rhodobacterales | Pararhodobacter | Pararhodobacter | 0 | 0 | 0.03123944688 | 1.78E-05 | 0.04558113236 | 0 | 0 | 0.06678789791 | 0 | 8.18E-04 | 5.00E-06 |
| MARD_SAMN08970498_REFG_MMP08970498 | 5 | -0.1791752801 | Rhodobacterales | Pararhodobacter | Pararhodobacter | 0 | 0 | 0.03899492814 | 0.001583206002 | 0.3874597821 | 0.9999996681 | 0 | 0.05601889822 | 0 | 8.75E-04 | 0.0487392121 |
| MARD_SAMN08979297_REFG_MMP08979297 | 2 | -0.04175596793 | Streptosporangia | Marinitenerispora | Marinitenerispora | 0 | 0 | 0.06470146679 | 6.25E-05 | 0.00112680377 | 0.07222273014 | 0 | 0.06410847261 | 0 | 0 | 0 |
| MARD_SAMN08997732_REFG_MMP08997732 | 6 | -0.2702666392 | Flavobacteriales | Winogradskyella | Winogradskyella | 0 | 2.33E-06 | 0.0262592632 | 0.05000249116 | 0.9519985269 | 4.31E-04 | 1.33E-04 | 0.9527813777 | 0 | 0.009255686443 | 0.03333334244 |
| MARD_SAMN08997828_REFG_MMP08997828 | 2 | -0.02275411593 | Balneolales | QGGB01 | QGGB01 sp0031 | 0 | 0 | 0.03584406113 | 1.17E-04 | 0.005061944297 | 8.66E-04 | 0 | 0.0461066214 | 0 | 7.77E-04 | 0.1386400075 |
| MARD_SAMN08998300_REFG_MMP08998300 | 1 | -0.1583218932 | Rhodobacterales | Salibaculum | Salibaculum grise | 5.58E-08 | 0.003310133459 | 0.9923679705 | 0.02262355447 | 0.999999708 | 0.9835198451 | 0.01666666667 | 0.9811547551 | 0 | 5.27E-05 | 0 |
| MARD_SAMN09010082_REFG_MMP09010082 | 2 | -0.1582505059 | Nevskiales | Abyssibacter | Abyssibacter pro | 0 | 0 | 0.04537210015 | 1.56E-04 | 0.009605109813 | 7.25E-04 | 0 | 0.04938502022 | 0 | 1.60E-04 | 0 |
| MARD_SAMN09041437_REFG_MMP09041437 | 5 | 0.06006200436 | Nitrococcales | E85 | E85 sp00314943 | 0.001212400593 | 0 | 0.04128900681 | 1.74E-04 | 0.007127420255 | 1.000000109 | 0 | 0.3441987741 | 0 | 0 | 0.5999999926 |
| MARD_SAMN09041732_REFG_MMP09041732 | 5 | -0.6449106996 | Bacillales | Bacillus | Bacillus subtilis | 0 | 0 | 0.02428054968 | 7.54E-07 | 0.003679221221 | 0.9833335351 | 0 | 0.06833665605 | 0 | 0.02796952053 | 0 |
| MARD_SAMN09064531_REFG_MMP09064531 | 4 | -0.352009936 | Caulobacterales | Litorimonas | Litorimonas taeai | 0 | 0 | 1.000000195 | 0.9833344381 | 0.005686512831 | 0.002687610337 | 0 | 1.000000195 | 0 | 0 | 0.008745779461 |
| MARD_SAMN09074685_REFG_MMP09074685 | 2 | -0.4251023412 | Pseudomonadale | Umboniibacter | Umboniibacter m | 0 | 0.0314955143 | 0.0507275611 | 5.25E-05 | 0.006392065646 | 2.85E-05 | 4.88E-05 | 0.1959370938 | 0 | 1.37E-05 | 0.5833334404 |
| MARD_SAMN09074692_REFG_MMP09074692 | 2 | -0.7808010338 | Bacillales | Cytobacillus | Cytobacillus ocea | 0 | 0 | 0.1073350483 | 1 | 0.1010521133 | 0.001789722706 | 0 | 0.06190802086 | 0 | 7.82E-04 | 0.003392910934 |
| MARD_SAMN09074742_REFG_MMP09074742 | 2 | -0.2239388991 | Rhodobacterales | Yoonia | Yoonia sediminili | 0 | 0 | 0.01778243811 | 0.01666667167 | 0.01740463958 | 0.08706335278 | 0 | 0.07143976691 | 0 | 7.55E-04 | 0.1389613511 |
| MARD_SAMN09074856_REFG_MMP09074856 | 2 | -0.0947017914 | DSM-26407 | Thioalbus | Thioalbus denitri | 1.21E-06 | 0 | 0.02934293924 | 2.46E-06 | 0.002392469014 | 0.01674599571 | 0 | 0.06595923076 | 0 | 0.01298262152 | 0 |
| MARD_SAMN09074857_REFG_MMP09074857 | 5 | -0.1773858143 | Arenicellales | Arenicella | Arenicella xantha | 3.21E-06 | 0 | 0.2178477408 | 8.66E-05 | 0.1148818101 | 0.983333596 | 0 | 0.07975789817 | 0 | 0.03062765236 | 0 |
| MARD_SAMN09074869_REFG_MMP09074869 | 5 | -0.09777939092 | Sphingomonadal | Parasphingopyxi | Parasphingopyxi | 0 | 0 | 0.1410576095 | 7.58E-05 | 0.002068021701 | 1 | 0 | 0.06302966172 | 0.01068160231 | 7.77E-04 | 0 |
| MARD_SAMN09074945_REFG_MMP09074945 | 2 | -0.2920837713 | Rhodobacterales | Pontivivens | Pontivivens insul | 0 | 0 | 0.1865584081 | 1.32E-04 | 0.09005452888 | 0 | 0 | 0.0757768447 | 0 | 0 | 0 |
| MARD_SAMN09074954_REFG_MMP09074954 | 5 | -0.3905866132 | Pseudomonadale | Marinomonas | Marinomonas aq | 0 | 0 | 0.05113027361 | 9.88E-05 | 0.01028512026 | 0.6833333333 | 0 | 0.07810678 | 0 | 6.38E-04 | 0 |
| MARD_SAMN09074955_REFG_MMP09074955 | 5 | -0.1970011908 | Pseudomonadale | Marinomonas | Marinomonas foli | 0 | 0 | 0.04587479801 | 5.98E-05 | 0.01585238256 | 1.000000091 | 0 | 0.06858550388 | 0 | 8.02E-04 | 0 |
| MARD_SAMN09074957_REFG_MMP09074957 | 5 | -0.2493591858 | Pseudomonadale | Marinomonas | Marinomonas rhi | 0 | 0 | 0.04256808517 | 1.16E-04 | 0.1173902582 | 0.8500000058 | 0 | 0.07949472995 | 0 | 7.99E-04 | 0 |
| MARD_SAMN09074958_REFG_MMP09074958 | 3 | -0.3279571779 | Pseudomonadale | Marinomonas | Marinomonas po | 0 | 0 | 0.01946399439 | 3.72E-05 | 0 | 0.9833333333 | 0 | 0.1276972788 | 0 | 0.9833595685 | 0 |
| MARD_SAMN09074965_REFG_MMP09074965 | 4 | -0.3975349172 | UBA8366 | Aestuariispira | Aestuariispira ins | 1.16E-05 | 0 | 1.000000178 | 8.63E-07 | 0.04720503712 | 4.54E-08 | 8.11E-05 | 0.03390048271 | 0 | 2.93E-05 | 0 |
| MARD_SAMN09078362_REFG_MMP09078362 | 6 | -0.1784843743 | Pseudomonadale | Marinospirillum | Marinospirillum p | 2.34E-06 | 0 | 0.03278898088 | 1.21E-04 | 1.000000293 | 1.000000293 | 0 | 0.04623850204 | 0 | 0.001616407531 | 0.03622988452 |
| MARD_SAMN09078458_REFG_MMP09078458 | 2 | -0.1417462388 | Cytophagales | Reichenbachiella | Reichenbachiella | 1.19E-06 | 0 | 0.02819003271 | 1.24E-04 | 0.00335267209 | 0.002240862051 | 0 | 0.08899652901 | 0 | 0 | 0 |
| MARD_SAMN09080319_REFG_MMP09080319 | 6 | -0.1966926378 | Rhodobacterales | Pararhodobacter | Pararhodobacter | 0 | 0 | 0.02059748617 | 2.96E-06 | 0.983333334 | 0 | 2.60E-04 | 0.1610416508 | 0 | 0 | 0.02465767162 |
| MARD_SAMN09083205_REFG_MMP09083205 | 8 | -0.06938306456 | Actinomycetales | Micrococcus | Micrococcus lute | 0 | 0 | 1 | 1 | 0 | 1 | 0 | 1 | 0 | 5.39E-04 | 0.07005413311 |
| MARD_SAMN09104480_REFG_MMP09104480 | 1 | -0.53292549 | Lactobacillales | Streptococcus | Streptococcus ini | 0.01158548203 | 0.003531112409 | 1.000000007 | 0.05013474691 | 0.05000000332 | 0.950492509 | 0 | 0.2761966666 | 0 | 1.32E-05 | 0.02972618508 |
| MARD_SAMN09195925_REFG_MMP09195925 | 5 | -0.1167030986 | Flavobacteriales | Brumimicrobium | Brumimicrobium | 4.89E-04 | 0 | 0.09952470506 | 9.81E-05 | 0.1039325643 | 1.000000201 | 0.01641544504 | 0.05506877971 | 0 | 2.70E-04 | 0 |
| MARD_SAMN09205056_REFG_MMP09205056 | 4 | -0.7074374823 | Enterobacterales | Vibrio | Vibrio albus | 0 | 0 | 0.9999998401 | 0.950000894 | 0.1411715915 | 0.03519340476 | 0 | 0.05814341162 | 0 | 0.03334720621 | 0.0657580017 |
| MARD_SAMN09205228_REFG_MMP09205228 | 1 | -0.2476178021 | Rhodobacterales | Litorivita | Litorivita pollutaq | 0 | 0 | 0.7076019865 | 3.02E-05 | 0.00907823538 | 1.000000003 | 0 | 0.05189279465 | 0 | 0.3166666667 | 0 |
| MARD_SAMN09206170_REFG_MMP09206170 | 2 | -0.4917293171 | Pseudomonadale | Acinetobacter | Acinetobacter rac | 0 | 0 | 0.007777550493 | 4.90E-05 | 0.145543215 | 4.82E-06 | 0 | 0.04720248872 | 0 | 0.001570605902 | 0 |
| MARD_SAMN09209463_REFG_MMP09209463 | 5 | -0.1591248989 | Rhodobacterales | Halovulum | Halovulum sedim | 0 | 0 | 0.02598034629 | 1.99E-06 | 6.40E-04 | 0.9500482117 | 0 | 0.04669921663 | 0 | 0.01118508886 | 0 |
| MARD_SAMN09214043_REFG_MMP09214043 | 2 | -0.2380912436 | Sphingomonadal | Sphingomonas | Sphingomonas p | 5.17E-06 | 0 | 0.04875786957 | 8.88E-05 | 0.03333333337 | 1.32E-04 | 0 | 0.06766306081 | 0 | 0.00157230205 | 0 |
| MARD_SAMN09217665_REFG_MMP09217665 | 2 | -0.1595721311 | Flavobacteriales | Muricauda | Muricauda koree | 0 | 0 | 0.03940309884 | 1.27E-04 | 0.003326637711 | 0 | 0 | 0.07381763937 | 0 | 7.90E-04 | 0 |
| MARD_SAMN09218204_REFG_MMP09218204 | 5 | -0.3618590516 | Pseudomonadale | Marinobacter_A | Marinobacter_A b | 0 | 0 | 0.04419513348 | 5.65E-07 | 0.02512133628 | 1 | 0.03396785003 | 0.9845111286 | 0 | 1.28E-05 | 0.001915320397 |
| MARD_SAMN09228017_REFG_MMP09228017 | 5 | -0.1942888549 | Rhodobacterales | Paracoccus | Paracoccus sedi | 0 | 0 | 0.03306889889 | 8.34E-05 | 0 | 1.000000168 | 0 | 0.1249437202 | 0 | 8.11E-04 | 0.02414077012 |
| MARD_SAMN09228018_REFG_MMP09228018 | 6 | -0.03058087755 | Rhodobacterales | Amaricoccus | Amaricoccus sp0 | 1.22E-07 | 0 | 0.0401215179 | 0.05000187143 | 0.95 | 1.24E-05 | 0 | 0.09843357854 | 0 | 5.52E-06 | 0 |
| MARD_SAMN09228019_REFG_MMP09228019 | 2 | -0.1608246811 | Flavobacteriales | Confluentibacter | Confluentibacter | 0 | 0 | 0.02636694525 | 1.14E-04 | 0.06337414424 | 0.1764707443 | 0 | 0.07492123462 | 0 | 7.98E-04 | 0 |
| MARD_SAMN09228020_REFG_MMP09228020 | 5 | -0.2839582365 | Pseudomonadale | Marinomonas | Marinomonas sh | 8.04E-07 | 0 | 0.0602552603 | 0.01666741843 | 0.01441819455 | 0.9999998209 | 8.12E-07 | 0.0302480828 | 1.82E-05 | 2.63E-05 | 1.03E-05 |
| MARD_SAMN09228104_REFG_MMP09228104 | 1 | -0.04089067344 | Rhodobacterales | Meridianimarinicc | Meridianimarinicc | 0 | 0 | 0.9999998491 | 5.37E-05 | 0.116536847 | 0.9833460056 | 0 | 0.03864798433 | 0.0230533668 | 0.01090316731 | 9.09E-04 |
| MARD_SAMN09228976_REFG_MMP09228976 | 5 | -0.2471061988 | Pseudomonadale | Marinimicrobium_ | Marinimicrobium_ | 0 | 1.30E-04 | 0.03106500622 | 6.37E-05 | 0.2604397427 | 1 | 0 | 0.08413332682 | 0 | 8.07E-04 | 0 |
| MARD_SAMN09230411_REFG_MMP09230411 | 2 | -0.3543685617 | Cytophagales | Algoriphagus | Algoriphagus litoi | 3.14E-08 | 0 | 0.04202750525 | 0.999999996 | 0.002313426145 | 0.004717742911 | 0 | 0.07934298377 | 0 | 8.10E-04 | 0.03665094808 |
| MARD_SAMN09236930_REFG_MMP09236930 | 2 | -0.126657542 | Rhizobiales | Aestuariivirga | Aestuariivirga lito | 0 | 0 | 0.04268731853 | 7.61E-05 | 0.003324335493 | 0 | 0 | 0.101052929 | 0 | 7.89E-04 | 0 |
| MARD_SAMN09240566_REFG_MMP09240566 | 4 | -0.4184520387 | Enterobacterales | Pseudoalteromor | Pseudoalteromor | 0 | 0 | 1 | 1.96E-04 | 0.009443331202 | 0 | 0 | 0.05879283956 | 0 | 7.52E-04 | 0 |
| MARD_SAMN09245781_REFG_MMP09245781 | 5 | -0.03291800167 | Mycobacteriales | Micromonospora | Micromonospora | 0 | 0 | 0.02631354518 | 0.01545645951 | 0.006449475657 | 1 | 0 | 1 | 0 | 0 | 0 |
| MARD_SAMN09245787_REFG_MMP09245787 | 2 | 0.005991356437 | Mycobacteriales | Micromonospora | Micromonospora | 0 | 0 | 0.1630549694 | 0.05000089229 | 0.0145101197 | 0.01065064302 | 2.10E-04 | 0.05449809287 | 7.21E-05 | 0.01673886862 | 0.01669215773 |
| MARD_SAMN09269770_REFG_MMP09269770 | 4 | -0.122996732 | Sphingomonadal | Alteriqipengyuan | Alteriqipengyuan | 1.04E-06 | 4.81E-05 | 1.000000229 | 0.01678573753 | 1.41E-04 | 0.03349178648 | 0 | 0.9980109119 | 0 | 0.05000000413 | 0.05797062092 |
| MARD_SAMN09269811_REFG_MMP09269811 | 1 | -0.07701812696 | Bradymonadales | Lujinxingia | Lujinxingia litorali | 0 | 0 | 0.9999997706 | 2.89E-04 | 0.1035903462 | 0.9999997706 | 0 | 0.01419382102 | 0 | 0.01087695882 | 0 |
| MARD_SAMN09279555_REFG_MMP09279555 | 4 | -0.1931991773 | Actinomycetales | OACT-916 | OACT-916 sp003 | 0 | 0 | 0.9717818787 | 6.78E-07 | 0.002830707934 | 4.00E-04 | 0 | 1 | 0 | 2.78E-05 | 0.03699464051 |
| MARD_SAMN09279561_REFG_MMP09279561 | 6 | -0.05583397626 | Rhodobacterales | Roseovarius | Roseovarius amc | 1.15E-06 | 0 | 0.0426966552 | 1.34E-04 | 1.000000179 | 0.004102654209 | 0 | 0.07900440298 | 0 | 0 | 5.60E-04 |
| MARD_SAMN09283747_REFG_MMP09283747 | 6 | -0.06174324738 | Mycoplasmatales | Metamycoplasma | Metamycoplasma | 0 | 0 | 0.05227263873 | 0.9999997674 | 0.9999997674 | 0.9988595378 | 0 | 0.06450990186 | 0 | 0 | 0.9999997674 |

| | | | | | | | | | | | | | | | | |
|---|---|---|---|---|---|---|---|---|---|---|---|---|---|---|---|---|
| MARD_SAMN09377736_REFG_MMP09377736 | 2 | -0.1723560163 | Flavobacteriales | Mesonia | Mesonia sp0032: | 0 | 0 | 0.08718098923 | 2.25E-06 | 0.01010013864 | 0.3839190293 | 0 | 0.05192611855 | 0 | 1.84E-05 | 0.00113129302 |
| MARD_SAMN09377739_REFG_MMP09377739 | 4 | -0.3905127453 | Pseudomonadale | Marinomonas | Marinomonas pie | 0 | 0 | 1 | 5.64E-05 | 0 | 3.04E-05 | 0 | 0.04961752108 | 0 | 0 | 0.02001839009 |
| MARD_SAMN09381170_REFG_MMP09381170 | 1 | -0.1168273949 | Francisellales | Facilibium | Facilibium subfla | 0 | 0 | 0.9999998024 | 0.000124103240 | 0.003361854023 | 0.9999998024 | 0 | 0.3721279141 | 0 | 0.006211090522 | 0 |
| MARD_SAMN09386782_REFG_MMP09386782 | 2 | -0.1582214097 | Flavobacteriales | Putridiphycobact | Putridiphycobact | 0 | 0 | 0.05513749341 | 4.88E-05 | 0.1081702878 | 5.86E-06 | 0 | 0.06592915026 | 0 | 0.00817093185 | 0.04323025239 |
| MARD_SAMN09389153_REFG_MMP09389153 | 2 | -0.2359367371 | Flavobacteriales | Winogradskyella | Winogradskyella | 3.30E-08 | 0 | 0.03227204587 | 0.9666677034 | 0.00108657301 | 0.06666667074 | 2.99E-04 | 0.07821750973 | 1.82E-04 | 3.97E-05 | 0 |
| MARD_SAMN09425313_REFG_MMP09425313 | 2 | -0.04150754825 | Rhodobacterales | Aquicoccus | Aquicoccus porpl | 0.005913430813 | 0 | 0.2717321219 | 5.52E-06 | 0.1116324123 | 0.03333342728 | 0 | 0.05190250709 | 0 | 3.91E-05 | 0 |
| MARD_SAMN09425405_REFG_MMP09425405 | 2 | -0.2772562113 | Pseudomonadale | Marinobacter | Marinobacter lito | 0 | 0 | 0.1029635373 | 1.86E-04 | 0.003288927629 | 0 | 4.64E-04 | 0.06589502249 | 0 | 9.58E-04 | 0 |
| MARD_SAMN09433089_REFG_MMP09433089 | 2 | -0.05770772342 | Desulfobacterale | Desulfobacter | Desulfobacter hy | 0 | 0 | 0.05521669812 | 4.18E-05 | 1.01E-04 | 0.003148488261 | 0 | 0.05465683974 | 0 | 3.61E-05 | 0 |
| MARD_SAMN09437379_REFG_MMP09437379 | 2 | -0.7725934091 | Enterobacterales | Vibrio | Vibrio tetraodonis | 0 | 0 | 0.1330021163 | 1.11E-05 | 0.006418444371 | 0 | 0 | 0.05963630608 | 0 | 7.95E-06 | 0 |
| MARD_SAMN09437435_REFG_MMP09437435 | 2 | -0.4272412178 | Enterobacterales | Vibrio | Vibrio maerlii | 0 | 0 | 0.1702191451 | 1.22E-04 | 0 | 1.78E-06 | 0 | 0.04347440576 | 0 | 0 | 0 |
| MARD_SAMN09437463_REFG_MMP09437463 | 2 | -0.579126314 | Enterobacterales | Vibrio | Vibrio rhodolitus | 0 | 0 | 0.03359573087 | 0 | 0.2301865534 | 4.52E-04 | 0 | 0.0391549662 | 0 | 0 | 0 |
| MARD_SAMN09469620_REFG_MMP09469620 | 3 | -0.333280064 | Enterobacterales | Algibacillus | Algibacillus agari | 0 | 0 | 0.05281992465 | 1.82E-04 | 0.08505167573 | 1 | 0 | 0.05496323345 | 0 | 1 | 0 |
| MARD_SAMN09478201_REFG_MMP09478201 | 2 | -0.02332142956 | Rhodobacterales | HLUCCA09 | HLUCCA09 sp00 | 0 | 0 | 0.02271781545 | 2.72E-06 | 0.08032749704 | 9.97E-08 | 0 | 0.05725217967 | 0 | 0.00164337833 | 0 |
| MARD_SAMN09478215_REFG_MMP09478215 | 7 | -0.1207273376 | Rhodobacterales | Tropicimonas | Tropicimonas sp( | 0 | 0 | 0.02058414048 | 1.02E-04 | 0.007356023294 | 5.61E-06 | 0 | 0.06470577778 | 0 | 1 | 0.03784300663 |
| MARD_SAMN09478228_REFG_MMP09478228 | 4 | -0.1280295922 | Rhizobiales | Pelagibacterium | Pelagibacterium | 0 | 0 | 1 | 3.72E-05 | 0.08708317841 | 0 | 0 | 0.05111182539 | 0 | 0 | 0 |
| MARD_SAMN09487630_REFG_MMP09487630 | 1 | -0.3460563157 | Enterobacterales | Aliidiomarina | Aliidiomarina cele | 0 | 0 | 0.9999998888 | 8.60E-05 | 1.79E-04 | 0.618005593 | 0.04321287802 | 0.3103650198 | 0 | 0.0181651876 | 0 |
| MARD_SAMN09499650_REFG_MMP09499650 | 2 | -0.2214045399 | Enterobacterales | Aliidiomarina | Aliidiomarina sp0 | 0 | 0 | 0.03859371802 | 0.002070467371 | 0.06926139914 | 1.05E-05 | 0 | 0.04886735094 | 0 | 0.001592397606 | 0 |
| MARD_SAMN09499816_REFG_MMP09499816 | 2 | -0.03776108546 | Mycobacteriales | Dietzia | Dietzia maris | 0 | 0 | 0.03206043399 | 1.38E-04 | 0.005834419009 | 0.002171493681 | 0 | 1 | 0 | 7.66E-04 | 0.01619274754 |
| MARD_SAMN09507747_REFG_MMP09507747 | 8 | -0.1582981406 | Rhodobacterales | Roseovarius | Roseovarius dice | 3.29E-04 | 0 | 0.134363875 | 1.000000207 | 0.007666284458 | 0.9833956136 | 0 | 0.9845700026 | 2.34E-05 | 7.77E-04 | 0.00273127919 |
| MARD_SAMN09508687_REFG_MMP09508687 | 5 | -0.09236623409 | Rhodobacterales | Roseovarius | Roseovarius sp0 | 0 | 0 | 0.02800945198 | 3.32E-05 | 0.003236064329 | 1 | 0 | 0.2492916382 | 0 | 8.01E-04 | 0 |
| MARD_SAMN09508719_REFG_MMP09508719 | 5 | -0.114413743 | Streptomycetales | Streptomyces | Streptomyces rer | 0 | 0 | 0.03685016186 | 1.88E-04 | 0.003177776477 | 1 | 0 | 0.1899865327 | 0 | 7.55E-04 | 0 |
| MARD_SAMN09508721_REFG_MMP09508721 | 2 | -0.1109890997 | Streptomycetales | Streptomyces | Streptomyces dia | 0 | 0 | 0.01202776576 | 5.89E-05 | 0.006742878556 | 0.005456346284 | 0 | 0.1380431158 | 0 | 8.03E-04 | 0 |
| MARD_SAMN09509732_REFG_MMP09509732 | 5 | -0.2259187868 | Pseudomonadale | Halomonas | Halomonas sulfic | 0 | 0 | 0.187067111 | 2.79E-05 | 0.03829151332 | 1.000000002 | 0 | 0.06030281984 | 0 | 5.37E-07 | 0 |
| MARD_SAMN09509804_REFG_MMP09509804 | 4 | -0.3124519767 | Pseudomonadale | Stutzerimonas | Stutzerimonas sp | 0 | 0 | 1 | 0.004461981371 | 0.04525337427 | 0.005072230899 | 0 | 0.05253182788 | 0 | 0 | 0 |
| MARD_SAMN09519150_REFG_MMP09519150 | 2 | -0.3739890727 | Flavobacteriales | Chryseobacteriur | Chryseobacteriur | 0 | 0 | 0.04644895799 | 7.80E-05 | 0 | 1.50E-06 | 0 | 0.09928510524 | 0 | 0 | 0 |
| MARD_SAMN09522130_REFG_MMP09522130 | 2 | -0.3061918491 | Enterobacterales | Psychromonas | Psychromonas sp | 0 | 0 | 0.149400304 | 2.00E-05 | 0.00342223702 | 0 | 0 | 0.05774681343 | 0 | 2.37E-05 | 0 |
| MARD_SAMN09522132_REFG_MMP09522132 | 2 | -0.4347452068 | Enterobacterales | Vibrio | Vibrio cortegader | 0 | 0 | 0.02969517223 | 6.30E-07 | 0.004932735671 | 0.001807903768 | 0 | 0.09943693292 | 0 | 1.31E-05 | 0 |
| MARD_SAMN09522133_REFG_MMP09522133 | 2 | -0.3872759505 | Rhodobacterales | OMPR01 | OMPR01 sp9003 | 0 | 0 | 0.03596404813 | 1.000000203 | 0.006530318276 | 0 | 0 | 0.05266274387 | 0 | 7.77E-04 | 0 |
| MARD_SAMN09522134_REFG_MMP09522134 | 2 | -0.3103470906 | Rhodobacterales | Ruegeria | Ruegeria sp0033 | 0 | 0 | 0.05923363395 | 1 | 0.002488677117 | 0 | 0 | 0.1300244045 | 0 | 7.77E-04 | 3.63E-06 |
| MARD_SAMN09522135_REFG_MMP09522135 | 4 | -0.3170500352 | Rhodobacterales | Yoonia | Yoonia sp003318 | 0 | 0 | 0.9837892045 | 8.24E-07 | 0.203966108 | 0.03528575546 | 0 | 0.9843834451 | 0 | 1.48E-04 | 0 |
| MARD_SAMN09522149_REFG_MMP09522149 | 2 | 0.01649641809 | Rhodobacterales | Rhodosalinus | Rhodosalinus se | 0 | 0 | 0.03283555081 | 3.85E-05 | 0.01632505341 | 0 | 0 | 0.06839854905 | 0 | 8.02E-04 | 0 |
| MARD_SAMN09522150_REFG_MMP09522150 | 6 | 0.04620723302 | Rhodobacterales | Rhodosalinus | Rhodosalinus ha | 0 | 0 | 0.02949678611 | 0.01666666234 | 0.9999998212 | 0.002156504515 | 0 | 0.08749840423 | 0 | 2.30E-04 | 0.02506426837 |
| MARD_SAMN09531221_REFG_MMP09531221 | 2 | -0.0822082983 | Chitinophagales | Lewinella_A | Lewinella_A sp00 | 0 | 0 | 0.04846388289 | 0.02112629928 | 0.003540585731 | 0 | 0 | 0.251775973 | 0 | 8.41E-04 | 1 |
| MARD_SAMN09531617_REFG_MMP09531617 | 4 | -0.4950578376 | Bacillales | Bacillus_D | Bacillus_D piezo | 0 | 0 | 0.9851537315 | 2.35E-07 | 0.01444258342 | 0.0171836965 | 0 | 1 | 2.47E-05 | 0.01666666667 | 0 |
| MARD_SAMN09579956_REFG_MMP09579956 | 2 | -0.1030090144 | Rhodobacterales | Rhodovulum | Rhodovulum visa | 0 | 0 | 0.02250893302 | 1.19E-04 | 0.004699643545 | 0.004285251076 | 0.07851160121 | 0.04399796577 | 0 | 7.65E-04 | 0 |
| MARD_SAMN09580333_REFG_MMP09580333 | 5 | -0.4125731265 | Rhizobiales | Cohaesibacter | Cohaesibacter in | 0 | 0 | 0.03121458177 | 0.01666667273 | 0.03223936625 | 1.000000006 | 0 | 0.05126256494 | 0 | 3.36E-05 | 8.49E-08 |
| MARD_SAMN09580375_REFG_MMP09580375 | 5 | -0.04103859629 | Rhodobacterales | Jannaschia | Jannaschia form | 0 | 0 | 0.04740410719 | 9.54E-05 | 0.003988707076 | 0.8333336524 | 0 | 0.08048197992 | 0 | 0 | 0.1500000205 |
| MARD_SAMN09625553_REFG_MMP09625553 | 2 | -0.3627198119 | Bacillales_G | Maribacillus | Maribacillus taea | 0 | 0 | 0.1110305629 | 1.10E-04 | 0.3528834964 | 7.90E-06 | 0.01205599821 | 0.07409800299 | 0.001349088589 | 0.02967831469 | 0 |
| MARD_SAMN09637267_REFG_MMP09637267 | 2 | -0.2309676938 | Pirellulales | Bremerella | Bremerella marin | 0 | 0 | 0.02864635062 | 7.51E-05 | 3.67E-06 | 6.53E-06 | 0 | 0.04304294465 | 0 | 0.008868980634 | 0 |
| MARD_SAMN09637346_REFG_MMP09637346 | 5 | -0.2317731142 | Pirellulales | Bremerella | Bremerella marin | 0 | 0 | 0.01280094086 | 1.12E-06 | 0.1321824119 | 0.9999997333 | 0 | 0.04237519632 | 0 | 3.70E-04 | 0 |
| MARD_SAMN09637541_REFG_MMP09637541 | 2 | -0.1481517837 | Pirellulales | Bremerella | Bremerella creme | 0 | 0 | 0.02721392528 | 9.03E-05 | 0.06671648858 | 0.002803489375 | 0 | 0.07049340377 | 0 | 0 | 0.06739420466 |
| MARD_SAMN09665230_REFG_MMP09665230 | 4 | -0.462061397 | Pseudomonadale | Salinibius | Salinibius halmop | 2.26E-06 | 0 | 1 | 1.21E-04 | 1 | 0 | 0 | 0.06388679779 | 3.11E-05 | 0 | 0 |
| MARD_SAMN09667312_REFG_MMP09667312 | 3 | -0.1035110338 | Flavobacteriales | Winogradskyella | Winogradskyella | 0 | 0 | 0.04044748485 | 1 | 0.2218336516 | 1 | 0 | 1 | 0 | 1 | 1 |
| MARD_SAMN09700386_REFG_MMP09700386 | 2 | -0.5333638622 | Enterobacterales | Shewanella | Shewanella algid | 0 | 0 | 0.05693774296 | 7.86E-05 | 0 | 0 | 0 | 0.06108344441 | 0 | 0.009238231662 | 0 |
| MARD_SAMN09700398_REFG_MMP09700398 | 4 | -0.3790239319 | Enterobacterales | Shewanella | Shewanella japo | 0 | 0 | 0.9999996161 | 1.55E-04 | 0.235328694 | 1.45E-07 | 0.002114205462 | 0.09653800474 | 0 | 0 | 0.9166662828 |
| MARD_SAMN09708543_REFG_MMP09708543 | 5 | -0.5835231617 | Bacillales | Planococcus | Planococcus sali | 0 | 0 | 0.1072734899 | 1.12E-04 | 0.0835271753 | 0.9833768256 | 0.0043767148 | 0.0662652997 | 0 | 7.69E-04 | 0 |
| MARD_SAMN09709391_REFG_MMP09709391 | 2 | -0.3950532615 | Pseudomonadale | Motiliproteus | Motiliproteus cor | 0 | 0 | 0.05793039021 | 1.14E-04 | 0.003282561797 | 0.2000011416 | 0 | 0.09002126961 | 0 | 0 | 0 |
| MARD_SAMN09710960_REFG_MMP09710960 | 5 | -0.7288547051 | Enterobacterales | Klebsiella | Klebsiella pneum | 0.1071428302 | 0 | 0.08750982303 | 2.74E-05 | 0.00636158357 | 1 | 0 | 0.2020283732 | 0 | 4.37E-05 | 0 |
| MARD_SAMN09713909_REFG_MMP09713909 | 5 | -0.03729005849 | Bradymonadales | Lujinxingia | Lujinxingia sedim | 0 | 0 | 0.1915831794 | 1.43E-04 | 0.008031420324 | 1.00000019 | 3.80E-06 | 1.00000019 | 0 | 0 | 0 |
| MARD_SAMN09714579_REFG_MMP09714579 | 5 | -0.1441698692 | Spirochaetales_E | Oceanispirochae | Oceanispirochae | 0 | 0 | 0.123036291 | 3.25E-05 | 0.0182297224 | 1.000000155 | 0 | 0.065581128 | 0 | 7.64E-04 | 0 |
| MARD_SAMN09721247_REFG_MMP09721247 | 6 | -0.2699803445 | Rhodobacterales | Paracoccus | Paracoccus sp00 | 4.76E-08 | 0 | 0.1283684759 | 1.08E-06 | 0.98333667 | 3.87E-04 | 8.84E-05 | 0.01877974693 | 0 | 1.29E-05 | 0.0323364162 |
| MARD_SAMN09725931_REFG_MMP09725931 | 2 | -0.2325188868 | Flavobacteriales | QXCW01 | QXCW01 sp0035 | 0 | 0 | 0.1752811016 | 9.08E-05 | 0.004487488083 | 0.05779615145 | 0 | 0.06063105641 | 0 | 0 | 0 |
| MARD_SAMN09736004_REFG_MMP09736004 | 2 | -0.4563674259 | Enterobacterales | Enterobacter | Enterobacter hor | 0 | 0 | 0.1174960609 | 1.27E-04 | 0.001637714789 | 0.09091046125 | 0 | 0.1207140773 | 0 | 0 | 0 |
| MARD_SAMN09736005_REFG_MMP09736005 | 2 | -0.4719596396 | Enterobacterales | Citrobacter | Citrobacter freun | 0.003477343007 | 0 | 0.01789697511 | 1.26E-05 | 0.07114727682 | 0 | 0.004928306016 | 0.07099415807 | 0 | 6.07E-05 | 0.03704699911 |
| MARD_SAMN09759695_REFG_MMP09759695 | 5 | -0.1951748181 | Rhodobacterales | Paroceanicella | Paroceanicella sp | 0 | 0 | 0.05313239192 | 1.14E-04 | 0.02753088216 | 1 | 0 | 0.06685914995 | 0 | 0 | 0 |
| MARD_SAMN09759696_REFG_MMP09759696 | 4 | -0.1768886023 | Sphingobacterial | Albibacterium | Albibacterium inc | 0 | 0 | 0.9999999969 | 1.21E-04 | 0.9999999969 | 8.23E-05 | 0 | 0.988723615 | 0 | 7.74E-04 | 0 |
| MARD_SAMN09762457_REFG_MMP09762457 | 2 | -0.1185310655 | Flavobacteriales | Brumimicrobium | Brumimicrobium | 0 | 0 | 0.1343731138 | 1.66E-05 | 0 | 1.45E-06 | 0 | 0.2944032025 | 0 | 0.01172000261 | 0 |
| MARD_SAMN09762562_REFG_MMP09762562 | 5 | -0.6320150095 | Enterobacterales | Thalassotalea | Thalassotalea eu | 0 | 0 | 0.1069893029 | 7.03E-05 | 0 | 0.9999998387 | 0 | 0.0556903512 | 0 | 8.00E-04 | 0 |
| MARD_SAMN09762563_REFG_MMP09762563 | 2 | -0.7428992266 | Enterobacterales | Thalassotalea | Thalassotalea eu | 0 | 0 | 0.1495671803 | 7.81E-05 | 0 | 2.79E-04 | 0 | 0.3138772855 | 0 | 0.01389655943 | 0 |
| MARD_SAMN09762564_REFG_MMP09762564 | 2 | -0.5820434313 | Enterobacterales | Thalassotalea | Thalassotalea eu | 1.55E-06 | 0 | 0.01549646693 | 0.03333487856 | 0.006762955561 | 7.84E-05 | 0 | 0.04940248007 | 0 | 0.01667319721 | 0.005963144751 |
| MARD_SAMN09762594_REFG_MMP09762594 | 6 | -0.5011233881 | Enterobacterales | Alteromonas | Alteromonas aes | 3.51E-06 | 0 | 0.05007192391 | 4.25E-05 | 1 | 6.25E-06 | 0.03703700812 | 0.06919177823 | 0.003499190978 | 3.04E-05 | 0 |
| MARD_SAMN09767239_REFG_MMP09767239 | 2 | -0.1399013647 | Flavobacteriales | Muricauda | Muricauda nanha | 0 | 0 | 0.04716541268 | 1.18E-04 | 0 | 4.87E-06 | 0 | 0.0373295517 | 0 | 0 | 0 |
| MARD_SAMN09767465_REFG_MMP09767465 | 5 | -0.9736402813 | Enterobacterales | Serratia | Serratia marcesc | 0 | 0 | 0.9999997087 | 2.44E-06 | 6.51E-04 | 0.9999997087 | 1.69E-04 | 0.06664067163 | 0.9500123324 | 4.82E-04 | 0 |
| MARD_SAMN09770279_REFG_MMP09770279 | 2 | -0.1087530361 | Sphingomonadal | Tsuneonella | Tsuneonella troits | 0 | 0 | 0.05268716073 | 0.3156479969 | 0 | 7.70E-06 | 0 | 0.05515014536 | 0 | 0 | 0 |
| MARD_SAMN09786826_REFG_MMP09786826 | 4 | -4.95E-04 | Mycobacteriales | Micromonospora | Micromonospora | 0 | 0 | 1.00000001 | 0.95000698 | 0.96751915 | 0.007433147279 | 0 | 0.05778917032 | 0 | 0.001564492874 | 0 |
| MARD_SAMN09835256_REFG_MMP09835256 | 2 | -0.151849488 | Flavobacteriales | Marixanthomona | Marixanthomona | 0 | 0 | 0.1281637432 | 7.77E-05 | 0 | 0.07000152986 | 0 | 0.04109931341 | 0 | 7.14E-07 | 0 |
| MARD_SAMN09835476_REFG_MMP09835476 | 6 | -0.5732624174 | Enterobacterales | Vibrio | Vibrio sinensis | 0 | 0 | 0.04399401472 | 9.46E-05 | 1 | 1 | 0 | 0.05940409974 | 0 | 0 | 0 |
| MARD_SAMN09842172_REFG_MMP09842172 | 6 | -0.3814528034 | Sphingobacterial | Mucilaginibacter | Mucilaginibacter | 3.17E-06 | 0 | 0.03647083212 | 1.03E-04 | 1 | 0.0027093768 | 0 | 0.07162257996 | 0 | 0 | 0 |
| MARD_SAMN09845332_REFG_MMP09845332 | 2 | -0.2332432304 | Flavobacteriales | Arenibacter | Arenibacter sp00 | 0 | 0 | 0.217190248 | 1.28E-04 | 0.002453834262 | 0.001744733862 | 0 | 0.113657187 | 0 | 0 | 0 |
| MARD_SAMN09901140_REFG_MMP09901140 | 2 | -0.3616861661 | Bacillales_D | Oceanobacillus | Oceanobacillus p | 0.004710407181 | 0 | 0.2774113961 | 0.05016833545 | 0.09252500003 | 1.62E-04 | 2.46E-04 | 0.08960762073 | 0 | 0 | 2.06E-04 |
| MARD_SAMN09901207_REFG_MMP09901207 | 8 | -0.07758841142 | Bacteroidales | Mariniphaga | Mariniphaga sedi | 0.001307846296 | 0 | 1 | 0.9666675356 | 0.005926893923 | 0.9667276614 | 0.01666666667 | 0.09681922942 | 0 | 4.54E-05 | 0.002340648504 |
| MARD_SAMN09908144_REFG_MMP09908144 | 2 | 0.07154676002 | Rhodobacterales | IMCC34051 | IMCC34051 sp00 | 0 | 0 | 0.1616668293 | 1.83E-05 | 0.003536726693 | 3.44E-07 | 0 | 0.05055095184 | 0 | 0 | 0 |
| MARD_SAMN09914392_REFG_MMP09914392 | 1 | -0.6038960447 | Enterobacterales | Motilimonas | Motilimonas pum | 2.99E-06 | 0 | 1 | 1.31E-04 | 0.002470862053 | 1 | 0 | 0.002915479454 | 0 | 0.004948197545 | 0 |

| | | | | | | | | | | | | | | | | |
|---|---|---|---|---|---|---|---|---|---|---|---|---|---|---|---|---|
| MARD_SAMN09915279_REFG_MMP09915279 | 4 | -0.1539562611 | Caulobacterales | Henriciella | Henriciella barba | 0 | 0 | 1.000000004 | 3.36E-06 | 0.001697241013 | 0.017567158 | 0 | 0.05626272904 | 0 | 2.51E-05 | 0 |
| MARD_SAMN09915280_REFG_MMP09915280 | 2 | -0.164626816 | Caulobacterales | Henriciella | Henriciella algico | 0 | 0 | 0.005806296383 | 9.52E-05 | 0.00169923185 | 7.05E-06 | 0 | 0.05919049699 | 0 | 8.06E-04 | 0 |
| MARD_SAMN09915281_REFG_MMP09915281 | 2 | -0.1502165873 | Caulobacterales | Henriciella | Henriciella mobili | 0 | 0 | 0.02923242492 | 1.28E-06 | 4.21E-04 | 0.002444772158 | 0 | 0.05659512472 | 0 | 1.34E-05 | 0 |
| MARD_SAMN09916314_REFG_MMP09916314 | 2 | -0.3973703496 | Enterobacterales | Litorilituus | Litorilituus sp003 | 0 | 0 | 0.2373902272 | 1.34E-04 | 0.003919184196 | 0.00211687747 | 0.004475662744 | 0.0684667383 | 0.001224871065 | 0 | 0.9999998507 |
| MARD_SAMN09920664_REFG_MMP09920664 | 2 | -0.5675447105 | Bacillales | Bacillus_A | Bacillus_A bomby | 7.51E-08 | 0.002126994564 | 0.03850664207 | 2.28E-06 | 0.01737984823 | 0 | 0 | 0.03868317019 | 0 | 0.004226170483 | 0 |
| MARD_SAMN09930780_REFG_MMP09930780 | 5 | -0.1541476533 | Sphingomonadal | Alteraurantiaciba | Alteraurantiaciba | 0 | 0 | 0.05319056945 | 1.78E-04 | 0.009453566068 | 0.9999999811 | 0 | 0.2685434301 | 0 | 0 | 0.01667215616 |
| MARD_SAMN09932323_REFG_MMP09932323 | 2 | -0.7634516834 | Bacillales | Bacillus | Bacillus lichenifor | 0 | 0 | 0.04549244007 | 0.106561095 | 0.003957769375 | 0.01666902055 | 0.02862660826 | 0.968597059 | 0 | 0.004187663038 | 0.004272274432 |
| MARD_SAMN09932619_REFG_MMP09932619 | 5 | -0.07598103059 | Actinomycetales | Dermacoccus | Dermacoccus ab | 0 | 0 | 0.04471386004 | 8.98E-05 | 0.007099398089 | 0.9833330506 | 0 | 0.07834115014 | 0 | 0 | 0 |
| MARD_SAMN09939629_REFG_MMP09939629 | 6 | -0.1968548009 | Sphingomonadal | Tsuneonella | Tsuneonella supr | 1.46E-07 | 0 | 0.05026546925 | 3.44E-06 | 0.9999999952 | 9.67E-05 | 0 | 0.05561162066 | 0 | 1.08E-07 | 0 |
| MARD_SAMN09939640_REFG_MMP09939640 | 6 | -0.3032609899 | Flavobacteriales | Muricauda | Muricauda aequ | 5.26E-06 | 0 | 0.02679042083 | 9.89E-05 | 1 | 3.16E-06 | 0 | 0.05531633235 | 0 | 0 | 0 |
| MARD_SAMN09939641_REFG_MMP09939641 | 6 | -0.3310177131 | Flavobacteriales | Muricauda | Muricauda mariti | 2.92E-06 | 0 | 0.03340945911 | 6.89E-05 | 1 | 0.002759340306 | 0 | 0.05089910793 | 0 | 5.84E-06 | 0 |
| MARD_SAMN09939643_REFG_MMP09939643 | 2 | -0.2805094058 | Flavobacteriales | Muricauda | Muricauda lutima | 0 | 0 | 0.03425743017 | 4.10E-05 | 0.001738604703 | 7.38E-06 | 0 | 0.05270462559 | 0 | 8.26E-04 | 0.03890231786 |
| MARD_SAMN09939799_REFG_MMP09939799 | 1 | -0.2411203362 | Rhodobacterales | Gemmobacter | Gemmobacter lu | 0 | 0 | 1 | 3.65E-05 | 0.08265297076 | 1 | 0.06608959703 | 0.0587141911 | 0.001878121045 | 7.94E-04 | 0.06860542994 |
| MARD_SAMN09989090_REFG_MMP09989090 | 4 | -0.1080549906 | Rhodobacterales | Paracoccus | Paracoccus aest | 0 | 0 | 0.9999996424 | 9.58E-05 | 0.0033443655920 | .001622117443 | 0 | 0.03853673277 | 0 | 7.94E-04 | 0 |
| MARD_SAMN10058827_REFG_MMP10058827 | 2 | -0.2604539561 | Pseudomonadale | Alcanivorax | Alcanivorax profu | 0 | 0 | 0.03728762769 | 3.63E-05 | 0.01364549946 | 0 | 0 | 0.04811006592 | 0 | 5.47E-04 | 0 |
| MARD_SAMN10075489_REFG_MMP10075489 | 1 | -0.3057033044 | Enterobacterales | Pseudoalteromor | Pseudoalteromor | 0 | 0 | 0.9999997674 | 1.01E-04 | 0.003435234596 | 0.9999997674 | 0 | 0.06190863885 | 0 | 8.14E-04 | 0.03805547862 |
| MARD_SAMN10076538_REFG_MMP10076538 | 1 | -0.08177547719 | Rhodobacterales | Chachezhania | Chachezhania ar | 0 | 0 | 1 | 9.44E-05 | 0 | 1 | 0 | 0.06398831735 | 0 | 0.009011607341 | 0 |
| MARD_SAMN10081109_REFG_MMP10081109 | 4 | -0.3590435512 | Enterobacterales | Parashewanella | Parashewanella | 0 | 0 | 1 | 4.45E-05 | 0 | 0.3833333333 | 0 | 0.05197618753 | 0 | 3.02E-04 | 0 |
| MARD_SAMN10081840_REFG_MMP10081840 | 8 | -0.5234514597 | Enterobacterales | Parashewanella | Parashewanella | 0 | 0 | 0.04159499982 | 1 | 0.003130220758 | 1 | 0 | 0.07140911849 | 0 | 0.001834047236 | 0 |
| MARD_SAMN10130764_REFG_MMP10130764 | 2 | -0.02045281819 | Rhodobacterales | Roseovarius | Roseovarius spo | 2.26E-06 | 0 | 0.4546094869 | 9.60E-05 | 0.1684136847 | 6.19E-05 | 0 | 0.02138796479 | 0 | 0.003065641172 | 0 |
| MARD_SAMN10130826_REFG_MMP10130826 | 5 | -0.2026581308 | Sphingomonadal | Altericroceibacte | Altericroceibacte | 0 | 0 | 0.04416914322 | 1.80E-04 | 0.00304953879 | 0.9999997385 | 0 | 0.01759212497 | 0 | 0.009977170305 | 0 |
| MARD_SAMN10130974_REFG_MMP10130974 | 2 | -0.1036521184 | Sphingomonadal | Alteraurantiaciba | Alteraurantiaciba | 2.25E-06 | 0 | 0.03523212002 | 6.11E-05 | 0.04071687205 | 4.97E-06 | 0 | 0.9844022513 | 0 | 0 | 0 |
| MARD_SAMN10134656_REFG_MMP10134656 | 2 | -0.1423357594 | Flavobacteriales | Aquimarina | Aquimarina later | 2.52E-06 | 0 | 0.03246548545 | 7.01E-05 | 0.2708466837 | 4.62E-06 | 0 | 0.0821115101 | 0 | 0 | 0 |
| MARD_SAMN10134657_REFG_MMP10134657 | 6 | -0.1546272866 | Flavobacteriales | Aquimarina | Aquimarina aggr | 0 | 0 | 0.06646161061 | 8.71E-05 | 1.000000239 | 5.11E-05 | 0 | 0.05159853237 | 0 | 5.12E-08 | 6.66E-04 |
| MARD_SAMN10134658_REFG_MMP10134658 | 2 | -0.09193927003 | Flavobacteriales | AU392 | AU392 sp003443 | 5.06E-08 | 0 | 0.03165877124 | 2.22E-06 | 0.03627363668 | 0 | 0 | 0.06339568637 | 0 | 1.10E-07 | 0.004928732715 |
| MARD_SAMN10134659_REFG_MMP10134659 | 5 | -0.1038077739 | Flavobacteriales | Aquimarina | Aquimarina sp00 | 0 | 0 | 0.03581322041 | 6.79E-08 | 0.08675442873 | 0.9834161697 | 0.01666666667 | 0.07357207135 | 0 | 0.0152487844 | 2.85E-05 |
| MARD_SAMN10134660_REFG_MMP10134660 | 2 | -0.4433341234 | Enterobacterales | Alteromonas | Alteromonas sp0 | 0 | 0 | 0.02942975035 | 4.04E-05 | 0.1299342591 | 0 | 0 | 0.05767188729 | 0 | 0.004774060635 | 0 |
| MARD_SAMN10141419_REFG_MMP10141419 | 5 | -0.02610278257 | Streptomycetales | Streptomyces | Streptomyces ho | 0 | 0 | 0.05336883305 | 3.17E-05 | 0.1404405626 | 1.000000006 | 0 | 0.07212375611 | 0 | 8.68E-04 | 0.09125607226 |
| MARD_SAMN10141426_REFG_MMP10141426 | 2 | -0.2036188476 | Streptomycetales | Streptomyces | Streptomyces kle | 0.00447090534 | 0 | 0.02947822839 | 0.03818453484 | 0.07680887639 | 5.13E-06 | 0 | 0.04485624447 | 0 | 3.31E-08 | 0 |
| MARD_SAMN10146989_REFG_MMP10146989 | 5 | -0.1078126168 | Flavobacteriales | Ulvibacterium | Ulvibacterium ma | 0.1111111878 | 0 | 0.0347000649 | 8.97E-05 | 0.01544513152 | 1.000000138 | 0 | 0.04310821435 | 0 | 7.67E-04 | 0.1516261175 |
| MARD_SAMN10169376_REFG_MMP10169376 | 2 | -0.09872265014 | Pseudomonadale | Haliea | Haliea alexandrii | 0 | 0 | 0.1385675546 | 1.03E-04 | 0 | 4.60E-05 | 0 | 1 | 0 | 0 | 0 |
| MARD_SAMN10172232_REFG_MMP10172232 | 2 | -0.384729383 | Bacillales_D | Oceanobacillus | Oceanobacillus p | 0 | 0 | 0.05275009448 | 1.48E-04 | 0.003319522457 | 6.45E-05 | 0 | 0.10632852 | 0 | 0 | 0 |
| MARD_SAMN10172241_REFG_MMP10172241 | 7 | -0.322603561 | Bacillales_D | Oceanobacillus | Oceanobacillus b | 0 | 0.001966711235 | 0.04420438476 | 3.72E-05 | 0 | 0 | 0 | 0.1630601904 | 0 | 1.000000219 | 0.06188333721 |
| MARD_SAMN10225537_REFG_MMP10225537 | 4 | -0.1857122788 | Actinomycetales | Rhodoglobus | Rhodoglobus am | 2.40E-07 | 0 | 0.9999998464 | 9.88E-05 | 0.1963045017 | 0 | 0 | 0.08476196823 | 0 | 5.13E-08 | 0 |
| MARD_SAMN10230918_REFG_MMP10230918 | 2 | -0.1072217247 | Flavobacteriales | Mesonia | Mesonia aquimar | 0 | 6.40E-06 | 0.1165886825 | 0.05000683781 | 0.005101136724 | 0.1000059479 | 0.001039640941 | 0.9547290677 | 5.55E-06 | 7.73E-04 | 0.03403534776 |
| MARD_SAMN10231401_REFG_MMP10231401 | 5 | -0.05953599725 | Rhodobacterales | Rhodophyticola | Rhodophyticola p | 0 | 0 | 0.0574306183 | 0.06026632287 | 0 | 1.000000411 | 0 | 1.000000411 | 0 | 0 | 0 |
| MARD_SAMN10231957_REFG_MMP10231957 | 4 | -0.00752203096 | Mycobacteriales | Micromonospora | Micromonospora | 0 | 0 | 1 | 0.03756273235 | 0.004828133773 | 0.003629757332 | 0 | 0.0677404235 | 0 | 7.87E-04 | 0 |
| MARD_SAMN10231959_REFG_MMP10231959 | 5 | -0.02612466529 | Mycobacteriales | Micromonospora | Micromonospora | 0 | 0 | 0.1342249613 | 2.44E-04 | 0.009334720674 | 0.9999997493 | 0 | 0.05424716004 | 0 | 2.90E-04 | 0 |
| MARD_SAMN10250232_REFG_MMP10250232 | 2 | -0.1912341397 | Flavobacteriales | Dokdonia | Dokdonia sinensi | 0 | 0 | 0.02629130812 | 0.01684062353 | 0.2577239233 | 5.94E-06 | 0 | 0.9840508993 | 0 | 9.15E-04 | 0 |
| MARD_SAMN10250233_REFG_MMP10250233 | 5 | -0.08736773524 | Propionibacterial | Arachnia | Arachnia antarcti | 0 | 0 | 0.03597859006 | 5.30E-05 | 0.05562550352 | 1.000000189 | 0 | 1.000000189 | 0 | 7.96E-04 | 0 |
| MARD_SAMN10250535_REFG_MMP10250535 | 4 | -0.04456713593 | Rhodobacterales | Oceanicola | Oceanicola lentu | 0.2611073431 | 0 | 1.000000162 | 0.03339711162 | 0.9846392762 | 0.001017701862 | 0 | 0.07740766039 | 0 | 5.95E-06 | 0 |
| MARD_SAMN10261828_SAGS_MMP10261828 | 5 | -0.04855218208 | Flavobacteriales | PRS1 | PRS1 sp0037091 | 0 | 0 | 0.04487910821 | 1.36E-05 | 0.3089166763 | 1.000000007 | 0 | 0.06356902948 | 0 | 7.84E-04 | 0 |
| MARD_SAMN10276034_REFG_MMP10276034 | 5 | -0.1491740893 | Streptomycetales | Streptomyces | Streptomyces sp | 0 | 0 | 0.02382653316 | 4.54E-05 | 0.02099704665 | 0.9833333333 | 0 | 0.07068385458 | 0 | 0 | 0 |
| MARD_SAMN10284718_REFG_MMP10284718 | 2 | -0.3614336131 | Pseudomonadale | Marinomonas | Marinomonas hw | 0 | 0 | 0.019411022 | 7.29E-07 | 1.08E-05 | 0 | 0 | 0.1963106273 | 0 | 0.01519307448 | 0 |
| MARD_SAMN10319551_REFG_MMP10319551 | 5 | -0.2473554208 | Pseudomonadale | Marortus | Marortus luteolus | 0 | 0.1131098249 | 0.03607515912 | 4.28E-05 | 0.1515935416 | 0.6500000366 | 0 | 0.06415409009 | 0 | 2.73E-04 | 0 |
| MARD_SAMN10320226_REFG_MMP10320226 | 5 | -0.2313184036 | Sphingomonadal | Tsuneonella | Tsuneonella flava | 0 | 0 | 0.06511105052 | 1.10E-04 | 0.003255674981 | 1 | 0 | 0.3561682306 | 0 | 0 | 0 |
| MARD_SAMN10362856_REFG_MMP10362856 | 3 | -0.5559166634 | Rhodobacterales | Pacificibacter | Pacificibacter ma | 0 | 0 | 1 | 1.87E-04 | 0.004881205001 | 0.9833333333 | 9.51E-04 | 0.09067720087 | 0 | 1 | 0 |
| MARD_SAMN10362944_REFG_MMP10362944 | 2 | -0.2825478328 | Pseudomonadale | Sinobacterium | Sinobacterium ca | 0 | 0 | 0.02545836769 | 0.1723967173 | 0 | 4.06E-06 | 0 | 0.05367175311 | 0 | 0.002352702411 | 0 |
| MARD_SAMN10362946_REFG_MMP10362946 | 5 | -0.2126323387 | Nautiliales | Cetia | Cetia pacifica | 0 | 0 | 0.03338891915 | 1.87E-06 | 5.49E-05 | 0.9833335034 | 0 | 0.3279169648 | 0 | 0.005079206381 | 0 |
| MARD_SAMN10363320_REFG_MMP10363320 | 4 | -0.5314463808 | Enterobacterales | Gallaecimonas | Gallaecimonas p | 0 | 0 | 1 | 0 | 0.1189380706 | 0.01821641731 | 0 | 0.03377043445 | 0 | 1.26E-05 | 0 |
| MARD_SAMN10363429_REFG_MMP10363429 | 2 | -0.2901886247 | Pseudomonadale | Marinimicrobium | Marinimicrobium | 0 | 0 | 0.3740911508 | 2.48E-04 | 0.002599223175 | 0 | 0 | 0.06350293198 | 0 | 0 | 0 |
| MARD_SAMN10363435_REFG_MMP10363435 | 7 | -0.5925363745 | Staphylococcales | Salinicoccus | Salinicoccus rose | 1.30E-06 | 0 | 1 | 8.87E-05 | 0.003665748925 | 0.004734816983 | 0 | 0.04558182676 | 0 | 1 | 1 |
| MARD_SAMN10397511_REFG_MMP10397511 | 2 | -0.3137584494 | Enterobacterales | Shewanella | Shewanella psyc | 0 | 0 | 0.02654906179 | 8.14E-05 | 0 | 0 | 0 | 0.03257947799 | 0 | 0 | 0 |
| MARD_SAMN10397593_REFG_MMP10397593 | 2 | -0.3505161669 | Enterobacterales | Shewanella | Shewanella vesic | 0 | 0 | 0.04893621887 | 1.27E-04 | 0.003246820521 | 0 | 0 | 0.1935132044 | 0 | 0 | 0 |
| MARD_SAMN10397596_REFG_MMP10397596 | 6 | -0.2963834759 | Enterobacterales | Shewanella | Shewanella frigid | 2.81E-06 | 0 | 0.02077402353 | 0.01683024417 | 0.9842503596 | 1.65E-06 | 0 | 0.06653119554 | 0 | 0.001199266263 | 0 |
| MARD_SAMN10411098_REFG_MMP10411098 | 5 | -0.00993847263 | Flavobacteriales | Aureibaculum | Aureibaculum ma | 0 | 0 | 0.03828568994 | 4.03E-05 | 0.3318329251 | 0.9666668122 | 0 | 0.1014673977 | 0 | 0 | 0.06001067321 |
| MARD_SAMN10412359_REFG_MMP10412359 | 1 | -0.4076204527 | Pseudomonadale | Pseudomonas_E | Pseudomonas_E | 0 | 0 | 1 | 1.23E-04 | 1 | 1 | 0 | 0.2883627525 | 0 | 0 | 0 |
| TARA_SAMEA2591057_METAG_LKPMBOAD | 1 | -0.07010863873 | Opitutales | MB11C04 | MB11C04 sp002 | 0 | 0 | 1.000000006 | 0.01679733888 | 0.01666666986 | 0.9833333365 | 0 | 0.9864190744 | 0 | 0.0172424104 | 0 |
| TARA_SAMEA2591057_METAG_MGBAEOCH | 8 | -0.07826026464 | Flavobacteriales | UBA11891 | UBA11891 sp003 | 0 | 0 | 0.999999882 | 0.999999882 | 0.999999882 | 0.999999882 | 0 | 0.999999882 | 0 | 0.999999882 | 0 |
| TARA_SAMEA2591057_METAG_MGIKKEGI | 6 | -0.3023782939 | UBA1280 | UBA2136 | UBA2136 sp0026 | 0 | 0 | 0.07863228632 | 0.01666835587 | 0.999999996 | 1.26E-04 | 7.09E-04 | 0.2986677241 | 2.41E-08 | 0 | 0.006727314426 |
| TARA_SAMEA2591057_METAG_PEOJEGGA | 2 | -0.1582200506 | Flavobacteriales | UBA952 | UBA952 sp00216 | 0 | 0 | 0.002722133761 | 1.81E-06 | 5.21E-05 | 0.01666666667 | 0 | 0.001198812852 | 0 | 3.66E-04 | 0 |
| TARA_SAMEA2591074_METAG_MMNGHOGL | 5 | -0.2338992329 | Rhodobacterales | Amylibacter | Amylibacter sp90 | 0 | 0 | 0.3699181302 | 0.01068446099 | 0.09848866256 | 0.9999998564 | 3.31E-04 | 0.1022955383 | 0 | 9.56E-04 | 0.008109314385 |
| TARA_SAMEA2591084_METAG_PFJBCBHJ | 6 | -0.05299944019 | Puniceispirillales | AAA536-G10 | AAA536-G10 sp0 | 0.2114912285 | 0 | 0.139205475 | 0 | 1 | 1 | 0 | 0.08875472758 | 0 | 4.68E-05 | 0.05082162528 |
| TARA_SAMEA2591098_METAG_EJOBFJID | 4 | -0.1101808682 | PCC-6307 | Synechococcus_ | Synechococcus_ | 1.39E-07 | 0 | 0.9999999901 | 5.09E-06 | 0.001135040475 | 0.04755570914 | 0 | 0.03503112485 | 1.87E-05 | 3.09E-04 | 0 |
| TARA_SAMEA2591098_METAG_EKCBDPJI | 5 | -0.1370994431 | Rhizobiales | Afipia | Afipia broomeae | 0 | 0.009647697315 | 0.03190121129 | 1.15E-04 | 0.1178622054 | 1 | 0 | 0.07329004253 | 0 | 7.62E-04 | 0 |
| TARA_SAMEA2591098_METAG_ELHAFCCE | 3 | -0.0789548772 | Pseudomonadale | UBA9145 | UBA9145 sp0024 | 0 | 0 | 0.2248821533 | 2.47E-04 | 0.05869217957 | 0.9834055454 | 0.06883784164 | 0.02805919792 | 0.01153601375 | 0.9999998663 | 0 |
| TARA_SAMEA2591098_METAG_OOHHFBGP | 7 | -0.1218043461 | Rhodobacterales | LGRT01 | LGRT01 sp01677 | 0 | 0 | 0.05118997245 | 0.003337940163 | 0.07367601923 | 0 | 0 | 0.06453583379 | 0 | 1 | 0 |
| TARA_SAMEA2591122_METAG_BAJAKJIO | 4 | 0.04579334054 | SAR86 | TMED112 | TMED112 sp003 | 0 | 1.48E-04 | 0.9838173012 | 0.01666667077 | 0.1777883361 | 0.01896194974 | 0 | 0.999999757 | 0 | 0 | 0 |
| TARA_SAMEA2591122_METAG_BNKOMGJI | 4 | 0.005018709072 | Flavobacteriales | Marisimplicoccus | Marisimplicoccus | 0 | 0 | 1 | 1.85E-04 | 1.01E-04 | 0.03795618066 | 0 | 0.2525785696 | 0 | 0.003727468187 | 0.05862831525 |
| TARA_SAMEA2591122_METAG_DCEMABAA | 4 | -0.02151445738 | Flavobacteriales | GCA-2692065 | GCA-2692065 sp | 9.77E-08 | 0 | 0.9836907665 | 5.86E-07 | 0.9889153706 | 0.006649751958 | 0 | 0.03865207963 | 0 | 0.007108882673 | 0.001522244182 |
| TARA_SAMEA2591122_METAG_EDCMBBCG | 5 | -0.07386376198 | TMED127 | GCA-002690875 | GCA-002690875 | 0 | 0 | 0.06820427911 | 1.43E-04 | 0.1157260446 | 0.9833335973 | 0 | 0.466141581 | 0 | 0.01666666667 | 0 |
| TARA_SAMEA2591122_METAG_OKANJJLB | 1 | -0.00116940211 | SAR86 | AG-339-G14 | AG-339-G14 sp0 | 0 | 0 | 1.000000175 | 1.03E-04 | 0.05701724929 | 1.000000175 | 0 | 0.06670553813 | 0 | 0.01341819588 | 0 |
| TARA_SAMEA2619376_METAG_EJDLPCAL | 3 | 0.008652603655 | Flavobacteriales | MAG-121220-bin | MAG-121220-bin | 0 | 0 | 0.008042856669 | 9.07E-07 | 0.9833333333 | 0.9999999956 | 0 | 0.9999999956 | 0 | 0.9999999956 | 0 |

| | | | | | | | | | | | | | | | | |
|---|---|---|---|---|---|---|---|---|---|---|---|---|---|---|---|---|
| TARA_SAMEA2619376_METAG_MCHHHBNN | 6 | -0.01334621962 | TMED25 | TMED25 | TMED25 sp0026 | 0 | 0 | 0.03745487109 | 9.31E-05 | 0.9999997538 | 0.009694143995 | 0 | 0.06810571965 | 0 | 0.002883719762 | 0.9999997538 |
| TARA_SAMEA2619399_METAG_OEMCHJKJ | 2 | -0.00581871237 | Flavobacteriales | BACL11 | BACL11 sp00273 | 0 | 0 | 0.04810359287 | 1.17E-04 | 0.127903588 | 5.18E-06 | 0 | 0.02945641651 | 0 | 0 | 0 |
| TARA_SAMEA2619531_METAG_CDJDEOGJ | 1 | -0.02755975358 | GCA-002705445 | GCA-2705445 | GCA-2705445 sp | 0 | 0 | 1 | 9.94E-05 | 0.00328994144 | 1 | 0 | 0.06325219437 | 0 | 0 | 0 |
| TARA_SAMEA2619531_METAG_GFEDGDLC | 4 | -0.07882865565 | Flavobacteriales | MS024-2A | | 0 | 0 | 0.999999722 | 1.43E-04 | 0.999999722 | 2.53E-04 | 0 | 0 | 4.42E-04 | 0.001235617187 | 0 |
| TARA_SAMEA2619531_METAG_HGKJGEEB | 5 | -0.1665912034 | Flavobacteriales | UBA11663 | UBA11663 sp002 | 0 | 0 | 0.05531135729 | 1.29E-04 | 0.02252175369 | 1 | 0.02982591646 | 0.06849104196 | 0.001427560616 | 2.45E-04 | 0.04051938793 |
| TARA_SAMEA2619531_METAG_KPGAFOOA | 2 | -0.00982288472 | Flavobacteriales | TMED96 | | 0 | 0 | 0.05632848242 | 6.75E-05 | 9.70E-04 | 0.01667264741 | 0 | 0.07360446746 | 0 | 7.77E-04 | 0 |
| TARA_SAMEA2619531_METAG_LEPIDGAG | 4 | 0.1011454066 | Kiritimatiellales | UBA1859 | UBA1859 sp002 | 0 | 0 | 1 | 0.03333486728 | 0.1688334639 | 4.79E-04 | 0 | 0.9688217397 | 6.06E-06 | 2.05E-04 | 0.04308387814 |
| TARA_SAMEA2619531_METAG_LLOIFEGM | 5 | -0.01120220033 | TMED127 | GCA-2691145 | GCA-2691145 sp | 1.10E-04 | 1.87E-04 | 0.02661582383 | 2.67E-06 | 0.1321343694 | 0.9538859641 | 6.28E-04 | 0.08126446044 | 0.06485701537 | 7.80E-05 | 0.03333340614 |
| TARA_SAMEA2619531_METAG_MBJLLBJC | 5 | -0.2304600702 | Puniceispirillales | UBA8309 | UBA8309 sp002 | 0 | 0 | 0.02728652507 | 1.47E-04 | 0.5472210316 | 1.000000281 | 0 | 1.000000281 | 0 | 0.02001296284 | 0.003359447084 |
| TARA_SAMEA2619548_METAG_ANCJELAM | 3 | 0.04946398665 | SCGC-AAA003-L | TMED6 | TMED6 sp00216 | 0 | 0 | 1.000000241 | 2.99E-04 | 0.003646772904 | 1.000000241 | 0 | 0.6439756811 | 0 | 1.000000241 | 0 |
| TARA_SAMEA2619548_METAG_JHDLHBGO | 7 | -0.06637863093 | Cytophagales | UBA4465 | UBA4465 sp002 | 0 | 0 | 1.000000236 | 3.64E-05 | 0 | 0.001465958946 | 0 | 0.2653272382 | 0 | 1.000000236 | 0 |
| TARA_SAMEA2619548_METAG_PAIGEGGM | 5 | -0.04581910896 | Actinomarinales | Actinomarina | Actinomarina sp0 | 0 | 0 | 0.02296278446 | 1.18E-04 | 0.09343709351 | 0.9666669175 | 0 | 0.05968982648 | 0 | 8.49E-04 | 0 |
| TARA_SAMEA2619625_METAG_IHHGDKNJ | 3 | -0.2332803973 | Parvibaculales | NORP139 | | 0 | 0 | 0.01187397302 | 9.03E-05 | 0.00707830194 | 1 | 0 | 0.04611693938 | 0 | 1 | 0.06516039964 |
| TARA_SAMEA2619667_METAG_BPBCELAF | 8 | -0.00939926274 | MED-G09 | MED-G09 | MED-G09 sp002 | 0 | 0 | 1 | 1 | 0.9833465666 | 1 | 0 | 0.9450943564 | 0 | 0.003665236416 | 0 |
| TARA_SAMEA2619677_METAG_DHGDAHEG | 2 | -0.1794344093 | Flavobacteriales | Marinirhabdus | Marinirhabdus sp | 0 | 0 | 0.04090172015 | 1.53E-04 | 0.007297327162 | 0.00449401261 | 0 | 0.06555346955 | 0 | 7.82E-04 | 0 |
| TARA_SAMEA2619677_METAG_DMIJGCPA | 2 | -0.07468074299 | Rhodobacterales | Alterinioella | Alterinioella nitra | 0 | 0 | 0.02641394097 | 0.01666666667 | 0.05618820708 | 0.4015613236 | 0 | 0.05205576052 | 0 | 0.006680094457 | 0.1990877094 |
| TARA_SAMEA2619678_METAG_AMMAMHCO | 1 | -0.02521957498 | SCGC-AAA003-L | GCA-2689995 | GCA-2689995 sp | 0 | 0 | 1.000000302 | 3.37E-04 | 0.06521209744 | 1.000000302 | 0 | 0.02134689228 | 0 | 0 | 0 |
| TARA_SAMEA2619678_METAG_EHMDLEPA | 2 | -0.06616304706 | Marinisomatales | TMED108 | TMED108 sp913 | 0 | 0 | 0.05469232865 | 5.45E-04 | 0.01438304828 | 0 | 0 | 1.000000302 | 0 | 0 | 0 |
| TARA_SAMEA2619766_METAG_CECBDPKB | 2 | -0.09920340152 | Flavobacteriales | UBA7446 | UBA7446 sp002 | 0 | 0.05775780838 | 0.02132653704 | 3.75E-05 | 0.009680409376 | 0 | 0 | 0.0476525284 | 0 | 0.001538388228 | 0 |
| TARA_SAMEA2619766_METAG_DFICPMHO | 4 | -0.04466030566 | Thalassobaculale | TMED26 | TMED26 sp0032 | 0 | 0 | 0.9841621843 | 8.53E-07 | 0.09068543463 | 2.91E-05 | 0 | 0.05373558671 | 0 | 0.015521251 | 0.05501526296 |
| TARA_SAMEA2619766_METAG_KBDCKIJJ | 6 | -0.1529896067 | Puniceispirillales | UBA5951 | UBA5951 sp016 | 0 | 0 | 0.05292372676 | 1.08E-06 | 0.9833468927 | 0.9833330711 | 0 | 0.02073632091 | 0 | 0.00245995788 | 0.03475469603 |
| TARA_SAMEA2619766_METAG_LMODONND | 5 | -0.01447084788 | TMED25 | TMED25 | TMED25 sp0027 | 0 | 0 | 0.0254228799 | 9.74E-06 | 0.1466834641 | 1 | 0 | 0.04003480985 | 0.006257286835 | 0.004484794689 | 0 |
| TARA_SAMEA2619766_METAG_NIKCOOOI | 4 | -0.0961020641 | PCC-6307 | RCC307 | RCC307 sp0024 | 0 | 0 | 1 | 7.53E-05 | 0.05861643886 | 0.02309947112 | 0 | 0.06952246422 | 0 | 4.42E-05 | 0 |
| TARA_SAMEA2619766_METAG_PGHMOJPG | 1 | -0.06346391598 | Opitutales | Coraliomargarita | Coraliomargarita | 0 | 0 | 1.000000313 | 1.62E-06 | 0.4838283932 | 1.000000313 | 0 | 0.05512998611 | 0 | 4.54E-05 | 0 |
| TARA_SAMEA2619782_METAG_DDMJHDOJ | 8 | -0.01700028331 | Burkholderiales | UBA7377 | UBA7377 sp002 | 0 | 3.28E-06 | 0.2183127292 | 0.9666688073 | 0.1902805028 | 0.9999999954 | 0 | 0.08471832017 | 9.64E-05 | 7.93E-04 | 9.92E-05 |
| TARA_SAMEA2619782_METAG_DJKHELNO | 4 | -0.03233226562 | Rhodospirillales_ | TMED8 | TMED8 sp00269 | 0 | 0 | 1 | 6.83E-05 | 0.003412650344 | 6.42E-06 | 0 | 1 | 0 | 8.10E-04 | 0.03629978291 |
| TARA_SAMEA2619782_METAG_PAKDINJI | 2 | 9.94E-04 | Marinisomatales | TMED108 | TMED108 sp002 | 0 | 0 | 0.161930204 | 1.11E-04 | 0.002668103898 | 6.24E-06 | 0 | 0.09753978132 | 0 | 7.47E-04 | 0 |
| TARA_SAMEA2619791_METAG_DBIFCBNB | 5 | -0.3654286488 | Rhodobacterales | Celeribacter | Celeribacter bae | 0.1081767883 | 0.007200699466 | 0.06801947434 | 1.43E-04 | 0.05623175739 | 0.9999998878 | 0 | 0.1285363647 | 0 | 0.0124947241 | 0 |
| TARA_SAMEA2619802_METAG_BHKIALJD | 5 | -0.07217363642 | Nanopelagicales | S36-B12 | S36-B12 sp0027 | 7.58E-08 | 0 | 0.0324043853 | 0.01666666667 | 0.09351572425 | 0.9843147289 | 0.006826722628 | 0.0461006708 | 0 | 2.36E-05 | 7.59E-06 |
| TARA_SAMEA2619815_METAG_CFAFOEAL | 5 | -0.1711032988 | Caulobacterales | Maricaulis | | 0 | 0 | 0.02767105766 | 0.01667092237 | 0.006651534855 | 1.000000257 | 2.10E-04 | 0.1361202692 | 0 | 5.29E-04 | 6.86E-06 |
| TARA_SAMEA2619815_METAG_EELKOFIF | 5 | -0.1904949854 | Sphingomonadal | Sphingobium | Sphingobium xer | 0 | 0 | 0.02885757675 | 5.72E-05 | 0.003243750188 | 1 | 0 | 0.03568248123 | 0 | 0.005722348996 | 0 |
| TARA_SAMEA2619815_METAG_KJEPBPMI | 4 | -0.2581811201 | Pseudomonadale | Alcanivorax | Alcanivorax gela | 0 | 2.56E-06 | 0.9999998576 | 4.98E-05 | 0.00959652477 | 0.01046083282 | 0 | 0.9999998576 | 0 | 7.85E-04 | 0.04035478493 |
| TARA_SAMEA2619815_METAG_MBLFNBKD | 7 | -0.05944809052 | Rhodospirillales_ | UBA4479 | | 0 | 0 | 0.9999998527 | 2.76E-05 | 4.96E-05 | 0.004021985445 | 0 | 0.2942677308 | 2.10E-06 | 0.9999998527 | 5.33E-04 |
| TARA_SAMEA2619815_METAG_MJBMGOIA | 5 | -0.2317141321 | Enterobacterales | Idiomarina | Idiomarina loihie | 0 | 0 | 0.04497418307 | 6.08E-05 | 0.01022774194 | 0.9999997875 | 0 | 0.06462722444 | 0 | 0.0126860722 | 0 |
| TARA_SAMEA2619815_METAG_OFMCCBEM | 6 | -0.1283431077 | Phycisphaerales | UBA1924 | | 0 | 0 | 0.100468868 | 4.81E-05 | 1.000000227 | 1.000000227 | 0 | 0.08076319519 | 0 | 0.01010487939 | 0 |
| TARA_SAMEA2619818_METAG_NFEOAGHJ | 3 | 0.02112062308 | UBA1144 | GCA-002715585 | GCA-002715585 | 0 | 0 | 0.01092410207 | 0.01666666153 | 0.1301722292 | 0.9999997123 | 0 | 0.9833330508 | 0 | 0.9999997123 | 0 |
| TARA_SAMEA2619840_METAG_EDOKFFDD | 2 | -0.1148713511 | Longimicrobiales | UBA2589 | UBA2589 sp002 | 1.70E-06 | 0 | 0.02164467592 | 1.47E-05 | 0.1151346774 | 0 | 0 | 0.03726278168 | 0 | 0.03522201737 | 1.39E-04 |
| TARA_SAMEA2619840_METAG_GHMKJPCJ | 4 | -0.02475792538 | Marinisomatales | GCA-002707905 | GCA-002707905 | 0.2000898742 | 0 | 0.9999999969 | 0.0166666636 | 0.9999999969 | 8.58E-08 | 0 | 0.04969563101 | 0 | 3.35E-05 | 0.02824437762 |
| TARA_SAMEA2619840_METAG_HFJDGBKI | 2 | -0.02248940472 | UBA2968 | UBA2968 | UBA2968 sp002 | 0 | 0.01422422596 | 0.02896270458 | 1.39E-04 | 0.1150363321 | 6.60E-06 | 0 | 0.07442522302 | 1.01E-06 | 0.009805612963 | 0.06524442809 |
| TARA_SAMEA2619840_METAG_KHCHLJMN | 2 | -0.01986040806 | SAR202 | UBA2962 | UBA2962 sp002 | 0 | 0 | 0.03358276188 | 0.01666666667 | 0.01125487554 | 0.01666666667 | 0 | 0.05957635625 | 0 | 2.63E-05 | 0 |
| TARA_SAMEA2619840_METAG_OBGEGELO | 2 | -0.01922760015 | UBA6615 | UBA8079 | UBA8079 sp002 | 0 | 0 | 0.02886570625 | 0 | 0.01764555058 | 0.01666667067 | 0 | 0.05138908255 | 0 | 0 | 0.07679025889 |
| TARA_SAMEA2619854_METAG_DAMFOOEB | 2 | -0.3877655202 | Pseudomonadale | Pseudomonas_E | Pseudomonas_E | 0 | 0 | 0.01323924489 | 0.04167324816 | 0.02874532784 | 7.35E-05 | 0 | 0.01210412391 | 0 | 1.20E-05 | 0 |
| TARA_SAMEA2619857_METAG_LKAHBEPF | 8 | -0.04271339275 | SAR324 | UBA1014 | UBA1014 sp001 | 0 | 0 | 0.9999999958 | 0.9999999958 | 0.04173393301 | 0.9999999958 | 0 | 0.04745613617 | 0.9833333333 | 0.9999999958 | 0 |
| TARA_SAMEA2619879_METAG_PFBFHBIF | 5 | 0.0162265654 | Flavobacteriales | GCA-002728855 | GCA-002728855 | 0 | 0.002646260585 | 0.03032175597 | 7.68E-05 | 0.009211865754 | 1.000000227 | 0 | 0.2599237135 | 0 | 0.006257045859 | 0 |
| TARA_SAMEA2619903_METAG_AMEKCAIK | 3 | -0.1556051834 | Cytophagales | Roseivirga | | 0 | 0 | 0.05141136 | 0.9999997352 | 0.1150979489 | 0.9999997352 | 0 | 0.04402662467 | 5.30E-07 | 0.9999997352 | 0.03286583294 |
| TARA_SAMEA2619907_METAG_EHDDCGIN | 3 | 0.005692248679 | GCA-002705445 | GCA-2716945 | | 0 | 0 | 0.03300742211 | 0.01681981396 | 0.003575611888 | 1 | 0 | 0.04832460753 | 0 | 1 | 0.01666666667 |
| TARA_SAMEA2619907_METAG_EMGKELAO | 1 | -0.1055131113 | Pseudomonadale | UBA9145 | UBA9145 sp002 | 0 | 0 | 0.9999996751 | 2.34E-05 | 0.9999996751 | 0.9999996751 | 0 | 0.9999996751 | 0 | 0.005477921171 | 1.30E-06 |
| TARA_SAMEA2619907_METAG_FFJLAFKD | 2 | 0.0395841694 | GCA-002720895 | | | 0 | 0 | 0.02745057381 | 0 | 0.004642584535 | 0.005888770919 | 0 | 0.01768329218 | 0 | 7.36E-04 | 0 |
| TARA_SAMEA2619907_METAG_FNHKDGPB | 6 | -0.1322970473 | Sphingomonadal | Alteriqipengyuan | Alteriqipengyuan | 0.2426846531 | 0 | 0.02288007604 | 7.32E-05 | 0.9999997967 | 0.004744957999 | 0 | 0.07068511101 | 0 | 7.76E-04 | 0 |
| TARA_SAMEA2619907_METAG_HJDFOLOM | 8 | -0.0446983035 | Anaerolineales | UBA11657 | UBA11657 sp002 | 0 | 0 | 0.9672381766 | 0.9833337843 | 0.008104735099 | 0.9500002816 | 1.11E-04 | 0.9834100491 | 0.001386951573 | 1.000000002 | 0 |
| TARA_SAMEA2619907_METAG_MFMGCAAC | 5 | -0.08533401353 | UBA6615 | UBA6615 | UBA6615 sp002 | 0 | 0 | 0.003631112049 | 0.01679415177 | 0.02573205845 | 0.9999998118 | 0.2423039477 | 0.04016990859 | 0 | 0.01751459066 | 0 |
| TARA_SAMEA2619907_METAG_NOGEJPAG | 1 | -0.1105482803 | UBA11872 | UBA11872 | UBA11872 sp002 | 0 | 0.003343525722 | 0.9999999959 | 0.0166683199 | 0.03957863335 | 0.9999999959 | 0 | 0.9843118532 | 0 | 0.002492530016 | 0 |
| TARA_SAMEA2619950_METAG_ADLKCPBP | 2 | -0.08730084685 | Flavobacteriales | UBA5081 | UBA5081 sp002 | 0 | 0 | 0.02576748808 | 1.05E-04 | 0.0298381258 | 0.006712395688 | 0 | 0.04220795313 | 0 | 0 | 0 |
| TARA_SAMEA2619952_METAG_NJJBGEID | 2 | -0.00580290195 | Burkholderiales | BACL14 | | 0 | 3.54E-04 | 0.03417378646 | 1.53E-06 | 0.05237586534 | 1.24E-07 | 9.76E-04 | 0.06849653678 | 0 | 0.003151021835 | 0.006676675298 |
| TARA_SAMEA2619970_METAG_CLAEKLCO | 5 | -0.03249997792 | Pseudomonadale | UBA2168 | UBA2168 sp002 | 2.63E-04 | 0 | 0.04165456069 | 0.03333337261 | 0.3417570345 | 0.9833337815 | 0 | 0.9834664047 | 0 | 3.99E-05 | 0 |
| TARA_SAMEA2619970_METAG_FKFPFEBP | 5 | -0.03566723608 | CAJXCL01 | CAJXCL01 | | 4.38E-06 | 0 | 0.02166721021 | 0 | 0.2712499345 | 0.9999998436 | 0 | 0.04093439101 | 0 | 0.003417351568 | 0 |
| TARA_SAMEA2619970_METAG_FKMCDENO | 5 | -0.00928920050 | UBA8248 | UBA8248 | UBA8248 sp002 | 0 | 0 | 0.1409204068 | 3.77E-05 | 0 | 1.000000199 | 0 | 0.368372361 | 0.001543955143 | 0.009777058201 | 0 |
| TARA_SAMEA2619970_METAG_GLICNMBG | 3 | 0.002026475414 | UBA828 | PTKB01 | PTKB01 sp9130 | 2.11E-06 | 0 | 0.02451043114 | 3.40E-05 | 0.4316129947 | 1 | 0 | 0.9833333333 | 0 | 1 | 0.06514894284 |
| TARA_SAMEA2619970_METAG_JMFNEFCE | 3 | -0.01901939411 | Marinisomatales | AB-629-J13 | | 8.45E-06 | 0 | 0.01262091767 | 4.18E-05 | 1 | 1 | 0 | 0 | 0.03689092147 | 1 | 0.001313261855 |
| TARA_SAMEA2619970_METAG_KIIHFBOD | 3 | -0.03753979419 | UBA3495 | UBA9611 | UBA9611 sp002 | 0 | 0 | 0.03219248396 | 1.25E-04 | 0.001707913229 | 1 | 0 | 0.07009043691 | 0 | 1 | 0 |
| TARA_SAMEA2619970_METAG_KLAFEPAG | 6 | -0.1091748404 | Nitrospinales | Nitromaritima | Nitromaritima sp0 | 0 | 0 | 0.01588017736 | 4.61E-05 | 1.000000004 | 0 | 0 | 0.05206494631 | 0 | 0.001660099114 | 0.03672209033 |
| TARA_SAMEA2619974_METAG_IOJPHMIM | 4 | -0.03690248487 | Acidimicrobiales | UBA3125 | UBA3125 sp002 | 0 | 0 | 1.000000203 | 1.000000203 | 0 | 0.007089302313 | 0 | 1.000000203 | 0 | 2.19E-05 | 0 |
| TARA_SAMEA2619974_METAG_PDHLOGOD | 5 | -0.229126977 | Cytophagales | Marinoscillum | Marinoscillum sp | 0 | 0 | 0.02175589545 | 0 | 0.003554504066 | 0.9999997074 | 0 | 0.05601549331 | 0 | 0.002571669996 | 0 |
| TARA_SAMEA2619992_METAG_FFIJIJBK | 5 | -0.1082719304 | Sneathiellales | Sneathiella | | 0 | 0 | 0.02462638411 | 1.05E-06 | 5.52E-05 | 0.9833333369 | 0 | 0.03237901739 | 0 | 0.002553211685 | 0 |
| TARA_SAMEA2620021_METAG_KBGMIOJP | 8 | -0.0315751398 | Acidimicrobiales | UBA8592 | UBA8592 sp002 | 0 | 0 | 0.9999997862 | 0.9833331195 | 8.61E-05 | 0.9999997862 | 0 | 0.07617172283 | 0 | 0 | 0 |
| TARA_SAMEA2620021_METAG_MLOOJBBB | 4 | -0.2954036986 | Micavibrionales | UBA2705 | UBA2705 sp002 | 7.76E-08 | 0 | 0.7648615125 | 0.02499278097 | 0.01964183817 | 0.005960853815 | 0 | 1.00000013 | 0 | 8.85E-04 | 0 |
| TARA_SAMEA2620035_METAG_GNMGNMHB | 2 | -0.07594258217 | Marinisomatales | UBA2126 | UBA2126 sp002 | 0 | 0 | 0.01941476017 | 2.80E-07 | 0.004588301572 | 0.01844270246 | 0 | 0.05020764473 | 0 | 1.33E-05 | 0 |
| TARA_SAMEA2620035_METAG_LBGPHFGM | 5 | -0.07318714067 | Marinisomatales | UBA2146 | UBA2146 sp002 | 2.84E-06 | 0 | 0.02727310728 | 1.28E-04 | 0.1131646049 | 1 | 0 | 0.03291871598 | 0 | 0.008428438024 | 0.04101923711 |
| TARA_SAMEA2620035_METAG_LBPLCHEF | 6 | -0.09820063375 | Rhodospirillales_ | Casp-alpha2 | Casp-alpha2 sp0 | 0 | 0 | 0.0258925889 | 3.68E-06 | 0.9842268447 | 8.35E-05 | 0 | 0.9523481211 | 0 | 0.003161968527 | 0.1289643162 |
| TARA_SAMEA2620035_METAG_LEKHLCMO | 2 | 0.00337876109 | Vicinamibacteral | UBA2161 | UBA2161 sp002 | 0 | 0 | 0.0187878303 | 9.02E-06 | 0.002037120231 | 0.1333333333 | 0 | 0.03398449921 | 0 | 1.83E-04 | 2.84E-04 |
| TARA_SAMEA2620039_METAG_BAKCDBNN | 5 | -0.1809188779 | Burkholderiales | Limnobacter | Limnobacter sp0 | 0 | 0 | 0.02470012865 | 3.43E-06 | 0.003149957913 | 0.9999996943 | 0 | 0.05601367435 | 0 | 0.01677191761 | 0 |
| TARA_SAMEA2620078_METAG_CJHBKDPI | 4 | -0.1384349386 | Flavobacteriales | Marixanthomona | Marixanthomona | 0 | 0 | 1 | 5.87E-05 | 0.07638164534 | 0 | 0 | 0.1022954958 | 0 | 8.26E-04 | 0.06727910918 |
| TARA_SAMEA2620097_METAG_HMMJHBHP | 8 | -0.04875525075 | Dehalococcoidal | GCA-2688105 | GCA-2688105 sp | 0.003255692875 | 0 | 0.1600723532 | 0.9999999979 | 6.55E-05 | 0.9999999979 | 0 | 0.9857713839 | 0 | 8.29E-04 | 0 |

| | | | | | | | | | | | | | | | | |
|---|---|---|---|---|---|---|---|---|---|---|---|---|---|---|---|---|
| TARA_SAMEA2620106_METAG_AFNJKAOM | 2 | -0.1936607155 | Micavibrionales | UBA2137 | UBA2137 sp002 | 0 | 0 | 0.05093071529 | 0.03333333585 | 0.01666666919 | 0 | 0 | 0.03268226638 | 0 | 0 | 0.3491045102 |
| TARA_SAMEA2620106_METAG_JMICEMNF | 8 | -0.3058094654 | Flavobacteriales | Polaribacter | Polaribacter sp0 | 0 | 0 | 0.9999999978 | 0.9999999978 | 2.68E-05 | 0.9999999978 | 3.30E-04 | 0.9707080374 | 3.50E-04 | 0.003594228232 | 0 |
| TARA_SAMEA2620230_METAG_HMHBGFAL | 5 | -0.05231283252 | Opitutales | MB11C04 | MB11C04 sp002 | 0 | 0 | 0.04582508485 | 1.09E-04 | 0.001576616435 | 0.9999996834 | 0 | 0.01757579077 | 0 | 7.47E-04 | 0 |
| TARA_SAMEA2620339_METAG_CMFLFIIF | 8 | -0.01298036201 | Flavobacteriales | UBA8444 | | 0 | 0 | 0.03821288149 | 0.9999996931 | 0.003247732771 | 0.9999996931 | 0 | 0.06839267305 | 0 | 7.71E-04 | 0 |
| TARA_SAMEA2620339_METAG_JDJMPEKP | 5 | -0.01115666137 | Flavobacteriales | MED-G14 | | 0 | 8.14E-04 | 0.03162707423 | 6.40E-05 | 0.6676257743 | 1 | 0 | 1 | 0 | 7.54E-04 | 0 |
| TARA_SAMEA2620339_METAG_NFMOIECC | 4 | -0.05211894233 | TMED127 | MED-G10 | MED-G10 sp902 | 3.03E-06 | 0 | 1 | 1.46E-05 | 0.004591051302 | 0.005975220818 | 0 | 0.06569406821 | 0 | 0.002267927179 | 0 |
| TARA_SAMEA2620384_METAG_KNFDGPKC | 4 | -0.185440269 | Caulobacterales | Brevundimonas | Brevundimonas | 0 | 0 | 0.6387639857 | 1.11E-06 | 0.001756850756 | 0.3833334034 | 0 | 0.04760209774 | 0 | 1.47E-04 | 0 |
| TARA_SAMEA2620384_METAG_PFNFNGAP | 4 | -0.227004311 | Caulobacterales | Hyphomonas | Hyphomonas sp | 0 | 0 | 1.000000004 | 1.50E-06 | 1.000000004 | 0 | 0 | 0.9844957425 | 0 | 3.08E-05 | 0 |
| TARA_SAMEA2620404_METAG_MDOJCKDP | 4 | 0.008443509969 | TMED127 | | | 0 | 0 | 0.9999997528 | 0.2090038003 | 2.52E-04 | 0.005027589086 | 0 | 0.03931581287 | 0.9833330819 | 0 | 0 |
| TARA_SAMEA2620570_METAG_EHBAIJHL | 5 | 0.0620528179 | Flavobacteriales | CAJWCP01 | | 0 | 0 | 0.02237175309 | 9.90E-07 | 0.07994248744 | 0.6666666658 | 0 | 0.0440862205 | 0 | 1.32E-05 | 0 |
| TARA_SAMEA2620663_METAG_MECILNPA | 4 | -0.2235917512 | Flavobacteriales | Aequorivita | Aequorivita sp00 | 0 | 0 | 0.9999999966 | 8.28E-07 | 0.0166666633 | 0 | 0 | 0.9894359794 | 0 | 0 | 0 |
| TARA_SAMEA2620666_METAG_AFGGNEFA | 4 | -0.145380196 | Sphingomonadal | Croceicoccus | Croceicoccus sp | 0 | 0 | 0.9849315604 | 1.52E-06 | 0.01107867442 | 3.90E-06 | 0.03021368646 | 1.000000202 | 0 | 1.30E-05 | 0 |
| TARA_SAMEA2620666_METAG_EDMBPCLD | 5 | -0.1721277098 | Micavibrionales | UBA2137 | UBA2137 sp002 | 0 | 0 | 0.04216432674 | 8.67E-05 | 2.39E-05 | 0.8999998816 | 0 | 0.2143520527 | 0 | 0.004383576389 | 0.03177682421 |
| TARA_SAMEA2620666_METAG_IPMNHCKD | 2 | -0.07031612545 | Flavobacteriales | Owenweeksia | Owenweeksia sp | 0 | 0 | 0.02872865685 | 4.57E-05 | 0.003402288489 | 0.004934509559 | 0 | 0.03771185456 | 0 | 8.13E-04 | 0.02089608959 |
| TARA_SAMEA2620672_METAG_GCIIDFDE | 7 | 0.02537160313 | Flavobacteriales | GCA-2693335 | GCA-2693335 sp | 2.49E-08 | 0 | 1.000000005 | 0.01666698119 | 1.000000005 | 0.02129669744 | 0 | 0.06270706605 | 0 | 0.9833341091 | 0 |
| TARA_SAMEA2620734_METAG_CDBINKLM | 2 | -0.0937754922 | UBA4151 | UBA4151 | UBA4151 sp002 | 1.04E-07 | 0 | 0.06932330985 | 8.75E-07 | 0.03526598377 | 0 | 0 | 0.06459865305 | 0 | 9.92E-06 | 0.07393458907 |
| TARA_SAMEA2620756_METAG_NONPAMNB | 3 | 0.06093758585 | SCGC-AAA003-I | GCA-002707645 | GCA-002707645 | 0 | 0 | 0.006105302497 | 9.43E-05 | 0.03744312766 | 0.9999997505 | 0 | 0.07097838271 | 0 | 0.9999997505 | 0.004514111776 |
| TARA_SAMEA2620812_METAG_BLLOGHKP | 2 | -0.1686027568 | Pseudomonadale | Pseudohongiella | Pseudohongiella | 3.42E-04 | 0 | 0.03773027228 | 5.30E-05 | 0.02234028078 | 0.007858913359 | 0 | 0.0917068039 | 0 | 8.09E-04 | 0.1089033897 |
| TARA_SAMEA2620812_METAG_HDCKCBGN | 5 | 0.003699396376 | Parvibaculales | Parvibaculum | Parvibaculum sp | 0 | 0 | 0.1117922191 | 2.29E-05 | 0.00343283089 | 1 | 0 | 0.09265845218 | 0 | 8.12E-04 | 0.03795662422 |
| TARA_SAMEA2620824_METAG_GEJHHNAG | 4 | -0.2749503605 | Flavobacteriales | Marinirhabdus | Marinirhabdus sp | 0 | 0 | 1 | 9.61E-05 | 1 | 2.05E-06 | 0 | 0.06716863619 | 0 | 0 | 0 |
| TARA_SAMEA2620824_METAG_IGMPJJKD | 3 | -0.1268476699 | Balneolales | Balneola | Balneola sp0023 | 0 | 0 | 0.3495406934 | 8.76E-05 | 0.003192256443 | 0.800000201 | 0 | 0.04997705161 | 0 | 1.00000016 | 0 |
| TARA_SAMEA2620824_METAG_MNBKNHDJ | 1 | -0.3156398337 | Pseudomonadale | UBA2679 | UBA2679 sp002 | 0 | 0 | 1.000000205 | 2.52E-05 | 5.38E-05 | 1.000000205 | 0 | 0.07617123998 | 0 | 0.01312114051 | 0.1344399265 |
| TARA_SAMEA2620824_METAG_NIGBPHMF | 6 | -0.1629141519 | Caulobacterales | Oceanicaulis | Oceanicaulis ale | 0 | 0 | 0.02550070899 | 3.92E-05 | 0.984793559 | 3.52E-06 | 0 | 0.997318571 | 0 | 0.01824769443 | 0 |
| TARA_SAMEA2620824_METAG_OAGJFFDB | 2 | -0.1315871288 | Rhodospirillales_ | UBA4479 | | 0 | 0 | 0.02940192593 | 1.12E-04 | 0.07383442108 | 0 | 0 | 0.07001242941 | 0 | 0.00686982616 | 0.002760309119 |
| TARA_SAMEA2620882_METAG_EKMOFPLF | 6 | -0.04317957728 | UBA8108 | UBA8108 | UBA8108 sp002 | 0 | 0 | 0.0445178195 | 1.03E-04 | 1.000000235 | 0 | 0 | 0.05840204809 | 0 | 0 | 0 |
| TARA_SAMEA2620925_METAG_LJBBMNFI | 4 | -0.1721486592 | Sphingomonadal | Qipengyuania | Qipengyuania cit | 3.93E-08 | 0 | 0.9999997875 | 0.06666783832 | 0.9999997875 | 0 | 0 | 0.1457704964 | 0 | 1.04E-05 | 0 |
| TARA_SAMEA2620947_METAG_GLLMAFGJ | 4 | 0.01313910379 | Parvibaculales | Parvibaculum | | 0 | 0 | 0.9677715476 | 0.0500000015 | 0.9676781768 | 4.97E-05 | 0 | 0.0295209717 | 0 | 0 | 0.004568052536 |
| TARA_SAMEA2620967_METAG_MHNHOBDL | 3 | -0.2163497171 | Flavobacteriales | Aequorivita | Aequorivita sp00 | 0 | 0 | 0.003110791069 | 1.07E-04 | 0.009029061315 | 0.999999783 | 0 | 0.09022509084 | 0 | 0.999999783 | 0.3201116396 |
| TARA_SAMEA2620970_METAG_GGBEMCNM | 2 | -0.02943270743 | Marinisomatales | Marinisoma | | 0 | 0.00306209757 | 0.03664942107 | 0.03342358936 | 0.111611009 | 0.01666765598 | 0 | 1.000000249 | 1.18E-07 | 7.64E-04 | 0 |
| TARA_SAMEA2620970_METAG_HEBCFGMN | 5 | -0.09833211664 | Pseudomonadale | UBA4582 | UBA4582 sp913 | 0 | 0 | 0.02344294299 | 1.27E-04 | 0.003345272616 | 1 | 0 | 1 | 3.86E-04 | 0.001851395532 | 0 |
| TARA_SAMEA2620980_METAG_EDCALDMI | 7 | -0.07228880293 | Rhodobacterales | Amylibacter | Amylibacter sp0 | 0 | 0 | 0.001569431892 | 2.09E-04 | 0.02689878758 | 3.75E-06 | 0.0466235651 | 0.03185740483 | 0 | 1 | 0 |
| TARA_SAMEA2620980_METAG_HOANNCOF | 2 | -0.03618255122 | Flavobacteriales | MAG-121220-bin8 | | 2.15E-04 | 0 | 0.02560187498 | 0.01682859926 | 0.1454800634 | 5.36E-06 | 0 | 0.065180109 | 0 | 0.001499968639 | 1.27E-04 |
| TARA_SAMEA2620980_METAG_MICENOFG | 2 | -0.03808434598 | Opitutales | MB11C04 | MB11C04 sp002 | 0 | 4.78E-05 | 0.0138782636 | 1.03E-04 | 3.95E-05 | 0 | 0.005272115045 | 0.06332318456 | 0 | 0 | 0 |
| TARA_SAMEA2620991_METAG_GHGOHFOA | 4 | -0.3269537656 | Flavobacteriales | Aequorivita | Aequorivita vladi | 0 | 0 | 1 | 1.42E-04 | 1 | 0.008813933863 | 0 | 0.1410490417 | 0 | 0 | 1 |
| TARA_SAMEA2620991_METAG_NLDPPCEE | 7 | -0.05867139763 | Flavobacteriales | Brumimicrobium | | 0 | 0 | 0.025762686 | 1.84E-04 | 0.0035049618 | 0.005973365184 | 0 | 0.07032812072 | 1.94E-05 | 1 | 0 |
| TARA_SAMEA2620995_METAG_BDJIHGBP | 2 | -0.09069533812 | Pseudomonadale | UBA11889 | UBA11889 sp913 | 0 | 0 | 0.03536498398 | 4.80E-05 | 0.001931686484 | 3.21E-05 | 0 | 0.06084094296 | 0 | 0.001018057752 | 8.61E-04 |
| TARA_SAMEA2621003_METAG_CKNNHJHJ | 3 | -0.06329201289 | UBA3495 | GCA-2712585 | GCA-2712585 sp | 0 | 0 | 1.000000135 | 8.14E-05 | 0.02383490429 | 1.000000135 | 0 | 0.06553797831 | 0 | 1.000000135 | 0 |
| TARA_SAMEA2621003_METAG_NGHODKEB | 4 | -0.05404692935 | Pseudomonadale | UBA9145 | UBA9145 sp002 | 6.65E-06 | 0.004522842551 | 0.9999997782 | 1.12E-04 | 0.006526094508 | 0 | 0 | 0.03383965423 | 0 | 0 | 0 |
| TARA_SAMEA2621020_METAG_ABKIOJHG | 3 | 4.96E-04 | Pelagibacterales | AG-414-E02 | | 0 | 0 | 0.03453689023 | 1.60E-04 | 0.9999997699 | 0.9999997699 | 0 | 0.2941511263 | 0 | 0.9999997699 | 0 |
| TARA_SAMEA2621021_METAG_BMDCHEJH | 2 | -0.01558382483 | Burkholderiales | BACL14 | | 9.55E-04 | 0 | 0.04210533582 | 1 | 0.007026222364 | 0.00525827775 | 0 | 1 | 0 | 0.00152773975 | 0.0379459263 |
| TARA_SAMEA2621044_METAG_PAAMBDEJ | 2 | -0.1504588251 | Bacteriovoracale | GCA-2706205 | GCA-2706205 sp | 2.94E-06 | 0 | 0.04731789677 | 0.9999999966 | 4.80E-05 | 8.32E-05 | 0 | 0.03383749534 | 0 | 4.02E-04 | 0.03391538579 |
| TARA_SAMEA2621075_METAG_KDKFOIAJ | 7 | -0.1828757695 | Bacteriovoracale | GCA-2707495 | GCA-2707495 sp | 5.35E-04 | 0 | 0.9999997003 | 0.00301338168 | 0.9833330336 | 1.18E-04 | 0 | 0.01894446341 | 2.66E-08 | 0.9837225132 | 0 |
| TARA_SAMEA2621076_METAG_PGLLOIOI | 4 | -0.03047612933 | Pedosphaerales | UBA1100 | UBA1100 sp913 | 0 | 0 | 1.000000225 | 1.07E-04 | 0.966873516 | 0.02720365211 | 0.01666667011 | 0.1912094055 | 0 | 0 | 0 |
| TARA_SAMEA2621085_METAG_NCMAHLFL | 2 | -0.209225754 | Flavobacteriales | UBA1494 | UBA1494 sp002 | 0 | 0 | 0.1775017104 | 7.12E-07 | 0.03432801472 | 9.65E-08 | 0 | 0.04659387345 | 0 | 1.31E-05 | 0 |
| TARA_SAMEA2621092_METAG_PINFDOMM | 6 | -0.345063989 | Pseudomonadale | Pseudomaricurvu | Pseudomaricurvu | 0 | 0 | 0.03698991047 | 6.75E-05 | 1 | 1 | 0 | 1 | 0 | 0.002322408729 | 0 |
| TARA_SAMEA2621099_METAG_MOEJCAFC | 4 | -0.338533726 | Bacteriovoracale | GCA-2712005 | GCA-2712005 sp | 0 | 0 | 0.9999997936 | 0.9999997936 | 0.0177395346 | 0.003628991555 | 0 | 0.2474613057 | 0 | 0 | 0 |
| TARA_SAMEA2621099_METAG_OCNFLOGF | 3 | -0.1103718095 | Flavobacteriales | GCA-2705995 | GCA-2705995 sp | 0 | 0 | 0.06931124065 | 1.17E-04 | 0.3050256776 | 0.8833333333 | 0 | 0.01138328424 | 0 | 0.9833969575 | 0 |
| TARA_SAMEA2621099_METAG_OLNLEHLG | 3 | -0.3712504115 | Pseudomonadale | Neptuniibacter | Neptuniibacter sp | 4.82E-06 | 0 | 0.01756664096 | 4.27E-05 | 1 | 1 | 0 | 1 | 0 | 1 | 0.06666534225 |
| TARA_SAMEA2621101_METAG_HCGFDFOF | 4 | -0.00717914471 | UBA3495 | UBA11650 | UBA11650 sp002 | 0 | 0 | 1 | 0.03341616901 | 0.1899352107 | 0.01776290601 | 0 | 0.04039716956 | 0 | 7.79E-04 | 0.003274629875 |
| TARA_SAMEA2621101_METAG_LEJMAEPD | 3 | -0.2712263244 | Bdellovibrionales | UBA6502 | UBA6502 sp002 | 0 | 0 | 0.02672608427 | 1.90E-06 | 0.118879183 | 0.9999997354 | 0 | 0.07122241346 | 0 | 0.9999997354 | 1.05E-06 |
| TARA_SAMEA2621107_METAG_MBDOFGMO | 2 | -0.2174175328 | Pseudomonadale | Pseudohongiella | Pseudohongiella | 0 | 0 | 0.03771615873 | 8.11E-07 | 0.004895600485 | 8.64E-08 | 0 | 0.1814033854 | 0 | 0 | 0 |
| TARA_SAMEA2621176_METAG_LNNLNJLH | 5 | -0.2739420333 | Pseudomonadale | Oleibacter | Oleibacter sp002 | 0.0171615123 | 0 | 0.09098010594 | 0 | 0.04343605368 | 1 | 0.003940144864 | 0.1493490232 | 0 | 7.69E-04 | 0 |
| TARA_SAMEA2621191_METAG_JOABIJEI | 2 | -0.138363929 | Rhodobacterales | Roseovarius | Roseovarius muc | 0 | 0 | 0.09565175438 | 4.61E-05 | 1.06E-06 | 0.003591876293 | 0 | 1 | 4.15E-04 | 8.01E-04 | 0.06936134843 |
| TARA_SAMEA2621191_METAG_KOACLFMB | 2 | -0.0908428602 | Rhodobacterales | Actibacterium_A | Actibacterium_A | 3.13E-06 | 0 | 0.1733464382 | 1.47E-04 | 0.009069171212 | 0.01666666233 | 0 | 0.0618150064 | 4.93E-04 | 0.01877025467 | 0 |
| TARA_SAMEA2621203_METAG_IEFLJMCG | 1 | -0.535307013 | Pseudomonadale | Acinetobacter | Acinetobacter ve | 0 | 0 | 1 | 1.04E-04 | 0.008234576429 | 1 | 0 | 0.1924195406 | 0 | 0 | 0 |
| TARA_SAMEA2621213_METAG_DIBPDECH | 6 | -0.00159816920 | Phycisphaerales | JANSXG01 | | 0.02685239798 | 0 | 0.1405544444 | 3.51E-05 | 0.9857014608 | 0.00534166207 | 0 | 0.08978313108 | 0 | 6.47E-04 | 0.005867567491 |
| TARA_SAMEA2621216_METAG_DIJMOGIJ | 6 | -0.05829479299 | WGA-4E | TMED15 | TMED15 sp0027 | 0.00841928012 | 0 | 0.05382678273 | 2.78E-06 | 0.9904890065 | 1.34E-04 | 0 | 0.06297286956 | 0.02160509695 | 7.61E-04 | 1.66E-04 |
| TARA_SAMEA2621221_METAG_HEHENJAP | 7 | -0.03200770718 | UBA1144 | | | 0 | 0.02905421671 | 1.000000239 | 8.18E-05 | 9.74E-05 | 2.94E-06 | 0 | 0.06436900052 | 0 | 1.000000239 | 0 |
| TARA_SAMEA2621222_METAG_DGLCEMDP | 5 | -0.0891428922 | Opitutales | UBA5691 | UBA5691 sp002 | 2.54E-06 | 0 | 0.02693490449 | 1.57E-04 | 0.02100032448 | 0.9999995769 | 0 | 0 | 0 | 7.45E-04 | 0.06964035887 |
| TARA_SAMEA2621232_METAG_JNKIOFLF | 3 | 0.004532640014 | AEGEAN-169 | MarineAlpha5-Bi | MarineAlpha5-Bi | 0 | 0 | 1.000000268 | 5.89E-06 | 1.000000268 | 1.000000268 | 0 | 1.000000268 | 0 | 1.000000268 | 0.08272800873 |
| TARA_SAMEA2621242_METAG_AIOKPNEJ | 6 | 0.04948015977 | Phycisphaerales | GCA-2706885 | GCA-2706885 sp | 0 | 0 | 0.03431466863 | 2.34E-06 | 1.000000219 | 2.94E-08 | 0 | 0.1367929412 | 0 | 0 | 0 |
| TARA_SAMEA2621242_METAG_AOBACFJI | 6 | -0.2478289385 | Flavobacteriales | Muricauda | Muricauda aurea | 7.68E-08 | 0 | 0.1021914737 | 6.86E-06 | 0.9842398945 | 0 | 0 | 0.1978883138 | 0 | 8.15E-05 | 0 |
| TARA_SAMEA2621242_METAG_GDMEHIBJ | 4 | -0.2001493726 | Flavobacteriales | Leeuwenhoekiell | Leeuwenhoekiell | 0 | 0 | 1 | 0.001789969069 | 1 | 0.004984402196 | 0 | 0.05750188574 | 0 | 3.13E-06 | 1.08E-04 |
| TARA_SAMEA2621242_METAG_HEAAMFCE | 4 | -0.01490803686 | UBA1113 | GCA-2707205 | GCA-2707205 sp | 0 | 0 | 1 | 1.48E-04 | 1 | 0.007586402409 | 0 | 0.06779877003 | 0 | 0.00160440566 | 0 |
| TARA_SAMEA2621242_METAG_HOBIHNJG | 4 | -0.2017006551 | Flavobacteriales | Flavobacterium | Flavobacterium c | 0 | 0 | 0.9999998003 | 0.9999998003 | 0.004995794896 | 0 | 0 | 0.05626710227 | 0 | 0.01288712967 | 0 |
| TARA_SAMEA2621247_METAG_FOKJHEGI | 5 | -0.1367984823 | Pseudomonadale | Litorivivens | Litorivivens sp00 | 0 | 0.006474484742 | 0.03300353437 | 2.88E-06 | 0.1706312762 | 0.9666861311 | 0 | 0.02090447554 | 0 | 0.00109894325 | 0 |
| TARA_SAMEA2621259_METAG_OOAMCINK | 7 | -0.03661402448 | Comchoanobacte | UBA6594 | UBA6594 sp002 | 0 | 0 | 0.983885309 | 1.73E-06 | 1 | 0.01772018055 | 0 | 1 | 0 | 1 | 0 |
| TARA_SAMEA2621277_METAG_JANMBIJC | 3 | -0.00553254511 | Acidimicrobiales | UBA8592 | UBA8592 sp002 | 0 | 0 | 0.02859587367 | 1.47E-04 | 1 | 1 | 0 | 1 | 0 | 1 | 0.06482975382 |
| TARA_SAMEA2621287_METAG_CBFAEDAG | 5 | -0.268018024 | Enterobacterales | UBA3532 | UBA3532 sp002 | 0.001476536445 | 0 | 0.02643306427 | 1.15E-05 | 8.44E-04 | 1.000000015 | 0 | 0.02998946303 | 0 | 7.07E-04 | 0 |
| TARA_SAMEA2621287_METAG_FFNGLIJO | 8 | -0.2161758851 | Burkholderiales | Limnobacter | Limnobacter sp0 | 0 | 0 | 1.000000188 | 1.000000188 | 0.09615564261 | 1.000000188 | 0.01764978645 | 0.04603469998 | 2.03E-04 | 7.18E-04 | 0 |
| TARA_SAMEA2621300_METAG_CCGLMDCH | 5 | -0.1610548805 | Planctomycetale | Gimesia | Gimesia sp0027 | 0 | 0 | 0.01982755271 | 0.002914543643 | 0.001994802529 | 1.000000159 | 0 | 0.1000272127 | 0 | 1.33E-05 | 0 |
| TARA_SAMEA2621300_METAG_EKFPEFJE | 4 | -0.04827533644 | Rhodospirillales_ | UBA1479 | UBA1479 sp002 | 0 | 0 | 1 | 7.83E-05 | 1 | 0 | 0 | 1 | 0 | 0 | 0 |
| TARA_SAMEA2621300_METAG_GGKHKFLB | 8 | -0.04335633599 | Balneolales | UBA2664 | UBA2664 sp002 | 0 | 6.28E-04 | 0.08520328893 | 0.9999998134 | 0.001739733677 | 0.9999998134 | 0 | 0.03619519274 | 0 | 7.90E-04 | 0 |

| | | | | | | | | | | | | | | | | |
|---|---|---|---|---|---|---|---|---|---|---|---|---|---|---|---|---|
| TARA_SAMEA2621401_METAG_PBJKAHMH | 1 | -0.1241008057 | Flavobacteriales | SCGC-AAA160-P02 | | 0 | 0 | 0.9683512933 | 0 | 0.06372756848 | 0.9500004828 | 0 | 0.06097360919 | 0 | 2.22E-04 | 5.35E-04 |
| TARA_SAMEA2621487_METAG_HMPOEMOI | 5 | -0.1174234713 | Flavobacteriales | UBA6049 | UBA6049 sp913 | 0 | 0 | 0.03119448932 | 9.60E-05 | 0.07561325147 | 0.9999997623 | 0 | 0.06732178723 | 0 | 0.01926616387 | 0 |
| TARA_SAMEA2621502_METAG_HFKILBIH | 2 | -0.2170924022 | Pseudomonadale | Marinobacter | Marinobacter ma | 0 | 0 | 0.01785701552 | 0.1544087062 | 0 | 0.002606009986 | 0 | 0.01011124268 | 0 | 0.01886386141 | 0 |
| TARA_SAMEA2621502_METAG_NLHALEFI | 4 | 0.005769696023 | Acidimicrobiales | UBA9382 | UBA9382 sp0034 | 0 | 0 | 0.9999997196 | 0.01666666667 | 0.9683096922 | 0 | 0 | 0.07706222576 | 0 | 0 | 0.06604925159 |
| TARA_SAMEA2621509_METAG_MPBCKPHO | 1 | -0.1800858867 | Rhodobacterales | Amylibacter | Amylibacter sp91 | 0 | 0 | 1.000000288 | 0.01666787872 | 0.06885634213 | 0.96666695 | 0 | 0.123378655 | 0 | 0 | 0 |
| TARA_SAMEA2621528_METAG_FCNHCPFO | 2 | -0.1638480917 | Pseudomonadale | Pseudohongiella | Pseudohongiella | 0 | 0 | 0.01754882121 | 0.9999997012 | 0.00347376631 | 1.95E-05 | 0 | 0.04030246393 | 0 | 1.89E-05 | 0.03337431521 |
| TARA_SAMEA2621528_METAG_GAEKHINC | 5 | -0.2951674036 | Rhodobacterales | Sulfitobacter | Sulfitobacter sp0 | 0 | 0 | 0.0173769009 | 3.06E-05 | 0.002636556482 | 0.9999999949 | 0 | 0.05491231819 | 0 | 0.01746165177 | 0.06774346646 |
| TARA_SAMEA2621528_METAG_LMPJOJBM | 4 | -0.1783199912 | Rhodobacterales | Sulfitobacter | | 0 | 0 | 1.000000205 | 1.35E-04 | 1.000000205 | 0.00537646405 | 0 | 0.17833181 | 0 | 0 | 0 |
| TARA_SAMEA2621536_METAG_AMHKKGFA | 3 | -0.162611112 | Flavobacteriales | UBA10364 | UBA10364 sp003 | 0 | 0 | 0.999999584 | 1.98E-04 | 0.09376067405 | 0.999999584 | 0 | 0.06841628568 | 0 | 0.999999584 | 0 |
| TARA_SAMEA2621536_METAG_ECBGEDBK | 8 | 0.01658743054 | Flavobacteriales | CAJIYG01 | CAJIYG01 sp913 | 0 | 0 | 1.000000303 | 1.000000303 | 0 | 1.000000303 | 0 | 0.03726406903 | 0 | 0 | 0 |
| TARA_SAMEA2621536_METAG_HLKLHGDE | 7 | -0.2636403567 | Flavobacteriales | UBA6772 | UBA6772 sp002 | 0.008003386116 | 0 | 0.090957872 | 0.0167917287 | 0.1743017746 | 4.09E-05 | 0 | 0.03941454776 | 0 | 1.000000005 | 0 |
| TARA_SAMEA2621536_METAG_JKOGBKOH | 4 | -0.05687239765 | Flavobacteriales | | | 0 | 0 | 1 | 1.35E-04 | 0.1178096272 | 0.003660721877 | 0 | 0.05453532757 | 0 | 0 | 0 |
| TARA_SAMEA2621536_METAG_KLKANNOC | 1 | -0.129303762 | Flavobacteriales | ASP10-05a | ASP10-05a sp00 | 0 | 0 | 1 | 0.01674910318 | 2.88E-04 | 0.9849042018 | 0 | 0.9865772158 | 0 | 7.68E-04 | 7.24E-05 |
| TARA_SAMEA2621551_METAG_HDAJGIEO | 2 | -0.06044388733 | UBA1151 | UBA1328 | UBA1328 sp002 | 0 | 0 | 0.007059405869 | 6.64E-05 | 0.07152590027 | 0 | 0 | 0.03542714389 | 0 | 0.01297546483 | 0 |
| TARA_SAMEA2621551_METAG_JDNJPCNE | 7 | -0.05527133481 | UBA8231 | UBA8231 | UBA8231 sp003 | 0 | 0 | 1.000000207 | 3.60E-05 | 1.000000207 | 0.005058553397 | 0 | 0.8872307937 | 0 | 1.000000207 | 0 |
| TARA_SAMEA2621551_METAG_KGBINJLE | 2 | -0.03528235028 | Verrucomicrobial | Roseibacillus_B | | 0 | 0 | 0.1005764918 | 2.47E-06 | 0.2128395668 | 0 | 0 | 0.05548428785 | 0 | 0.002404944022 | 0.02201014129 |
| TARA_SAMEA2621551_METAG_KNGBNMOK | 4 | -0.3197250939 | Enterobacterales | Colwellia | Colwellia sp0035 | 3.46E-08 | 0 | 1.000000159 | 1.56E-04 | 0.9700848172 | 0.03947122394 | 0 | 1.000000159 | 0.01099514847 | 0.0599217931 | 9.74E-06 |
| TARA_SAMEA2621551_METAG_MMPJBINL | 6 | -0.0776802583 | Flavobacteriales | Polaribacter | Polaribacter sp9 | 0 | 0 | 0.4099823829 | 0.01670881079 | 1.000000216 | 0.05477005178 | 0.003742353666 | 1.000000216 | 0 | 8.45E-04 | 0.06270981094 |
| TARA_SAMEA2621551_METAG_MNJLJAEC | 7 | -0.04444420878 | SAR324 | UBA8110 | UBA8110 sp002 | 0 | 0 | 0.04269851083 | 2.18E-06 | 0.01120185741 | 1.04E-07 | 0 | 0.07864766007 | 0 | 0.9999999953 | 0.2559031816 |
| TARA_SAMEA2621551_METAG_PDBJEGFH | 5 | -0.08391499376 | Pirellulales | GCA-2718815 | | 2.42E-06 | 0 | 0.02644210465 | 3.33E-05 | 0 | 1.000000303 | 0 | 0.05323841762 | 0 | 0.01376023609 | 0 |
| TARA_SAMEA2621564_METAG_GKALJALI | 5 | -0.1316140259 | Sphingomonadal | Parasphingorhab | Parasphingorhab | 0 | 0.01896229621 | 0.1931920745 | 1.61E-06 | 5.32E-04 | 0.9671536992 | 1.09E-04 | 0.06316350028 | 2.30E-04 | 0 | 0.05099259745 |
| TARA_SAMEA2621779_METAG_IOENAFMJ | 4 | -0.02920106695 | NS11-12g | UBA9320 | UBA9320 sp913 | 0 | 0 | 1.000000201 | 9.08E-05 | 1.000000201 | 0.009996671927 | 0 | 0.06383111626 | 0 | 0 | 0 |
| TARA_SAMEA2621812_METAG_BHEJFGLB | 6 | -0.08695035487 | Woeseiales | UBA1847 | UBA1847 sp002 | 0 | 0 | 0.1875233584 | 0.0333351426 | 0.9999999951 | 0.9833333285 | 0 | 0.0570698082 | 0 | 0.001461135456 | 1.64E-07 |
| TARA_SAMEA2621812_METAG_EINIOJLI | 8 | -0.1132653747 | Flavobacteriales | MS024-2A | MS024-2A sp002 | 0.1617765055 | 0 | 0.04477551522 | 1.00000038 | 0.005229716018 | 1.00000038 | 0 | 0.9995834032 | 0 | 8.29E-04 | 0.01299553614 |
| TARA_SAMEA2621812_METAG_MHCAACBH | 2 | -0.06804141436 | Flavobacteriales | UBA8444 | | 0.001419089203 | 0 | 0.04871555198 | 0.01666883549 | 0.002714146496 | 0.01669334782 | 0 | 0.1041743172 | 0 | 0.008281366246 | 0 |
| TARA_SAMEA2621812_METAG_MOHOOAMM | 1 | -0.1080161264 | PCC-6307 | Synechococcus_ | Synechococcus_ | 0 | 0 | 1.000000216 | 1.57E-04 | 1.000000216 | 1.000000216 | 0 | 0.08400576683 | 0 | 0.02042061957 | 0 |
| TARA_SAMEA2621812_METAG_NHHPCCBJ | 1 | -0.1147147067 | Phycisphaerales | UBA1668 | UBA1668 sp002 | 0 | 0 | 0.9801964676 | 1.23E-06 | 0.9842727178 | 1 | 0 | 0.06259915857 | 0.001070252026 | 5.15E-05 | 0 |
| TARA_SAMEA2621839_METAG_CHKPNCKL | 4 | -0.05941555027 | GCA-002705445 | GCA-2720195 | GCA-2720195 sp | 0 | 9.11E-04 | 0.9839082428 | 0.0166678419 | 0.09012359271 | 0.01666671487 | 0 | 0.07699811843 | 0 | 2.53E-05 | 0 |
| TARA_SAMEA2621839_METAG_ECLMIDKH | 4 | -0.01789259163 | Verrucomicrobial | Roseibacillus_B | Roseibacillus_B | 0 | 0 | 0.9965661771 | 1.02E-06 | 1.000000215 | 0.007824152208 | 0 | 0.9846712452 | 0 | 1.43E-04 | 0 |
| TARA_SAMEA2621839_METAG_EOEPIHHM | 4 | -0.1256480883 | Flavobacteriales | UBA952 | UBA952 sp9130 | 0 | 0 | 1 | 5.69E-05 | 1 | 5.64E-06 | 0 | 1 | 0 | 0.001608702422 | 0 |
| TARA_SAMEA2621839_METAG_FOCILNML | 6 | -0.1278707186 | Opitutales | UBA7441 | UBA7441 sp913 | 7.22E-08 | 0 | 0.1353519312 | 0 | 0.9833333333 | 8.66E-08 | 0 | 0.06634548996 | 0 | 7.54E-05 | 0.06611555592 |
| TARA_SAMEA2621839_METAG_HILMGKBK | 7 | -0.04096551371 | Burkholderiales | BACL14 | | 0 | 0 | 0.9999998138 | 1.29E-04 | 0.9999998138 | 0.06150812452 | 0.001223505074 | 0.06685341936 | 0 | 0.9999998138 | 0 |
| TARA_SAMEA2621839_METAG_NEIJCBGE | 6 | -0.00412628815 | SAR86 | TMED112 | TMED112 sp002 | 0 | 0 | 0.01682674195 | 1.37E-04 | 0.9999996911 | 0 | 0 | 0.01081199297 | 0 | 0 | 0 |
| TARA_SAMEA2621859_METAG_BLBOAJGJ | 2 | -0.07234122797 | Flavobacteriales | UBA7446 | UBA7446 sp002 | 0.01974743555 | 0 | 0.1250681732 | 1.32E-04 | 0.05679942527 | 0.005904801522 | 0 | 0.06221291097 | 0 | 0 | 0 |
| TARA_SAMEA2621859_METAG_GAEHGPFE | 7 | -0.07859130943 | Opitutales | MB11C04 | MB11C04 sp002 | 0 | 0 | 1 | 1.06E-04 | 0.0434324786 | 0 | 0 | 0.06146563563 | 0 | 1 | 1 |
| TARA_SAMEA2621990_METAG_CKNIILJE | 5 | 0.02493799835 | Pelagibacterales | HIMB114 | HIMB114 sp0032 | 0 | 0 | 0.03267076792 | 0 | 0.342887265 | 0.9999996095 | 0 | 0.04514555809 | 0 | 0.008282685899 | 0 |
| TARA_SAMEA2621990_METAG_OMFNCAAD | 1 | -0.01576159751 | Flavobacteriales | MED-G13 | MED-G13 sp902 | 0 | 0 | 0.9999998411 | 0.01666666667 | 0.01666666667 | 0.9999998411 | 0.02951622619 | 0.9977704696 | 0 | 2.00E-05 | 0 |
| TARA_SAMEA2622097_METAG_DKMMMDIM | 1 | -0.1001270051 | Flavobacteriales | UBA952 | UBA952 sp9130 | 0 | 0 | 1.000000207 | 1.63E-04 | 1.000000207 | 1.000000207 | 0 | 0.2073678056 | 0 | 0 | 0 |
| TARA_SAMEA2622097_METAG_HGADEAJE | 1 | -0.03574799951 | Flavobacteriales | SP287 | SP287 sp002721 | 0 | 0 | 1.000000258 | 1.84E-04 | 0.008587738726 | 1.000000258 | 0 | 1.000000258 | 0 | 0.002335869063 | 0.1094100421 |
| TARA_SAMEA2622097_METAG_NFKPCCND | 2 | -0.00175400190 | Verrucomicrobial | Roseibacillus_B | Roseibacillus_B | 0 | 0 | 0.03325845562 | 9.48E-05 | 0.05816214643 | 0.005047644962 | 0 | 0.07708381041 | 0 | 0 | 0.03497735618 |
| TARA_SAMEA2622119_METAG_AHPOGDIA | 2 | -0.3678870095 | Cytophagales | Fabibacter | Fabibacter therm | 0 | 0 | 0.04052310999 | 8.91E-07 | 0.001056755187 | 7.83E-05 | 0 | 0.08457614049 | 0 | 0.005413038324 | 0 |
| TARA_SAMEA2622119_METAG_CKEFGNEL | 4 | -0.07916518743 | Rhodospirillales_ | UBA1479 | UBA1479 sp913 | 0 | 0 | 1 | 1 | 0.006824298735 | 0 | 0 | 1 | 0 | 0.0015617316 | 0.06773021611 |
| TARA_SAMEA2622149_METAG_FEMNCMMF | 6 | -0.07834607413 | Nitrospinales | Nitromaritima | Nitromaritima sp | 0 | 0 | 0.0319490792 | 7.91E-05 | 1 | 1 | 0 | 0.05093138261 | 0 | 0.007816715583 | 0 |
| TARA_SAMEA2622149_METAG_KGNNLAEL | 2 | -0.0624331093 | UBA1144 | NP136 | NP136 sp002730 | 0 | 0 | 0.0426500942 | 1 | 0 | 0 | 0 | 0 | 0 | 4.04E-05 | 0 |
| TARA_SAMEA2622149_METAG_MMILIKLJ | 6 | -0.1165561522 | Phycisphaerales | GCA-002718515 | GCA-002718515 | 0 | 0 | 0.0295407376 | 2.26E-04 | 0.9999996923 | 0 | 0 | 0.9999996923 | 0 | 0.001537285165 | 0 |
| TARA_SAMEA2622149_METAG_PDNNHHPM | 3 | -0.03343240645 | Longimicrobiales | UBA1138 | UBA1138 sp002 | 0 | 0 | 0.01976399589 | 1.94E-05 | 0.9999996771 | 0.9999996771 | 0 | 0.9999996771 | 0 | 0.9999996771 | 0 |
| TARA_SAMEA2622197_METAG_CGIMDGAD | 4 | -0.1012235502 | SAR324 | Arctic96AD-7 | Arctic96AD-7 sp | 1 | 0 | 1 | 1.36E-04 | 1 | 0.00450900144 | 0 | 1 | 0 | 0.004165757632 | 0 |
| TARA_SAMEA2622219_METAG_FGDKILBA | 6 | 0.0230537645 | Burkholderiales | BACL14 | | 0 | 0 | 0.04414583774 | 1.02E-04 | 1 | 0.00340749163 | 0 | 0.08054681592 | 0 | 7.18E-04 | 0 |
| TARA_SAMEA2622219_METAG_GIOFANNH | 8 | 0.0410589668 | Flavobacteriales | GCA-2697505 | | 0 | 0 | 0.09463368132 | 1 | 0.002984183693 | 1 | 0 | 0.04531157078 | 0 | 0.001415694249 | 0 |
| TARA_SAMEA2622219_METAG_GJKCCBDG | 5 | -0.1043244627 | Rhodobacterales | MED-G52 | MED-G52 sp002 | 0 | 0 | 0.03720225279 | 3.34E-07 | 0.4220440336 | 0.9833333715 | 0 | 0.08722966709 | 0 | 9.08E-04 | 0 |
| TARA_SAMEA2622316_METAG_PCEAOEII | 2 | -0.1226359983 | Verrucomicrobial | Arctic95D-9 | Arctic95D-9 sp0 | 9.67E-04 | 0 | 0.04531100481 | 8.19E-07 | 0.1192697665 | 0.01844105448 | 9.78E-04 | 0.0315202986 | 0 | 0.004358334383 | 0 |
| TARA_SAMEA2622336_METAG_ELNAHGFM | 6 | -0.05418768416 | TMED25 | TMED25 | TMED25 sp9130 | 0 | 0 | 0.03853432729 | 3.14E-05 | 0.9858719416 | 1.17E-06 | 0 | 0.07513389765 | 0 | 0.01678334195 | 0 |
| TARA_SAMEA2622336_METAG_NINLODAE | 3 | -0.04334814011 | SAR86 | TMED112 | TMED112 sp902 | 0 | 0 | 1 | 8.80E-05 | 0.2430452187 | 1 | 0 | 0.7569095861 | 0 | 1 | 0.02920432935 |
| TARA_SAMEA2622362_METAG_ILIBDBIG | 3 | 0.006655624487 | Pelagibacterales | CACPJE01 | | 0.04205998019 | 0 | 0.003724764447 | 1.73E-06 | 0.9999999966 | 0.9833333333 | 0 | 0.04184547549 | 0 | 0.9999999966 | 0.03395496748 |
| TARA_SAMEA2622376_METAG_GLDFILLI | 7 | -0.09214655892 | Rhodobacterales | MED-G52 | MED-G52 sp902 | 1.00000016 | 0 | 1.00000016 | 1.45E-04 | 0 | 0.004599136615 | 0 | 0.1421480724 | 0 | 1.00000016 | 0.02808640396 |
| TARA_SAMEA2622376_METAG_KBAMCEAM | 6 | -0.1677082854 | Flavobacteriales | CAJXFJ01 | CAJXFJ01 sp913 | 0 | 0 | 0.04608405832 | 0.03347576273 | 0.9999999857 | 0.0499999857 | 0 | 0.09764653835 | 1.00E-07 | 0.01817005675 | 0 |
| TARA_SAMEA2622478_METAG_HAEEGDCL | 4 | -0.04616907531 | SAR324 | Arctic96AD-7 | | 0 | 0 | 0.9999999025 | 1.45E-06 | 6.58E-04 | 0.001088949524 | 0 | 0.03402994015 | 0 | 0.004708068064 | 2.13E-04 |
| TARA_SAMEA2622499_METAG_LMOJHPFB | 2 | 0.1147128416 | Cytophagales | | | 0 | 0 | 0.03915948458 | 1 | 0 | 0 | 0 | 0.04329474643 | 0 | 4.37E-04 | 0 |
| TARA_SAMEA2622518_METAG_CDPNIGGJ | 6 | 0.005755161519 | UBA1144 | GCA-002715585 | GCA-002715585 | 0 | 0 | 0.07013888921 | 1.54E-04 | 1 | 0.003880775545 | 0 | 0.07619225149 | 0 | 0 | 0.04540276113 |
| TARA_SAMEA2622545_METAG_HPKFPAAP | 5 | -0.06681642392 | SCGC-AAA003-L08 | | | 0.01259944903 | 0 | 0.02428611131 | 3.51E-05 | 0.00994647863 | 1 | 0.003260164725 | 0.07606346937 | 0 | 0.02721595539 | 0.2447880251 |
| TARA_SAMEA2622545_METAG_ODBBDHAI | 4 | -4.28E-04 | PCC-6307 | Prochlorococcus_A | | 0 | 0 | 0.9846500765 | 3.94E-06 | 0.1303106992 | 0.01667094695 | 3.41E-05 | 0.9688393438 | 0 | 0.02609153229 | 0 |
| TARA_SAMEA2622545_METAG_OECGPLDK | 4 | -0.1604055899 | Pirellulales | UBA721 | UBA721 sp0027 | 0 | 0 | 1 | 8.76E-05 | 1 | 0 | 6.26E-06 | 0.06864169457 | 0 | 0.004658438996 | 0.03722084791 |
| TARA_SAMEA2622568_METAG_BPIONGFC | 5 | -0.176232124 | Flavobacteriales | Mesonia | Mesonia oceanic | 0 | 0.1020487728 | 0.03508467132 | 1.28E-04 | 0.08858349383 | 1.000000188 | 0 | 0.04985704549 | 0 | 7.99E-04 | 5.05E-05 |
| TARA_SAMEA2622656_METAG_FODMCDIL | 5 | -0.06220130091 | Pseudomonadale | PGZG01 | | 0 | 0 | 0.02673100562 | 5.89E-07 | 5.36E-05 | 1.000000172 | 0 | 0.97053088 | 0 | 0.01666667174 | 0 |
| TARA_SAMEA2622656_METAG_JDDFFOHG | 4 | -0.04557466123 | Burkholderiales | ALPHA2B | ALPHA2B sp901 | 0 | 0 | 1 | 1.99E-04 | 2.19E-04 | 8.46E-07 | 0 | 0.06837588766 | 0 | 0.01096296021 | 0 |
| TARA_SAMEA2622657_METAG_BLIPHDIP | 6 | -0.02647416047 | GCA-002705445 | GCA-2705445 | GCA-2705445 sp | 0 | 0 | 0.03209422323 | 1.000000283 | 1.000000283 | 0.005162494454 | 0 | 1.000000283 | 0 | 0 | 0 |
| TARA_SAMEA2622658_METAG_MEGADAMM | 4 | 0.04209712973 | Flavobacteriales | GCA-2697505 | | 7.81E-08 | 0 | 1.000000233 | 0.01678365007 | 1.000000233 | 0.01666666667 | 0 | 0.9835374364 | 0 | 1.24E-05 | 0.0367477019 |
| TARA_SAMEA2622658_METAG_PNALDEAO | 1 | 0.02727695361 | Cytophagales | MED-G16 | | 2.13E-08 | 0 | 1 | 0.1645274599 | 0.006495446894 | 0.9833333333 | 0 | 0.05794945897 | 0 | 0.01235097799 | 0.001157213657 |
| TARA_SAMEA2622667_METAG_BMCGBAMJ | 2 | -0.09031633558 | Phycisphaerales | UBA1924 | UBA1924 sp002 | 0 | 0 | 0.03314722312 | 6.28E-05 | 0.001509699089 | 0.004378259672 | 0 | 0.05610695444 | 0 | 0 | 0 |
| TARA_SAMEA2622667_METAG_IKEGAHGN | 2 | -0.1741005867 | Pseudomonadale | Litorivivens | Litorivivens sp00 | 0 | 0 | 0.04691370141 | 1.22E-04 | 0.003411619485 | 0 | 0 | 0.06666619932 | 0 | 0 | 0 |
| TARA_SAMEA2622673_METAG_EOOHAFPO | 5 | -0.1201722089 | Pseudomonadale | UBA11889 | UBA11889 sp002 | 0 | 0 | 0.03355954956 | 1.04E-04 | 1.76E-04 | 1 | 0 | 0.01765619766 | 0 | 0.01264782986 | 0.009882604529 |
| TARA_SAMEA2622678_METAG_LCCFEJME | 2 | -0.1242054182 | Legionellales | GCA-2863045 | GCA-2863045 sp | 0 | 0 | 0.03011358384 | 1.74E-04 | 0.003220737267 | 6.19E-06 | 0 | 0.06961427669 | 0 | 0.007370720482 | 0.0954195384 |
| TARA_SAMEA2622685_METAG_HEEHHBLE | 5 | -0.3204062277 | Rhodobacterales | Sulfitobacter | Sulfitobacter sp0 | 0 | 0 | 0.2795123133 | 1.43E-04 | 0.001223846916 | 1.000000005 | 0 | 0.4772909918 | 0 | 7.97E-04 | 0.04503566869 |

| | | | | | | | | | | | | | | | | |
|---|---|---|---|---|---|---|---|---|---|---|---|---|---|---|---|---|
| TARA_SAMEA2622685_METAG_JPLIAAIG | 2 | -0.276724628 | Pseudomonadale | Marinobacter_A | Marinobacter_A s | 0 | 0 | 0.02160850276 | 2.08E-06 | 4.57E-04 | 0.001039776275 | 1.16E-05 | 0.3016123921 | 0 | 0.001592243294 | 2.49E-05 |
| TARA_SAMEA2622690_METAG_BHOFBPFK | 2 | 0.007788857554 | Pelagibacterales | GCA-2704625 | | 0 | 0.01775094089 | 0.1965572975 | 1.68E-05 | 0.1125251699 | 0.4003649462 | 0.02577457571 | 0.04362935698 | 0.002003903842 | 4.43E-04 | 0.02660087879 |
| TARA_SAMEA2622694_METAG_JHIJAEDG | 1 | -0.3484644852 | Enterobacterales | Pseudidiomarina | Pseudidiomarina | 3.72E-04 | 0 | 0.9999996869 | 1.41E-06 | 0.2448204424 | 0.983333024 | 1.15E-04 | 0.05297787747 | 0 | 0.0025692281 | 0 |
| TARA_SAMEA2622695_METAG_BPMNPGCH | 4 | 0.03911317739 | SCGC-AAA003-L | TMED161 | | 0 | 0 | 0.9999998378 | 0 | 0 | 0.07999187725 | 0 | 0.1258990291 | 3.24E-07 | 0.001756576742 | 8.94E-05 |
| TARA_SAMEA2622695_METAG_GIMEDFFN | 5 | 0.0227842574 | GCA-2716485 | GCA-002719695 | GCA-002719695 | 0 | 0 | 0.07320065756 | 1.80E-04 | 0.006859439177 | 1.000000227 | 0 | 0.2182966672 | 0 | 0.001337552035 | 0 |
| TARA_SAMEA2622696_METAG_BAOOJBHG | 5 | -0.0150214479 | PCC-6307 | Prochlorococcus_A | | 0 | 0 | 0.02982728491 | 8.64E-05 | 0.2862733519 | 1.00000032 | 0 | 0.06981793149 | 0 | 8.46E-04 | 0.03316111833 |
| TARA_SAMEA2622696_METAG_HCJFGCBE | 5 | -0.07214716947 | Flavobacteriales | GCA-2716065 | GCA-2716065 sp | 0 | 1.65E-04 | 0.05119438175 | 3.77E-05 | 0.01987027772 | 1.000000006 | 0 | 0.05489592997 | 0 | 1.21E-05 | 0.004099465806 |
| TARA_SAMEA2622696_METAG_KGIODGLG | 2 | 0.05109134139 | SCGC-AAA003-L | GCA-2717625 | | 4.02E-08 | 0 | 0.02855517594 | 0 | 0.03333333668 | 1.20E-04 | 0 | 0.02962211974 | 0 | 0.2804930177 | 0.04195908132 |
| TARA_SAMEA2622696_METAG_LJCKGMCC | 1 | -0.07455011316 | UBA9615 | UBA6601 | UBA6601 sp002 | 0 | 0 | 0.9853302006 | 1.24E-06 | 0.03277466873 | 0.9833498412 | 0 | 0.04396095609 | 0 | 2.63E-04 | 0.04328475591 |
| TARA_SAMEA2622710_METAG_DBGJLHNK | 4 | 0.05137146243 | TMED109 | TMED109 | TMED109 sp002 | 0.3111141712 | 0 | 1.000000202 | 1.28E-04 | 0.07340729199 | 0 | 0 | 1.000000202 | 0 | 0.007239044968 | 0 |
| TARA_SAMEA2622710_METAG_KOHPLGIJ | 2 | -0.4143960973 | Flavobacteriales | UBA974 | UBA974 sp9025 | 4.93E-06 | 0 | 0.05130174589 | 1.000000247 | 0.07090468165 | 0 | 0 | 1.000000247 | 0 | 0 | 0 |
| TARA_SAMEA2622715_METAG_BAPMEKGD | 4 | -0.02860414609 | Pseudomonadale | UBA9145 | UBA9145 sp002 | 0 | 0 | 0.999999683 | 1.02E-04 | 0.001574722783 | 4.44E-06 | 0 | 0.0586451108 | 0 | 0.00381762504 | 0 |
| TARA_SAMEA2622716_METAG_PGFFMCOD | 2 | -0.01127963873 | Flavobacteriales | TMED113 | TMED113 sp002 | 0 | 0 | 0.1273796776 | 1.58E-04 | 0.1281250863 | 0 | 0 | 0.0445316169 | 0 | 7.14E-04 | 0 |
| TARA_SAMEA2622725_METAG_ALFDIHAE | 5 | -0.2346441576 | Nitrosococcales | Methylophaga | Methylophaga sp | 3.52E-06 | 3.62E-04 | 0.007494663185 | 7.22E-05 | 0 | 1 | 0 | 0.0401132061 | 3.24E-04 | 0.01518971843 | 0 |
| TARA_SAMEA2622733_METAG_CKLGIHPC | 3 | -0.00232098659 | Pelagibacterales | AG-414-E02 | | 6.65E-06 | 0 | 0.01666666667 | 0.9999996976 | 0.9999996976 | 0.9999996976 | 0 | 0.05929351096 | 0 | 0.9999996976 | 0 |
| TARA_SAMEA2622733_METAG_DHIHELHE | 3 | -0.05553428143 | Pseudomonadale | UBA9145 | UBA9145 sp002 | 0 | 0 | 0.983647256 | 3.01E-06 | 0.2357310104 | 0.8337196919 | 0 | 0.01193140633 | 0 | 0.9999999949 | 0.9833333333 |
| TARA_SAMEA2622733_METAG_EELCBJME | 6 | -0.03469111884 | Burkholderiales | GCA-2721545 | GCA-2721545 sp | 0 | 0 | 0.250804465 | 1.33E-05 | 0.9999997927 | 0.9999997927 | 0 | 0.05950750679 | 0 | 7.61E-04 | 0.001060805636 |
| TARA_SAMEA2622733_METAG_OECEHKPE | 1 | -0.0652412557 | Marinisomatales | GCA-2715465 | | 1.32E-05 | 0 | 0.9999997392 | 0.001919062191 | 0.03614039784 | 0.9834137718 | 2.21E-04 | 0.05054756421 | 0.01071019082 | 2.62E-05 | 0 |
| TARA_SAMEA2622736_METAG_DIOOGNDP | 2 | -0.5825431509 | Pseudomonadale | Pseudomonas_E | Pseudomonas_E | 0 | 0 | 0.15149299 | 1.13E-04 | 0.01513145634 | 0 | 0 | 0.02441492598 | 0 | 0.001950797435 | 0 |
| TARA_SAMEA2622737_METAG_BCPAJEON | 4 | -0.1016322702 | Rhodospirillales_ | TMED8 | TMED8 sp00272 | 0 | 2.88E-05 | 0.9837237199 | 3.62E-06 | 0.03213076022 | 2.46E-07 | 0 | 0.9834350938 | 0 | 0.03943096964 | 0 |
| TARA_SAMEA2622737_METAG_HMOCFPAL | 4 | -0.03236187161 | Burkholderiales | GCA-2721545 | | 0.2720609608 | 0.01393685592 | 1 | 0 | 0.01942553889 | 0.002517859767 | 0 | 0.06609046148 | 0 | 1.50E-06 | 0 |
| TARA_SAMEA2622737_METAG_HPGMPLOF | 3 | -0.1581469075 | Pirellulales | UBA721 | UBA721 sp0023 | 3.32E-06 | 0 | 0.03831110346 | 0 | 1 | 1 | 0 | 1 | 0 | 1 | 0 |
| TARA_SAMEA2622737_METAG_NMILOMPH | 3 | -0.04620593247 | UBA1151 | GCA-2717385 | GCA-2717385 sp | 0 | 0 | 0.1170908831 | 0.9999998205 | 0 | 0.9999998205 | 0 | 0.05492893775 | 0 | 0.9999998205 | 0 |
| TARA_SAMEA2622737_METAG_OHMFMGLP | 1 | -0.00524623337 | Marinisomatales | UBA2134 | UBA2134 sp002 | 1.37E-06 | 0 | 1 | 2.52E-04 | 1 | 1 | 0.009862059881 | 1 | 0.009180812315 | 0.001065218885 | 0 |
| TARA_SAMEA2622737_METAG_PCAKDJOK | 3 | 0.009623207419 | Pelagibacterales | MED-G40 | | 0.003073202242 | 0 | 0.01499176735 | 8.22E-05 | 1.000000321 | 0.9833337922 | 2.30E-05 | 0.9839311397 | 0.001341268314 | 1.000000321 | 0.03612292811 |
| TARA_SAMEA2622738_METAG_PMOIGPEK | 4 | -0.08097740395 | SAR202 | GCA-2719675 | GCA-2719675 sp | 0 | 0 | 1.000000298 | 1.91E-05 | 1.000000298 | 0 | 0 | 1.000000298 | 0 | 0.008705488078 | 0.06724745004 |
| TARA_SAMEA2622764_METAG_EHKPLJDK | 1 | -0.00704056049 | Marinisomatales | Marinisoma | Marinisoma sp90 | 0 | 0 | 0.9999996899 | 1.41E-04 | 0.002359980289 | 0.9999996899 | 0 | 0.9843672229 | 0 | 0 | 0 |
| TARA_SAMEA2622765_METAG_CGEGLBBL | 4 | -0.05052916892 | Pirellulales | UBA7805 | UBA7805 sp002 | 0 | 0 | 0.9999997435 | 0.1622833471 | 0 | 0.009752701195 | 0 | 0.02466548845 | 0 | 7.77E-04 | 0 |
| TARA_SAMEA2622765_METAG_CKKHFEEO | 2 | -0.1705947183 | Phycisphaerales | UBA7800 | UBA7800 sp002 | 0 | 0 | 0.08712284392 | 6.19E-05 | 0.03495882146 | 0.007776328006 | 2.91E-05 | 0.0748282833 | 0 | 0 | 0 |
| TARA_SAMEA2622765_METAG_ICPMIAHH | 4 | -0.06896710173 | Opitutales | MB11C04 | MB11C04 sp002 | 0 | 0 | 1.000000003 | 0.003456551516 | 0.03188028807 | 0.00200908234 | 0 | 0.0661148732 | 0 | 3.90E-05 | 0.02896186923 |
| TARA_SAMEA2622765_METAG_LCCLODIH | 2 | -0.00370145605 | Verrucomicrobial | Roseibacillus_B | Roseibacillus_B | 0 | 0 | 0.174530282 | 1.52E-04 | 0.003357472687 | 4.48E-04 | 0 | 0.005125043649 | 0 | 0.001589901725 | 0.03722448131 |
| TARA_SAMEA2622765_METAG_MHBAPCGP | 8 | -0.09649447201 | Rhizobiales | Stappia | Stappia sp00272 | 0.07907462515 | 0 | 1.000000175 | 1.000000175 | 0.002568343545 | 1.000000175 | 0 | 0.04783574029 | 0 | 8.09E-04 | 0 |
| TARA_SAMEA2622765_METAG_OGNPIJOJ | 7 | -0.1420296345 | Flavobacteriales | Muricauda_A | Muricauda_A luta | 0 | 0 | 0.03801589238 | 2.02E-04 | 1.000000317 | 0.0167444835 | 0 | 0.03609580637 | 0 | 1.000000317 | 0.006303642654 |
| TARA_SAMEA2622766_METAG_APIKJLNH | 1 | -8.72E-04 | MED-G09 | MED-G09 | MED-G09 sp003 | 0 | 0 | 0.9999997687 | 2.34E-04 | 0.9999997687 | 0.9999997687 | 0 | 0.07091043147 | 0 | 0.004322994253 | 0 |
| TARA_SAMEA2622766_METAG_DIHIBLAE | 6 | -0.1173204082 | UBA8366 | UBA8366 | UBA8366 sp002 | 0 | 0 | 0.3200503064 | 7.57E-05 | 1.000000195 | 1.000000195 | 0 | 0.04476175156 | 0.06550552541 | 7.47E-04 | 0 |
| TARA_SAMEA2622766_METAG_FDFJEPOA | 3 | 0.009584899006 | UBA1151 | GCA-002708145 | GCA-002708145 | 6.46E-04 | 0.002124335737 | 0.09493113804 | 2.89E-05 | 0.06619892086 | 0.999999833 | 0 | 0.06956487465 | 0 | 0.8999998358 | 0.03396106798 |
| TARA_SAMEA2622766_METAG_HMICCCOP | 1 | -0.3703166739 | Rhodospirillales_ | Thalassospira | Thalassospira sp | 0 | 0 | 0.9999997203 | 4.64E-05 | 0.111873605 | 0.9999997203 | 0 | 0.03524711264 | 5.93E-04 | 0.02454696646 | 0 |
| TARA_SAMEA2622766_METAG_ICABGDLO | 2 | 0.04261063856 | Pelagibacterales | Pelagibacter | Pelagibacter sp9 | 0 | 0 | 0.5230217861 | 8.19E-05 | 0.01426222704 | 0.0190362554 | 0 | 0.04635075046 | 0 | 0.005669214392 | 0 |
| TARA_SAMEA2622796_METAG_BOADKHEC | 3 | -0.05264092166 | UBA2963 | UBA2963 | UBA2963 sp003 | 0 | 0 | 0.01834836774 | 1.45E-04 | 0.003681220983 | 1.000000317 | 0 | 1.000000317 | 0 | 1.000000317 | 0 |
| TARA_SAMEA2622796_METAG_EAKHFFIO | 8 | -0.2052397408 | UBA1151 | UBA1328 | UBA1328 sp002 | 0 | 0 | 0.9999998816 | 0.9999998816 | 0.9999998816 | 0.9999998816 | 0 | 0.9999998816 | 0 | 0.01172298931 | 0.03739396763 |
| TARA_SAMEA2622796_METAG_FBJMAKDD | 2 | 0.01476103763 | Marinisomatales | Marinisoma | Marinisoma sp00 | 0 | 0 | 0.02251775372 | 6.94E-05 | 0.007422537056 | 0.003649225297 | 0 | 0.05151114661 | 0 | 0 | 4.92E-06 |
| TARA_SAMEA2622796_METAG_NAJLOALF | 2 | 0.07753513072 | SCGC-AAA003-L | TMED6 | | 0 | 0 | 0.05340358213 | 1.000000002 | 0.001817254353 | 0.02918060909 | 0 | 0.04193218037 | 0 | 1.18E-04 | 1.93E-06 |
| TARA_SAMEA2622796_METAG_NEBCDNLH | 1 | 0.01929225602 | Acidimicrobiales | UBA9410 | UBA9410 sp002 | 0 | 0 | 0.9999996851 | 7.62E-05 | 7.93E-04 | 0.9999996851 | 0 | 0.09025396412 | 0 | 0.01957111962 | 0 |
| TARA_SAMEA2622796_METAG_NPJHAAOM | 5 | -0.2614201715 | Caulobacterales | Ponticaulis | Ponticaulis sp002 | 1.34E-06 | 0 | 0.01666505403 | 7.17E-05 | 0.113397297 | 0.9999998881 | 0 | 0.4595296164 | 0.001195035422 | 0.002408354718 | 0.004588494202 |
| TARA_SAMEA2622800_METAG_LIMMEGDN | 7 | 0.03979473386 | UBA1144 | GCA-002715585 | GCA-002715585 | 0 | 0 | 0.9046014143 | 3.87E-05 | 0.001639159009 | 4.24E-06 | 0 | 0.03857820649 | 0 | 0.9875746425 | 0.08032686789 |
| TARA_SAMEA2622801_METAG_ECAHMIGO | 6 | -0.05548169487 | MPNO01 | UBA2964 | UBA2964 sp002 | 0 | 2.14E-05 | 0.0278333661 | 0.1663283792 | 0.9500488205 | 0.01666667026 | 0 | 0.05363839917 | 0 | 0 | 0.03787521111 |
| TARA_SAMEA2622801_METAG_EMDBLGIB | 4 | -0.01320760149 | UBA8366 | UBA6544 | UBA6544 sp002 | 0 | 0 | 0.9999998895 | 0 | 0.007428550831 | 0.002020497817 | 0 | 0.0351344673 | 0 | 9.91E-04 | 0.001159083854 |
| TARA_SAMEA2622801_METAG_GKAHAABC | 2 | -0.2111357485 | Pseudomonadale | UBA6940 | UBA6940 sp002 | 0 | 0 | 0.04369715427 | 1.02E-04 | 0.003044830861 | 0 | 0 | 0.03572780561 | 0 | 0.02163211841 | 0 |
| TARA_SAMEA2622801_METAG_GLDMIMJA | 5 | -0.00664411696 | Pseudomonadale | GCA-002728675 | GCA-002728675 | 0 | 0 | 0.03826819914 | 0.01666667164 | 0.04135231747 | 1.000000307 | 0 | 0.0532742027 | 0 | 9.44E-04 | 0.02668362803 |
| TARA_SAMEA2622801_METAG_LEMAIJLG | 6 | 0.0210129358 | Pedosphaerales | UBA1096 | UBA1096 sp002 | 0 | 0 | 5.32E-04 | 1.78E-06 | 0.9667203329 | 1.54E-05 | 0 | 0.07469866424 | 0 | 0.009898781622 | 0 |
| TARA_SAMEA2622817_METAG_ANKMHOAM | 6 | 0.008044922918 | Phycisphaerales | UBA854 | UBA854 sp0022 | 0 | 0 | 0.05524398794 | 6.92E-05 | 1.000000191 | 0.01115142913 | 0 | 0.09673251999 | 0 | 7.82E-04 | 0.06846793175 |
| TARA_SAMEA2622822_METAG_DAPOALAA | 7 | -0.04385732198 | Marinisomatales | TMED80 | TMED80 sp9130 | 1.53E-06 | 0 | 0.05165929008 | 0.9833337085 | 0.9833337085 | 0.01666665885 | 0 | 0.02082756986 | 0 | 1.000000367 | 0 |
| TARA_SAMEA2622822_METAG_GPEDNHIC | 2 | 0.001947141436 | Acidimicrobiales | UBA8592 | | 0 | 0 | 0.007044364372 | 0.01678836239 | 0.03652887239 | 0.2863962892 | 0 | 0.04715867602 | 0 | 0.008434715135 | 0 |
| TARA_SAMEA2622822_METAG_IEDPHKED | 3 | -0.02938338743 | GCA-2717565 | GCA-2717565 | GCA-2717565 sp | 0 | 0 | 0.04812958047 | 1 | 0.01233627575 | 1 | 0.001835316616 | 0.08841877372 | 0 | 1 | 0.100000505 |
| TARA_SAMEA2622822_METAG_KGNHGHNB | 2 | 0.004771867862 | SAR86 | AEGEAN-183 | | 0.9666667252 | 5.83E-04 | 0.04010709955 | 0.03335343901 | 0.07173028316 | 0.03333333333 | 0 | 1 | 0 | 0.004012054926 | 0 |
| TARA_SAMEA2622822_METAG_OGCALNKE | 3 | 0.05564184435 | GCA-002720895 | GCA-002720895 | GCA-002720895 | 0.01407782445 | 0 | 0.9999999978 | 9.93E-07 | 0.9999999978 | 0.9197881027 | 0 | 0.09017656593 | 0 | 0.9999999978 | 2.44E-04 |
| TARA_SAMEA2622822_METAG_PPEPLHJD | 5 | -0.06760867308 | UBA9160 | UBA6930 | UBA6930 sp002 | 0 | 0 | 0.0434105522 | 0.0166974004 | 0.003185966731 | 0.9666667735 | 0 | 0.04791747213 | 0 | 0.009024482563 | 0 |
| TARA_SAMEA2622823_METAG_BGCLLDIO | 1 | 0.003956896633 | Arenicellales | UBA868 | UBA868 sp0023 | 4.38E-06 | 0 | 0.999999687 | 1.26E-04 | 0.006875553729 | 0.999999687 | 0 | 0.999999687 | 0 | 0.01437012636 | 0 |
| TARA_SAMEA2622823_METAG_BHGDDDOD | 2 | -0.04501502417 | Arenicellales | GCA-2721105 | GCA-2721105 sp | 0 | 0 | 0.03237831256 | 0.0167144382 | 0.3071272921 | 0.004900256655 | 0 | 0.07650945492 | 0 | 0 | 0.01993432492 |
| TARA_SAMEA2622823_METAG_CFMOJEHO | 2 | -0.00759092670 | UBA2968 | UBA6620 | UBA6620 sp002 | 8.79E-04 | 0 | 0.02165060685 | 0.01666666667 | 0.09404831764 | 0 | 0 | 0.9841080762 | 0 | 4.40E-05 | 0 |
| TARA_SAMEA2622823_METAG_GOFKMFMO | 8 | -0.1365319664 | Opitutales | UBA2993 | UBA2993 sp002 | 0.2475645572 | 0 | 1 | 1 | 1 | 1 | 0.001988980203 | 1 | 0 | 1 | 0 |
| TARA_SAMEA2622823_METAG_HHHENNDP | 7 | -0.2375523872 | UBA1280 | UBA12223 | UBA12223 sp00 | 0 | 6.40E-05 | 0.03333332782 | 2.65E-06 | 0.1202384145 | 0.03333332782 | 0.001517882476 | 0.06060645201 | 1.16E-05 | 0.9999997887 | 0.05290520418 |
| TARA_SAMEA2622823_METAG_KADFMIEB | 3 | -0.04973027716 | Verrucomicrobial | EC70 | EC70 sp0027088 | 0 | 0 | 0.934586953 | 5.09E-06 | 9.17E-04 | 0.9833330529 | 0.002633839479 | 0.06340013673 | 0 | 0.966815286 | 0.001366144429 |
| TARA_SAMEA2622823_METAG_MLHCOEJP | 3 | -0.03933175296 | Woeseiales | SZUA-117 | SZUA-117 sp913 | 2.26E-06 | 0 | 0.0303579873 | 1 | 1 | 1 | 0 | 0.02782518889 | 0 | 1 | 0 |
| TARA_SAMEA2622837_METAG_AFJNLKBG | 2 | 0.006628420876 | MED-G09 | | | 0 | 0.01123022657 | 0.03378767119 | 4.85E-05 | 0.006327118846 | 0 | 0 | 0.05680928926 | 0 | 0.01720527898 | 0 |
| TARA_SAMEA2622837_METAG_BCCMNEFM | 3 | -0.06188935249 | Anaerolineales | UBA11657 | UBA11657 sp002 | 0 | 0 | 0.9999995905 | 9.83E-05 | 0.2570161792 | 0.9999995905 | 0 | 0.03642479432 | 0 | 0.9999995905 | 0 |
| TARA_SAMEA2622837_METAG_BEOLKKCE | 2 | -0.00756144663 | UBA1127 | GCA-2717925 | GCA-2717925 sp | 0 | 0 | 0.02316735132 | 1.30E-04 | 0.1146366741 | 0.005511144086 | 0 | 0.04029900888 | 0 | 0.00267100073 | 0 |
| TARA_SAMEA2622837_METAG_CJEHOFFG | 4 | -0.05264930731 | Pirellulales | UBA9264 | UBA9264 sp002 | 0.009094741805 | 0 | 0.9839620772 | 0.9999999937 | 1.63E-04 | 0.01666666036 | 0 | 0.00256374066 | 0 | 8.54E-05 | 0 |
| TARA_SAMEA2622837_METAG_FJHMIHLF | 1 | -0.06208958335 | Vicinamibacterales | | | 2.68E-06 | 0 | 0.9839820928 | 9.64E-05 | 0.9866819212 | 0.9833333333 | 2.20E-04 | 0.04858805225 | 0.01666666667 | 0.0227332003 | 0 |
| TARA_SAMEA2622837_METAG_HBAPHKBN | 4 | -0.1030697027 | KS3-K002 | CAJWLK01 | CAJWLK01 sp91 | 0.2086207965 | 0 | 1 | 0.01666666667 | 0.1971870716 | 0.006089588025 | 0 | 0.04495519587 | 0 | 0.001016325733 | 0 |
| TARA_SAMEA2622837_METAG_INCIHHJP | 3 | -0.06963414027 | Marinisomatales | TCS55 | TCS55 sp001577 | 0 | 0 | 0.07019363651 | 1.000000004 | 0.02022917835 | 0.9834060097 | 0.002631959807 | 0.9970219437 | 0 | 1.000000004 | 0.07268348145 |
| TARA_SAMEA2622837_METAG_JMDHKPDK | 6 | -0.0684049743 | Opitutales | JABJCN01 | JABJCN01 sp913 | 0 | 0 | 0.02292458523 | 1.19E-04 | 1 | 1 | 0 | 0.04819531482 | 0 | 7.45E-04 | 0.001468880563 |
| TARA_SAMEA2622837_METAG_LKADBGOG | 3 | -0.01031217461 | Thalassobaculale | GCA-2722775 | GCA-2722775 sp | 3.13E-06 | 0 | 0.02808614808 | 1.26E-04 | 5.48E-06 | 1 | 0.01497223101 | 1 | 0 | 1 | 0 |

| | | | | | | | | | | | | | | | | |
|---|---|---|---|---|---|---|---|---|---|---|---|---|---|---|---|---|
| TARA_SAMEA2622837_METAG_MANGALKA | 3 | -0.05431564148 | UBA2985 | VFHP01 | VFHP01 sp0093 | 5.86E-08 | 0 | 0.01392161783 | 7.28E-06 | 1.000000004 | 1.000000004 | 0 | 0.04775036418 | 0 | 1.000000004 | 0 |
| TARA_SAMEA2622837_METAG_MEOGPOJG | 3 | -0.032356832 | UBA1151 | GCA-2717385 | GCA-2717385 sp | 4.31E-06 | 0 | 1.00000024 | 1.32E-04 | 1.00000024 | 1.00000024 | 0 | 0 | 0.001452302295 | 1.00000024 | 0 |
| TARA_SAMEA2622837_METAG_NAOINJEI | 1 | -0.0445865011 | Vicinamibacterale | DTWD01 | | 0 | 0 | 0.999999773 | 8.17E-06 | 0.01162539183 | 0.999999773 | 0.001349108338 | 0.01630777828 | 0 | 0.01028127683 | 0 |
| TARA_SAMEA2622837_METAG_POOBDMOF | 3 | 0.07619347892 | Acidimicrobiales | S20-B6 | S20-B6 sp00234 | 6.50E-08 | 0 | 0.3213601103 | 1.40E-04 | 0.1906144884 | 0.9999997363 | 0 | 0.9841426597 | 0 | 0.9999997363 | 0 |
| TARA_SAMEA2622843_METAG_AHKNABLD | 4 | -0.1395521333 | Flavobacteriales | GCA-002722245 | GCA-002722245 | 4.98E-04 | 0 | 0.9999999961 | 0.01678993959 | 0.01738944651 | 0.0128835284 | 1.16E-04 | 0.19958158 | 0 | 0.004140505348 | 0.01600580739 |
| TARA_SAMEA2622843_METAG_BGMDGEHN | 4 | -0.1877390601 | Flavobacteriales | Winogradskyella | Winogradskyella | 0 | 0 | 1 | 0.01666666667 | 0 | 0 | 0 | 1 | 0 | 6.99E-05 | 0.9833333333 |
| TARA_SAMEA2622843_METAG_KHMHMHPH | 7 | -0.03040858702 | TMED127 | GCA-2715305 | GCA-2715305 sp | 0 | 0 | 1 | 1.51E-04 | 0.007022136901 | 0 | 0 | 1 | 0 | 1 | 0 |
| TARA_SAMEA2622843_METAG_NCDLFJCE | 5 | -0.081891773 | Phycisphaerales | UBA1668 | UBA1668 sp002 | 1.43E-04 | 0 | 0.01369701796 | 1.42E-06 | 0 | 0.9999998298 | 0 | 0.9846148151 | 0 | 1.40E-05 | 2.56E-04 |
| TARA_SAMEA2622901_METAG_OJIJLFFE | 2 | -0.02138518132 | Flavobacteriales | GCA-002727735 | | 8.60E-07 | 0 | 0.04434711015 | 9.12E-05 | 0.005899766414 | 6.20E-06 | 0 | 0.08075258278 | 0 | 8.08E-04 | 0 |
| TARA_SAMEA2622923_METAG_DLBMCHKI | 2 | -0.0151063394 | Pedosphaerales | UBA1100 | UBA1100 sp002 | 0.1172729849 | 0 | 0.05864779389 | 8.48E-05 | 0.1051474316 | 0.001167589555 | 4.07E-04 | 0.03853355619 | 8.50E-07 | 7.37E-04 | 4.78E-06 |
| TARA_SAMEA2622923_METAG_DNGLHAAI | 8 | -0.1644465205 | Desulfobacterale | Desulfobacter | Desulfobacter sp | 0 | 0 | 1 | 1 | 7.32E-04 | 1 | 0 | 1 | 0 | 0 | 0 |
| TARA_SAMEA2622923_METAG_JAPLMJLG | 8 | -0.07841603604 | Thalassobaculale | GCA-002701455 | GCA-002701455 | 5.49E-08 | 0 | 0.07045613001 | 0.9666677194 | 3.82E-04 | 0.933343262 | 0 | 0.08149652296 | 0 | 8.06E-04 | 0 |
| TARA_SAMEA2623059_METAG_FJBFNHMP | 4 | -0.02508218347 | TMED109_A | GCA-2732015 | | 3.92E-08 | 0 | 1.000000007 | 0.6722220742 | 0.9834272324 | 0 | 0 | 0.9999499453 | 0 | 1.34E-05 | 0 |
| TARA_SAMEA2623079_METAG_CJOCBLOE | 1 | -0.00719397247 | Verrucomicrobial | EC70 | EC70 sp9025916 | 4.94E-06 | 0 | 1 | 1.24E-04 | 1 | 1 | 0 | 0.05934474676 | 0 | 0.01371892209 | 0 |
| TARA_SAMEA2623079_METAG_NJMDAEIA | 7 | -0.00106935809 | Pelagibacterales | | | 0 | 0 | 0.9932650954 | 1 | 0.001726845345 | 0 | 0 | 1 | 0 | 1 | 0 |
| TARA_SAMEA2623098_METAG_BFNDOAOF | 3 | -0.02757895901 | Woeseiales | PBBK01 | PBBK01 sp0027 | 0 | 0.01212898815 | 0.9840577579 | 1.82E-04 | 0.1359556315 | 0.9834560009 | 0 | 0.06955242463 | 0 | 1 | 0 |
| TARA_SAMEA2623098_METAG_HGPGGPGN | 2 | -0.057991083 | Pseudomonadale | UBA9145 | UBA9145 sp002 | 4.47E-06 | 0 | 0.03991504329 | 1.000000009 | 0 | 0 | 0 | 0.9666858289 | 0 | 2.58E-07 | 0 |
| TARA_SAMEA2623116_METAG_BHDMLDOE | 2 | -0.04803419114 | Flavobacteriales | MED-G21 | | 0.002709692104 | 0 | 0.2015436387 | 7.35E-05 | 0.0323779627 | 0 | 0 | 0.06092317188 | 0 | 0 | 0 |
| TARA_SAMEA2623116_METAG_CBIBJFOP | 7 | -0.00223155002 | Pelagibacterales | | | 0 | 0 | 1.000000297 | 0.01666666667 | 0.06756052524 | 8.50E-04 | 0.9833336305 | 1.000000297 | 0 | 1.000000297 | 0 |
| TARA_SAMEA2623116_METAG_HDAFLMCO | 4 | -0.1735383661 | Opitutales | GCA-2690565 | GCA-2690565 sp | 0 | 0 | 1 | 1.17E-04 | 1 | 0.002746452151 | 0 | 0.1033231381 | 0 | 0.001611598225 | 0 |
| TARA_SAMEA2623116_METAG_HIAOIBEN | 7 | 0 | Flavobacteriales | MAG-121220-bin8 | | 0 | 0 | 0.02661953188 | 1.35E-04 | 0.04313840094 | 6.14E-06 | 0 | 0.08174590067 | 0 | 1 | 0 |
| TARA_SAMEA2623116_METAG_LCHCHCLK | 2 | -0.00480707685 | Flavobacteriales | GCA-2697505 | | 0.00349570418 | 0 | 0.111312637 | 1.17E-04 | 0.0092970084 | 0 | 0 | 0.2772440734 | 0 | 1.49E-05 | 0 |
| TARA_SAMEA2623116_METAG_MOMOJFAF | 2 | -0.02632540482 | Nanopelagicales | S36-B12 | S36-B12 sp0027 | 0 | 0 | 0.05295289082 | 1.24E-06 | 0.005388465141 | 0.2499999964 | 0.07866030593 | 0.9222878309 | 0 | 1.23E-05 | 0.01646837889 |
| TARA_SAMEA2623135_METAG_BLHAGEJF | 7 | -0.0336020006 | Flavobacteriales | Marivariicella | Marivariicella sp | 0 | 0 | 1 | 1.32E-04 | 4.02E-04 | 0.3333333333 | 0 | 0.7704558281 | 0 | 1 | 0 |
| TARA_SAMEA2623135_METAG_DJILNNCH | 2 | -0.5933801151 | Enterobacterales | Alteromonas | Alteromonas aus | 0.04342441081 | 0 | 0.05975166759 | 1 | 0.001235896066 | 0 | 0 | 0.0579748299 | 0 | 0.002411856105 | 0 |
| TARA_SAMEA2623135_METAG_ILGMCAFG | 4 | -0.06573403974 | Acidimicrobiales | UBA11606 | UBA11606 sp002 | 9.32E-04 | 0 | 0.9999999943 | 1.50E-04 | 0.1715147724 | 0.01666666098 | 0.03241786092 | 0.06022616223 | 0 | 1.94E-04 | 3.15E-04 |
| TARA_SAMEA2623135_METAG_KCHDOGHJ | 7 | -0.07438323354 | Flavobacteriales | MS024-2A | | 0 | 0 | 0.03312057176 | 9.15E-07 | 0.00500977982 | 0.03333333047 | 0 | 0.2151907209 | 0 | 1.000000179 | 0.01458372255 |
| TARA_SAMEA2623155_METAG_MLJGDHKJ | 4 | -0.01853675942 | Vicinamibacterale | UBA823 | UBA823 sp0022 | 0 | 0 | 0.9999997703 | 7.72E-05 | 0.00656806603 | 0.00240133714 | 0 | 0.09986134644 | 0 | 0.004572006226 | 6.60E-04 |
| TARA_SAMEA2623275_METAG_KPGDNKKP | 5 | -0.0636992044 | Flavobacteriales | SP287 | | 7.66E-08 | 0 | 0.02801714592 | 0 | 0.01680487379 | 0.966703765 | 0 | 0.9835575534 | 0 | 0.0384026278 | 0 |
| TARA_SAMEA2623295_METAG_JONIEMEH | 2 | -0.03445419071 | Verrucomicrobial | Roseibacillus_B | Roseibacillus_B | 0 | 0 | 0.01740036563 | 0.05000246224 | 0.1244155822 | 0.05000009174 | 0 | 0.919313111 | 0 | 1.03E-04 | 0 |
| TARA_SAMEA2623295_METAG_MNECEEFO | 7 | -0.04520630035 | UBA8231 | UBA8231 | UBA8231 sp002 | 0 | 0 | 0.0235346532 | 1.89E-04 | 0 | 0.005248253798 | 0 | 0.05557423806 | 0 | 1.000000204 | 0 |
| TARA_SAMEA2623295_METAG_PHBBDDEJ | 6 | -0.03195329012 | Acidimicrobiales | UBA3125 | UBA3125 sp002 | 0 | 0.001098164612 | 0.01103911926 | 7.60E-05 | 1 | 0.00434848487 | 0 | 0.06633493736 | 0 | 0.002128406412 | 0 |
| TARA_SAMEA2623314_METAG_ALHGJMCB | 3 | 0.01664903441 | Pedosphaerales | UBA1096 | UBA1096 sp913 | 0 | 0 | 0.02911279079 | 3.07E-05 | 0.9654461325 | 0.9833333334 | 0 | 0 | 0 | 0.9833333322 | 0.0621579015 |
| TARA_SAMEA2623314_METAG_BIIFJDAP | 4 | -0.04108753715 | SAR202 | UBA2160 | | 0 | 0 | 1.000000251 | 0.01209938537 | 0.9164809502 | 3.18E-04 | 0 | 0.9204026915 | 0 | 7.10E-05 | 0.002472128451 |
| TARA_SAMEA2623314_METAG_DGDHOOEI | 6 | -0.0549668492 | Marinisomatales | UBA2126 | UBA2126 sp002 | 0 | 0 | 0.01549793082 | 3.74E-05 | 0.9888888889 | 0.9833333333 | 0 | 0.06137108428 | 0 | 3.27E-05 | 0 |
| TARA_SAMEA2623314_METAG_HLLPALBA | 7 | -0.04394273965 | Phycisphaerales | UBA12567 | UBA12567 sp00 | 0 | 0 | 1.000000181 | 0.4916728676 | 0.001521229292 | 2.38E-04 | 0 | 0.06387499313 | 0 | 1.000000181 | 0.03156565085 |
| TARA_SAMEA2623370_METAG_KLNICJAN | 8 | -0.06553157046 | Rhodobacterales | CACKLF01 | CACKLF01 sp00 | 0 | 0 | 1.000000198 | 1.000000198 | 0 | 1.000000198 | 0 | 1.000000198 | 0 | 0 | 0.1325275523 |
| TARA_SAMEA2623390_METAG_BKPADPMA | 3 | -0.02861121825 | UBA890 | UBA890 | UBA890 sp0027 | 0 | 0 | 0.03326764776 | 1.31E-04 | 0.9555783607 | 1.000000001 | 0 | 0.07581676776 | 0 | 1.000000001 | 0.3064314798 |
| TARA_SAMEA2623390_METAG_IFCMLOCO | 2 | -0.07464171287 | Vicinamibacterale | UBA11600 | UBA11600 sp002 | 0 | 0 | 0.04505881731 | 1.000000011 | 0.003927445645 | 0.02802659753 | 0 | 1.000000011 | 0 | 7.88E-04 | 0 |
| TARA_SAMEA2623390_METAG_KCIBEPJC | 7 | -0.0255810536 | Dehalococcoidale | GCA-2685815 | GCA-2685815 sp | 0 | 0 | 0.02817973328 | 1.54E-04 | 0.1842318242 | 0.00562910812 | 0 | 0 | 0 | 0.9999997243 | 0 |
| TARA_SAMEA2623390_METAG_LNAPLDHH | 7 | -0.04854292921 | SAR202 | UBA11996 | UBA11996 sp002 | 0 | 0 | 0.01640018612 | 2.87E-06 | 9.66E-05 | 0 | 0 | 0.9884315324 | 0 | 0.9999999966 | 0.2522424548 |
| TARA_SAMEA2623390_METAG_NACNCLMF | 4 | -0.06890120941 | Pseudomonadale | GCA-002726655 | GCA-002726655 | 0 | 0 | 0.967364829 | 0.03333332797 | 0.967003944 | 0.003154779923 | 0 | 0.9676845874 | 0 | 0.0013943521 | 0.0353989261 |
| TARA_SAMEA2623403_METAG_IGDKFJNA | 2 | -0.170800329 | Flavobacteriales | Salegentibacter | Salegentibacter | 0 | 0 | 0.09383101441 | 9.20E-07 | 0.03318219137 | 0.01666666667 | 0 | 0.9679479312 | 0 | 2.47E-05 | 3.44E-05 |
| TARA_SAMEA2623426_METAG_CDDIFCLD | 7 | -0.01326978296 | Opitutales | MB11C04 | | 0 | 0 | 0.04839279239 | 1.28E-04 | 0.1185932492 | 0.04913816297 | 0.1501207069 | 0.07914989979 | 1.72E-04 | 1 | 0.05007581651 |
| TARA_SAMEA2623446_METAG_EANBFFNL | 2 | -0.1169419955 | Rhodobacterales | LFER01 | LFER01 sp0015 | 0 | 0 | 0.1918173063 | 1.10E-04 | 0.001538484169 | 0.001312211047 | 0 | 0.06541528545 | 0 | 0.02198353701 | 6.28E-04 |
| TARA_SAMEA2623463_METAG_BLGGBFLI | 8 | 0.07231152512 | SCGC-AAA003-L | TMED161 | TMED161 sp002 | 0 | 0 | 1.0000003 | 1.0000003 | 0.0220956946 | 1.0000003 | 0 | 0.07495806277 | 1.0000003 | 1.0000003 | 0 |
| TARA_SAMEA2623463_METAG_PPLLMEMB | 4 | -0.09122929791 | PCC-6307 | RCC307 | RCC307 sp9130 | 1.95E-06 | 0.001583266884 | 1 | 9.00E-05 | 1 | 9.37E-05 | 0 | 0.0835697365 | 0 | 0.3352911413 | 0 |
| TARA_SAMEA2623513_METAG_FFOGFDIA | 6 | -0.3021970539 | Pseudomonadale | Oleibacter | Oleibacter sp002 | 0 | 0 | 0.05040914188 | 4.18E-05 | 1.000000337 | 1.000000337 | 0 | 1.000000337 | 0 | 0.006873759134 | 0 |
| TARA_SAMEA2623627_METAG_JBIOOEFI | 5 | 0.01697732915 | UBA1151 | TMED-70 | TMED-70 sp009 | 0.244077948 | 0 | 0.1192353857 | 4.90E-05 | 0.03315130372 | 0.9999996973 | 2.72E-04 | 0.01703655604 | 0 | 0.003648920043 | 0.06442734253 |
| TARA_SAMEA2623649_METAG_EJBMLLJL | 4 | -0.08214726401 | Marinisomatales | GCA-002701945 | GCA-002701945 | 0 | 0 | 1.000000179 | 4.43E-05 | 1.000000179 | 0 | 0 | 0.06626791193 | 0 | 0 | 0.05865808327 |
| TARA_SAMEA2623649_METAG_JNHJPEEE | 7 | -0.09957056632 | Pseudomonadale | UBA11889 | UBA11889 sp002 | 0.3010026729 | 0 | 1.000000004 | 0.05000000439 | 0.9853274537 | 0.005502625316 | 0 | 0.07945977129 | 0 | 0.9833599053 | 0 |
| TARA_SAMEA2623673_METAG_BFAMLMFE | 1 | 0.02973022524 | Flavobacteriales | CAJWCP01 | CAJWCP01 sp9 | 0 | 0 | 0.9999998171 | 9.84E-05 | 0.3328470608 | 0.9999998171 | 0 | 0.06779103704 | 0 | 1.68E-05 | 0 |
| TARA_SAMEA2623734_METAG_PJGEFNLF | 7 | -0.01263704168 | Flavobacteriales | GCA-002697625 | GCA-002697625 | 0 | 0 | 1.000000274 | 1.00E-04 | 1.000000274 | 2.06E-08 | 0 | 0.07922315179 | 0 | 1.000000274 | 0.03549101844 |
| TARA_SAMEA2623756_METAG_BNJLBMPF | 4 | 0.05017079967 | Flavobacteriales | GCA-2706465 | | 0.05231499482 | 0 | 1.000000003 | 0.05238544848 | 2.83E-05 | 0.00408849336 | 0 | 0.04249352737 | 0 | 1.34E-05 | 0.03600998975 |
| TARA_SAMEA2623756_METAG_EILEHNFN | 6 | -0.0896656368 | Tepidiformales | JAKUNN01 | JAKUNN01 sp02 | 0 | 0 | 0.0282691448 | 0.01666907389 | 0.95 | 0 | 0 | 0.1041947297 | 0 | 0 | 0 |
| TARA_SAMEA2623756_METAG_FDADPHPN | 8 | -0.01561874073 | UBA1135 | UBA1135 | UBA1135 sp003 | 0.1070288419 | 0 | 1.000000193 | 0.9666688681 | 0.9833335264 | 0.9681935174 | 0 | 0.06864159882 | 0 | 2.81E-05 | 0.03479117845 |
| TARA_SAMEA2623756_METAG_HDODALJM | 2 | -0.1322071756 | JAAXHH01 | JAAXHH01 | | 0.03445802413 | 0 | 0.04171106025 | 4.04E-05 | 0.009357917224 | 4.00E-06 | 0 | 0.06003435038 | 0 | 0.004815215649 | 0 |
| TARA_SAMEA2623756_METAG_MIHFNHCC | 7 | -0.05700687106 | UBA3495 | VEXD01 | | 1.78E-06 | 0 | 0.1368256656 | 9.92E-05 | 0.9999999944 | 0.02355112262 | 0 | 0.01898442993 | 0 | 0.9999999944 | 0 |
| TARA_SAMEA2623756_METAG_NFIBGMOG | 3 | 0.0143288822 | UBA3495 | | | 1.03E-06 | 0 | 0.007998374541 | 1.21E-04 | 0.9999998969 | 0.9999998969 | 0 | 0.9999998969 | 0 | 0.9999998969 | 0 |
| TARA_SAMEA2623774_METAG_IHINPAGN | 2 | -0.02802227166 | Flavobacteriales | | | 0.1677114064 | 0 | 0.02240072191 | 1.11E-04 | 8.76E-04 | 0.01097751656 | 0 | 0.07329838131 | 0.01315387176 | 7.94E-04 | 0 |
| TARA_SAMEA2623826_METAG_FBPPGNKK | 2 | -0.1555437494 | Micavibrionales | UBA9664 | UBA9664 sp002 | 0 | 0 | 0.01256430099 | 0.01667113343 | 0.007127486381 | 0.03423322339 | 0 | 0.07093471647 | 0 | 2.58E-05 | 0.03378421368 |
| TARA_SAMEA2623850_METAG_NADLKMKF | 3 | -0.03050306775 | Actinomarinales | Actinomarina | Actinomarina sp | 0 | 0 | 0.03932175006 | 0.0168301164 | 0.9834537858 | 0.9999999956 | 0 | 0.9999999956 | 0.03728295076 | 0.9999999956 | 0 |
| TARA_SAMEA2623886_METAG_LLPNBDAJ | 5 | -0.07360414912 | Acidimicrobiales | MedAcidi-G1 | MedAcidi-G1 sp | 0 | 0 | 0.2608588556 | 1.33E-04 | 0.1995541659 | 1 | 0 | 0.04429027845 | 0.0056685921 | 7.91E-04 | 0 |
| TARA_SAMEA2623907_METAG_JBHOJJHF | 4 | -0.1933238845 | Rhodobacterales | Pacificitalea | Pacificitalea man | 0 | 0.09865710459 | 0.9860250336 | 8.78E-05 | 0.009403297127 | 0.05216396093 | 7.66E-05 | 0.2023863998 | 0 | 8.20E-04 | 0.02799804014 |
| TARA_SAMEA4396424_METAG_CMFOJHLK | 1 | -0.07138552725 | Marinisomatales | | | 0.01207460729 | 0 | 0.9999999949 | 9.01E-04 | 0.9999999949 | 0.5679213043 | 0.006481022918 | 0.2956327061 | 0 | 7.98E-04 | 0.02141290971 |
| TARA_SAMEA4396424_METAG_HIKNENFF | 8 | -0.00929590543 | Flavobacteriales | GCA-2683775 | GCA-2683775 sp | 0 | 0.02451879772 | 1.000000165 | 1.000000165 | 8.63E-04 | 1.000000165 | 0 | 0.06501987615 | 0 | 0 | 0 |
| TARA_SAMEA4396424_METAG_MEHMBKLF | 1 | -0.1049838178 | Opitutales | | | 0 | 0 | 1.000000227 | 1.47E-04 | 2.38E-04 | 1.000000227 | 0 | 1.000000227 | 0 | 8.03E-04 | 0.05332850969 |
| TARA_SAMEA4396468_METAG_DHGJMAAG | 7 | -0.03285648416 | Planctomycetales | UBA8437 | | 5.49E-04 | 0 | 0.09816320871 | 0.0166666631 | 0.0888923353 | 0.01666673378 | 2.40E-04 | 0.0657263778 | 9.63E-07 | 0.9666961183 | 1.58E-05 |
| TARA_SAMEA4396468_METAG_DOOIBEFB | 5 | -0.03658414286 | Arenicellales | REDSEA-S09-B1 | REDSEA-S09-B1 | 0 | 0 | 0.03189397302 | 1.09E-04 | 0.1211314202 | 0.9999997775 | 0 | 0.0470731672 | 0 | 0.01816270478 | 0.09815994598 |
| TARA_SAMEA4396468_METAG_PDKKHEEF | 6 | -0.09057325417 | SAR202 | UBA11996 | | 0.1906744887 | 0 | 0.0498292174 | 9.63E-05 | 1 | 0.004564070136 | 0.004216334856 | 0.09421022541 | 0.02639292945 | 0.00216309177 | 0.067611508 |
| TARA_SAMEA4396468_METAG_PMKCLFEO | 2 | 0.01615506634 | Verrucomicrobial | EC70 | EC70 sp0142393 | 0 | 0 | 0.03770955097 | 0.9999997127 | 0.008564408247 | 0.01666666244 | 0 | 0.1022953363 | 0 | 8.08E-04 | 0.03619107772 |
| TARA_SAMEA4396489_METAG_CIGGPLLK | 2 | -0.2789256309 | Rhizobiales | Devosia | | 0 | 0 | 0.03066727508 | 1.05E-05 | 0.003239405547 | 0 | 0 | 0.07223119493 | 0 | 0.01986966685 | 0 |
| TARA_SAMEA4396489_METAG_DJIFJGEM | 8 | -0.244084768 | Caulobacterales | Litorimonas | | 0 | 0 | 0.1046429817 | 0.9833332311 | 0.1446223196 | 0.9833334415 | 0 | 0.06945968398 | 0 | 1.29E-05 | 0.03495179421 |

| | | | | | | | | | | | | | | | | |
|---|---|---|---|---|---|---|---|---|---|---|---|---|---|---|---|---|
| TARA_SAMEA4396489_METAG_FPKKKDHE | 2 | -0.1063308655 | Rhizobiales | Jiella | | 0 | 0.006583887298 | 0.03759810213 | 0.0333786638 | 0.2160772041 | 1.46E-05 | 0 | 0.05623248676 | 0 | 7.47E-04 | 0 |
| TARA_SAMEA4396489_METAG_HMLIJBGP | 2 | -0.2339558761 | Enterobacterales | Idiomarina | Idiomarina abyss | 0 | 0 | 0.0294704048 | 1.28E-06 | 0.01741964311 | 2.95E-04 | 0.004108275553 | 0.2706087786 | 0 | 0 | 1.36E-04 |
| TARA_SAMEA4396489_METAG_JENLENIG | 6 | -0.2659064598 | Cytophagales | Algoriphagus | | 0 | 0 | 0.1422854423 | 1.30E-04 | 1.000000003 | 0.03333876652 | 0 | 0.2360222827 | 0 | 7.73E-04 | 0 |
| TARA_SAMEA4396489_METAG_KGPCBANM | 2 | -0.00896839312 | Xanthomonadale | Arenimonas | | 0 | 0 | 0.1211452421 | 0.01673799629 | 0.01578816506 | 0.01794836597 | 0 | 0.03440963056 | 0 | 7.64E-04 | 3.10E-07 |
| TARA_SAMEA4396489_METAG_MGODDFLG | 6 | -0.05576604243 | Actinomycetales | Pseudolysinimon | Pseudolysinimon | 1.52E-06 | 0 | 0.06040689296 | 1.00E-04 | 0.9999998098 | 0 | 0 | 0.07650260441 | 0 | 0 | 0 |
| TARA_SAMEA4396489_METAG_OCBFFKCH | 4 | -0.3348481091 | Flavobacteriales | Arenibacter | Arenibacter algic | 0 | 0 | 0.9999997762 | 6.09E-05 | 0.1264528335 | 0.002291182016 | 0 | 0.07875300248 | 0 | 0 | 0 |
| TARA_SAMEA4396489_METAG_OFBBDPAO | 6 | -0.1088973989 | Planctomycetales | Rubinisphaera | Rubinisphaera sp | 3.06E-06 | 0 | 0.03915228403 | 1.28E-04 | 0.9840272779 | 0.002841561124 | 0 | 0.2445521826 | 0 | 0 | 0.0263043265 |
| TARA_SAMEA4396489_METAG_PBAFPEGL | 2 | -0.04416677066 | Flavobacteriales | Olleya | Olleya marilimos | 0 | 0 | 0.03086618249 | 0.9999998261 | 0.01434144799 | 0.04213774253 | 0.08306053846 | 0.04072349322 | 0 | 0.001591639164 | 0.03570266469 |
| TARA_SAMEA4396496_METAG_FNFDMJKC | 2 | -0.01384695577 | Burkholderiales | BACL14 | | 0 | 0 | 0.1173330966 | 1.49E-04 | 0 | 0 | 0 | 1 | 0 | 7.52E-04 | 0.002854689685 |
| TARA_SAMEA4396496_METAG_HGBPLAFP | 2 | -0.2077565386 | Rhodobacterales | Amylibacter | Amylibacter sp00 | 0 | 0 | 0.05351905571 | 1.99E-04 | 0.001776263427 | 0.00435269773 | 5.02E-04 | 0.2604026049 | 0.03055923148 | 7.78E-04 | 0.001777286227 |
| TARA_SAMEA4396496_METAG_PFACBJFH | 6 | -0.1420891826 | PCC-6307 | Parasynechococ | Parasynechococ | 0 | 0 | 0.02240365992 | 1.69E-06 | 0.9833336124 | 1.35E-04 | 0 | 0.09650294915 | 0 | 0 | 1.45E-04 |
| TARA_SAMEA4396515_METAG_ADNJBJOC | 4 | -0.1218047704 | Flavobacteriales | Dokdonia | Dokdonia sp913 | 0 | 0 | 0.9838680263 | 0.9500012244 | 0.005620519784 | 0.03805291966 | 0 | 0.2229974384 | 0 | 7.99E-04 | 0.03627746962 |
| TARA_SAMEA4396515_METAG_BDMNHPLO | 2 | -0.1659853293 | Rhodobacterales | EhC02 | | 0 | 0 | 0.04334338949 | 6.44E-05 | 0.006831266638 | 0 | 0 | 0.1403533074 | 0 | 0 | 0 |
| TARA_SAMEA4396515_METAG_FELBDCME | 5 | -0.1707597152 | Flavobacteriales | Algorimicrobium | Algorimicrobium | 0 | 0 | 0.05424355495 | 1.30E-04 | 0.001387657176 | 0.9999997753 | 0 | 0.03123621695 | 0 | 7.43E-04 | 0.009380167063 |
| TARA_SAMEA4396515_METAG_FFGDBLLE | 2 | -0.154244959 | Thalassobaculale | Nisaea | | 0 | 0 | 0.02408208779 | 0.01667056278 | 0.08700538746 | 2.27E-04 | 0 | 0.03668226833 | 0 | 0.00372394212 | 0 |
| TARA_SAMEA4396515_METAG_GBPKHNMJ | 5 | -0.1607463041 | Cytophagales | Marivirga | | 0 | 0 | 0.03042439924 | 1.90E-06 | 0.1569297286 | 1 | 0 | 0.04212317483 | 0 | 1.34E-05 | 0 |
| TARA_SAMEA4396515_METAG_JCDCCEGE | 5 | -0.2085605016 | Flavobacteriales | Nonlabens | Nonlabens ulvan | 0 | 0 | 0.07018019408 | 1.40E-04 | 0.04302385761 | 1 | 0 | 0.0540196476 | 0 | 7.99E-04 | 0.005419546281 |
| TARA_SAMEA4396515_METAG_LLNDKEAP | 2 | -0.1018989435 | Caulobacterales | Brevundimonas | | 3.44E-06 | 0 | 0.0259105412 | 0.01675319268 | 0.08673291834 | 0 | 1.55E-05 | 0.06020253138 | 0 | 1.08E-07 | 0 |
| TARA_SAMEA4396538_METAG_CAOPAICI | 4 | -0.1594699011 | Rhodobacterales | Planktomarina | Planktomarina sp | 0 | 0 | 0.9999997602 | 2.01E-04 | 0.9999997602 | 0.06398868298 | 0 | 0.08156314743 | 0 | 7.63E-04 | 0 |
| TARA_SAMEA4396941_METAG_BMNNLBNP | 6 | -0.1338742537 | Pseudomonadale | Halioglobus | | 4.58E-04 | 0 | 0.03178926975 | 0.01666666934 | 0.9521057703 | 5.05E-05 | 4.38E-04 | 0.07759232887 | 0 | 2.53E-05 | 0.2074974278 |
| TARA_SAMEA4396941_METAG_CLEKMJLB | 2 | -0.09899816674 | Burkholderiales | JACKLP01 | | 0 | 0 | 0.1014217229 | 0.005372578142 | 8.47E-05 | 7.15E-04 | 0 | 0.9840183269 | 0 | 0.0166666697 | 0.02947798164 |
| TARA_SAMEA4396941_METAG_EHPFJEKM | 2 | -0.1559282344 | Rhodobacterales | G020447305 | G020447305 sp0 | 0 | 0 | 0.03195040368 | 6.15E-05 | 0.003347547436 | 4.93E-06 | 0 | 0.1535827924 | 0 | 7.89E-04 | 0 |
| TARA_SAMEA4396941_METAG_EJKIMIHL | 2 | -0.2361811065 | Pseudomonadale | Halomonas | Halomonas sp00 | 0 | 0 | 0.03352937717 | 1.48E-06 | 0.001725785726 | 0.00530077502 | 0 | 0.07990904158 | 0 | 1.26E-05 | 0 |
| TARA_SAMEA4396941_METAG_FIKOMDNM | 3 | -0.1546319827 | Pseudomonadale | Halioglobus | | 0 | 0 | 0.02079549042 | 2.44E-06 | 0.07491274345 | 1.000000004 | 0 | 0.03728769264 | 0 | 1.000000004 | 0.005550743978 |
| TARA_SAMEA4396941_METAG_HEODMAFM | 2 | -0.1152301854 | Burkholderiales | Thiobacillus | | 0 | 0 | 0.1506413516 | 4.00E-06 | 0.07302645642 | 0.01666666667 | 0 | 0.03830220692 | 0 | 2.46E-05 | 0 |
| TARA_SAMEA4396941_METAG_IJEMMHFI | 2 | -0.07532458101 | Burkholderiales | Azoarcus_D | | 0 | 0 | 0.03457861671 | 0.05005676956 | 6.02E-05 | 0.005433035585 | 0 | 0.9655588986 | 0 | 0.004753886677 | 0 |
| TARA_SAMEA4396941_METAG_JFDOBMBJ | 2 | -0.09139561048 | Woeseiales | UBA1844 | | 0 | 0 | 0.08219017007 | 1.40E-04 | 0.08287469319 | 9.04E-06 | 0 | 0.09723991016 | 0 | 0 | 0.05314678028 |
| TARA_SAMEA4396941_METAG_KAPCODLD | 2 | -0.06305518666 | Verrucomicrobial | JAGRPD01 | | 0 | 0 | 0.04187035502 | 5.93E-05 | 0.07167617443 | 0 | 0 | 0.03220545112 | 0 | 7.53E-04 | 0 |
| TARA_SAMEA4396941_METAG_LBLFLNAI | 2 | -0.2614184674 | Micavibrionales | UBA2705 | | 0.01910582609 | 0 | 0.1524223369 | 1.79E-05 | 0.0552588641 | 0.2297965246 | 0 | 0.06871135827 | 0 | 9.19E-05 | 2.91E-05 |
| TARA_SAMEA4396941_METAG_NMOKIAAF | 1 | -0.199954868 | Rhodobacterales | Sulfitobacter | | 0 | 0 | 1 | 1.14E-04 | 1 | 1 | 0 | 1 | 0 | 0 | 0 |
| TARA_SAMEA4396988_METAG_BLBNHPMH | 2 | -0.08267082795 | Flavobacteriales | Crocinitomix | Crocinitomix cat | 0.128338966 | 0 | 0.06174129862 | 1.07E-06 | 0.03768058044 | 0.001658937524 | 0.003401134294 | 0.04003993637 | 0 | 0 | 0 |
| TARA_SAMEA4396988_METAG_CABPDNKF | 6 | -0.2598241259 | Bacteroidales | Sunxiuqinia | | 0 | 0 | 0.02477682228 | 0.9666675632 | 0.9841664406 | 1.89E-08 | 0 | 0.09001129014 | 0 | 0 | 0 |
| TARA_SAMEA4396988_METAG_MIJAAAFI | 4 | -0.2270448409 | Flavobacteriales | CAJXZP01 | | 0 | 0 | 1 | 1.39E-04 | 1 | 0 | 1 | 0.08413198248 | 0 | 0.01514709406 | 0 |
| TARA_SAMEA4396988_METAG_NHAKENJI | 6 | -0.2853440221 | Flavobacteriales | Aequorivita | | 2.21E-06 | 0 | 0.03654231664 | 1.05E-04 | 0.9999997238 | 1.66E-06 | 0 | 0.06953167018 | 0 | 7.61E-04 | 0 |
| TARA_SAMEA4396988_METAG_OJLLLIJD | 8 | -0.1259673941 | Flavobacteriales | Yeosuana | | 5.97E-08 | 0 | 0.03018482185 | 1.000000214 | 0.1187175927 | 0.9833336708 | 0.00335679913 | 0.9833335534 | 2.15E-05 | 2.31E-05 | 1.14E-04 |
| TARA_SAMEA4397004_METAG_GDCOLFIP | 4 | -0.00191020356 | Marinisomatales | TMED80 | | 0 | 0 | 1.0000003 | 2.59E-04 | 0.03808622081 | 2.40E-06 | 0 | 1.0000003 | 0 | 0.003999065185 | 0 |
| TARA_SAMEA4397004_METAG_GIHJMGDM | 5 | -0.0677398745 | Pseudomonadale | UBA9145 | UBA9145 sp012 | 0 | 0 | 0.03646248937 | 1.17E-04 | 0 | 1 | 0 | 0.04105827106 | 0 | 0.01069496453 | 0.008230928585 |
| TARA_SAMEA4397004_METAG_JFHKLEJO | 4 | -0.01793426697 | Marinisomatales | UBA8229 | UBA8229 sp003 | 0 | 0 | 1.000000001 | 0.03342553938 | 0.0541549398 | 0.05265355579 | 0 | 0.0598128548 | 0 | 4.45E-05 | 0.03641744868 |
| TARA_SAMEA4397004_METAG_OEMGLDGO | 5 | -0.07610992723 | Woeseiales | SP4260 | | 0 | 0 | 0.02934758551 | 7.47E-05 | 0.07364923699 | 1.000000168 | 0 | 1.000000168 | 0 | 0.002481691617 | 0.03603449597 |
| TARA_SAMEA4397023_METAG_ACFKGOCL | 6 | -0.3123965389 | Rhizobiales | Pseudorhizobium | Pseudorhizobium | 0 | 0 | 0.1660164701 | 1.59E-06 | 0.9699413336 | 0 | 0 | 0.06821214372 | 0 | 0 | 0 |
| TARA_SAMEA4397023_METAG_BIAHDCHN | 6 | -0.1339375205 | Flavobacteriales | Maribacter | | 3.10E-05 | 0 | 0.1463296263 | 1.54E-06 | 0.9837448787 | 5.77E-05 | 0 | 0.0590864487 | 0 | 0.01666667024 | 0 |
| TARA_SAMEA4397023_METAG_CDAJJLGH | 3 | -0.2681094702 | Pseudomonadale | Halopseudomona | Halopseudomona | 0 | 0 | 1.000000264 | 0.01666750414 | 0.2712331229 | 1.000000264 | 0 | 0.05790283971 | 4.00E-05 | 0.9835114182 | 5.32E-04 |
| TARA_SAMEA4397023_METAG_DCKGLPKH | 6 | -0.03777935196 | Xanthomonadale | Arenimonas | | 0 | 0 | 0.02890377323 | 0.03333558384 | 0.9671255242 | 7.57E-05 | 0 | 0.0794571814 | 0 | 0.01674860512 | 0 |
| TARA_SAMEA4397023_METAG_DPNJCNNI | 5 | -0.1982772789 | Flavobacteriales | Leeuwenhoekiell | Leeuwenhoekiell | 0 | 0 | 0.09146768147 | 4.29E-06 | 0.00798659247 | 0.9500008248 | 0 | 0.06501437724 | 0 | 6.64E-05 | 0 |
| TARA_SAMEA4397023_METAG_FBMOJIEP | 2 | -0.2208804247 | Pseudomonadale | Halopseudomona | Halopseudomona | 0 | 0 | 0.1099000885 | 7.18E-05 | 0.001013965043 | 0.0121008117 | 0 | 0.06600903221 | 0 | 7.91E-04 | 0 |
| TARA_SAMEA4397023_METAG_FMECILJE | 5 | -0.1484099193 | Caulobacterales | Hyphomonas | Hyphomonas adh | 8.26E-07 | 0 | 0.0424731315 | 1.23E-04 | 0.002202243189 | 1.000000207 | 0.02702272215 | 0.1169866289 | 0 | 0.002726980309 | 0 |
| TARA_SAMEA4397023_METAG_GPOLAJDG | 4 | -0.2923863099 | Rhodobacterales | Yoonia | | 2.52E-04 | 0 | 0.9861917491 | 0.08333398618 | 0.07785844693 | 0.00395491825 | 0 | 0.06202234326 | 0 | 6.18E-05 | 0.03227805185 |
| TARA_SAMEA4397023_METAG_JDCBPCFM | 6 | -0.1937898688 | Rhizobiales | Hoeflea | | 0 | 0 | 0.0258977015 | 1.08E-04 | 0.9834476203 | 5.81E-08 | 0 | 1.000000286 | 0 | 0.001598582469 | 0 |
| TARA_SAMEA4397023_METAG_KOLKIBFJ | 6 | -0.1947611685 | Flavobacteriales | Dokdonia | | 0 | 0 | 0.03628583166 | 1.69E-04 | 1 | 1 | 0 | 0.06511445192 | 0 | 0 | 0.05428052433 |
| TARA_SAMEA4397034_METAG_ANPPJHKJ | 6 | -0.234155303 | Flavobacteriales | Hel1-33-131 | | 0 | 0 | 0.5729534757 | 1.66E-05 | 1 | 0 | 0 | 0.0595586209 | 0 | 0 | 0.1596398759 |
| TARA_SAMEA4397034_METAG_GPHNGABH | 2 | -0.1643492035 | Flavobacteriales | NORP294 | NORP294 sp905 | 9.02E-06 | 0 | 0.022319803 | 0.03333700766 | 0.03493631485 | 3.08E-04 | 0 | 0.04707308707 | 0 | 1.08E-05 | 0 |
| TARA_SAMEA4397051_METAG_ANCHFCML | 5 | -0.1590618632 | Flavobacteriales | Yeosuana | | 0 | 0 | 0.03643428575 | 1.18E-04 | 0.02105471977 | 1 | 0.01099911335 | 0.04796684339 | 0 | 0 | 0.00399377608 |
| TARA_SAMEA4397051_METAG_BDIOPMOI | 5 | -0.3876638868 | Rhodospirillales_ | Thalassospira | | 0 | 0 | 0.1602305436 | 1.08E-04 | 0 | 1 | 0 | 0.1419082837 | 0 | 0.001479163704 | 0 |
| TARA_SAMEA4397051_METAG_GINDMBMO | 6 | -0.1272452667 | Campylobacteral | Arcobacter | | 1.53E-08 | 0 | 0.03151564033 | 4.35E-06 | 0.9505828609 | 0 | 0 | 0.05258182438 | 0 | 1.39E-05 | 0 |
| TARA_SAMEA4397101_METAG_GMHNMKNJ | 2 | -0.03885197 | Puniceispirillales | CAJJBL01 | | 0 | 0 | 0.02295070679 | 1.10E-04 | 0.01075751472 | 0.1309441272 | 0 | 0.06940859312 | 0 | 0.001951728744 | 0.01376611033 |
| TARA_SAMEA4397118_METAG_CEBIIJJH | 3 | -0.2001597906 | Flavobacteriales | Planktosalinus | | 0 | 0 | 0.0211278751 | 8.82E-05 | 0.3349121073 | 1 | 0 | 0.07994634491 | 0 | 1 | 0 |
| TARA_SAMEA4397118_METAG_DAPHGPDF | 4 | -0.164964479 | Pseudomonadale | Pseudohongiella | | 0 | 0 | 0.9999998053 | 9.34E-05 | 0.9999998053 | 0.008753234432 | 0 | 0.7656686069 | 0 | 0 | 0.0201350865 |
| TARA_SAMEA4397118_METAG_DDLCHGOL | 2 | -0.06670860321 | Flavobacteriales | | | 0 | 0 | 0.1600786496 | 3.97E-06 | 0.01762977348 | 1.74E-04 | 0 | 0.09822657536 | 0 | 0 | 8.63E-04 |
| TARA_SAMEA4397118_METAG_GHBEGJFO | 5 | -0.0925914863 | Sphingomonadal | Qipengyuania | | 0 | 0 | 0.0486162149 | 0.01666666149 | 0.002686047839 | 0.9999999948 | 0 | 0.05044566361 | 0 | 3.46E-04 | 0.03077428916 |
| TARA_SAMEA4397118_METAG_HIJBLIPC | 2 | -0.1527591151 | Pseudomonadale | UBA4582 | | 0 | 0 | 0.06893588777 | 1.000000186 | 0.01480470416 | 0.01088500439 | 0 | 0.02532278162 | 0 | 8.23E-04 | 0 |
| TARA_SAMEA4397118_METAG_NDKAGOCA | 4 | -0.2610454393 | Kiloniellales | | | 0 | 0 | 1.000000004 | 1.37E-04 | 1.000000004 | 0.007244380583 | 0 | 0.05210622386 | 0 | 0 | 0 |
| TARA_SAMEA4397118_METAG_OKNDPBEJ | 2 | -0.07317460048 | Pseudomonadale | Halioglobus | | 0 | 0.005551973505 | 0.03587390383 | 1.08E-04 | 0.2074940894 | 0.001315699111 | 0 | 0.07896958141 | 0 | 9.83E-04 | 0.001609149071 |
| TARA_SAMEA4397118_METAG_PGCEFNEO | 1 | -0.1107497483 | Pseudomonadale | Halioglobus | Halioglobus sp0 | 0.001367180825 | 0 | 1 | 2.04E-06 | 7.15E-04 | 1 | 2.11E-05 | 0.04577357514 | 3.45E-04 | 1.39E-05 | 6.53E-05 |
| TARA_SAMEA4397157_METAG_AHGIHHGL | 1 | -0.262633081 | Rhodospirillales_A | | | 0 | 0.002858525819 | 1 | 7.52E-05 | 2.54E-04 | 1 | 0 | 0.1937593449 | 0 | 0 | 0 |
| TARA_SAMEA4397157_METAG_OEPPPCEN | 5 | -0.1274771961 | Flavobacteriales | UBA2108 | | 0 | 0 | 0.03406245775 | 1.26E-06 | 0.1958304793 | 1.000000004 | 0 | 0.05258770509 | 0 | 1.35E-05 | 0 |
| TARA_SAMEA4397174_METAG_DPMFAFMH | 5 | -0.07844971657 | Flavobacteriales | Pontimicrobium | | 0 | 0 | 0.03240558585 | 2.13E-06 | 0.1307675408 | 1.000000256 | 0 | 0.9841780919 | 0 | 0.002847240277 | 0 |
| TARA_SAMEA4397174_METAG_EHJFDMGO | 1 | -0.1027613941 | Pseudomonadale | PGZG01 | PGZG01 sp0180 | 0 | 0 | 1.000000234 | 0.2105522767 | 0 | 1.000000234 | 0 | 0.04200339628 | 0 | 0.005896655782 | 0 |
| TARA_SAMEA4397174_METAG_HBDIMJIM | 5 | -0.201466224 | Pseudomonadale | GM7GCV1 | | 0 | 0 | 0.0257765737 | 5.69E-07 | 0 | 1.000000003 | 0 | 0.3136458531 | 0 | 1.34E-05 | 0 |
| TARA_SAMEA4397174_METAG_KJLHPIFN | 1 | -0.05862371506 | Phycisphaerales | UBA1924 | | 0 | 0 | 1 | 2.34E-06 | 0.03635586951 | 1 | 0 | 0.05392218914 | 0 | 2.37E-05 | 0 |
| TARA_SAMEA4397174_METAG_KMPDDMPD | 4 | -0.05605207664 | Flavobacteriales | GCA-002733185 | GCA-002733185 | 0 | 0 | 1.000000272 | 1.12E-04 | 0.1066895481 | 0 | 0 | 0.08019072575 | 0 | 3.29E-06 | 3.96E-04 |
| TARA_SAMEA4397174_METAG_KNDEEOOG | 5 | -0.1072125116 | Flavobacteriales | SCGC-AAA160-P02 | | 0 | 0 | 0.04087650989 | 9.76E-05 | 0.2084970182 | 1 | 0 | 0.02670563211 | 0 | 7.93E-04 | 0 |
| TARA_SAMEA4397174_METAG_LMPNIEMG | 1 | -0.1384510427 | Pseudomonadale | Haliea | Haliea sp002416 | 0 | 0 | 1 | 1.28E-04 | 1 | 1 | 0 | 0.07931316516 | 0 | 0.004279045972 | 0 |
| TARA_SAMEA4397174_METAG_MAHICFFA | 1 | -0.1461069399 | Phycisphaerales | UBA1924 | | 0 | 0 | 0.9999998008 | 1.06E-04 | 0.0882509545 | 0.9999998008 | 0 | 0.9999998008 | 0 | 0.001582817762 | 0 |

| | | | | | | | | | | | | | | | | |
|---|---|---|---|---|---|---|---|---|---|---|---|---|---|---|---|---|
| TARA_SAMEA4397174_METAG_OLMHMLCA | 4 | -0.1629018231 | Rhodobacterales | SW4 | SW4 sp0027328 | 2.68E-06 | 0 | 1 | 0.01443190623 | 0 | 0.08841653865 | 0 | 4.14E-04 | 0 | 0 | 0 |
| TARA_SAMEA4397196_METAG_BMMPCMEC | 2 | -0.2024249527 | Flavobacteriales | Crocinitomix | Crocinitomix sp9 | 0.3062024002 | 0 | 0.05454406844 | 1.59E-04 | 0.1415554999 | 0.003600954857 | 0 | 0.06790278079 | 0 | 0 | 0 |
| TARA_SAMEA4397196_METAG_DBBDHPDM | 6 | -0.2673721185 | Flavobacteriales | UBA974 | | 0 | 0 | 0.03180144243 | 2.12E-06 | 1.000000004 | 6.64E-04 | 0 | 0.07918056005 | 0 | 2.61E-05 | 0.0365738731 |
| TARA_SAMEA4397196_METAG_EFAOIEIE | 7 | -0.1491188962 | Opitutales | | | 0.1153873433 | 0 | 1 | 2.96E-05 | 0.007493631834 | 0.004804515762 | 0.01056091831 | 0.01647611224 | 0 | 1 | 0.03490280274 |
| TARA_SAMEA4397196_METAG_GOGNMPAD | 4 | -0.1109750525 | Verrucomicrobial | Luteolibacter | Luteolibacter sp9 | 0 | 0 | 0.9999997925 | 0.01676756452 | 0.01666667171 | 0.01610100839 | 0.003646844908 | 0.2967422628 | 0 | 0 | 0 |
| TARA_SAMEA4397196_METAG_OINJKPLG | 2 | -0.1375216418 | Cytophagales | UBA4465 | UBA4465 sp905 | 5.66E-08 | 0 | 0.02630727905 | 0.9999999999 | 0.003387930987 | 0.06948514318 | 0 | 0.9838749422 | 0 | 0.003539119564 | 0 |
| TARA_SAMEA4397215_METAG_AJJFEKGE | 2 | -0.1169109979 | Rhodobacterales | Alterinioella | | 0 | 0 | 0.1456446836 | 0 | 5.16E-05 | 0.0166666613 | 0 | 0.04973301682 | 0.0166666613 | 0.008222334821 | 0 |
| TARA_SAMEA4397215_METAG_CJAEGIAG | 2 | -0.07492787545 | Parvibaculales | Parvibaculum | | 0 | 0 | 0.04452416309 | 1.61E-06 | 0.001472928219 | 0.01666666667 | 0 | 0.03129819444 | 0 | 0.007600106698 | 0 |
| TARA_SAMEA4397215_METAG_DPBONLII | 2 | -0.2021858942 | Pseudomonadale | PZPK01 | | 0 | 0 | 0.04383027609 | 0.01666687428 | 0.04821412619 | 5.16E-04 | 0.03141877635 | 1.000000221 | 1.98E-05 | 5.59E-06 | 0 |
| TARA_SAMEA4397215_METAG_GIHNDBCM | 2 | -0.1125635207 | Flavobacteriales | SCGC-AAA160-F | SCGC-AAA160-F | 0 | 0 | 0.02639784365 | 1.19E-06 | 0.004231768288 | 0.03333332831 | 0 | 0.9842923816 | 0 | 5.32E-05 | 0 |
| TARA_SAMEA4397215_METAG_HKNGHGKC | 7 | -0.08346508003 | Flavobacteriales | Winogradskyella | | 0 | 0 | 0.01554941561 | 1.23E-06 | 0.2437535502 | 4.35E-05 | 0 | 0.02764634693 | 0 | 1.000000192 | 0 |
| TARA_SAMEA4397215_METAG_KLMFFEGC | 8 | -0.207567893 | Methylococcales | Cycloclasticus | | 0 | 0 | 0.9999997974 | 0.9999997974 | 3.65E-06 | 0.9999997974 | 0 | 0.5131405748 | 0 | 0.2544786982 | 0 |
| TARA_SAMEA4397239_METAG_IKJKOEDG | 3 | -0.03150650933 | Flavobacteriales | Urechidicola | | 0.08226329179 | 0 | 3.33E-04 | 1.77E-04 | 0.02704265234 | 1 | 0.001830459096 | 0.002544006768 | 0 | 0.9837335603 | 0.09067318674 |
| TARA_SAMEA4397255_METAG_BAGPGNGF | 4 | -0.26231008 | Micavibrionales | UBA1672 | | 0 | 0 | 1.000000012 | 7.59E-05 | 0.1306459537 | 0.001354792833 | 0 | 0.005692807201 | 0 | 3.98E-04 | 0.03447768953 |
| TARA_SAMEA4397255_METAG_KKHJAOMA | 2 | -0.1063487538 | Flavobacteriales | Algibacter_B | | 0.001267907222 | 0 | 0.02979265485 | 0 | 0 | 3.67E-04 | 3.82E-06 | 0.1675725284 | 6.65E-05 | 0.0225177955 | 0 |
| TARA_SAMEA4397255_METAG_LHBNFFOA | 4 | -0.1555644544 | Planctomycetales | Gimesia | | 0.2433032968 | 0 | 1 | 1.25E-06 | 1 | 1.06E-07 | 0 | 0.9684247045 | 7.45E-05 | 0.0166671669 | 0 |
| TARA_SAMEA4397255_METAG_OLNPCAAC | 4 | -0.06696761697 | Phycisphaerales | UBA1924 | | 0.02031411534 | 0 | 1.000000137 | 0.03333332982 | 8.96E-05 | 0 | 2.67E-05 | 0.2633948623 | 0 | 0.06666666049 | 0.03203221867 |
| TARA_SAMEA4397255_METAG_PFEJDDLC | 4 | -0.1113134923 | Rhodospirillales_ | HT1-32 | HT1-32 sp00964 | 0 | 0.01259633465 | 1 | 5.70E-05 | 0.08053972356 | 0.004918315599 | 0 | 0.06493738994 | 0 | 0.02132029595 | 0.0657082841 |
| TARA_SAMEA4397295_METAG_BEKHEJEB | 2 | -0.06763353284 | Planctomycetales | Symmachiella | Symmachiella dy | 0 | 0 | 0.03240516143 | 3.83E-06 | 0.0139600952 | 0.01824870598 | 0 | 0.00718201348 | 0 | 0 | 0 |
| TARA_SAMEA4397295_METAG_EHHOEIHM | 6 | -0.2784653086 | Sphingomonadal | Kordiimonas | | 0 | 0 | 0.06154523524 | 7.89E-05 | 0.5681238939 | 0.01071306798 | 0 | 0.4592689152 | 0 | 0.004760821159 | 0.01161322153 |
| TARA_SAMEA4397295_METAG_GHOELJED | 2 | -0.1243877686 | Woeseiales | UBA1847 | | 0 | 0 | 0.07069872041 | 9.18E-07 | 0.01695930279 | 0 | 0 | 0.06278315274 | 0 | 8.14E-04 | 0.07645416118 |
| TARA_SAMEA4397295_METAG_HEGEJJIG | 5 | -0.230578016 | Haliangiales | NORP324 | | 0 | 0 | 0.03238471574 | 2.75E-06 | 0.1147357955 | 0.9833330807 | 0 | 0.0571171279 | 0 | 0 | 0.03592635592 |
| TARA_SAMEA4397295_METAG_JMJMPGEN | 2 | -0.05630192562 | Balneolales | Balneola | | 0 | 0 | 0.2362124278 | 2.11E-04 | 0.06530658925 | 7.04E-06 | 0 | 0.05039734905 | 0 | 7.88E-04 | 0 |
| TARA_SAMEA4397295_METAG_KBDLACKE | 2 | -0.3543740387 | Rhodobacterales | Amylibacter | | 0.1465964659 | 0 | 0.0137033288 | 6.94E-05 | 0 | 0 | 0 | 1 | 0 | 0 | 0 |
| TARA_SAMEA4397295_METAG_LIPLIBEG | 3 | -0.08890941816 | Cytophagales | | | 0 | 0 | 0.01951214017 | 6.39E-07 | 0.3094044889 | 1.000000164 | 0.001204516463 | 0.9839298172 | 0 | 1.000000164 | 0.02853065613 |
| TARA_SAMEA4397295_METAG_MLOPDMFJ | 2 | -0.1438720814 | Methylococcales | Cycloclasticus | Cycloclasticus pu | 0 | 0 | 0.105852527 | 4.88E-06 | 0.0189850759 | 7.18E-06 | 0 | 0.04211090218 | 0 | 0.001545282694 | 0 |
| TARA_SAMEA4397295_METAG_NDHNHJHC | 4 | -0.2189489498 | Rhizobiales | GCA-2746425 | | 0 | 0 | 1 | 6.06E-05 | 0.07743333034 | 0 | 0 | 0.04858298914 | 0 | 0.002010191254 | 0.03364460806 |
| TARA_SAMEA4397295_METAG_NFMMNHJF | 2 | -0.07130359282 | Ga0077536 | | | 5.39E-08 | 0 | 0.05409919446 | 2.78E-06 | 0.1138749397 | 8.24E-05 | 0 | 0.0672349716 | 0 | 0 | 0 |
| TARA_SAMEA4397295_METAG_NOICPOFE | 2 | -0.09613978423 | Verrucomicrobiales | | | 0 | 0.002553779055 | 0.03636586318 | 9.22E-05 | 0.0074876437 | 9.44E-07 | 2.01E-05 | 0.2428682546 | 0 | 2.71E-06 | 4.97E-04 |
| TARA_SAMEA4397295_METAG_PDFOFGBN | 4 | -0.2426552854 | Flavobacteriales | Owenweeksia | | 0 | 0 | 1.000000181 | 8.74E-07 | 0.1037154181 | 0.03637126123 | 0 | 0.05476718987 | 0 | 0.02319662542 | 0 |
| TARA_SAMEA4397311_METAG_CNLAMCIL | 5 | -0.4264844979 | Actinomycetales | Aquiluna | Aquiluna sp9051 | 0.00344818502 | 0 | 0.136596643 | 1.74E-06 | 0.05630403583 | 0.966708116 | 1.79E-04 | 0.02639907366 | 0 | 0.001562150522 | 0 |
| TARA_SAMEA4397330_METAG_AFKOBOMF | 2 | -0.1103766666 | Flavobacteriales | Ulvibacter | | 0 | 0 | 0.1516351337 | 1.08E-04 | 0.001237054077 | 0.001162280498 | 0.02777778609 | 0.06141333428 | 0 | 7.91E-04 | 0.01457695951 |
| TARA_SAMEA4397330_METAG_AJIEMPFL | 2 | -0.2271752162 | Rhodobacterales | Sulfitobacter | | 0 | 0 | 0.02498541775 | 4.94E-06 | 0.001170809358 | 0.03434954424 | 5.08E-04 | 0.02158523625 | 0 | 0.0103915081 | 7.77E-05 |
| TARA_SAMEA4397330_METAG_BNOLGDGH | 6 | -0.1208302929 | Parvibaculales | Mf105b01 | Mf105b01 sp002 | 0 | 0 | 0.1211138989 | 1.52E-06 | 0.9833333333 | 2.95E-04 | 0 | 0.06328309599 | 0 | 5.89E-08 | 0 |
| TARA_SAMEA4397330_METAG_DLAKABPD | 4 | -0.1719776105 | Flavobacteriales | Flavobacterium | | 0 | 1 | 1 | 4.40E-05 | 0.04402641434 | 2.32E-06 | 0 | 1 | 0 | 3.36E-05 | 0.001512138825 |
| TARA_SAMEA4397330_METAG_EKBCNCBM | 4 | -0.1792469043 | Balneolales | Balneola | | 4.68E-06 | 0 | 1 | 4.06E-05 | 0.02048243418 | 4.29E-06 | 0 | 0.2091771079 | 0 | 0 | 0 |
| TARA_SAMEA4397330_METAG_FBBFIGGC | 4 | -0.08012770122 | Flavobacteriales | Polaribacter_A | | 0 | 0 | 1 | 0.1644263881 | 3.20E-04 | 2.48E-04 | 6.75E-04 | 0.016062673 | 0 | 0.01801509539 | 4.04E-05 |
| TARA_SAMEA4397330_METAG_GDIMMCKF | 5 | -0.04201268258 | Parvibaculales | Parvibaculum | | 0 | 0 | 0.1691198514 | 8.69E-08 | 0.01385592041 | 0.9833693506 | 0 | 0.03546606024 | 0 | 4.51E-05 | 0 |
| TARA_SAMEA4397330_METAG_GFBBFFPE | 1 | -0.1153743141 | Rhizobiales | JAJFHR01 | | 0 | 0 | 0.9999998214 | 1.20E-06 | 0.08119943138 | 0.9999998214 | 0 | 0.06940512467 | 0 | 1.29E-05 | 5.42E-08 |
| TARA_SAMEA4397330_METAG_HKBLOPMM | 3 | -0.2457343421 | Rhodobacterales | Pseudorhodobacter | | 0 | 0 | 0.07034573807 | 0.9999998646 | 0.003214894823 | 0.9999998646 | 7.87E-05 | 0.0525949894 | 3.93E-04 | 0.9999998646 | 0.03717754188 |
| TARA_SAMEA4397330_METAG_HPNADKDC | 3 | -0.2899916263 | Rhodobacterales | Sulfitobacter | | 3.88E-04 | 0 | 0.05143593915 | 1.25E-04 | 0.003711246308 | 0.9999999974 | 3.88E-04 | 0.06990786133 | 0 | 0.9834984502 | 1.67E-05 |
| TARA_SAMEA4397330_METAG_INBJHHPP | 6 | -0.02275523719 | UBA6615 | UBA8079 | | 0 | 0 | 0.03178931315 | 1.25E-04 | 1.000000224 | 0 | 0 | 0.02568716798 | 0 | 7.06E-04 | 0.3240570139 |
| TARA_SAMEA4397330_METAG_KGKHFCBB | 5 | -0.09825114975 | Pseudomonadale | Porticoccus | | 0 | 0 | 0.02716598444 | 0.1110980542 | 0 | 1 | 0 | 0.003906036177 | 0 | 0.001846553744 | 0.03419012021 |
| TARA_SAMEA4397330_METAG_KHDOKGAD | 2 | -0.03340527965 | Minwuiales | Minwuia | | 0 | 0 | 0.03790761168 | 1.82E-06 | 0 | 8.48E-04 | 0.004471859843 | 0.05420019542 | 0 | 0.01639132405 | 0 |
| TARA_SAMEA4397330_METAG_KLOKAHBC | 5 | -0.1988655708 | Pseudomonadale | Halioglobus | | 1.07E-08 | 0 | 0.03268969802 | 0.003217057541 | 7.94E-04 | 0.9833426963 | 0 | 1.000000114 | 0 | 8.04E-04 | 0.006557660237 |
| TARA_SAMEA4397330_METAG_LHLFCDNB | 5 | -0.1265680684 | Acidimicrobiales | Ilumatobacter_A | | 0 | 0 | 0.05901729131 | 2.50E-06 | 0.02505550811 | 1.000000008 | 0 | 0.03969256379 | 0 | 8.43E-05 | 0 |
| TARA_SAMEA4397330_METAG_MGNNFKFN | 5 | -0.07463210922 | UBA4486 | UBA7359 | | 3.16E-06 | 0 | 0.04954497611 | 6.28E-05 | 0.00463547892 | 1 | 0 | 0.08545281081 | 0 | 0 | 1 |
| TARA_SAMEA4397330_METAG_MHHDJKOH | 2 | -0.07581806622 | MPNO01 | UBA830 | | 0 | 0 | 0.04381778549 | 4.54E-06 | 0.09101698564 | 0.0166667765 | 0 | 0.0205182213 | 0 | 5.20E-05 | 0.00263928198 |
| TARA_SAMEA4397330_METAG_NDOPFAKP | 4 | -0.1233883094 | Pseudomonadale | Haliea | | 0 | 0 | 1.000000004 | 1.000000004 | 0.3086301866 | 0.03333333734 | 5.14E-04 | 0.07284909751 | 0 | 7.84E-04 | 2.40E-05 |
| TARA_SAMEA4397330_METAG_ONJLDILD | 7 | -0.08089561613 | Acidimicrobiales | Ilumatobacter_A | | 0 | 0 | 0.03298148603 | 0.01666667193 | 0.01857440956 | 0.03333334267 | 0 | 0.05727181237 | 0 | 1.000000203 | 0 |
| TARA_SAMEA4397330_METAG_PICKGHED | 7 | -0.1648813006 | Pseudomonadale | UBA4582 | | 0 | 0 | 0.5730024797 | 0.03335491266 | 0.9667617717 | 0.03515460715 | 0 | 0.04783640811 | 0 | 1.000000286 | 0.06569030751 |
| TARA_SAMEA4397355_METAG_ADHJNKOG | 6 | -0.03382261687 | UBA1151 | GCA-2730555 | | 7.65E-08 | 0 | 0.002287928402 | 6.97E-05 | 0.9833847701 | 1.00000031 | 0 | 0.05420002018 | 0 | 0.05841921747 | 0 |
| TARA_SAMEA4397355_METAG_PKFDAOLJ | 5 | -0.04291807118 | Flavobacteriales | DUAL01 | DUAL01 sp0129 | 0 | 0 | 0.04441482711 | 1.01E-04 | 0.2633941061 | 0.9999996839 | 0 | 0.02992432301 | 0 | 0.001495900026 | 0 |
| TARA_SAMEA4397363_METAG_HEOAEBLP | 7 | -0.00566912892 | Phycisphaerales | JAJRRS01 | | 0 | 0 | 0.1594288471 | 3.67E-06 | 0.1642925885 | 0.01674563088 | 0 | 0.02688399104 | 1.26E-05 | 1.000000008 | 0 |
| TARA_SAMEA4397363_METAG_IDONDPJC | 5 | -0.2028809736 | Methylococcales | Cycloclasticus | Cycloclasticus sp | 5.91E-08 | 0 | 0.01823684816 | 1.66E-06 | 6.25E-04 | 0.9999997869 | 0 | 0.9606694319 | 0 | 7.51E-07 | 0 |
| TARA_SAMEA4397363_METAG_IPJBEJHI | 2 | -0.03121078351 | UBA2386 | GCA-2684655 | | 0 | 0 | 0.1047465012 | 6.08E-05 | 0 | 4.71E-06 | 0 | 0.1373205629 | 0 | 0.01814497212 | 0 |
| TARA_SAMEA4397390_METAG_EJBOOLAH | 6 | -0.1784376864 | Flavobacteriales | ASP10-05a | ASP10-05a sp90 | 0 | 0 | 0.1434750955 | 3.49E-06 | 0.9839167715 | 0.003565586948 | 0 | 0.0425974636 | 0 | 3.72E-04 | 0.02969605657 |
| TARA_SAMEA4397390_METAG_OAJNPHCG | 2 | -0.05205172045 | Flavobacteriales | GCA-002722245 | GCA-002722245 | 0 | 1.66E-05 | 0.04087830351 | 3.00E-04 | 2.15E-05 | 0.03531806131 | 0 | 0.07408283234 | 0 | 0.01745367501 | 0 |
| TARA_SAMEA4397409_METAG_KPGCMJHM | 4 | -0.2385104181 | Pseudomonadale | Marinobacter | Marinobacter sp0 | 0 | 0 | 1 | 0 | 0.2809736816 | 8.90E-07 | 0 | 0.07696947537 | 0 | 0 | 0.05772421518 |
| TARA_SAMEA4397426_METAG_APFFFHCD | 8 | -0.2289622345 | Opitutales | Lentimonas | | 8.22E-04 | 0 | 0.3431322683 | 1.000000196 | 0.01086655683 | 0.9669537534 | 1.39E-04 | 0.05855157636 | 5.99E-04 | 2.74E-04 | 3.43E-05 |
| TARA_SAMEA4397426_METAG_PDFLOHJD | 2 | -0.1412249193 | Burkholderiales | GCA-2401735 | | 0.02945586118 | 0 | 0.08679327695 | 1.53E-06 | 0.01085614289 | 0 | 0.01073607407 | 0.02292526725 | 0 | 0.01666666667 | 0.002135044405 |
| TARA_SAMEA4397445_METAG_AIAEOEKP | 2 | -0.1885829854 | Pseudomonadale | Halioglobus | | 0 | 0 | 0.05832597474 | 6.28E-05 | 0.0178626769 | 5.63E-06 | 0 | 0.02319902182 | 0 | 7.84E-04 | 0 |
| TARA_SAMEA4397445_METAG_CEKAOEIO | 1 | -0.3225925708 | Enterobacterales | Glaciecola | | 0 | 0.001757986462 | 0.9999997834 | 2.17E-05 | 0.002752403047 | 0.9999997834 | 0 | 0.04238180963 | 0 | 7.57E-04 | 0.001079738914 |
| TARA_SAMEA4397445_METAG_CHDHEFAE | 5 | -0.1278494485 | Rhodobacterales | Pseudooceanicol | Pseudooceanicol | 0 | 0 | 0.0333889112 | 0.03333333333 | 0.0639908298 | 1 | 0 | 0.04438763981 | 0 | 2.71E-05 | 0.03156910878 |
| TARA_SAMEA4397445_METAG_GIPEALJA | 6 | -0.2685531053 | Cytophagales | Dyadobacter | | 0 | 0 | 0.09134473345 | 0.01666666667 | 1.0000002 | 4.03E-04 | 0 | 0.05066636743 | 0 | 0 | 0 |
| TARA_SAMEA4397445_METAG_HLJHHDAL | 1 | -0.1397531527 | Cytophagales | Cyclobacterium | Cyclobacterium a | 0 | 0 | 1.000000194 | 0.03343953486 | 2.35E-05 | 1.000000194 | 0 | 0.08152210264 | 0 | 7.74E-04 | 0 |
| TARA_SAMEA4397445_METAG_HMENGOCD | 8 | -0.2581307822 | Micavibrionales | UBA2705 | | 0 | 0 | 1 | 1 | 0.003166888076 | 1 | 0 | 0.05632149895 | 0 | 7.52E-04 | 7.76E-04 |
| TARA_SAMEA4397445_METAG_JDEGCGLB | 4 | -0.08093973265 | Micavibrionales | CSBR16-224 | CSBR16-224 sp0 | 0 | 0 | 0.968453559 | 0.03333333389 | 0.003843353857 | 1.81E-07 | 0 | 0.05265170802 | 0 | 0.01666666361 | 0 |
| TARA_SAMEA4397445_METAG_OIJNNMDI | 2 | -0.05848067034 | Verrucomicrobial | JAGRPD01 | | 0 | 0 | 0.04079183231 | 8.46E-05 | 5.08E-05 | 0 | 0 | 0.2172748479 | 0 | 7.54E-04 | 0 |
| TARA_SAMEA4397472_METAG_CJMGHPKL | 3 | -0.1698448006 | Flavobacteriales | MAG-120531 | | 0 | 0 | 0.9999996251 | 9.30E-05 | 0.2434012175 | 0.9999996251 | 0 | 0.1594521277 | 0 | 0.9999996251 | 0 |
| TARA_SAMEA4397472_METAG_EEFNKHIK | 4 | -0.09784527857 | Rickettsiales | | | 0 | 0 | 1 | 0.01666666667 | 0 | 0 | 0 | 0.2783255148 | 0 | 7.15E-05 | 0 |
| TARA_SAMEA4397472_METAG_FANKLAJP | 8 | -0.1102046755 | Pseudomonadale | UBA4582 | UBA4582 sp0132 | 0 | 0 | 1 | 1 | 0.006317923466 | 1 | 0 | 1 | 0 | 0 | 0 |
| TARA_SAMEA4397472_METAG_KPDDLJNG | 6 | -0.2138738024 | Rhodobacterales | SW4 | | 0 | 0 | 0.053345672 | 0.9833348154 | 0.9846398045 | 8.57E-05 | 0 | 0.06918985085 | 0 | 0 | 0.03376340545 |

| | | | | | | | | | | | | | | | | |
|---|---|---|---|---|---|---|---|---|---|---|---|---|---|---|---|---|
| TARA_SAMEA4397472_METAG_ODCNGKGH | 2 | -0.01414320852 | AEGEAN-169 | AEGEAN-169 | | 0.2122116222 | 0.00165069 | 0.04005706303 | 0.01666666667 | 0.05101343926 | 0.01736211372 | 0 | 0.01051504796 | 0 | 0 | 0 |
| TARA_SAMEA4397472_METAG_PLMEHPPN | 1 | -0.1926979947 | Rhodobacterales | UBA10365 | UBA10365 sp003 | 0 | 0 | 1 | 1.69E-04 | 0.001116795408 | 1 | 0 | 0.3070819275 | 0 | 0 | 0.02721285926 |
| TARA_SAMEA4397489_METAG_CGBPAIPC | 8 | -0.140063555 | Flavobacteriales | NORP294 | | 0 | 0 | 0.03609528997 | 1.000000185 | 0.01971973908 | 1.000000185 | 0.04992382026 | 0.0905645908 | 0 | 7.87E-04 | 0 |
| TARA_SAMEA4397489_METAG_EFNJEAPH | 2 | -0.2908004585 | Pseudomonadale | Pseudomonas_E | Pseudomonas_E | 0.003252591245 | 0 | 0.04020331263 | 0.9833333333 | 0.01470201678 | 2.29E-05 | 0 | 0.07678442041 | 0 | 7.85E-04 | 0.001084197082 |
| TARA_SAMEA4397489_METAG_FLBNDNPC | 4 | -0.1292272901 | Flavobacteriales | Cellulophaga | | 0 | 0 | 0.9704729884 | 0.9666712402 | 0.1079537375 | 6.58E-05 | 0 | 0.07685088411 | 0 | 1.63E-04 | 0.03494425063 |
| TARA_SAMEA4397489_METAG_KBFAAALH | 2 | -0.2298364338 | Flavobacteriales | Croceimicrobium | | 0 | 0 | 0.02804775536 | 8.90E-05 | 0 | 6.07E-06 | 0 | 0.1925416841 | 0 | 0.002082879278 | 0 |
| TARA_SAMEA4397489_METAG_KHFCPKOF | 5 | -0.06236994415 | DSM-16500 | 2-12-FULL-45-12 | | 0 | 0 | 0.02270326146 | 1.99E-04 | 0.04597873186 | 1 | 0 | 0 | 0 | 7.41E-04 | 0 |
| TARA_SAMEA4397489_METAG_LJGOPMMD | 8 | -0.1199548499 | Rhodospirillales | UBA11612 | | 0 | 0 | 0.03966851436 | 0.9999999975 | 0.01637559615 | 0.9999999975 | 0 | 0.06072650908 | 0 | 0 | 0 |
| TARA_SAMEA4397489_METAG_NCPEBONE | 1 | -0.05881553103 | Sphingomonadal | Parasphingorhab | Parasphingorhab | 0 | 0 | 1 | 3.65E-06 | 0.003278716462 | 1 | 0 | 0.06879157357 | 0 | 7.77E-04 | 0 |
| TARA_SAMEA4397489_METAG_NELFBNMP | 2 | -0.1891742281 | Caulobacterales | Brevundimonas | | 0.002038842897 | 0 | 0.01857953614 | 0.05 | 0.07993621268 | 1.15E-04 | 0 | 0.008324134732 | 0 | 2.52E-08 | 2.51E-08 |
| TARA_SAMEA4397489_METAG_OBKGIEFA | 7 | -0.1791473149 | Caulobacterales | Henriciella | | 0 | 0 | 1.00000023 | 1.00000023 | 0.966751876 | 0.04699101363 | 0 | 1.00000023 | 0 | 1.00000023 | 6.19E-04 |
| TARA_SAMEA4397489_METAG_OJJKGFOM | 2 | -0.2406540556 | UBA1280 | | | 7.86E-08 | 0 | 0.1225181921 | 0.01666666339 | 0.02131818288 | 0 | 0.01189622363 | 0.06669940162 | 0 | 5.23E-04 | 0.01666666339 |
| TARA_SAMEA4397489_METAG_PCMCDNHN | 1 | -0.03637837036 | Flavobacteriales | JABDMV01 | | 0 | 0 | 1 | 1.74E-04 | 0 | 1 | 0 | 0.01102824639 | 0 | 7.87E-04 | 0 |
| TARA_SAMEA4397489_METAG_PJKGBEKD | 5 | -0.1358206726 | Flavobacteriales | Cellulophaga | | 0.001261441006 | 0 | 0.02281363176 | 8.55E-05 | 0.004191935547 | 1.000000165 | 0 | 0.1292584398 | 0 | 7.97E-04 | 0 |
| TARA_SAMEA4397489_METAG_PKJFCHJO | 5 | -0.3750567984 | Rhodospirillales_ | Thalassospira | Thalassospira all | 4.67E-04 | 0 | 0.02908409733 | 0.01667192628 | 0.1072934793 | 1.000000174 | 0 | 0.08697117337 | 0 | 0.01116274128 | 0.03014781114 |
| TARA_SAMEA4397489_METAG_PMFBIFBE | 2 | -0.2453720346 | Sphingomonadal | Sphingobium | Sphingobium sp0 | 0 | 0 | 0.04562434354 | 1.57E-04 | 0.003198701824 | 1.81E-05 | 0 | 0.0471305314 | 0 | 7.60E-04 | 0 |
| TARA_SAMEA4397516_METAG_GNHIGIFJ | 4 | 5.63E-04 | Pelagibacterales | SYDM01 | | 0 | 0 | 0.9836562089 | 0.01674253108 | 0.2363448635 | 1.64E-04 | 0 | 0.9845118258 | 0 | 0.001578542108 | 0 |
| TARA_SAMEA4397516_METAG_GOKBCAFM | 2 | -0.1157719624 | Flavobacteriales | Hel1-33-131 | Hel1-33-131 sp0 | 0 | 0 | 0.04332952907 | 1.39E-04 | 0 | 0 | 3.00E-06 | 0.07359082172 | 0 | 0 | 0 |
| TARA_SAMEA4397516_METAG_HGNBAOFA | 6 | 0.01032260767 | Flavobacteriales | GCA-002697625 | GCA-002697625 | 0 | 0 | 0.08768384576 | 3.37E-05 | 0.9999997947 | 0.007538294549 | 0 | 0.9999997947 | 0 | 0.001645293657 | 0 |
| TARA_SAMEA4397516_METAG_IHKMLABH | 5 | -0.04856725591 | Flavobacteriales | | | 0 | 0 | 0.02295701069 | 1.25E-04 | 0.04286151527 | 1 | 0 | 0.02870729186 | 0 | 0.001199263797 | 0 |
| TARA_SAMEA4397536_METAG_EOIKDEPL | 5 | -0.1093594808 | Legionellales | JAALKW01 | | 0 | 0 | 0.2347755498 | 1.35E-06 | 0.0599574275 | 0.988942366 | 0 | 0.06821913965 | 0 | 1.67E-04 | 0 |
| TARA_SAMEA4397536_METAG_GKHJOLJD | 2 | -0.1740839559 | Sphingomonadal | Parasphingorhabdus | | 0 | 0 | 0.03616832635 | 1.05E-04 | 0.003364789375 | 0 | 0 | 0.3774364281 | 0 | 0.01345441836 | 0 |
| TARA_SAMEA4397536_METAG_HNHPOKJG | 2 | -0.542229442 | Enterobacterales | Arsukibacterium | | 0 | 0 | 0.1802154529 | 1.60E-04 | 0.008084141511 | 0.007361647423 | 0 | 0.9999998291 | 0 | 7.91E-04 | 0 |
| TARA_SAMEA4397536_METAG_IBGBAAJA | 2 | -0.2429093428 | Rhodobacterales | Sulfitobacter | Sulfitobacter sp0 | 3.50E-07 | 0 | 0.09022232562 | 0.9999998201 | 0.001819103846 | 0.03796015896 | 0 | 0.05952802832 | 5.47E-06 | 0.002411238502 | 0.03690774496 |
| TARA_SAMEA4397536_METAG_MBMFBPNN | 4 | -0.1515486509 | Rhodobacterales | 188UL27-1 | | 1.27E-04 | 0 | 1.000000142 | 0.9833351974 | 0.05969831659 | 0.01940300028 | 0 | 0.053113821 | 0 | 9.71E-05 | 7.66E-08 |
| TARA_SAMEA4397569_METAG_MBIHDIFA | 5 | -0.05591822284 | Rhodobacterales | GCA-2697345 | GCA-2697345 sp | 0 | 0 | 0.03240955719 | 0.01666666667 | 0.2401557403 | 0.9999997623 | 0 | 0.06055561379 | 0 | 0 | 0 |
| TARA_SAMEA4397586_METAG_AALDNBDA | 1 | -0.1535506833 | Burkholderiales | Pusillimonas_A | Pusillimonas_A n | 0 | 0 | 1.000000199 | 1.78E-06 | 0.535351726 | 1.000000199 | 0 | 0.17623938 | 0 | 1.28E-05 | 0.01873728632 |
| TARA_SAMEA4397586_METAG_DHBKNKAI | 2 | -0.186015678 | Pseudomonadale | PZPK01 | | 0.003043881207 | 0 | 0.1249178985 | 4.11E-05 | 0.09288041728 | 7.77E-04 | 0 | 0.06377483499 | 0 | 7.73E-04 | 0 |
| TARA_SAMEA4397586_METAG_KHMNOPDN | 2 | -0.05903409493 | Flavobacteriales | Flavobacterium | | 0 | 0 | 0.110240498 | 0.0166678416 | 0.07790135291 | 0.003176277989 | 0 | 0.9687039147 | 0.02051114367 | 1.12E-05 | 0.0352096658 |
| TARA_SAMEA4397586_METAG_MJKFHMCN | 2 | -0.2145652845 | Cytophagales | Algoriphagus | Algoriphagus aqu | 0 | 0.007430984332 | 0.04000724649 | 8.00E-05 | 1.03E-04 | 0.0216026284 | 0 | 0.1186996512 | 0 | 0.001851735738 | 0.09125505126 |
| TARA_SAMEA4397586_METAG_MLANKLIE | 5 | -0.1110992923 | Sphingomonadal | Sphingopyxis | Sphingopyxis bau | 7.41E-08 | 0 | 0.03320758049 | 0 | 0.00691269181 | 0.9833330503 | 0 | 0.04677806813 | 0 | 8.22E-04 | 0.03619768379 |
| TARA_SAMEA4397586_METAG_NMBCHNOJ | 2 | -0.1249644617 | Caulobacterales | Hyphomonas | | 6.23E-08 | 0 | 0.08938678961 | 8.79E-07 | 0.004938153304 | 0 | 0 | 0.2322669019 | 0 | 1.23E-05 | 0 |
| TARA_SAMEA4397586_METAG_OLNPJHKL | 2 | -0.1567032079 | Verrucomicrobial | Arctic95D-9 | | 0 | 0 | 0.1213464381 | 3.35E-07 | 0.06891024682 | 0.03333333616 | 0 | 0.03449308481 | 0 | 0.02286383062 | 0 |
| TARA_SAMEA4397611_METAG_DKOCJOBL | 7 | -0.0784605355 | Marinisomatales | GCA-2722105 | GCA-2722105 sp | 0 | 0 | 0.03240012123 | 1.87E-04 | 0.003344317149 | 0 | 0 | 0.04598003959 | 0 | 0.999999702 | 0.999999702 |
| TARA_SAMEA4397611_METAG_GJFPHMJM | 5 | -0.1840895123 | Pseudomonadale | UBA9926 | UBA9926 sp905 | 0 | 0 | 0.04286249389 | 1.18E-04 | 5.29E-04 | 0.9833333927 | 0 | 0.09757640024 | 0 | 0.008371925553 | 0.06942945972 |
| TARA_SAMEA4397611_METAG_OBHMADEI | 2 | -0.06595871829 | Pseudomonadales | | | 0 | 0 | 0.1240978828 | 3.39E-05 | 0.003152105479 | 0.003551779659 | 0 | 1 | 0 | 7.47E-04 | 0 |
| TARA_SAMEA4397666_METAG_HIJOACGG | 1 | -0.1794907871 | Sphingomonadal | Qipengyuania | Qipengyuania sp | 0 | 0 | 0.9999998477 | 3.81E-06 | 0.007150329276 | 0.9999998477 | 0.003274516572 | 0.9845188352 | 0 | 3.41E-05 | 0.9999998477 |
| TARA_SAMEA4397666_METAG_KIBCBFEH | 5 | -0.5187654408 | Pseudomonadale | Pseudomonas_E | Pseudomonas_E | 0 | 0 | 0.1904724639 | 1.51E-04 | 0.06930025358 | 0.8835346523 | 0 | 0.07895255921 | 4.43E-06 | 8.85E-05 | 0.06507298667 |
| TARA_SAMEA4397705_METAG_PADFJKCF | 4 | -0.03074687826 | Puniceispirillales | CAJJBL01 | | 1.47E-08 | 1.70E-04 | 0.9999998217 | 0.01666666667 | 0.966666492 | 3.29E-04 | 0 | 0.3263796689 | 0 | 0 | 0 |
| TARA_SAMEA4397730_METAG_CNEJGHKJ | 6 | -0.05478151335 | Marinisomatales | TCS55 | TCS55 sp905182 | 0 | 0 | 0.02301647506 | 1.34E-04 | 0.9999998117 | 0.9999998117 | 3.77E-06 | 0.04446231849 | 0 | 0 | 0 |
| TARA_SAMEA4397730_METAG_NDMAPFEA | 5 | -0.05658163024 | UBA1151 | GCA-2730555 | | 1.59E-07 | 0 | 0.1368987451 | 1.39E-06 | 8.35E-05 | 0.966844629 | 0 | 0.06282287084 | 0.9666669215 | 0.03333333775 | 0 |
| TARA_SAMEA4397747_METAG_FPBHCOLE | 2 | -0.145885435 | Sphingomonadal | Parasphingorhabdus | | 0 | 0 | 0.03121555169 | 1.11E-04 | 0.006891894891 | 0 | 0 | 0.1885662674 | 0 | 8.21E-04 | 0.02039469456 |
| TARA_SAMEA4397747_METAG_HMNAJNKB | 3 | -0.2269740303 | Pseudomonadale | Halomonas | Halomonas maris | 0 | 0 | 0.02966299272 | 1.58E-04 | 0.006557073315 | 1 | 0 | 0.1757846188 | 0 | 1 | 0.06068513123 |
| TARA_SAMEA4397756_METAG_ALFJIJFA | 4 | -0.261991661 | Pseudomonadale | UBA9145 | UBA9145 sp0014 | 0 | 4.28E-05 | 0.9863142179 | 0.01680159438 | 1 | 0.006619957436 | 7.58E-04 | 1 | 0 | 0.02381851034 | 0.1416010925 |
| TARA_SAMEA4397756_METAG_NKKEIPND | 2 | -0.1303260498 | Flavobacteriales | UBA952 | UBA952 sp90518 | 0 | 0 | 0.03932801522 | 0.01666666667 | 0.1378564004 | 0 | 0 | 0.05863324892 | 0 | 0.003211714913 | 0 |
| TARA_SAMEA4397798_METAG_ALIFFGIC | 4 | -0.1029284141 | Pseudomonadale | Halioglobus | Halioglobus sp91 | 1.21E-04 | 0 | 0.983886062 | 1.000000007 | 0.0498733138 | 0.03333333995 | 0 | 0.9840482002 | 0 | 0 | 0 |
| TARA_SAMEA4397798_METAG_MCJEHICI | 5 | -0.1363516633 | PS1 | Pseudothioglobu | Pseudothioglobu | 6.73E-08 | 0 | 0.1155624041 | 0.01666709453 | 0.01990570863 | 0.9833333296 | 0.001099157981 | 0.9999999963 | 0 | 2.58E-05 | 0.006497490086 |
| TARA_SAMEA4397798_METAG_MDFHKJJK | 8 | -0.1874739857 | Rhodobacterales | Planktomarina | Planktomarina sp | 1.40E-04 | 0 | 0.1563524296 | 0.966670142 | 0.04462698255 | 1.000000161 | 0 | 0.06959252306 | 0 | 3.62E-05 | 4.36E-06 |
| TARA_SAMEA4397815_METAG_FBKIEFLJ | 5 | -0.2412089305 | Rhodobacterales | Sulfitobacter | Sulfitobacter sp0 | 0 | 0 | 0.2241755435 | 4.94E-05 | 0.1686801994 | 0.9002717808 | 0 | 0.07226071452 | 0 | 0.002934144708 | 0.01934657767 |
| TARA_SAMEA4397842_METAG_DGOAJPAF | 3 | -0.2040281988 | Flavobacteriales | UBA6772 | | 7.66E-06 | 0 | 1 | 4.79E-07 | 0.004601054586 | 1 | 0 | 0.05475138404 | 0 | 1 | 0 |
| TARA_SAMEA4397842_METAG_EKIJPONI | 3 | -0.1910490911 | Burkholderiales | RS62 | RS62 sp0004964 | 0 | 0 | 0.9999995996 | 1.04E-05 | 0.005587444467 | 0.9999995996 | 0 | 0.9999995996 | 0 | 0.9999995996 | 0 |
| TARA_SAMEA4397842_METAG_ILGPDOJP | 2 | -0.1471034385 | NS11-12g | UBA9320 | | 4.42E-09 | 0 | 0.07921668126 | 0.05000000251 | 0.006396434293 | 0.05000000251 | 0 | 0.9533140111 | 0 | 0.001478667953 | 0.03326991767 |
| TARA_SAMEA4397842_METAG_OIMALLNB | 6 | 0.01365280966 | Pelagibacterales | CACCCN01 | CACCCN01 sp90 | 0 | 0.02161328587 | 0.185296357 | 0.06666766773 | 0.833838969 | 0.1286616585 | 0 | 0.0399427739 | 0 | 0.001812033792 | 0.02462086622 |
| TARA_SAMEA4397859_METAG_BFDFGKDA | 3 | -0.0868938116 | Woeseiales | UBA1847 | | 0 | 0 | 1 | 8.63E-05 | 0.1467253061 | 0.9833333333 | 0 | 0.06745188886 | 0 | 0.983733211 | 0 |
| TARA_SAMEA4397859_METAG_FGINHFMC | 5 | 0.03015744895 | Phycisphaerales | UBA6054 | | 0 | 0 | 0.02052404932 | 1.94E-06 | 6.19E-04 | 0.977934806 | 4.34E-05 | 0.05652892431 | 0 | 1.79E-04 | 0.00364788266 |
| TARA_SAMEA4397859_METAG_GLPLBGPK | 5 | -0.04065011283 | Sphingomonadal | Sphingosinicella | | 0 | 0 | 0.04520117743 | 8.52E-06 | 6.33E-04 | 0.9833954541 | 0 | 0.05943562763 | 0 | 1.05E-04 | 0.03290546508 |
| TARA_SAMEA4397859_METAG_IKEAOLKI | 1 | -0.2019129527 | Pseudomonadale | Halopseudomonas | | 0 | 0 | 1.000000222 | 0 | 0.004045233722 | 0.9669331675 | 0 | 0.06857175207 | 0 | 1.25E-05 | 0.04589060331 |
| TARA_SAMEA4397920_METAG_ABHPMBCP | 2 | -0.06902652171 | Pseudomonadale | HB2-32-21 | | 0 | 1.90E-05 | 0.02100140775 | 5.92E-06 | 0.189954014 | 0.03368784051 | 8.39E-05 | 0.03338826144 | 7.07E-05 | 0.01873929934 | 0.05378944633 |
| TARA_SAMEA4397920_METAG_ADLLGMMD | 8 | -0.1220876956 | Nevskiales | Panacagrimonas | | 0 | 0 | 0.073651063 | 1 | 0.03444616868 | 1 | 0 | 0.05468942659 | 0 | 7.69E-04 | 0 |
| TARA_SAMEA4397920_METAG_AOALJNEL | 5 | -0.2730933609 | Rhodobacterales | Sulfitobacter | Sulfitobacter sp0 | 0 | 0 | 0.03974910287 | 6.38E-05 | 0 | 0.9999998364 | 0 | 0.1733838766 | 0.02316726132 | 7.89E-04 | 0 |
| TARA_SAMEA4397920_METAG_DNIDHFPG | 1 | -0.02121084525 | MPNO01 | | | 0 | 0.005059329816 | 1.000000181 | 3.74E-05 | 0.1583060383 | 1.000000181 | 0 | 0.05990439419 | 0 | 7.46E-04 | 1.39E-04 |
| TARA_SAMEA4397920_METAG_GFPOHCNM | 2 | -0.2216189958 | Rhizobiales | Hoeflea | Hoeflea alexandr | 0 | 0 | 0.02751368774 | 8.36E-05 | 0.001146246836 | 0 | 0 | 0.2296491527 | 0 | 8.20E-04 | 0 |
| TARA_SAMEA4397920_METAG_HCFGADBA | 5 | -0.05411031377 | MPNO01 | | | 0 | 0 | 0.1716978024 | 1.40E-04 | 0.04952940736 | 1 | 0 | 0.08136460634 | 0 | 0.001403167234 | 1.49E-04 |
| TARA_SAMEA4397920_METAG_HMEDABHJ | 2 | -0.2476546231 | Rhodobacterales | Loktanella | | 0 | 0 | 0.02316019692 | 0.008507341546 | 0 | 0 | 0 | 0.1098830824 | 0 | 0.004714608119 | 0.03363080776 |
| TARA_SAMEA4397920_METAG_KIGOFCII | 5 | -0.08500271254 | Flavobacteriales | GCA-002733185 | | 0 | 0 | 0.02829431238 | 0 | 0.2000033503 | 1.000000236 | 0 | 0.05082944256 | 0 | 0.002723165951 | 0 |
| TARA_SAMEA4397930_METAG_FEBNIJHN | 5 | -0.270210554 | Verrucomicrobial | Arctic95D-9 | | 0 | 1.05E-04 | 0.2651232761 | 0.01687996931 | 8.02E-04 | 0.983333668 | 0 | 1.000000301 | 0 | 0.03382784365 | 0 |
| TARA_SAMEA4397930_METAG_NDLBHLEN | 2 | -0.08021600767 | Opitutales | MB11C04 | | 8.55E-04 | 0 | 0.03997200162 | 8.21E-05 | 0.2000511785 | 0.01243147229 | 0 | 0.03728183198 | 0 | 0.01355332517 | 0 |
| TARA_SAMEA4397930_METAG_NILIJJOB | 4 | -0.05518162563 | Burkholderiales | UBA3031 | UBA3031 sp9130 | 0 | 0 | 0.9999998093 | 8.93E-05 | 0.2114851769 | 0.004724988181 | 0.01478370535 | 0.03478938464 | 0 | 0.001474174953 | 0.006259808365 |
| TARA_SAMEA4397930_METAG_NJJBNGCP | 7 | -0.08335239925 | Pseudomonadale | HIMB30 | | 0 | 0 | 0.00832990975 | 0.01667026308 | 0.01111160249 | 0.01666666667 | 0.05735803197 | 0.07724305497 | 1.31E-05 | 1.000000208 | 0 |
| TARA_SAMEA4397930_METAG_ONGLCALM | 7 | -0.2157541438 | Opitutales | Synoicihabitans | Synoicihabitans s | 0 | 0 | 0.9999997884 | 1.60E-04 | 0.9999997884 | 0.005398617231 | 0 | 0.04177464148 | 0 | 0.9999997884 | 0 |
| TARA_SAMEA4397948_METAG_BNMPBJDB | 2 | -0.2026278381 | Sphingomonadal | Parasphingorhab | Parasphingorhab | 0 | 0 | 0.02085543441 | 2.87E-06 | 0.09992371808 | 1.89E-07 | 0 | 0.04634314635 | 0 | 2.70E-05 | 0 |
| TARA_SAMEA4397948_METAG_CJELGJCM | 5 | -0.06197966044 | Rhodospirillales_ | UBA1479 | | 0 | 0 | 0.02235496412 | 0.004069092365 | 6.82E-05 | 1.000000005 | 0 | 0.117175357 | 0 | 3.59E-04 | 0 |
| TARA_SAMEA4397948_METAG_FJPCIEMI | 2 | -0.1161610491 | Flavobacteriales | Ulvibacter | | 0 | 0 | 0.02078339239 | 1.12E-04 | 0 | 0.01666666379 | 0 | 0.2072628259 | 0 | 7.83E-04 | 0 |

| | | | | | | | | | | | | | | | | |
|---|---|---|---|---|---|---|---|---|---|---|---|---|---|---|---|---|
| TARA_SAMEA4397983_METAG_NHFCNOHN | 2 | -0.1503921013 | Flavobacteriales | UBA974 | | 0 | 0 | 0.03570750755 | 4.09E-07 | 0.005073195608 | 0.01317494941 | 0 | 0.9844404426 | 0 | 6.19E-05 | 0 |
| TARA_SAMEA4398001_METAG_DCPOPHPO | 2 | -0.2259114509 | Pseudomonadale | Halopseudomona | Halopseudomona | 0 | 0 | 0.08749102644 | 7.44E-05 | 0.0034424538 | 0.001299670886 | 0 | 0.05917394153 | 0 | 8.16E-04 | 0.04187079558 |
| TARA_SAMEA4398001_METAG_HHDPNOOC | 4 | -0.283096581 | Rhizobiales | SPNT01 | | 0 | 0 | 0.999999995 | 0.9833333532 | 0.01481455988 | 4.50E-05 | 0 | 0.07488928515 | 0 | 4.63E-05 | 0 |
| TARA_SAMEA4398001_METAG_KDFKCPOD | 2 | -0.1273226572 | Caulobacterales | Hyphomonas | | 8.14E-04 | 0 | 0.1704496819 | 2.38E-05 | 0.009300676973 | 0.007168948291 | 0 | 0.06105002368 | 0 | 7.90E-04 | 0 |
| TARA_SAMEA4398009_METAG_CAOMJLNE | 6 | -0.01198883707 | Burkholderiales | BACL14 | BACL14 sp9130 | 0 | 0 | 0.3045331434 | 4.53E-05 | 1 | 1 | 0 | 1 | 0 | 1.16E-05 | 0 |
| TARA_SAMEA4398009_METAG_LEDOCBPK | 7 | -0.08883254818 | Pseudomonadale | UBA4421 | UBA4421 sp9051 | 0 | 0 | 1 | 0.01674790225 | 0.00298836571 | 4.41E-06 | 0 | 0.2590115441 | 0 | 1 | 0.02623644434 |
| TARA_SAMEA4398027_METAG_BKLIPBOK | 5 | -0.1907613002 | Flavobacteriales | UBA2040 | | 0 | 0 | 0.02131071421 | 3.39E-05 | 0.003375163327 | 1.000000215 | 0 | 0.2768837195 | 0 | 0 | 0 |
| TARA_SAMEA4398027_METAG_DOLBIGEE | 5 | -0.1058176076 | MPNO01 | | | 0 | 0.001317302635 | 0.03753598843 | 9.45E-05 | 0.005173303579 | 1.000000068 | 0 | 0.6070763461 | 0 | 0.4337729687 | 8.57E-04 |
| TARA_SAMEA4398027_METAG_JFPJLACK | 2 | -0.1928465733 | Flavobacteriales | Maribacter | | 0.02468699912 | 0 | 0.03350125269 | 6.27E-05 | 0 | 0 | 0 | 0.0317635613 | 0 | 0 | 0 |
| TARA_SAMEA4398027_METAG_PHDMBMMK | 7 | -0.1113591382 | Chitinophagales | CAJXUU01 | | 0 | 0 | 0.03322166488 | 1 | 0.2844702366 | 0.01941594538 | 0 | 0.06267633174 | 0 | 1 | 0.04224130363 |
| TARA_SAMEA4398067_METAG_CAMNPADC | 4 | -0.1607960803 | Flavobacteriales | Arcticimaribacter | Arcticimaribacter | 0 | 0 | 0.9838271556 | 0.01858793498 | 1.45E-04 | 7.78E-09 | 0 | 0.05258205323 | 0 | 1.30E-05 | 0 |
| TARA_SAMEA4398067_METAG_CBDGHFLB | 5 | -0.1669097186 | Pseudomonadale | UBA4582 | UBA4582 sp9051 | 0 | 0 | 0.1546401757 | 1.23E-04 | 0 | 1 | 0 | 1 | 0 | 7.47E-04 | 0 |
| TARA_SAMEA4398067_METAG_KGGLPANE | 8 | -0.2958089344 | Enterobacterales | JALJPB01 | | 5.89E-04 | 0 | 0.09550179174 | 0.9833332466 | 0.007175405391 | 0.999999823 | 5.31E-04 | 0.999999823 | 2.96E-06 | 3.06E-05 | 0.0349814004 |
| TARA_SAMEA4398094_METAG_EINFCOHD | 1 | -0.1196958051 | Flavobacteriales | UA16 | | 0 | 0 | 0.9999998087 | 3.79E-05 | 0.003500367579 | 0.9999998087 | 0 | 0.05779222034 | 0 | 8.25E-04 | 0.9999998087 |
| TARA_SAMEA4398094_METAG_FKONMKBF | 5 | -0.1625284762 | Sphingomonadal | Pseudemcibacte | Pseudemcibacte | 0 | 0.005291922738 | 0.03211693109 | 2.38E-04 | 0.04498314325 | 1 | 0 | 0.07312813233 | 0 | 0 | 0 |
| TARA_SAMEA4398094_METAG_HLGDABHH | 2 | -0.1144422097 | Pseudomonadale | UBA9145 | | 0 | 0 | 0.02051433677 | 0.03333333333 | 0.3467799713 | 0 | 0 | 0.9679009417 | 0 | 2.41E-04 | 0.03434880515 |
| TARA_SAMEA4398094_METAG_OJPBDFHD | 2 | -0.1287797249 | Flavobacteriales | UBA8316 | UBA8316 sp9051 | 0 | 0 | 0.02233342393 | 1 | 0.005156008917 | 5.45E-06 | 0 | 0.0515134805 | 0 | 7.82E-04 | 0 |
| TARA_SAMEA4398111_METAG_GFHMNJCF | 5 | -0.2037985249 | Rhodobacterales | Pseudosulfitobac | Pseudosulfitobac | 0 | 0 | 0.02342337323 | 8.06E-05 | 0.09128314278 | 1 | 0 | 0.07183616008 | 0 | 7.85E-04 | 0 |
| TARA_SAMEA4398111_METAG_JEHKNIFP | 2 | -0.2384143482 | Pseudomonadale | Marinomonas | | 0 | 0 | 0.04476746243 | 1.01E-04 | 4.24E-04 | 0 | 0 | 0.04720337684 | 0 | 0 | 0 |
| TARA_SAMEA4398111_METAG_MPNOCJLE | 2 | -0.2299941159 | Flavobacteriales | Muricauda | Muricauda sp002 | 0 | 0 | 0.03972777846 | 0 | 0.001693334519 | 0.004899798732 | 0 | 0.05183458851 | 0 | 8.46E-04 | 0 |
| TARA_SAMEA4398168_METAG_EEEDAEIF | 2 | -0.2380412748 | Pseudomonadale | Halopseudomona | Halopseudomona | 0 | 0 | 0.01496798597 | 0.03333752522 | 0.175564165 | 0 | 0 | 0.03745776879 | 0.01666666183 | 4.01E-05 | 0.003639789544 |
| TARA_SAMEA4398193_METAG_GAPAJFEN | 4 | -0.00444574501 | Phycisphaerales | GCA-2732755 | | 0 | 0 | 0.9704506523 | 2.97E-05 | 0.009183788512 | 0.03799817573 | 0 | 0.06701910003 | 0 | 8.11E-04 | 0.3163642209 |
| TARA_SAMEA4398219_METAG_MOHHODOE | 3 | -0.1330950822 | UBA796 | UBA796 | | 0 | 0 | 0.03254465949 | 1.33E-04 | 0.08631675465 | 1 | 0.04719817475 | 0.09428428184 | 0 | 1 | 0.06975111867 |
| TARA_SAMEA4398219_METAG_PMMHAMPC | 4 | -0.230475202 | Flavobacteriales | UBA3442 | UBA3442 sp0023 | 0 | 0 | 0.999999789 | 9.65E-05 | 0.01311920798 | 8.90E-04 | 0 | 0.07271824182 | 0 | 0 | 0.154300235 |
| TARA_SAMEA4398238_METAG_AANIECHH | 8 | -0.2971390696 | Enterobacterales | Paraglaciecola | | 0 | 0 | 0.9999998054 | 0.9999998054 | 0.00341629902 | 0.9999998054 | 0 | 0.9999998054 | 0 | 9.31E-04 | 0.0417063279 |
| TARA_SAMEA4398238_METAG_BBIMKEDM | 5 | -0.08324447671 | Francisellales | | | 0 | 0 | 0.04739967972 | 1.14E-05 | 0 | 1 | 0 | 0.06301291108 | 0 | 4.16E-05 | 0.148629449 |
| TARA_SAMEA4398238_METAG_CLIBKFHJ | 2 | -0.3078824893 | Enterobacterales | Paraglaciecola | Paraglaciecola sp | 0 | 0 | 0.0293227679 | 0 | 0.009523519085 | 0.03333333675 | 0 | 0.0542912711 | 0 | 0 | 0.03367827072 |
| TARA_SAMEA4398238_METAG_DNPPHHEA | 6 | -0.02968401425 | GCA-2731375 | | | 0 | 0 | 0.1481224486 | 6.95E-05 | 1 | 0 | 0 | 0.06659552981 | 0 | 1.57E-05 | 0.03943008036 |
| TARA_SAMEA4398238_METAG_GONOIKHO | 7 | -0.1711350299 | Sphingomonadal | Novosphingobiur | Novosphingobiur | 0 | 0 | 0.03755005386 | 0 | 3.04E-04 | 0.001785055499 | 0 | 0.2485525721 | 0 | 0.9999998264 | 0 |
| TARA_SAMEA4398238_METAG_IPMCJFCD | 5 | -0.1463071807 | Opitutales | Synoicihabitans | | 0 | 0 | 0.05633751941 | 1.52E-04 | 0.09528208855 | 0.9999998555 | 0.05130816295 | 0.06206013179 | 0.01207201597 | 0.02033878224 | 3.94E-04 |
| TARA_SAMEA4398238_METAG_KPHFHLOE | 4 | -0.2138500543 | Pirellulales | Novipirellula | | 8.99E-07 | 0 | 1 | 3.84E-04 | 0.1013888762 | 0.01666666667 | 0 | 0.05290314634 | 0 | 0.002892644316 | 0 |
| TARA_SAMEA4398238_METAG_MBGMDCJJ | 2 | -0.1973913925 | Caulobacterales | Maricaulis | | 0 | 0 | 0.02155586246 | 1.98E-06 | 0.001702267689 | 0.003620893145 | 0 | 0.2080843894 | 0 | 0 | 0 |
| TARA_SAMEA4398238_METAG_MKMOEGDP | 6 | -0.2106767138 | Pseudomonadale | Halomonas | Halomonas glaci | 0 | 0 | 0.04327387225 | 8.07E-05 | 1 | 0 | 0 | 0.06116628426 | 0 | 0 | 0.03799642418 |
| TARA_SAMEA4398247_METAG_MCKJBFJL | 2 | -0.08325636885 | Flavobacteriales | Polaribacter | Polaribacter sp90 | 0 | 0 | 0.03115427802 | 8.67E-05 | 0.042278993 | 0.4333333333 | 0 | 0.05848168737 | 0 | 8.00E-04 | 0 |
| TARA_SAMEA4398266_METAG_CIJKCIBB | 4 | -0.05956540934 | JAGOUU01 | | | 0 | 0 | 1.000000268 | 0 | 0.1074229308 | 0 | 0 | 1.000000268 | 0 | 0 | 0 |
| TARA_SAMEA4398266_METAG_FKIOLFBB | 6 | -0.1579899138 | Flavobacteriales | Dokdonia | | 0 | 0 | 0.03909882443 | 1.33E-06 | 0.9833331598 | 0.001254138616 | 0 | 0.325028558 | 0 | 0 | 0.02928528578 |
| TARA_SAMEA4398266_METAG_KLOMINHN | 5 | -0.1018228544 | Flavobacteriales | Maribacter | Maribacter stanie | 0 | 0 | 0.05944092664 | 7.65E-05 | 0.001208974099 | 0.9833333333 | 0 | 0.1473955375 | 0 | 0 | 2.90E-05 |
| TARA_SAMEA4398296_METAG_BLGABLEO | 1 | -0.1745593605 | Chitinophagales | JAFMKE01 | | 0 | 0 | 1 | 1.94E-04 | 0 | 1 | 0 | 0.2848228494 | 0 | 7.86E-04 | 0 |
| TARA_SAMEA4398296_METAG_PCOLMPAO | 4 | -0.1121542064 | Flavobacteriales | SCGC-AAA160-F | SCGC-AAA160-F | 0 | 0 | 0.9999998514 | 1.15E-04 | 0.01985970225 | 1.35E-07 | 0 | 0.06809453674 | 5.15E-07 | 0 | 2.20E-05 |
| TARA_SAMEA4398315_METAG_ANIFHKHO | 2 | -0.2607692412 | Pseudomonadale | Alcanivorax | Alcanivorax mobi | 0 | 0 | 0.04568233892 | 7.51E-05 | 0.03982386474 | 0 | 0 | 0.07165649548 | 0 | 0 | 0.03697665686 |
| TARA_SAMEA4398315_METAG_BJAKHDDB | 4 | 0.01565357954 | UBA6615 | | | 0 | 0 | 0.9999997691 | 0.9999997691 | 0.05154041857 | 2.08E-05 | 0 | 0.06689761033 | 0 | 0.001218377549 | 0.04727861426 |
| TARA_SAMEA4398315_METAG_BPOJMCNG | 2 | -0.1474845841 | Rhodobacterales | Paracoccus | Paracoccus marc | 4.01E-08 | 0 | 0.0365032669 | 0.03333333333 | 0.001486771579 | 0.03333333333 | 0 | 0.08956467868 | 0 | 2.58E-05 | 1.63E-07 |
| TARA_SAMEA4398315_METAG_HKMENHED | 6 | -0.240607561 | Pseudomonadale | Psychrobacter | Psychrobacter ni | 0 | 2.40E-04 | 0.03922182857 | 0.06666666598 | 0.9504253025 | 1.01E-04 | 0 | 0.08543963504 | 0 | 0 | 0 |
| TARA_SAMEA4398315_METAG_IKMBHIOE | 2 | -0.3536699474 | Cytophagales | Imperialibacter | | 0 | 0 | 0.03557229487 | 1.71E-05 | 0.003258410744 | 0 | 0 | 0.04552720988 | 0 | 0.01895163842 | 0 |
| TARA_SAMEA4398315_METAG_JIDPDINC | 2 | -0.49810583 | Enterobacterales | Pseudoalteromor | Pseudoalteromor | 0 | 0 | 0.07198770896 | 0.01688407306 | 0.06738131901 | 0 | 8.16E-08 | 0.02987632412 | 0 | 0 | 0 |
| TARA_SAMEA4398315_METAG_OCCFADGL | 7 | -0.05110496474 | Flavobacteriales | Polaribacter | | 0 | 0 | 0.03914864838 | 1.57E-04 | 0.0726599467 | 0 | 0 | 0.03887041008 | 0 | 1 | 0.06414977171 |
| TARA_SAMEA4398315_METAG_PALEGFHE | 5 | -0.05187959323 | UBA8366 | MarineAlpha4-Bin1 | | 0 | 0 | 0.02563236844 | 1.13E-04 | 0.04673150782 | 1.000000293 | 0 | 0.06957424197 | 0 | 0.01498893093 | 0 |
| TARA_SAMEA4398315_METAG_POBJPKPL | 1 | -0.1794703016 | Sphingomonadal | Parasphingorhabdus | | 1.25E-07 | 0 | 1.000000187 | 1.27E-04 | 0.001148787688 | 0.9834450414 | 0 | 0.2899086032 | 2.99E-08 | 0.007562552439 | 0.05829847001 |
| TARA_SAMEA4398360_METAG_AEECNLEP | 2 | -0.2156257813 | Cytophagales | Roseivirga | | 0 | 0 | 0.02920069823 | 1 | 0.003631230739 | 0 | 0 | 0.05419189851 | 0 | 0.001431899757 | 0 |
| TARA_SAMEA4398360_METAG_BPNDFBAG | 6 | -0.03983317205 | Oceanibaculales | Oceanibaculum | Oceanibaculum r | 0 | 0 | 0.03970788183 | 1.78E-04 | 1 | 0 | 0 | 0.007087161633 | 0 | 0 | 0 |
| TARA_SAMEA4398360_METAG_CBLJDGJE | 5 | -0.242381644 | Balneolales | Balneola | Balneola sp9130 | 0 | 0 | 0.07455550483 | 6.66E-05 | 0.00442684403 | 1.000000231 | 0 | 0.03609185343 | 0 | 7.67E-04 | 0 |
| TARA_SAMEA4398360_METAG_FOIOEKLJ | 5 | -0.224420803 | Pseudomonadale | Zhongshania | | 0 | 0 | 0.1458804695 | 4.47E-06 | 0.01689786014 | 1.00000023 | 0 | 0.03685720893 | 0 | 2.81E-04 | 3.66E-04 |
| TARA_SAMEA4398360_METAG_GEECGHMB | 5 | -0.3257490558 | Enterobacterales | Cognaticolwellia | Cognaticolwellia | 0 | 0 | 0.02897120649 | 0 | 0.01207864593 | 0.9999998014 | 0 | 0.2591773593 | 0 | 1.75E-05 | 0 |
| TARA_SAMEA4398360_METAG_GJHCJNNP | 2 | -0.1395347042 | Burkholderiales | CAISIP01 | | 2.44E-04 | 0.001211652727 | 0.03330993941 | 0.0168241412 | 0.004582846051 | 0.05059810064 | 0 | 0.2499214259 | 0 | 6.93E-04 | 0.01051431001 |
| TARA_SAMEA4398370_METAG_CFILGFGC | 3 | -0.06151080285 | Woeseiales | SP4260 | | 0 | 0 | 0.9999999938 | 0.03344274268 | 0.9668380044 | 0.9834134558 | 3.46E-05 | 0.06482627993 | 0 | 0.9999999938 | 0.0457298807 |
| TARA_SAMEA4398370_METAG_EHKIJLNC | 3 | -0.01432386478 | TMED25 | CAJIYR01 | CAJIYR01 sp018 | 0 | 0 | 1 | 7.55E-05 | 0.493697225 | 1 | 0 | 1 | 0 | 1 | 0 |
| TARA_SAMEA4398370_METAG_ILDJJAMF | 5 | 0.003855155567 | SAR86 | GCA-2707915 | GCA-2707915 sp | 0 | 0 | 0.004101087818 | 3.18E-07 | 0 | 1 | 0 | 0.1800736908 | 0 | 0.003643591149 | 0 |
| TARA_SAMEA4398387_METAG_EACEGFLE | 2 | -0.1638941969 | Flavobacteriales | | | 0 | 0.009550182784 | 0.06182007564 | 7.64E-05 | 0.00682689143 | 0 | 0 | 0.1625543656 | 0 | 0.005413291871 | 0 |
| TARA_SAMEA4398387_METAG_GHIDNKEH | 5 | -0.3597960695 | Enterobacterales | Psychromonas | | 6.74E-08 | 0 | 0.08981762435 | 0.01129775094 | 8.26E-05 | 0.9833331844 | 0 | 0.04084636517 | 0 | 0.002010248041 | 0 |
| TARA_SAMEA4398387_METAG_ILGDMCFO | 2 | -0.1291316039 | Thalassobaculale | Nisaea | | 0 | 0 | 0.223923579 | 5.76E-05 | 0 | 0.003678327884 | 0 | 1 | 0 | 0 | 0 |
| TARA_SAMEA4398387_METAG_KHFOIAHK | 6 | -0.1548997088 | Cytophagales | Cyclobacterium | | 0 | 0 | 0.05030688665 | 7.86E-07 | 1 | 8.64E-05 | 0 | 0.07881429655 | 0 | 0.0195254834 | 0 |
| TARA_SAMEA4398387_METAG_MKDFJCJD | 5 | -0.1109597251 | Pseudomonadale | PZPK01 | | 0 | 0.003596855931 | 0.04097985211 | 1.23E-04 | 0 | 1 | 0 | 0.2255452017 | 0 | 0 | 0 |
| TARA_SAMEA4398387_METAG_NJONMNKJ | 7 | -0.2904877441 | Enterobacterales | Colwellia | | 0 | 0 | 0.04099254581 | 2.02E-06 | 0.04716184075 | 8.62E-04 | 0 | 0.07959022196 | 0 | 0.9499997329 | 0.06343256161 |
| TARA_SAMEA4398387_METAG_OKMNOAGP | 5 | -0.04838135424 | Flavobacteriales | Polaribacter | | 0 | 3.82E-05 | 0.1354950201 | 3.51E-06 | 0.03076111091 | 0.9833331354 | 0 | 0.1112379757 | 0 | 0.02230136225 | 0 |
| TARA_SAMEA4398387_METAG_PBBIBPIA | 3 | -0.2922183422 | Pseudomonadale | Oceanobacter | Oceanobacter sp | 0 | 0 | 0.03371617129 | 1.32E-04 | 0.02603511177 | 1.00000025 | 0.01882291495 | 0.08980892722 | 0 | 1.00000025 | 0 |
| TARA_SAMEA4398413_METAG_BBKHKGHP | 3 | -0.00721324758 | Pelagibacterales | AG-414-E02 | | 3.59E-06 | 0 | 0.04086938498 | 1 | 0.2569402594 | 1 | 0 | 0.04962026408 | 0 | 1 | 0.0369749802 |
| TARA_SAMEA4398413_METAG_MPJICBCG | 7 | -0.06931624125 | SCGC-AAA003-L | JABHHG01 | JABHHG01 sp01 | 0 | 0 | 1.000000193 | 1.16E-04 | 0.002593962277 | 0.002380944585 | 0 | 1.000000193 | 0 | 1.000000193 | 0.05557334983 |
| TARA_SAMEA4398413_METAG_PHKIEMHL | 2 | -0.02147573311 | Acidimicrobiales | UBA8592 | UBA8592 sp0224 | 0 | 0 | 0.04350455984 | 1.02E-04 | 0.3222224896 | 0.009723832139 | 0 | 0.06141943566 | 0 | 7.16E-07 | 0 |
| TARA_SAMEA4398432_METAG_DPPBFELL | 5 | -0.3792796113 | Enterobacterales | Psychromonas | | 0 | 0 | 0.04118427068 | 8.26E-05 | 0.1386704673 | 1.000000004 | 0.01451962305 | 0.063428454 | 0 | 0.001582181835 | 0.02617560144 |
| TARA_SAMEA4398432_METAG_GFLEHPMN | 6 | -0.2396094422 | Burkholderiales | Azoarcus_C | Azoarcus_C com | 0 | 0 | 0.02803204194 | 8.19E-05 | 1.000000217 | 3.47E-06 | 0 | 0.0718866816 | 0 | 0 | 0 |
| TARA_SAMEA4398432_METAG_GMCKOCBG | 4 | -0.1615761525 | Flavobacteriales | Maribacter_A | Maribacter_A lute | 0 | 0 | 1 | 7.64E-05 | 0 | 0 | 0 | 0.2187875264 | 0 | 0 | 0.008983090682 |
| TARA_SAMEA4398432_METAG_HCGEJLHD | 1 | -0.06891135522 | Bacteroidales | Ancylomarina | | 0 | 0 | 1.0000002 | 6.22E-05 | 0.128536343 | 1.0000002 | 0 | 0.04980265934 | 0 | 8.04E-04 | 0 |
| TARA_SAMEA4398432_METAG_LDDJDMHE | 1 | -0.02110496111 | Flavobacteriales | Lutibacter | | 0 | 0 | 0.9999999949 | 5.03E-06 | 0.08736338638 | 0.6875491216 | 0 | 0.04766754562 | 0 | 2.81E-04 | 0 |

| | | | | | | | | | | | | | | | | |
|---|---|---|---|---|---|---|---|---|---|---|---|---|---|---|---|---|
| TARA_SAMEA4398432_METAG_LPCKCMMB | 1 | -0.2387007185 | Caulobacterales | Hyphomonas | Hyphomonas sp0 | 2.67E-06 | 0.02391605273 | 0.9851195135 | 1.60E-04 | 0.01356176393 | 1 | 0.002220525067 | 0.984509563 | 0.001329212127 | 0.001904048839 | 0.009403424076 |
| TARA_SAMEA4398432_METAG_MGKICFLD | 2 | -0.3654616648 | Rhodobacterales | Celeribacter | Celeribacter bael | 0 | 0 | 0.02358869117 | 2.92E-08 | 0.00350098285 | 0.005433757977 | 0 | 0.9720401181 | 0 | 7.93E-04 | 0.01666666667 |
| TARA_SAMN05326640_METAG_ANE00014 | 8 | 0.04031557402 | UBA1144 | TMED126 | | 1.55E-04 | 0 | 0.07051038135 | 1.00000019 | 0.0146003275 | 1.00000019 | 0 | 0.06981760974 | 0 | 0.002325909003 | 0.03823387954 |
| TARA_SAMN05326641_METAG_ANW00005 | 2 | -0.09225365342 | Methylococcales | Cycloclasticus | Cycloclasticus sp | 0 | 0 | 0.02881293996 | 1.78E-06 | 6.77E-04 | 0 | 0 | 0.07973374021 | 0 | 0.0119929679 | 0.335372808 |
| TARA_SAMN05326641_METAG_ANW00009 | 7 | -0.1179989282 | Pseudomonadale | Halioglobus | Halioglobus sp0 | 0 | 0 | 1.000000399 | 1.35E-04 | 1.000000399 | 0.004600477782 | 0 | 0.05072987351 | 0 | 1.000000399 | 0.006144621479 |
| TARA_SAMN05326641_METAG_ANW00010 | 5 | -0.2529985918 | Flavobacteriales | Aequorivita | Aequorivita aquir | 0 | 0 | 0.09987976573 | 1.12E-04 | 0.003333262769 | 1 | 0 | 0.07650752998 | 0 | 7.92E-04 | 0 |
| TARA_SAMN05326641_METAG_ANW00013 | 2 | 0.003123692252 | SCGC-AAA003-L08 | | | 5.73E-06 | 0 | 0.03218136206 | 1.59E-06 | 0.223840639 | 0.08303710669 | 0.002658395254 | 0.08446727619 | 0 | 0.004566986719 | 8.73E-05 |
| TARA_SAMN05326641_METAG_ANW00019 | 2 | -0.08576083907 | Pseudomonadale | Ketobacter | Ketobacter sp002 | 0 | 0 | 0.1789649865 | 4.39E-05 | 0.07024497153 | 0.005948357591 | 0 | 1 | 0 | 7.96E-04 | 0 |
| TARA_SAMN05326641_METAG_ANW00032 | 1 | -0.0454425055 | Rhodobacterales | GCA-002705045 | | 0 | 0 | 1.000000287 | 2.06E-06 | 0.01315207591 | 1.000000287 | 0 | 0.101210582 | 0 | 0.0338831578 | 0.09174305942 |
| TARA_SAMN05326641_METAG_ANW00035 | 2 | -0.1108107684 | Pirellulales | UBA1268 | | 0 | 0 | 0.03332522294 | 1.60E-06 | 0.01646048456 | 0 | 0 | 0.03882686904 | 0 | 4.14E-05 | 0 |
| TARA_SAMN05326643_METAG_ASW00001 | 5 | -0.01151570613 | Pseudomonadale | UBA7434 | UBA7434 sp002 | 3.01E-06 | 0 | 0.02685249766 | 8.82E-05 | 0 | 1 | 8.32E-04 | 0.179404133 | 0 | 8.01E-04 | 0.03157109224 |
| TARA_SAMN05326643_METAG_ASW00003 | 1 | -0.06140672635 | Cyanobacteriales | Atelocyanobacte | Atelocyanobacte | 0 | 0 | 1 | 1.49E-04 | 0 | 1 | 2.25E-06 | 0.05487800631 | 0 | 0 | 0.03769115752 |
| TARA_SAMN05326644_METAG_PON00004 | 5 | -0.05993906674 | Coxiellales | UBA9148 | UBA9148 sp002 | 0 | 0 | 0.03161136928 | 1.56E-04 | 0.01273550821 | 1.000000206 | 0 | 0.03604027265 | 0 | 0.01489611755 | 0 |
| TARA_SAMN05326644_METAG_PON00018 | 7 | -0.0227972218 | Comchoanobacte | UBA7916 | UBA7916 sp002 | 0.001766417568 | 0 | 1.000000199 | 6.59E-07 | 1.000000199 | 0.01666666337 | 0 | 0.3200469677 | 0 | 1.000000199 | 0 |
| TARA_SAMN05326644_METAG_PON00029 | 2 | -0.1152982296 | Verrucomicrobial | SW10 | SW10 sp002729 | 0 | 0 | 0.03900563176 | 0 | 0 | 8.38E-05 | 0 | 0.06590659336 | 0 | 7.75E-07 | 0 |
| TARA_SAMN05326645_METAG_PSE00001 | 2 | -0.2831545473 | Balneolales | UBA7797 | UBA7797 sp002 | 0 | 0 | 0.02129152991 | 0 | 4.50E-05 | 0.01666666177 | 0 | 0.04824030619 | 0 | 2.35E-04 | 0 |
| TARA_SAMN05326645_METAG_PSE00002 | 6 | -0.1767646072 | Caulobacterales | Maricaulis | Maricaulis sp002 | 1.16E-06 | 0 | 0.02685614675 | 1.54E-04 | 0.9999997088 | 4.47E-04 | 0 | 0.0167452754 | 0 | 5.82E-07 | 0 |
| TARA_SAMN05326645_METAG_PSE00009 | 2 | -0.2718977116 | Flavobacteriales | Croceimicrobium | Croceimicrobium | 0 | 0 | 0.1328070644 | 9.83E-05 | 0.01328195507 | 0.005386542728 | 0 | 0.02786579683 | 0 | 7.59E-04 | 0 |
| TARA_SAMN05326645_METAG_PSE00010 | 1 | -0.1210209867 | Rhizobiales | Stappia | Stappia sp00270 | 0 | 0 | 1.000000286 | 1.35E-06 | 0.4332939015 | 1.000000286 | 0 | 0.04088669788 | 0.001854570536 | 0.01342275363 | 0 |
| TARA_SAMN05326645_METAG_PSE00013 | 2 | -0.1211840746 | Flavobacteriales | Winogradskyella | Winogradskyella | 0 | 0 | 0.03305517061 | 0.01671440533 | 5.31E-05 | 0.01667315093 | 0 | 0.2617953257 | 0 | 0.007795171524 | 0 |
| TARA_SAMN05326645_METAG_PSE00015 | 2 | 0.01332794178 | Pseudomonadale | UBA7803 | UBA7803 sp002 | 0.01740047198 | 0 | 0.02260676211 | 3.92E-05 | 0.004476518711 | 6.36E-06 | 0 | 0.0619958196 | 0 | 3.28E-05 | 0.001769365312 |
| TARA_SAMN05326645_METAG_PSE00020 | 1 | -0.05056066005 | Flavobacteriales | GCA-2862585 | | 0 | 0 | 0.9999997784 | 1.08E-04 | 0.9999997784 | 0.9999997784 | 0 | 0.05762528109 | 0 | 0.001819998312 | 0 |
| TARA_SAMN05326645_METAG_PSE00023 | 5 | -0.0291985409 | Phycisphaerales | UBA1926 | UBA1926 sp002 | 0 | 0 | 0.03222224211 | 0.01676563413 | 0.01898629857 | 0.983820196 | 0 | 0.07525073216 | 0.001376569642 | 7.65E-04 | 0 |
| TARA_SAMN05326645_METAG_PSE00025 | 2 | -0.1090476905 | Rhodobacterales | LFER01 | LFER01 sp9026 | 0 | 0 | 0.02127695899 | 1.38E-05 | 0.009372056751 | 0 | 0 | 0.2901227969 | 0 | 0.01108541636 | 0 |
| TARA_SAMN05326645_METAG_PSE00044 | 3 | -0.02065694843 | Verrucomicrobial | Roseibacillus_B | | 0 | 0 | 1.000000356 | 1.07E-05 | 0.004021251628 | 1.000000356 | 0 | 0.07644222094 | 0 | 1.000000356 | 0 |
| TARA_SAMN05326645_METAG_PSE00073 | 5 | 0.01530857114 | Pelagibacterales | Pelagibacter | | 1.37E-07 | 0 | 0.0248414697 | 3.23E-06 | 0.0032080472 | 0.9835935844 | 0 | 0.06841193655 | 0 | 0.01100527944 | 4.42E-04 |
| TARA_SAMN05326646_METAG_PSW00002 | 7 | -0.06935122494 | Rhodospirillales_ | GCA-2717285 | GCA-2717285 sp | 0 | 0.00505260541 | 0.02372384423 | 4.76E-05 | 0.1226752489 | 0 | 0 | 0.03643291138 | 0 | 1.00000027 | 0 |
| TARA_SAMN05326646_METAG_PSW00011 | 4 | -0.03721070026 | Chromatiales | JAHDBG01 | | 0.004474356389 | 0 | 0.9695423046 | 3.85E-06 | 0.9859123046 | 0 | 0 | 0.01926162267 | 0 | 3.97E-05 | 0 |
| TARA_SAMN05326646_METAG_PSW00012 | 5 | -0.1369680799 | Flavobacteriales | GCA-002722245 | GCA-002722245 | 0 | 0 | 0.06833088045 | 1.91E-04 | 0.007011225466 | 1.000000242 | 0 | 0.07715785837 | 0 | 0 | 0 |
| TARA_SAMN05326646_METAG_PSW00016 | 2 | -0.1649074937 | Bdellovibrionales | UBA6776 | UBA6776 sp002 | 0 | 0 | 0.02702408835 | 0.005078581395 | 0.08825975256 | 4.32E-06 | 0 | 0.0400527875 | 0 | 0 | 0 |
| TARA_SAMN05326646_METAG_PSW00017 | 8 | -0.05795432334 | Mycobacteriales | Mycobacterium | Mycobacterium p | 0 | 1.22E-04 | 0.04023666024 | 0.9500024276 | 0 | 0.9833333364 | 0 | 0.09683972896 | 0 | 6.90E-04 | 2.94E-06 |
| TARA_SAMN05326646_METAG_PSW00020 | 4 | 0.02723631244 | Ferrovibrionales | | | 0.08687371025 | 0 | 0.9999997901 | 5.55E-06 | 0.002666557424 | 1.87E-04 | 0 | 0.06007722112 | 0 | 5.14E-05 | 0 |
| TARA_SAMN05326646_METAG_PSW00024 | 1 | -0.1631748867 | UBA1280 | | | 0.09048140503 | 0 | 1 | 0.01666666667 | 1 | 0.9841521696 | 0.02753470546 | 0.9841439304 | 0 | 0.006713107224 | 0.001208655278 |
| TARA_SAMN05326646_METAG_PSW00037 | 7 | -0.0469980635 | Ga0077536_A | TMED69 | | 0 | 0 | 5.61E-04 | 9.10E-07 | 0.001723688995 | 7.75E-08 | 0 | 0.001104889613 | 0 | 0.9999999952 | 0 |
| TARA_SAMN05326646_METAG_PSW00059 | 7 | -0.00841956418 | TMED109_A | GCA-2720125 | GCA-2720125 sp | 0 | 1.92E-04 | 0.9837531368 | 0.01680200725 | 0.1276382169 | 0.01666666667 | 0 | 0.9843379886 | 0 | 1.000000297 | 0 |
| TARA_SAMN05326646_METAG_PSW00077 | 8 | -0.03356874371 | Puniceispirillales | HIMB100 | HIMB100 sp0027 | 0 | 0 | 1.000000185 | 0.9833369369 | 0.983333518 | 0.9674448872 | 9.39E-04 | 0.1789324153 | 0 | 7.43E-04 | 0.05608671793 |
| TARA_SAMN05326646_METAG_PSW00093 | 5 | -0.1102191907 | UBA6191 | | | 8.09E-08 | 0 | 0.02611149381 | 0 | 0.1909823201 | 1.000000004 | 0 | 0.04616412118 | 0 | 1.23E-04 | 0 |
| TARA_SAMN05326647_METAG_ION00002 | 2 | -0.2155041696 | Caulobacterales | Brevundimonas | Brevundimonas a | 0 | 0 | 0.03354358558 | 3.43E-05 | 0.001250055301 | 0.001514172627 | 0 | 0.05821015947 | 0 | 0.009140535304 | 0.1556217911 |
| TARA_SAMN05326647_METAG_ION00006 | 3 | 0.02674759468 | Flavobacteriales | GCA-2716345 | | 0 | 0 | 0.0478101466 | 2.77E-07 | 0.9875801187 | 0.9999999947 | 0 | 0.04135675123 | 0 | 0.9833333333 | 0 |
| TARA_SAMN05326647_METAG_ION00007 | 1 | -0.09379890474 | Pirellulales | CACORA01 | CACORA01 sp9 | 0 | 0 | 0.9840192563 | 1.19E-04 | 0.9834864965 | 1 | 0 | 0.05793471235 | 0 | 5.42E-04 | 0 |
| TARA_SAMN05326647_METAG_ION00015 | 1 | -0.07158301039 | Actinomycetales | Microbacterium | Microbacterium g | 0 | 0 | 1 | 0 | 1 | 1 | 0.01616685011 | 1 | 0 | 0.007296326325 | 0.0217398377 |
| TARA_SAMN05326647_METAG_ION00022 | 3 | -0.03857408555 | TMED127 | | | 0 | 0 | 1.000000201 | 1.40E-04 | 0 | 1.000000201 | 0 | 0.0666092929 | 0 | 1.000000201 | 0.04504397995 |
| TARA_SAMN05326648_METAG_IOS00003 | 5 | -0.08838191696 | Caulobacterales | Marinicaulis | Marinicaulis sp0 | 0 | 0 | 0.0108134851 | 3.87E-05 | 0.004314350824 | 1 | 0 | 0.05612523577 | 0 | 8.38E-04 | 0 |
| TARA_SAMN05326648_METAG_IOS00004 | 4 | -0.134871119 | UBA8366 | GCA-002724395 | GCA-002724395 | 0 | 0 | 0.9984885093 | 9.53E-05 | 1.000000291 | 0 | 0 | 0.07092880132 | 0 | 0.001457166537 | 0.06229154544 |
| TARA_SAMN05326650_METAG_MED00002 | 7 | -0.01521902211 | UBA6522 | TMED78 | TMED78 sp0021 | 7.92E-04 | 0 | 0.03208534508 | 1.31E-04 | 0.003039990033 | 8.40E-04 | 0 | 0.07212179957 | 0 | 1 | 0.002260749571 |
| TARA_SAMN05326650_METAG_MED00005 | 2 | -0.00327617657 | SCGC-AAA003-L | TMED198 | TMED198 sp002 | 0 | 0 | 0.06230236185 | 1.79E-05 | 0.004116148795 | 0.001843381292 | 0 | 0.0684241824 | 0 | 7.71E-06 | 0 |
| TARA_SAMN05326650_METAG_MED00020 | 5 | -0.01083480315 | TMED109_A | MED722 | MED722 sp0026 | 0 | 0 | 0.02170571663 | 0.01682195237 | 0.02796746212 | 0.9999999964 | 0 | 0.09459997443 | 0 | 0.02708988247 | 0.01666666306 |
| TARA_SAMN05326650_METAG_MED00027 | 1 | -0.07075111873 | SCGC-AAA003-L08 | | | 0 | 0 | 1 | 1.82E-04 | 0.0718412061 | 1 | 0 | 0.08780416407 | 0 | 0.01830602188 | 0 |
| TARA_SAMN05326650_METAG_MED00034 | 7 | -0.1411130718 | Puniceispirillales | UBA12202 | UBA12202 sp002 | 0 | 0 | 0.9999998144 | 1.65E-04 | 0.04437102205 | 0.01587200551 | 0 | 0.9999998144 | 0 | 0.9999998144 | 0 |
| TARA_SAMN05326650_METAG_MED00040 | 1 | -0.03168497199 | Marinisomatales | NZWG01 | | 0 | 0 | 1 | 1.98E-05 | 0.5680057077 | 1 | 0 | 0.1123684142 | 0.03418970755 | 0 | 0 |
| TARA_SAMN05326650_METAG_MED00044 | 1 | -0.06547089885 | Opitutales | MB11C04 | MB11C04 sp002 | 0 | 0.001705148173 | 0.9618227621 | 1.21E-04 | 0.3389350307 | 0.9834016977 | 0 | 0.08790380136 | 0 | 0.02168830657 | 0 |
| TARA_SAMN05326650_METAG_MED00045 | 1 | -0.104909244 | Pseudomonadale | Luminiphilus | Luminiphilus sp0 | 0 | 0 | 0.9999997654 | 1.07E-04 | 0.1981118906 | 0.9999997654 | 0 | 0.08071492805 | 0 | 0.009650993734 | 0.06084564909 |
| TARA_SAMN05326650_METAG_MED00050 | 5 | 0.00460403836 | TMED109_A | GCA-2720125 | | 0 | 0 | 0.02705321585 | 0 | 3.70E-06 | 0.9999996913 | 0.0326899456 | 0.9999996913 | 0 | 0.00869587497 | 0 |
| TARA_SAMN05326650_METAG_MED00066 | 2 | -0.03028527968 | TMED127 | | | 0 | 0 | 0.01666760847 | 1.01E-04 | 0.2030279081 | 0 | 1.90E-06 | 0.9999995185 | 0 | 0.02551341146 | 0.07854531265 |
| TARA_SAMN05326650_METAG_MED00083 | 7 | -0.0694014399 | Parvibaculales | UBA7378 | UBA7378 sp913 | 0.05486326639 | 0.001821473657 | 0.04190519735 | 0.4916683349 | 0.03171442758 | 0.002049758666 | 0.04742288024 | 0.9842680817 | 1.79E-04 | 1.000000206 | 0 |
| TARA_SAMN05326651_METAG_RED00002 | 2 | -0.03841742327 | Marinisomatales | GCA-2722105 | GCA-2722105 sp | 0 | 0 | 0.04650415954 | 2.10E-04 | 0.002937104813 | 6.11E-06 | 0 | 0.08703769963 | 0 | 6.97E-04 | 0 |
| TARA_SAMN05326651_METAG_RED00008 | 3 | -0.07523877384 | Vicinamibacterale | DTWD01 | | 0 | 0 | 0.02117835611 | 7.20E-05 | 0.9999996634 | 0.9999996634 | 0 | 0.9999996634 | 0 | 0.9999996634 | 0 |
| TARA_SAMN05326651_METAG_RED00019 | 4 | -0.07042053249 | Anaerolineales | UBA11657 | UBA11657 sp002 | 0 | 0 | 0.9839134451 | 6.60E-05 | 0.1470860872 | 0.003469123259 | 0 | 0.05079912063 | 0 | 0.01363580317 | 0.002516130847 |
| TARA_SAMN05326651_METAG_RED00025 | 1 | -0.0490888665 | Pseudomonadale | UBA7455 | | 0 | 0 | 1 | 1.07E-04 | 0 | 1 | 0 | 1 | 0 | 0 | 0.03810255401 |
| TARA_SAMN05326651_METAG_RED00039 | 1 | -0.07514046756 | UBA1151 | UBA1328 | UBA1328 sp009 | 3.07E-06 | 0 | 1.000000176 | 1.14E-04 | 0.1741057312 | 1.000000176 | 0 | 0.9692527831 | 0 | 0.003499748223 | 0 |
