## Supplemental Table S2 for "Emergent Metabolic Niches for Marine Heterotrophs"

| ***Supplemental Table S2*** | | |
|---|---|---|
| Metabolite | Classification | CarveMe Reaction |
| 10 Phenyldecanoic acid | Carboxylic Acid | EX_phedca_e |
| 2 4 6 Trinitrotoluene | Other | EX_tnt_e |
| 2-Oxoglutarate | Ketones/Aldehydes | EX_akg_e |
| 4-Aminobenzoate | Amino Acids/Derivatives | EX_4abz_e |
| 4-Aminobutanoate | Amino Acids/Derivatives | EX_4abut_e |
| Acetaldehyde | Ketones/Aldehydes | EX_acald_e |
| Adenosine | Nucleobases/Nucleosides/Nucleotides/Derivatives | EX_adn_e |
| Ala-Gln | Peptides | EX_ala_gln_e |
| Ala-His | Peptides | EX_ala_his_e |
| Ala-L-Thr-L | Peptides | EX_ala_L_thr__L_e |
| Ala-Leu | Peptides | EX_ala_leu_e |
| Arsenate | Inorganic | EX_aso4_e |
| D Valine | Amino Acids/Derivatives | EX_val__D_e |
| Dimethyl sulfoxide | Organic Sulfur | EX_dmso_e |
| Ethanol | Alcohol | EX_etoh_e |
| Gly asn L C6H11N3O4 | Peptides | EX_gly_asn__L_e |
| Gly pro L C7H12N2O3 | Peptides | EX_gly_pro__L_e |
| Gly-Cys | Peptides | EX_gly_cys_e |
| Gly-Gln | Peptides | EX_gly_gln_e |
| Gly-Leu | Peptides | EX_gly_leu_e |
| Gly-Met | Peptides | EX_gly_met_e |
| Gly-Phe | Peptides | EX_gly_phe_e |
| Gly-Tyr | Peptides | EX_gly_tyr_e |
| Glycerophosphoglycerol | Other | EX_g3pg_e |
| Glycine-glycine-glutamine tripeptide | Peptides | EX_glyglygln_e |
| Glycylphenylalanine | Peptides | EX_glyphe_e |
| H2O H2O | Inorganic | EX_h2o_e |
| Hydrogen cyanide | Other | EX_cyan_e |
| Hydrogen peroxide | Inorganic | EX_h2o2_e |
| L Glycinylmethionine | Peptides | EX_glymet_e |

| | | |
|---|---|---|
| L alaninyltryptophan | Peptides | EX_alatrp_e |
| L glycinylglutamine | Peptides | EX_glygln_e |
| L histidinylhistidine | Peptides | EX_hishis_e |
| L methionine R oxide C5H11NO3S | Amino Acids/Derivatives | EX_metox__R_e |
| L-Alanine | Amino Acids/Derivatives | EX_ala__L_e |
| L-Arginine | Amino Acids/Derivatives | EX_arg__L_e |
| L-Asparagine | Amino Acids/Derivatives | EX_asn__L_e |
| L-Glutamate | Amino Acids/Derivatives | EX_glu__L_e |
| L-Glutamine | Amino Acids/Derivatives | EX_gln__L_e |
| L-Lysine | Amino Acids/Derivatives | EX_lys__L_e |
| L-Phenylalanine | Amino Acids/Derivatives | EX_phe__L_e |
| L-Proline | Amino Acids/Derivatives | EX_pro__L_e |
| L-Threonine | Amino Acids/Derivatives | EX_thr__L_e |
| L-Tryptophan | Amino Acids/Derivatives | EX_trp__L_e |
| L-Tyrosine | Amino Acids/Derivatives | EX_tyr__L_e |
| L-alanine-D-glutamate-meso-2,6-diaminoheptanedioate | Other | EX_LalaDgluMdap_e |
| Lysine-glutamine-glycine tripeptide | Peptides | EX_lysglugly_e |
| Met L ala L C8H16N2O3S | Peptides | EX_met_L_ala__L_e |
| Nitrate | Inorganic | EX_no3_e |
| Nitrite | Inorganic | EX_no2_e |
| O2 O2 | Inorganic | EX_o2_e |
| Phosphoenolpyruvate | Carboxylic Acid | EX_pep_e |
| Putrescine | Amines/Amides | EX_ptrc_e |
| Sulfur | Inorganic | EX_s_e |
| Tetrathionate | Organic Sulfur | EX_tet_e |
| Thiamin | B Vitamins | EX_thm_e |
| Trithionate | Organic Sulfur | EX_tton_e |
| Uracil | Nucleobases/Nucleosides/Nucleotides/Derivatives | EX_ura_e |
| Xanthine | Nucleobases/Nucleosides/Nucleotides/Derivatives | EX_xan_e |
| 3 Hydroxy 10 phenyldecanoic acid | Carboxylic Acid | EX_R_3hpdeca_e |
| 2-Oxobutanoate | Carboxylic Acid | EX_2obut_e |
| Alpha-L-Arabinan (3 subunits) | Nucleobases/Nucleosides/Nucleotides/Derivatives | EX_araban__L_e |

| | | |
|---|---|---|
| Chorismate | Carboxylic Acid | EX_chor_e |
| Cytidine | Nucleobases/Nucleosides/Nucleotides/Derivatives | EX_cytd_e |
| D-Serine | Amino Acids/Derivatives | EX_ser__D_e |
| Deoxycytidine | Nucleobases/Nucleosides/Nucleotides/Derivatives | EX_dcyt_e |
| Diacetylchitobiose | Carbohydrates/Derivatives | EX_dachi_e |
| Fe(III)hydroxamate | Other | EX_fe3hox_e |
| Folate | B Vitamins | EX_fol_e |
| Formamide | Amines/Amides | EX_frmd_e |
| Glycine betaine | Amino Acids/Derivatives | EX_glyb_e |
| Guanine | Nucleobases/Nucleosides/Nucleotides/Derivatives | EX_gua_e |
| Iron(III) chelated carboxymycobactin T (R=8 carbon, final carbon is c | Other | EX_fcmcbtt_e |
| L Methionine S oxide C5H11NO3S | Amino Acids/Derivatives | EX_metox_e |
| L alaninylthreonine | Peptides | EX_alathr_e |
| L-Citrulline | Amino Acids/Derivatives | EX_citr__L_e |
| L-Leucine | Amino Acids/Derivatives | EX_leu__L_e |
| Maltotriose C18H32O16 | Carbohydrates/Derivatives | EX_malttr_e |
| N,N'-diacetylchitobiose | Carbohydrates/Derivatives | EX_chtbs_e |
| NMN C11H14N2O8P | Nucleobases/Nucleosides/Nucleotides/Derivatives | EX_nmn_e |
| Pyridoxine | B Vitamins | EX_pydxn_e |
| S-Methyl-L-methionine | Amino Acids/Derivatives | EX_mmet_e |
| Thymidine C10H14N2O5 | Nucleobases/Nucleosides/Nucleotides/Derivatives | EX_thymd_e |
| UDP-N-acetyl-3-O-(1-carboxyvinyl)-D-glucosamine | Nucleobases/Nucleosides/Nucleotides/Derivatives | EX_uaccg_e |
| Undecaprenyl phosphate | Phospholipids/Fatty Acids/Triglycerides | EX_udcpp_e |
| Urea CH4N2O | Amines/Amides | EX_urea_e |
| 1,4-alpha-D-glucan | Carbohydrates/Derivatives | EX_14glucan_e |
| 1-deoxy-D-xylulose | Carbohydrates/Derivatives | EX_dxyl_e |
| Ammonium | Inorganic | EX_nh4_e |
| Carbon monoxide | Inorganic | EX_co_e |
| D-Fructose | Carbohydrates/Derivatives | EX_fru_e |
| L histidinylglycine | Amino Acids/Derivatives | EX_hisgly_e |
| N-Acetyl-D-glucosamine 1-phosphate | Carbohydrates/Derivatives | EX_acgam1p_e |
| Shikimate | Carboxylic Acid | EX_skm_e |

| | | |
|---|---|---|
| Sulfate | Inorganic | EX_so4_e |
| Thiosulfate | Organic Sulfur | EX_tsul_e |
| 2',3'-Cyclic CMP | Nucleobases/Nucleosides/Nucleotides/Derivatives | EX_23ccmp_e |
| 4-Aminobutanal | Ketones/Aldehydes | EX_4abutn_e |
| 5-Methylcytosine | Nucleobases/Nucleosides/Nucleotides/Derivatives | EX_5mcsn_e |
| Adenine | Nucleobases/Nucleosides/Nucleotides/Derivatives | EX_ade_e |
| Alpha,alpha'-Trehalose 6-phosphate | Carbohydrates/Derivatives | EX_tre6p_e |
| Choline C5H14NO | Amines/Amides | EX_chol_e |
| Creatinine | Amines/Amides | EX_crtn_e |
| D Asparagine | Amino Acids/Derivatives | EX_asn__D_e |
| D-Glucose 1-phosphate | Carbohydrates/Derivatives | EX_g1p_e |
| D-Glucuronate | Carbohydrates/Derivatives | EX_glcur_e |
| D-Tagatose | Carbohydrates/Derivatives | EX_tag__D_e |
| Deoxyuridine | Nucleobases/Nucleosides/Nucleotides/Derivatives | EX_duri_e |
| Glycine | Amino Acids/Derivatives | EX_gly_e |
| H+ | Inorganic | EX_h_e |
| Hydrogen sulfide | Inorganic | EX_h2s_e |
| L alaninylhistidine | Peptides | EX_alahis_e |
| L glycinylserine | Peptides | EX_glyser_e |
| L-Homoserine | Amino Acids/Derivatives | EX_hom__L_e |
| L-Methionine Sulfoxide | Amino Acids/Derivatives | EX_metsox_S__L_e |
| L-Prolinylglycine | Peptides | EX_progly_e |
| L-alanine-L-glutamate | Peptides | EX_LalaLglu_e |
| Leucylleucine | Peptides | EX_leuleu_e |
| Maltoheptaose | Carbohydrates/Derivatives | EX_malthp_e |
| Phenylethyl alcohol | Alcohol | EX_pea_e |
| Proline-histidine-glutamine tripeptide | Peptides | EX_prohisglu_e |
| Raffinose C18H32O16 | Carbohydrates/Derivatives | EX_raffin_e |
| Reduced glutathione | Organic Sulfur | EX_gthrd_e |
| Uridine | Nucleobases/Nucleosides/Nucleotides/Derivatives | EX_uri_e |
| 3 Hydroxypentanoic acid | Phospholipids/Fatty Acids/Triglycerides | EX_R_3hpt_e |
| R R 2 3 Butanediol C4H10O2 | Alcohol | EX_btd_RR_e |

| | | |
|---|---|---|
| 2',3'-Cyclic GMP | Nucleobases/Nucleosides/Nucleotides/Derivatives | EX_23cgmp_e |
| 2,3-diaminopropionate | Amino Acids/Derivatives | EX_23dappa_e |
| 2,5-diketo-D-gluconate | Carboxylic Acid | EX_25dkglcn_e |
| 2-Dehydro-D-gluconate | Carboxylic Acid | EX_2dhglcn_e |
| 3 CMP C9H12N3O8P | Nucleobases/Nucleosides/Nucleotides/Derivatives | EX_3cmp_e |
| 3-Phospho-D-glycerate | Carboxylic Acid | EX_3pg_e |
| 4 Hydroxyphenylacetic acid C8H8O3 | Carboxylic Acid | EX_4hoxpac_e |
| Arbutin C12H16O7 | Carbohydrates/Derivatives | EX_arbt_e |
| Benzoate | Carboxylic Acid | EX_bz_e |
| Catechol | Alcohol | EX_catechol_e |
| Cellobiose | Carbohydrates/Derivatives | EX_cellb_e |
| Cis-Aconitate | Carboxylic Acid | EX_acon_C_e |
| Coenzyme A | Other | EX_coa_e |
| Creatine cytosol | Amino Acids/Derivatives | EX_creat_e |
| D-Galacturonate | Carbohydrates/Derivatives | EX_galur_e |
| D-Glucosamine | Carbohydrates/Derivatives | EX_gam_e |
| D-Glucose | Carbohydrates/Derivatives | EX_glc__D_e |
| D-Glyceraldehyde | Carbohydrates/Derivatives | EX_glyald_e |
| D-Methionine | Amino Acids/Derivatives | EX_met__D_e |
| D-O-Phosphoserine | Amino Acids/Derivatives | EX_pser__D_e |
| D-Proline | Amino Acids/Derivatives | EX_pro__D_e |
| D-Xylose | Carbohydrates/Derivatives | EX_xyl__D_e |
| D-phenylalanine | Amino Acids/Derivatives | EX_phe__D_e |
| Dextrin C12H20O10 | Carbohydrates/Derivatives | EX_dextrin_e |
| Dihydroxyacetone | Carbohydrates/Derivatives | EX_dha_e |
| Dihydroxyacetone phosphate | Carbohydrates/Derivatives | EX_dhap_e |
| Dopamine | Amines/Amides | EX_dopa_e |
| Fe2+ mitochondria | Inorganic | EX_fe2_e |
| Fumarate | Carboxylic Acid | EX_fum_e |
| Galactitol | Carbohydrates/Derivatives | EX_galt_e |
| Glycerol 3-phosphate | Carbohydrates/Derivatives | EX_glyc3p_e |
| Isocitrate | Carboxylic Acid | EX_icit_e |

| | | |
|---|---|---|
| L-Arabinose | Carbohydrates/Derivatives | EX_arab__L_e |
| L-tartrate | Carboxylic Acid | EX_tartr__L_e |
| Linoleic acid (all cis C18:2) n-6 | Phospholipids/Fatty Acids/Triglycerides | EX_lnlc_e |
| Meso-2,6-Diaminoheptanedioate | Other | EX_26dap__M_e |
| N-Acetyl-D-glucosamine(anhydrous)N-Acetylmuramic acid | Other | EX_anhgm_e |
| N-Formimidoyl-L-glutamate | Amino Acids/Derivatives | EX_forglu_e |
| Nitric oxide | Inorganic | EX_no_e |
| Octacosanoyl-CoA | Other | EX_octscoa_e |
| Selenate | Inorganic | EX_sel_e |
| Succinate | Carboxylic Acid | EX_succ_e |
| Thymine C5H6N2O2 | Nucleobases/Nucleosides/Nucleotides/Derivatives | EX_thym_e |
| Trans 4 Hydroxy L proline C5H9NO3 | Amino Acids/Derivatives | EX_4hpro_LT_e |
| Trimethylamine N-oxide | Amines/Amides | EX_tmao_e |
| Undecaprenyl diphosphate | Phospholipids/Fatty Acids/Triglycerides | EX_udcpdp_e |
| 3 hydroxynonanoic acid | Carboxylic Acid | EX_R_3hnonaa_e |
| 2',3'-Cyclic AMP | Nucleobases/Nucleosides/Nucleotides/Derivatives | EX_23camp_e |
| 3 AMP C10H12N5O7P | Nucleobases/Nucleosides/Nucleotides/Derivatives | EX_3amp_e |
| 3-Methylbutanoic acid | Carboxylic Acid | EX_3mb_e |
| 4-Hydroxyphenylacetate | Carboxylic Acid | EX_4hphac_e |
| 8 Phenyloctanoic acid | Carboxylic Acid | EX_pheocta_e |
| Beta D glucose 6 phosphate C6H11O9P | Carbohydrates/Derivatives | EX_g6p_B_e |
| Cu+ | Inorganic | EX_cu_e |
| D-Arginine | Amino Acids/Derivatives | EX_arg__D_e |
| D-Galactarate | Carboxylic Acid | EX_galct__D_e |
| D-Lysine | Amino Acids/Derivatives | EX_lys__D_e |
| D-Mannitol | Carbohydrates/Derivatives | EX_mnl_e |
| D-tartrate | Carboxylic Acid | EX_tartr__D_e |
| DCMP C9H12N3O7P | Nucleobases/Nucleosides/Nucleotides/Derivatives | EX_dcmp_e |
| DTMP C10H13N2O8P | Nucleobases/Nucleosides/Nucleotides/Derivatives | EX_dtmp_e |
| Deoxyadenosine | Nucleobases/Nucleosides/Nucleotides/Derivatives | EX_dad_2_e |
| Fructoselysine | Other | EX_frulys_e |
| Glycerol | Alcohol | EX_glyc_e |

| | | |
|---|---|---|
| Glycerophosphoserine | Phospholipids/Fatty Acids/Triglycerides | EX_g3ps_e |
| Glycolate C2H3O3 | Carboxylic Acid | EX_glyclt_e |
| Heptanoate | Phospholipids/Fatty Acids/Triglycerides | EX_hpta_e |
| Hydrogen | Inorganic | EX_h2_e |
| Hydroxylamine | Amines/Amides | EX_ham_e |
| Indole | Other | EX_indole_e |
| L-Histidine | Amino Acids/Derivatives | EX_his__L_e |
| L-Isoleucine | Amino Acids/Derivatives | EX_ile__L_e |
| L-Serine | Amino Acids/Derivatives | EX_ser__L_e |
| L-Valine | Amino Acids/Derivatives | EX_val__L_e |
| L-alanine-D-glutamate-meso-2,6-diaminoheptanedioate-D-alanine | Other | EX_LalaDgluMdapDala_e |
| L-methionine-R-sulfoxide | Amino Acids/Derivatives | EX_metsox_R__L_e |
| Methanol | Alcohol | EX_meoh_e |
| N-Acetylneuraminate | Carbohydrates/Derivatives | EX_acnam_e |
| Nonanoate C9H17O2 | Phospholipids/Fatty Acids/Triglycerides | EX_nona_e |
| Octadecanoate (n-C18:0) | Phospholipids/Fatty Acids/Triglycerides | EX_ocdca_e |
| Octadecenoate (n-C18:1) | Phospholipids/Fatty Acids/Triglycerides | EX_ocdcea_e |
| Ornithine | Amino Acids/Derivatives | EX_orn_e |
| Petroselaidic acid | Phospholipids/Fatty Acids/Triglycerides | EX_ptsla_e |
| Pyruvate | Carboxylic Acid | EX_pyr_e |
| Sulfite | Inorganic | EX_so3_e |
| Superoxide anion | Inorganic | EX_o2s_e |
| Triacylglycerol hexadecanoate | Phospholipids/Fatty Acids/Triglycerides | EX_tag160_e |
| Triacylglycerol nC181d9 | Phospholipids/Fatty Acids/Triglycerides | EX_tag181d9_e |
| UDP-D-glucuronate | Carbohydrates/Derivatives | EX_udpglcur_e |
| L-fuculose | Carbohydrates/Derivatives | EX_fuc_e |
| Propanal | Ketones/Aldehydes | EX_ppal_e |
| L-Malate | Carboxylic Acid | EX_mal__L_e |
| 3 Hydroxy 10 phenyldecanoic acid | Carboxylic Acid | EX_R_3hpdeca_e |
| UMP C9H11N2O9P | Nucleobases/Nucleosides/Nucleotides/Derivatives | EX_ump_e |
| 3 Hydroxy 4Z decenic acid | Carboxylic Acid | EX_R3hdec4e_e |
| 3 Hydroxypentanoic acid | Carboxylic Acid | EX_R_3hpt_e |

| | | |
|---|---|---|
| R R 2 3 Butanediol C4H10O2 | Alcohol | EX_btd_RR_e |
| 4-Hydroxybenzoate | Carboxylic Acid | EX_4hbz_e |
| Cytosine | Nucleobases/Nucleosides/Nucleotides/Derivatives | EX_csn_e |
| D-Mannose | Carbohydrates/Derivatives | EX_man_e |
| D-Mannose 1-phosphate | Carbohydrates/Derivatives | EX_man1p_e |
| Diacetyl C4H6O2 | Ketones/Aldehydes | EX_diact_e |
| Lactose C12H22O11 | Carbohydrates/Derivatives | EX_lcts_e |
| Orotate C5H3N2O4 | Nucleobases/Nucleosides/Nucleotides/Derivatives | EX_orot_e |
| 3 hydroxynonanoic acid | Carboxylic Acid | EX_R_3hnonaa_e |
| Cys Gly C5H10N2O3S | Peptides | EX_cgly_e |
| D-Ornithine | Amino Acids/Derivatives | EX_orn__D_e |
| (R)-Propane-1,2-diol | Alcohol | EX_12ppd__R_e |
| 1-Propanol | Alcohol | EX_ppoh_e |
| 2-3-dihydroxybenzoylserine trimer-Fe-III | Other | EX_fe3dhbzs3_e |
| 2-Aminoethylphosphonate | Other | EX_2ameph_e |
| 3-Oxoadipate | Carboxylic Acid | EX_3oxoadp_e |
| 4-aminobenzoate-glutamate | Peptides | EX_abg4_e |
| D-Glucose 6-phosphate | Carbohydrates/Derivatives | EX_g6p_e |
| DAMP C10H12N5O6P | Nucleobases/Nucleosides/Nucleotides/Derivatives | EX_damp_e |
| DGMP C10H12N5O7P | Nucleobases/Nucleosides/Nucleotides/Derivatives | EX_dgmp_e |
| DUMP C9H11N2O8P | Nucleobases/Nucleosides/Nucleotides/Derivatives | EX_dump_e |
| Decanoate (n-C10:0) | Phospholipids/Fatty Acids/Triglycerides | EX_dca_e |
| Glycolaldehyde | Ketones/Aldehydes | EX_gcald_e |
| Hexadecanoate (n-C16:0) | Phospholipids/Fatty Acids/Triglycerides | EX_hdca_e |
| Hexadecenoate (n-C16:1) | Phospholipids/Fatty Acids/Triglycerides | EX_hdcea_e |
| Inosine | Nucleobases/Nucleosides/Nucleotides/Derivatives | EX_ins_e |
| Pentanoate | Phospholipids/Fatty Acids/Triglycerides | EX_pta_e |
| Phenylacetaldehyde | Ketones/Aldehydes | EX_pacald_e |
| Phosphate | Inorganic | EX_pi_e |
| Phosphotyrosine | Amino Acids/Derivatives | EX_tyrp_e |
| Starch C12H20O10 | Carbohydrates/Derivatives | EX_starch_e |
| Sucrose C12H22O11 | Carbohydrates/Derivatives | EX_sucr_e |

| | | |
|---|---|---|
| UDP-N-acetyl-D-galactosamine | Nucleobases/Nucleosides/Nucleotides/Derivatives | EX_udpacgal_e |
| Alpha L Arabinan C15H24O12 | Carbohydrates/Derivatives | EX_Larab_e |
| Calcium | Inorganic | EX_ca2_e |
| Chloride | Inorganic | EX_cl_e |
| Co2+ | Inorganic | EX_cobalt2_e |
| Copper | Inorganic | EX_cu2_e |
| Iron bound extracellular staphyloferrin B | Other | EX_istfrnB_e |
| Potassium | Inorganic | EX_k_e |
| Magnesium | Inorganic | EX_mg2_e |
| Manganese | Inorganic | EX_mn2_e |
| Pyridoxamine | B Vitamins | EX_pydam_e |
| Zinc | Inorganic | EX_zn2_e |
| Iron bound extracellular staphyloferrin A | Other | EX_istfrnA_e |
| L-Methionine | Amino Acids/Derivatives | EX_met__L_e |
| Sn-Glycero-3-phosphocholine | Phospholipids/Fatty Acids/Triglycerides | EX_g3pc_e |
| 2 methylbutyraldehyde C5H10O | Ketones/Aldehydes | EX_2mbald_e |
| Citrate | Carboxylic Acid | EX_cit_e |
| Deoxyguanosine | Nucleobases/Nucleosides/Nucleotides/Derivatives | EX_dgsn_e |
| (R)-Pantothenate | B Vitamins | EX_pnto__R_e |
| Riboflavin C17H20N4O6 | B Vitamins | EX_ribflv_e |
| 4-Amino-5-hydroxymethyl-2-methylpyrimidine | Other | EX_4ahmmp_e |
| D-Alanyl-D-alanine | Peptides | EX_alaala_e |
| Iron (Fe3+) | Inorganic | EX_fe3_e |
| Hypoxanthine | Nucleobases/Nucleosides/Nucleotides/Derivatives | EX_hxan_e |
| Diphosphate | Inorganic | EX_ppi_e |
| Pseudouridine | Nucleobases/Nucleosides/Nucleotides/Derivatives | EX_psuri_e |
| Trehalose | Carbohydrates/Derivatives | EX_tre_e |
| Ferrypyoverdine P putida KT2440 specific | Other | EX_fe3pyovd_kt_e |
| 2-Dehydro-3-deoxy-D-gluconate | Carboxylic Acid | EX_2ddglcn_e |
| D Arabinose C5H10O5 | Ketones/Aldehydes | EX_arab__D_e |
| Methanethiol CH4S | Organic Sulfur | EX_ch4s_e |
| L-Cystathionine | Amino Acids/Derivatives | EX_cyst__L_e |

| | | |
|---|---|---|
| Myo-Inositol | Carbohydrates/Derivatives | EX_inost_e |
| Serine-glutamine-glycine tripeptide | Peptides | EX_serglugly_e |
| Fe(III)dicitrate | Other | EX_fe3dcit_e |
| D-Glucosamine 6-phosphate | Carbohydrates/Derivatives | EX_gam6p_e |
| Guanosine | Nucleobases/Nucleosides/Nucleotides/Derivatives | EX_gsn_e |
| Nicotinate | B Vitamins | EX_nac_e |
| D Tyrosine | Amino Acids/Derivatives | EX_tyr__D_e |
| D-Galactonate | Carboxylic Acid | EX_galctn__D_e |
| 3 UMP C9H11N2O9P | Nucleobases/Nucleosides/Nucleotides/Derivatives | EX_3ump_e |
| L-Cysteine | Amino Acids/Derivatives | EX_cys__L_e |
| Maltose C12H22O11 | Carbohydrates/Derivatives | EX_malt_e |
| Maltohexaose | Carbohydrates/Derivatives | EX_malthx_e |
| GMP C10H12N5O8P | Nucleobases/Nucleosides/Nucleotides/Derivatives | EX_gmp_e |
| D-Glycerate 2-phosphate | Carbohydrates/Derivatives | EX_2pg_e |
| D-Mannose 6-phosphate | Carbohydrates/Derivatives | EX_man6p_e |
| N-Acetyl-D-mannosamine | Carbohydrates/Derivatives | EX_acmana_e |
| 2-Phosphoglycolate | Carboxylic Acid | EX_2pglyc_e |
| Aminoimidazole-riboside | Nucleobases/Nucleosides/Nucleotides/Derivatives | EX_airs_e |
| Phenethylamine | Amines/Amides | EX_peamn_e |
| Pyridoxal | B Vitamins | EX_pydx_e |
| (S)-Propane-1,2-diol | Alcohol | EX_12ppd__S_e |
| Guanosine 3 phosphate C10H12N5O8P | Nucleobases/Nucleosides/Nucleotides/Derivatives | EX_3gmp_e |
| AMP C10H12N5O7P | Nucleobases/Nucleosides/Nucleotides/Derivatives | EX_amp_e |
| Formate | Carboxylic Acid | EX_for_e |
| Deoxyribose C5H10O4 | Carbohydrates/Derivatives | EX_drib_e |
| Gly asp L C6H9N2O5 | Peptides | EX_gly_asp__L_e |
| Fe-enterobactin | Other | EX_feenter_e |
| Sn-Glycero-3-phosphoethanolamine | Phospholipids/Fatty Acids/Triglycerides | EX_g3pe_e |
| Sn-Glycero-3-phospho-1-inositol | Phospholipids/Fatty Acids/Triglycerides | EX_g3pi_e |
| D-Fructose 6-phosphate | Carbohydrates/Derivatives | EX_f6p_e |
| L-Fucose | Carbohydrates/Derivatives | EX_fuc__L_e |
| 4-Hydroxy-benzyl alcohol | Alcohol | EX_4hba_e |

| | | |
|---|---|---|
| L Ornithine C5H13N2O2 | Amino Acids/Derivatives | EX_orn__L_e |
| D-Sorbitol | Carbohydrates/Derivatives | EX_sbt__D_e |
| N-Acetyl-D-glucosamine | Carbohydrates/Derivatives | EX_acgam_e |
| L-Aspartate | Amino Acids/Derivatives | EX_asp__L_e |
| Ethanolamine | Amines/Amides | EX_etha_e |
| Allantoin | Other | EX_alltn_e |
| Propionate (n-C3:0) | Carboxylic Acid | EX_ppa_e |
| 4-aminobenzoyl-glutamate | Amino Acids/Derivatives | EX_4abzglu_e |
| Tagatose | Carbohydrates/Derivatives | EX_tgt_e |
| Methanesulfonate | Organic Sulfur | EX_mso3_e |
| D-Glucuronate 1-phosphate | Carbohydrates/Derivatives | EX_glcur1p_e |
| D-Ribose | Carbohydrates/Derivatives | EX_rib__D_e |
| Maltotetraose | Carbohydrates/Derivatives | EX_maltttr_e |
| Agmatine | Amines/Amides | EX_agm_e |
| L-Carnosine | Peptides | EX_carn_e |
| Toluene | Other | EX_tol_e |
| Xanthosine | Nucleobases/Nucleosides/Nucleotides/Derivatives | EX_xtsn_e |
| Salicin C13H18O7 | Carbohydrates/Derivatives | EX_salcn_e |
| CMP C9H12N3O8P | Nucleobases/Nucleosides/Nucleotides/Derivatives | EX_cmp_e |
| L-Threonine O-3-phosphate | Amino Acids/Derivatives | EX_thrp_e |
| Nitrous oxide | Inorganic | EX_n2o_e |
| Benzaldehyde | Ketones/Aldehydes | EX_bzal_e |
| Deoxyinosine | Nucleobases/Nucleosides/Nucleotides/Derivatives | EX_din_e |
| N Ribosylnicotinamide C11H15N2O5 | Nucleobases/Nucleosides/Nucleotides/Derivatives | EX_rnam_e |
| Nicotinamide | B Vitamins | EX_ncam_e |
| 2',3'-Cyclic UMP | Nucleobases/Nucleosides/Nucleotides/Derivatives | EX_23cump_e |
| D-Alanine | Amino Acids/Derivatives | EX_ala__D_e |
| DIMP C10H12N4O7P | Nucleobases/Nucleosides/Nucleotides/Derivatives | EX_dimp_e |
| L alaninylleucine | Peptides | EX_alaleu_e |
| L-Ascorbate | Other | EX_ascb__L_e |
| UDP-N-acetyl-D-glucosamine | Nucleobases/Nucleosides/Nucleotides/Derivatives | EX_uacgam_e |
| Oxidized glutathione | Organic Sulfur | EX_gthox_e |

| | | |
|---|---|---|
| UDPgalactose | Nucleobases/Nucleosides/Nucleotides/Derivatives | EX_udpgal_e |
| L-Lyxose | Carbohydrates/Derivatives | EX_lyx__L_e |
| Propanoyl phosphate | Other | EX_ppap_e |
| Formaldehyde | Ketones/Aldehydes | EX_fald_e |
| Indole 3 acetaldehyde C10H9NO | Ketones/Aldehydes | EX_id3acald_e |
| 6-Phospho-D-gluconate | Carboxylic Acid | EX_6pgc_e |
| L-Xylulose | Carbohydrates/Derivatives | EX_xylu__L_e |
| D-Mannitol 1-phosphate | Carbohydrates/Derivatives | EX_mnl1p_e |
| D-Glutamate | Amino Acids/Derivatives | EX_glu__D_e |
| IMP C10H11N4O8P | Nucleobases/Nucleosides/Nucleotides/Derivatives | EX_imp_e |
| UDPglucose | Nucleobases/Nucleosides/Nucleotides/Derivatives | EX_udpg_e |
| Phenylacetic acid | Carboxylic Acid | EX_pac_e |
| 3 Hydroxy 8 phenyloctanoic acid | Phospholipids/Fatty Acids/Triglycerides | NA |
| Acetate | Carboxylic Acid | EX_ac_e |
| 5-Methylthio-D-ribose | Organic Sulfur | EX_5mtr_e |
| Acetoacetate | Carboxylic Acid | EX_acac_e |
| Tetradecanoate (n-C14:0) | Phospholipids/Fatty Acids/Triglycerides | EX_ttdca_e |
| Alpha D-glucose | Carbohydrates/Derivatives | EX_glc__aD_e |
| Quinate | Carboxylic Acid | EX_quin_e |
| L Ectoine | Carboxylic Acid | EX_ecto__L_e |
| Ala L asp L C7H11N2O5 | Peptides | EX_ala_L_asp__L_e |
| 6 Phenylhexanoic acid | Carboxylic Acid | EX_phehxa_e |
| D-Glucarate | Carboxylic Acid | EX_glcr_e |
| Choline sulfate | Organic Sulfur | EX_chols_e |
| L-Lactate | Carboxylic Acid | EX_lac__L_e |
| 9 Phenylnonanoic acid | Carboxylic Acid | EX_phenona_e |
| D Leucine | Amino Acids/Derivatives | EX_leu__D_e |
| Oxaloacetate | Carboxylic Acid | EX_oaa_e |
| Psicoselysine | Amino Acids/Derivatives | EX_psclys_e |
| D-Gluconate | Carboxylic Acid | EX_glcn_e |
| 4-Hydroxy-L-threonine | Amino Acids/Derivatives | EX_4hthr_e |
| Beta-Alanine | Amino Acids/Derivatives | EX_ala_B_e |

| | | |
|---|---|---|
| L Sorbose C6H12O6 | Carbohydrates/Derivatives | EX_srb__L_e |
| Beta Alaninamide | Amino Acids/Derivatives | EX_balamd_e |
| Butyrate (n-C4:0) | Phospholipids/Fatty Acids/Triglycerides | EX_but_e |
| 3 hydroxy 5Z 8Z tetradecedienic acid | Phospholipids/Fatty Acids/Triglycerides | NA |
| Benzyl alcohol | Alcohol | EX_bzalc_e |
| 2(alpha-D-Mannosyl)-D-glycerate | Carbohydrates/Derivatives | EX_manglyc_e |
| Maltopentaose | Carbohydrates/Derivatives | EX_maltpt_e |
| Glycylglycine C4H8N2O3 | Peptides | EX_glygly_e |
| Beta D-Galactose | Carbohydrates/Derivatives | EX_gal_bD_e |
| 1 4 Diguanidinobutane | Amines/Amides | EX_dgudbutn_e |
| L Arginine phosphate C6H14N4O5P | Amino Acids/Derivatives | EX_argp_e |
| R Acetoin C4H8O2 | Ketones/Aldehydes | NA |
| N-Acetyl-L-glutamate | Amino Acids/Derivatives | EX_acglu_e |
| L-Rhamnose | Carbohydrates/Derivatives | EX_rmn_e |
| Beta-1,3/1,4-glucan (Barley, n=6, Glc beta1->3,4 Glc) | Carbohydrates/Derivatives | EX_glucan6_e |
| Xylotriose | Carbohydrates/Derivatives | EX_xyl3_e |
| Arbutin 6-phosphate | Carbohydrates/Derivatives | EX_arbt6p_e |
| Xanthosine 5'-phosphate | Nucleobases/Nucleosides/Nucleotides/Derivatives | EX_xmp_e |
| Salmochelin-S2-Fe-III | Other | EX_salchs2fe_e |
| O-Phospho-L-serine | Amino Acids/Derivatives | EX_pser__L_e |
| Ala L glu L C8H13N2O5 | Peptides | EX_ala_L_glu__L_e |
| N-Acetylmuramate | Carbohydrates/Derivatives | EX_acmum_e |
| L-alanine-D-glutamate | Peptides | EX_LalaDglu_e |
| Ferric 2,3-dihydroxybenzoylserine | Other | EX_fe3dhbzs_e |
| Vaccenic acid | Phospholipids/Fatty Acids/Triglycerides | EX_vacc_e |
| Urate C5H4N4O3 | Nucleobases/Nucleosides/Nucleotides/Derivatives | EX_urate_e |
| Glycerol 2-phosphate | Other | EX_glyc2p_e |
| Hexanoate (n-C6:0) | Phospholipids/Fatty Acids/Triglycerides | EX_hxa_e |
| 3 hydroxyheptanoic acid | Carboxylic Acid | NA |
| Tyramine | Amino Acids/Derivatives | EX_tym_e |
| Cellulose (n=4 repeating units) | Carbohydrates/Derivatives | EX_cell4_e |
| (R)-mevalonate | Carboxylic Acid | EX_mevR_e |

| | | |
|---|---|---|
| D-Galactose | Carbohydrates/Derivatives | EX_gal_e |
| L-Galactonate | Carboxylic Acid | EX_galctn__L_e |
| L glycinylglutamate | Peptides | EX_glyglu_e |
| 4-Hydroxybenzaldehyde | Ketones/Aldehydes | EX_4hbald_e |
| Triacylglycerol octadecanoate | Phospholipids/Fatty Acids/Triglycerides | EX_tag180_e |
| Triacylglycerol nC182d9d12 | Phospholipids/Fatty Acids/Triglycerides | EX_tag182d9d12_e |
| Dimethyl sulfone | Organic Sulfur | EX_dmso2_e |
| Beta alanylL leucine | Peptides | EX_balaleu_e |
| Myo-Inositol hexakisphosphate | Other | EX_minohp_e |
| 5-Dehydro-D-gluconate | Carbohydrates/Derivatives | EX_5dglcn_e |
| L-Idonate | Carboxylic Acid | EX_idon__L_e |
| L-Carnitine | Amines/Amides | EX_crn_e |
| L-Homocysteine | Amino Acids/Derivatives | EX_hcys__L_e |
| D-Lactate | Carboxylic Acid | EX_lac__D_e |
| Salmochelin-S4-Fe-III | Other | EX_salchs4fe_e |
| 3 Hydroxydodecanoic 6 en acid | Carboxylic Acid | NA |
| Isethionic acid | Organic Sulfur | EX_isetac_e |
| Salicylate | Carboxylic Acid | EX_salc_e |
| Ethanesulfonate | Organic Sulfur | EX_ethso3_e |
| Glycol | Alcohol | EX_glycol_e |
| Ribitol | Carbohydrates/Derivatives | EX_rbt_e |
| Coniferol | Other | EX_confrl_e |
| Isethionate C2H5O4S | Organic Sulfur | EX_istnt_e |
| 2-Oxoarginine | Carboxylic Acid | EX_5g2oxpt_e |
| Xylan (4 backbone units, 1 glcur side chain) | Carbohydrates/Derivatives | EX_xylan4_e |
| Phosphonate | Other | EX_ppat_e |
| 1-O-methyl-Beta-D-glucuronate | Carbohydrates/Derivatives | EX_metglcur_e |
| D Galactarate C6H8O8 | Carboxylic Acid | EX_galctr__D_e |
| D Galactosamine C6H13NO5 | Carbohydrates/Derivatives | EX_galam_e |
| Galactomannan(n=6 repeat units mannose, alpha-1,4 man) | Carbohydrates/Derivatives | EX_galman6_e |
| 3 Hydroxyphenylacetic acid C8H8O3 | Carboxylic Acid | EX_3hoxpac_e |
| Galactomannan(n=4 repeat units mannose, alpha-1,4 man) | Carbohydrates/Derivatives | EX_galman4_e |

| | | |
|---|---|---|
| D-Xylonate | Carboxylic Acid | EX_dxylnt_e |
| L-gulonate | Carboxylic Acid | EX_guln__L_e |
| Xylan (8 backbone units, 2 glcur side chain) | Carbohydrates/Derivatives | EX_xylan8_e |
| Beta Methylglucoside C7H14O6 | Other | EX_mbdg_e |
| D-Fructuronate | Carboxylic Acid | EX_fruur_e |
| Gallic acid | Alcohol | EX_ga_e |
| (R)-Glycerate | Carboxylic Acid | EX_glyc__R_e |
| Mannotriose (beta-1,4) | Carbohydrates/Derivatives | EX_mantr_e |
| GTP C10H12N5O14P3 | Nucleobases/Nucleosides/Nucleotides/Derivatives | EX_gtp_e |
| Ethanesulfonate C2H5O3S | Organic Sulfur | EX_eths_e |
