## Supplemental Table S3 for "Emergent Metabolic Niches for Marine Heterotrophs"

| Supplemental Table S3 | | | | | | | | | | | |
|---|---|---|---|---|---|---|---|---|---|---|---|
| Full Region Name | Region Identifier | Number of Stations | Oceanographic Category | 1 | 2 | 3 | 4 | 5 | 6 | 7 | 8 |
| Atlantic Ocean North | AON | 263 | Oligotrophic Open Ocean | 0.260364076 | 0.0349331189 | 0.161649979 | 0.1365709009 | 0.02481738114 | 0.04945085427 | 0.1255909017 | 0.2066227881 |
| Atlantic Ocean South | AOS | 117 | Oligotrophic Open Ocean | 0.2560422656 | 0.04364614741 | 0.1260065851 | 0.1414345622 | 0.06010333221 | 0.04128340442 | 0.1590016308 | 0.1724820722 |
| Baltic Sea | Baltic Sea | 55 | Coastal | 0.3264895027 | 0.03862604166 | 0.07295514286 | 0.2931048935 | 0.08146425573 | 0.04965877839 | 0.03661489623 | 0.1010864889 |
| Pelagic Baltic Sea | Baltic_Pelagic | 4 | Coastal | 0.2421887128 | 0.04636411151 | 0.03542523909 | 0.3239646337 | 0.1915216583 | 0.05826592593 | 0.04568347217 | 0.05658624654 |
| Black Sea | Black Sea | 5 | Coastal | 0.102164708 | 0.1045024354 | 0.0406608424 | 0.2402517605 | 0.05463539411 | 0.1386252567 | 0.2846484886 | 0.03451111441 |
| Chesapeake/Delaware Bay | CB | 36 | Estuarine | 0.1299348408 | 0.1427070879 | 0.06460987342 | 0.1999216267 | 0.1248936071 | 0.06323995123 | 0.1007246705 | 0.1739683424 |
| Chesapeake Bay | Chesapeake Bay | 15 | Estuarine | 0.1724961711 | 0.3985819176 | 0.01916731784 | 0.05382289659 | 0.08540220485 | 0.1993124604 | 0.007120240991 | 0.0640967906 |
| Columbia River | Columbia River | 4 | Estuarine | 0.2218225934 | 0.1936499244 | 0.01807052537 | 0.2835425615 | 0.1079347336 | 0.07465891859 | 0.05623520316 | 0.04408554006 |
| Gulf of Mexico | GOM | 7 | Coastal | 0.1182063714 | 0.02910215687 | 0.200741243 | 0.07998169773 | 0.1050340337 | 0.005717683904 | 0.0294296813 | 0.4317871322 |
| Station ALOHA | HOT | 299 | Oligotrophic Open Ocean | 0.1878806525 | 0.02282639957 | 0.1522887557 | 0.1596116893 | 0.1193697459 | 0.03692357894 | 0.1113352106 | 0.2097639675 |
| Indian Ocean North | ION | 9 | Oligotrophic Open Ocean | 0.2175502052 | 0.01917380234 | 0.106411588 | 0.1147630334 | 0.02729155058 | 0.01115583153 | 0.1562701367 | 0.3473838523 |
| Indian Ocean South | IOS | 12 | Oligotrophic Open Ocean | 0.2173185265 | 0.01690134505 | 0.1133546909 | 0.129895977 | 0.01794501066 | 0.02088969215 | 0.1966703767 | 0.287024381 |
| Mediterranean Sea | MED | 7 | Oligotrophic Seas | 0.386542599 | 0.0282757865 | 0.1131949239 | 0.1106030917 | 0.01623620146 | 0.0747713184 | 0.08497365299 | 0.185402426 |
| San Pedro Ocean Time-series | NPAC | 12 | Coastal | 0.2287294645 | 0.04868169073 | 0.01914442086 | 0.1421650938 | 0.1500951996 | 0.1777866626 | 0.1312195948 | 0.1021778732 |
| Pacific Ocean North | PON | 78 | Oligotrophic Open Ocean | 0.2335037556 | 0.03235575028 | 0.1667013573 | 0.09586769766 | 0.0209050433 | 0.05132718872 | 0.1507461121 | 0.248593095 |
| Pacific Ocean South | POS | 207 | Oligotrophic Open Ocean | 0.3343296372 | 0.02380094532 | 0.1233919427 | 0.07684942256 | 0.02569257387 | 0.05868610026 | 0.1062032291 | 0.251046149 |
| Pearl River | Pearl_river | 15 | Estuarine | 0.1422177718 | 0.1677922755 | 0.01846245669 | 0.1468304336 | 0.1183244855 | 0.05899260951 | 0.2054204745 | 0.141959493 |
| Red Sea | RED | 6 | Oligotrophic Seas | 0.1449686598 | 0.02031445109 | 0.1265289505 | 0.09973935276 | 0.02174299215 | 0.01888908478 | 0.1422482784 | 0.4255682304 |
| San Francisco Bay | SFBay | 8 | Estuarine | 0.2950774723 | 0.08208865936 | 0.06524683363 | 0.1526398205 | 0.1554155723 | 0.05593076443 | 0.05199932558 | 0.1416015519 |
| Saanich Inlet | SI | 4 | Coastal | 0.1527334592 | 0.01710858158 | 0.01666965497 | 0.2760001872 | 0.03513987358 | 0.333944066 | 0.155419953 | 0.01298422445 |
| Southern Ocean | SOC | 3 | Southern Ocean | 0.1095835443 | 0.02091484749 | 0.2958038819 | 0.06167178951 | 0.1228618317 | 0.2580164272 | 0.04720747022 | 0.08394020774 |
| Sapelo Island | Sapelo | 5 | Coastal | 0.1212185323 | 0.05057639249 | 0.03054432166 | 0.2198176077 | 0.1495611145 | 0.2425214243 | 0.09144833999 | 0.0943122671 |
| Yquina Bay | Yaquina Bay | 32 | Estuarine | 0.2446371889 | 0.1094614481 | 0.01474587385 | 0.1458108956 | 0.2045267358 | 0.02327114573 | 0.1395819538 | 0.1179647582 |
