## Supplemental Table S4 for "Emergent Metabolic Niches for Marine Heterotrophs"

| ***Supplemental Table S4*** | | | | | | | | | |
|---|---|---|---|---|---|---|---|---|---|
| Oceanographic Category | Number of Stations | Cluster 1 | Cluster 2 | Cluster 3 | Cluster 4 | Cluster 5 | Cluster 6 | Cluster 7 | Cluster 8 |
| Coastal | 92 | 0.2333759115 | 0.04388963309 | 0.070819109 | 0.2061396253 | 0.108399806 | 0.1047314634 | 0.08444503354 | 0.1481994182 |
| Estuarine | 110 | 0.2270870975 | 0.1272328312 | 0.02284734109 | 0.1533877098 | 0.1824165949 | 0.03895003324 | 0.1274704525 | 0.1206079398 |
| Oligotrophic Open Ocean | 985 | 0.2649355317 | 0.03022121037 | 0.1440054204 | 0.120178338 | 0.04317272676 | 0.04770935714 | 0.1267005569 | 0.2230768587 |
| Oligotrophic Seas | 13 | 0.2935536776 | 0.02531296426 | 0.1184094804 | 0.1065445174 | 0.01841808644 | 0.05262707659 | 0.1070790739 | 0.2780551234 |
| Southern Ocean | 3 | 0.1096757406 | 0.02059858057 | 0.2956828003 | 0.06128628782 | 0.123368813 | 0.2590647262 | 0.04688827452 | 0.08343477704 |
