## Supplemental Table S5 for "Emergent Metabolic Niches for Marine Heterotrophs"

| ***Supplemental Table S5*** | |
|---|---|
| Cluster Comparison | p value |
| 2-1 | 3.58E-08 |
| 3-1 | 0.954627773 |
| 4-1 | 0.7981970848 |
| 5-1 | 0.00255195585 |
| 6-1 | 0.9752095145 |
| 7-1 | 0.7596861371 |
| 8-1 | 0.9999507727 |
| 3-2 | 3.76E-09 |
| 4-2 | 1.49E-05 |
| 5-2 | 0.2200741497 |
| 6-2 | 8.68E-05 |
| 7-2 | 2.39E-09 |
| 8-2 | 1.38E-05 |
| 4-3 | 0.1893939319 |
| 5-3 | 1.21E-04 |
| 6-3 | 0.4932582833 |
| 7-3 | 0.9996558229 |
| 8-3 | 0.9988789317 |
| 5-4 | 0.1662492488 |
| 6-4 | 0.9999237676 |
| 7-4 | 0.0708115924 |
| 8-4 | 0.7433751924 |
| 6-5 | 0.1405184432 |
| 7-5 | 3.80E-05 |
| 8-5 | 0.0129446364 |
| 7-6 | 0.2429095454 |
| 8-6 | 0.9333372143 |
| 8-7 | 0.9675410519 |
