## Supplemental Table S6 for "Emergent Metabolic Niches for Marine Heterotrophs"

| Compared Cluster | Compared Cluster | All | SAR86 | Rhodobacterales | Sphingomonadales | Pseudomonadales | Pelagibacterales | PCC-6307 | Other | Opitutales | Marinisomatales | Flavobacteriales | Enterobacterales_ | Cytophagales | Caulobacterales | Burkholderiales | AEGEAN-169 | Acidimicrobiales |
|---|---|---|---|---|---|---|---|---|---|---|---|---|---|---|---|---|---|---|
| 1 | 2 | 0.7989463806 | 0.419934988 | 0.6798227429 | 0.2306112051 | 0.6746929884 | 0.4773527086 | NA | 0.7957854867 | 0.2828065455 | 0.5017846823 | 0.8194350004 | 0.5985862613 | 0.6349437833 | 0.3565019071 | 0.5099224448 | NA | 0.310235858 |
| 1 | 3 | 0.8449069262 | 0.2632276416 | 0.5056242347 | NA | 0.5002261996 | 0.4709132612 | NA | 0.7938459516 | NA | 0.6293544173 | 0.8369632959 | 0.4351541698 | 0.5563268661 | NA | 0.2931603789 | NA | 0.2074897885 |
| 1 | 4 | 0.8374670148 | 0.4297683537 | 0.5563185811 | 0.1714592874 | 0.5421856642 | 0.5026069283 | 0.1509902179 | 0.7996332645 | 0.3514454067 | 0.5440512896 | 0.7559486032 | 0.483912468 | 0.3959026933 | 0.2814624012 | 0.3617681563 | NA | 0.2570967376 |
| 1 | 5 | 0.8273881078 | 0.419586122 | 0.64904809 | 0.3811065555 | 0.6712320447 | 0.548007071 | 0.2030960023 | 0.7707895637 | 0.373598963 | 0.6052199602 | 0.7722569704 | 0.4256902337 | 0.5653985143 | 0.1184490025 | 0.1648871452 | NA | 0.2831065655 |
| 1 | 6 | 0.8072746396 | 0.4530544281 | 0.4921690524 | 0.2279495746 | 0.5759056211 | 0.6362354159 | 0.0788583383 | 0.7761890888 | 0.3719415665 | 0.6281718016 | 0.7740310431 | 0.4510219097 | 0.5138685703 | 0.3167761564 | 0.3996652961 | NA | 0.1034685001 |
| 1 | 7 | 0.8284811378 | 0.2717142999 | 0.5636680126 | 0.1675379425 | 0.6187599301 | 0.4591261744 | NA | 0.8476480842 | 0.3926586807 | 0.5819787979 | 0.7716632485 | 0.3981662095 | 0.5582184196 | 0.1752095222 | 0.2906773984 | NA | 0.3186494112 |
| 1 | 8 | 0.8025676012 | NA | 0.5519781709 | NA | 0.5284642577 | 0.3886255026 | NA | 0.763360858 | 0.3297410309 | 0.5531373024 | 0.7541276217 | 0.4212360978 | 0.519285202 | 0.2866186798 | 0.2744484246 | NA | 0.24097009 |
| 2 | 3 | 0.7753917575 | 0.5118632317 | 0.718740046 | NA | 0.692592144 | 0.4707812071 | NA | 0.8123838305 | NA | 0.5401253104 | 0.8803586364 | 0.6586849689 | 0.6478453875 | NA | 0.4859125614 | 0.3869579434 | 0.2948750257 |
| 2 | 4 | 0.7971437573 | 0.4545585513 | 0.6796560884 | 0.3406110704 | 0.6083652377 | 0.3602496386 | NA | 0.78514117 | 0.3217269182 | 0.6299499273 | 0.7540445328 | 0.5726030469 | 0.6462606788 | 0.2746297717 | 0.5293958783 | 0.285030961 | 0.4240909219 |
| 2 | 5 | 0.7815839052 | 0.1901007742 | 0.6268313527 | 0.4071877599 | 0.6257848144 | 0.3350928724 | NA | 0.7683380246 | 0.2839986086 | 0.6067393422 | 0.794308424 | 0.5156766772 | 0.688791573 | 0.3808265924 | 0.5517795086 | NA | 0.3216001987 |
| 2 | 6 | 0.7672163844 | 0.3944503963 | 0.6928328276 | 0.3283340633 | 0.7147321701 | 0.4453222156 | NA | 0.8035640121 | 0.3802590966 | 0.4874662161 | 0.7490028739 | 0.6060413718 | 0.7190599442 | 0.3523624241 | 0.4648796618 | NA | 0.3358679712 |
| 2 | 7 | 0.8182488084 | 0.3697157502 | 0.7279790044 | 0.2679120898 | 0.7345622182 | 0.3430925608 | NA | 0.8466829062 | 0.3454710543 | 0.5160543323 | 0.8731450438 | 0.6501075625 | 0.6690906286 | 0.3182494044 | 0.4724879861 | NA | 0.3538352847 |
| 2 | 8 | 0.7759789824 | NA | 0.7274648547 | NA | 0.7628696561 | 0.3653849363 | NA | 0.8099697828 | 0.3348436356 | 0.541426301 | 0.8164143562 | 0.6322367191 | 0.6822730899 | 0.4128487408 | 0.5853829384 | 0.2825010419 | 0.3249714673 |
| 3 | 4 | 0.7699774504 | 0.3647487462 | 0.4826725125 | NA | 0.559281826 | 0.4451363087 | NA | 0.7990082502 | NA | 0.5311200619 | 0.827902019 | 0.474285692 | 0.5500840545 | NA | 0.3372276723 | 0.2134583592 | 0.3346907496 |
| 3 | 5 | 0.8005220294 | 0.51161623 | 0.6561794281 | NA | 0.6519393921 | 0.5029203892 | NA | 0.8309012651 | NA | 0.4230251014 | 0.8229458332 | 0.5423144102 | 0.4840340912 | NA | 0.2885446846 | NA | 0.3482176065 |
| 3 | 6 | 0.8038511276 | 0.3742012978 | 0.3985678554 | NA | 0.4870015383 | 0.5161299109 | NA | 0.8339858651 | NA | 0.3026397824 | 0.8256617785 | 0.3600393236 | 0.5466056466 | NA | 0.4107034504 | NA | 0.2127988338 |
| 3 | 7 | 0.8328499794 | 0.3699981868 | 0.4836077392 | NA | 0.6244325638 | 0.3768307567 | NA | 0.8413235545 | NA | 0.4188953042 | 0.7026120424 | 0.3167015314 | 0.5069461465 | NA | 0.3133125603 | NA | 0.3857189715 |
| 3 | 8 | 0.839173913 | NA | 0.5118207932 | NA | 0.52800107 | 0.3419329524 | NA | 0.8123958707 | NA | 0.3426534235 | 0.8141863942 | 0.3551894426 | 0.3811972141 | NA | 0.3702973425 | 0.1975741237 | 0.1271596998 |
| 4 | 5 | 0.8054401875 | 0.4542334676 | 0.6252754927 | 0.4417518079 | 0.6552917957 | 0.3646194935 | 0.2268708199 | 0.820358932 | 0.4148525 | 0.5170619488 | 0.7453194857 | 0.5087736845 | 0.5981732011 | 0.3364825249 | 0.3580073714 | NA | 0.3911091983 |
| 4 | 6 | 0.776378572 | 0.2443776429 | 0.4720506072 | 0.2079518586 | 0.5397134423 | 0.3867835999 | 0.208245337 | 0.7929137349 | 0.3966782391 | 0.5534476042 | 0.7639408708 | 0.3740318418 | 0.6334023476 | 0.4001167119 | 0.4124534428 | NA | 0.1862937212 |
| 4 | 7 | 0.8470900655 | 0.3361594081 | 0.5812653899 | 0.1953549534 | 0.6836054325 | 0.3148459792 | NA | 0.8271434307 | 0.4170547426 | 0.4398395121 | 0.8407593966 | 0.4019573629 | 0.5076542497 | 0.2744829059 | 0.433978349 | NA | 0.4292904735 |
| 4 | 8 | 0.8296276331 | NA | 0.5719254017 | NA | 0.622013092 | 0.3130659759 | NA | 0.8003775477 | 0.3559266925 | 0.4509515464 | 0.8212280869 | 0.3798933625 | 0.5791752934 | 0.2696650028 | 0.4253309071 | 0.07236015052 | 0.3674325049 |
| 5 | 6 | 0.7731743455 | 0.3940497041 | 0.6180651188 | 0.4210611582 | 0.6617231965 | 0.4143125415 | 0.2110337168 | 0.804462254 | 0.4540700912 | 0.4295547009 | 0.7626585364 | 0.4837541878 | 0.5701366663 | 0.3473000526 | 0.3962272704 | NA | 0.2875641882 |
| 5 | 7 | 0.8213275671 | 0.3692816198 | 0.6191690564 | 0.4619380236 | 0.6891596317 | 0.3564739823 | NA | 0.8496707082 | 0.3480095863 | 0.428401947 | 0.8399067521 | 0.5071648359 | 0.4892756343 | 0.2157658041 | 0.2857961357 | NA | 0.1947282702 |
| 5 | 8 | 0.8014882207 | NA | 0.6954833865 | NA | 0.7194139957 | 0.403449297 | NA | 0.8232325315 | 0.4164543152 | 0.3491246104 | 0.8422886729 | 0.4872995913 | 0.483532697 | 0.3198311627 | 0.1454428583 | NA | 0.3531067669 |
| 6 | 7 | 0.8263292313 | 0.3584939241 | 0.4971340895 | 0.2004271299 | 0.6191598773 | 0.4637551904 | NA | 0.8138129115 | 0.4924795628 | 0.4380289316 | 0.7984529734 | 0.3613923192 | 0.5631402135 | 0.2550594807 | 0.2955740392 | NA | 0.323277235 |
| 6 | 8 | 0.8025747538 | NA | 0.525490284 | NA | 0.5228598714 | 0.5041097403 | NA | 0.7894396186 | 0.2447075546 | 0.3265831172 | 0.7551851273 | 0.3016343117 | 0.4747754633 | 0.2349104434 | 0.4579447508 | NA | 0.246285364 |
| 7 | 8 | 0.8017535806 | NA | 0.5695308447 | NA | 0.4938226938 | 0.2609551847 | NA | 0.8021425009 | 0.4586596489 | 0.367495209 | 0.7518382072 | 0.2851961553 | 0.4861619174 | 0.2190639824 | 0.3717501163 | NA | 0.3168465495 |
